# Phylogenetic Analysis of Beta-Lactamases Reveals Distinct Evolutionary Patterns of Chromosomal and Plasmid-Encoded BLs and the Mosaic Role of VIM Linking NDM and IMP

**DOI:** 10.1101/2025.10.08.681124

**Authors:** Ananya Anurag Anand, Vidushi Yadav, K Kshitiz, PC Sreedevi, Sarfraz Anwar, Sintu Kumar Samanta

## Abstract

Beta-lactamases (BLs) are a major driver of antibiotic resistance in pathogens like *Acinetobacter baumannii*, Enterobacteriaceae and *Pseudomonas aeruginosa*. This study explores the structural stability and functional divergence of metallo-beta-lactamases (MBLs) and AmpC through study of protein-protein interaction (PPI) network analysis, phylogenetics and identification of conserved domains and motifs. The study also involves analysis of co-evolutionary dynamics of BLs with related genes. PPI analysis identified NagZ as central interacting partner for BLs in *P. aeruginosa* and Enterobacteriaceae, suggesting its pivotal role in BL expression, and highlighting its probable role as a potential drug target. In contrast, *A. baumannii* exhibited such a high diversity in protein interactions, that considering a single protein as topmost interacting partner was difficult. Phylogenetic analysis revealed strong co-evolutionary trends between functionally-related proteins. *A. baumannii* and Enterobacteriaceae were found to be closely related in case of not just the chromosomally-encoded MBL-fold containing proteins, but also the true MBLs which are plasmid-encoded. On the other hand, similar evolutionary patterns for chromosomally-encoded BLs like AmpC and related genes was found in *A. baumannii* and *P. aeruginosa*. Analysis of PFAM domains showed that catalytic and substrate binding domains are more conserved than accessory ones. Several motifs, such as, ASNGLI were found to be conserved across all MBLs and thus carry the potential to be significant drug targets. Finally, inter-BL analysis revealed that VIM acts as an evolutionary link by acting like a mosaic between IMP and NDM as supported by intermediate GC content, positioning in phylogenetic trees and shared sequence features.

## 1. Introduction

The emergence and spread of antibiotic resistance represent one of the most pressing challenges to modern medicine, with beta-lactamases (BLs) playing a central role (**Bush and Bradford, 2016; Amod et al., 2025**). BLs are enzymes that hydrolyze the beta-lactam ring of the beta-lactam antibiotics such as penicillin, cephalosporin and carbapenem, rendering them ineffective and severely limiting treatment options for multidrug-resistant (MDR) bacterial infection **(Arer and Anurag Anand et al., 2024)**. World Health Organization (WHO), in its 2017 priority pathogen list, identified carbapenem-resistant *Acinetobacter baumannii*, Enterobacteriaceae and *Pseudomonas aeruginosa*, as critical threats due to their ability to acquire and disseminate BL genes through both vertical inheritance and horizontal gene transfer (HGT) (https://www.who.int/publications/i/item/9789240093461). We, thus, aimed to study the evolution and transfer patterns of BLs in and across these pathogens.

BLs are broadly classified into four molecular classes (A, B, C, and D) based on their amino acid sequences and catalytic mechanisms **(Anurag Anand et al., 2024; Arer and Kar, 2022)**. Class A, C, and D BLs are serine-based enzymes that utilize a serine residue at their active site to hydrolyze beta-lactams. In contrast, Class B BLs, also known as metallo-beta-lactamases (MBLs), are zinc-dependent and require divalent metal ions for activity **(Bush and Jacoby, 2010)**. MBLs, such as New Delhi metallo-beta-lactamase (NDM), Verona integron-encoded metallo-beta-lactamase (VIM), and imipenemase (IMP), are particularly concerning due to their plasmid borne-nature facilitating high HGT rates, broad substrate specificity and resistance to nearly all beta-lactams, including carbapenems **(Anurag Anand et al., 2025a)**. AmpC-type BLs, which are majorly chromosomally encoded, confer resistance to cephalosporins and penicillins and are well known for their inducibility in response to antibiotic exposure **(Liu et al., 2022)**. Plasmid-encoded MBLs of several types have been largely identified in *P. aeruginosa*, *A. baumannii*, and Enterobacteriaceae, indicating extensive HGT between environmental and clinical bacterial reservoirs. In contrast, chromosomally encoded BLs like AmpC are more conserved within specific bacterial lineages, reflecting host-specific evolutionary pressures **(Imkamp et al., 2022; Schotte et al., 2024)**. The ability of pathogens to harbor both chromosomal and plasmid-encoded BLs further complicates treatment and increases the likelihood of multidrug resistance. Thus, it becomes important for us to identify conserved evolutionary patterns in order to target multiple BLs.

Recently, phylogenetic studies have been showing great success in analysis of several enzymes and their associated proteins by providing insights into the evolutionary pressures shaping these interactions and revealing conserved and divergent motifs that may serve as drug targets **(Salahuddin and Khan, 2014)**. Therefore, this study aims to address the growing challenge of BL-mediated antibiotic resistance by studying evolutionary patterns, and identifying conserved and divergent motifs in BLs **(Anurag Anand et al., 2025b)**. Through study of protein-protein interaction (PPI) network analysis, multiple sequence alignment (MSA), phylogenetic analysis and motif identification, we uncover co-evolutionary relationships and functional interdependence between BLs and associated proteins. We also evaluate the evolutionary clustering of beta-lactamases based on amino acid composition (AAC), atomic and bond composition (ABC), and physicochemical properties (APP), providing insights into the structural and functional mechanisms driving resistance **(Brandt and Barrangou, 2016)**.

Our findings reveal that *A. baumannii* exhibits a higher diversity of interacting proteins, reflecting a more complex and adaptable resistance strategy. This increased diversity is likely driven by its high capacity for HGT and its ability to integrate foreign genetic material, thereby expanding its repertoire of beta-lactamase-associated proteins **(Mei et al., 2019)**. In contrast, *P. aeruginosa* and Enterobacteriaceae demonstrate more conserved interaction patterns, indicating a streamlined and specialized approach to resistance. This study identifies NagZ as the topmost interacting partner for beta-lactamases in *P. aeruginosa* and Enterobacteriaceae. NagZ plays a role in the regulation of chromosomally-encoded BL (AmpC) expression. This BL expression is regulated via a complex network of interacting proteins such as AmpR, AmpD, and NagZ. To be more specific, NagZ serves a role in AmpC expression through the AmpR-AmpD regulatory cascade and is also a central player in the peptidoglycan recycling pathway. Most importantly, our study suggests a strong co-evolutionary link between NagZ, AmpC, and AmpR indicating that these proteins have undergone coordinated genetic changes to enhance resistance efficiency under antibiotic pressure **(Bedhomme et al., 2017)**. Further, we found that chromosomally-encoded MBL-fold containing proteins and plasmid-encoded MBLs show similar evolutionary patterns for *A. baumannii* and Enterobacteriaceae, whereas chromosomally-encoded BLs like AmpC and related show similar evolutionary patterns across *A. baumannii* and *P. aeruginosa*. Comparative analysis of AAC, ABC, and APP clusters using Principal Component Analysis (PCA) reveals distinct clustering patterns among NDM, VIM, IMP, and AmpC BLs. NDM and VIM form a cohesive cluster due to conserved hydrophobic and charged residues, reflecting their evolutionary stability and broad substrate specificity. In contrast, IMP shows greater structural divergence, indicating functional differentiation and adaptation to diverse environmental niches. GC content analysis further highlights differences in structural stability, with NDM exhibiting high GC content and lower mutational plasticity, while IMP displaying low GC content and higher evolutionary variability **(Brandt and Barrangou, 2016)**. VIM was found to lie between NDM and IMP in GC content analysis. Motif analysis using MEME and TOMTOM identifies conserved motifs and mutational hotspots across different beta-lactamase classes **(Nystrom and McKay, 2021)**. Highly conserved motifs in NDM and VIM suggest a strong evolutionary link, while the partial conservation of motifs in IMP reflects a mix of conserved and functionally distinct regions. Mutational hotspots affecting Zn² coordination and enzyme stability are identified, highlighting key structural vulnerabilities that could serve as drug targets. Lastly, inter-BL phylogenetic analysis combined with study of GC content, conserved domains and motifs reveals the mosaic role of VIM linking IMP and NDM, due to shared sequence physiochemical properties, motifs and functional domains from both the MBLs.

## 2. Methodology

### 2.1 Protein-protein interaction study using STRING

STRING database was utilized to search for interacting partners of BLs and related proteins **(Szklarczyk et al, 2025)**. By related proteins, we mean proteins which have domains similar to those lying within BLs. The interacting partners of BLs and related proteins were identified using STRING. Further, FASTA sequences of the same were obtained for further analysis. The keyword used for protein search was ‘beta-lactamases’ along with organism-specific search filters: ‘*Pseudomonas aeruginosa* group’, ‘*Acinetobacter baumannii’* and ‘Enterobacteriaceae’ for *P. aeruginosa*, *A. baumannii* and Enterobacteriaceae respectively. Topmost interacting partners found associated with most BLs and related proteins were identified. Those proteins which were found to interact with at least 5 BLs and related proteins were selected for studying their co-evolutionary dynamics with BLs and related proteins.

### 2.2 Selection of beta-lactamase sequences for BLAST

The sequence of NDM-1 was sourced from high quality PDB structure with PDB ID 8B1W, that can be accessed at https://doi.org/10.2210/pdb8b1w/pdb **(Bersani et al., 2023)**. The sequences chosen for sequence analysis of VIM-2, and IMP-1 were obtained from NCBI accession ACY78406.1 and CRP48843.1, respectively. Subsequently, homologous sequences from NCBI were obtained using the NCBI BLASTP facility. All the initial sequences were from the bacteria *P. aeruginosa* and were considered as the starting point.

### 2.3 Sequence similarity search of BLs, related proteins and their interacting partners

BLASTP was performed against the non-redundant (nr) database using NCBI BLAST facility **(Camacho et al., 2009)**. For each protein, the initial number of sequences obtained using BLASTP were 5000. These were then subjected to an algorithm that could identify the unique organisms and select one representative sequence (the topmost hit) for each organism or species. For VIM, NDM and IMP, top 100 sequences were studied (multiple sequences of the same species were also retained). Since the query sequence was from *P. aeruginosa,* therefore, as expected, the topmost sequences were from *Pseudomonas* itself. Thus, the sequences of *A. baumannii* and Enetrobacteriaceae were added to the list when not found in top hits for any of the proteins. This was done by performing BLASTP with organism-specific search. The keywords or filters used while performing organism-specific search were Taxid: 562 (*Escherichia coli*), Taxid: 91347 (Enterobacteriaceae) and Taxid: 470 (*A. baumannii)*. Later, BLASTP was also performed against the clustered nr database, to study the representatives of different MBL clusters.

### 2.4 Multiple Sequence Alignment and Mutational Analysis

Multiple sequence alignment (MSA) for each of the protein datasets was performed. MEGA software (version 11) was used for the alignment, with the Muscle algorithm as the alignment method **(Edgar et al., 2004; Kumar et al., 2008; Tamura et al., 2021)**. The gap open penalty was set to -2.90, while the gap extension penalty was set at 0.00. To account for sequence hydrophobicity, a multiplier of 1.20 was applied. The algorithm was run for a maximum of 16 iterations, ensuring sufficient refinement in the alignment. UPGMA (Unweighted Pair Group Method with Arithmetic Mean) was used to guide the clustering of sequences **(Sokal and Michener, 1958).** This method continued to be applied in subsequent iterations, maintaining consistency in alignment structure. To enhance alignment precision, the minimum diagonal length (Lambda) was set to 24. This systematic approach with MEGA and the Muscle algorithm allowed us to produce multiple sequence alignments, which were subsequently visualized using itol (itol can be accessed at https://itol.embl.de/) (used for construction of phylogenetic tree using newick data extracted from MEGA analysis). These were further analyzed in the Jalview software (version 2.11.4.0) **(Waterhouse et al., 2009)**. Through Jalview, we identified conserved domains present across all species in the study. Additionally, it enabled us to detect specific amino acid mutations occurring within these conserved domains. While studying the alignment of protein sequences using Jalview, color-coding proved useful in refining the alignment.

Sequence logos were then generated using the WebLogo software to visualize residue conservation across all aligned blocks in our MSA **(Crooks et al., 2004)**. These logos illustrate highly conserved domains, with minimal mutations observed within the conserved region.

### 2.5 Phylogenetic analysis

Maximum likelihood (ML) tree was built to perform the phylogenetic analysis and study the evolutionary dynamics of the proteins **(Felsenstein, 1981)**. Our aim was to construct a phylogenetic tree using bootstrap sampling. We first prepared and aligned the sequences of interest. Next, we selected the best-fit substitution model using tools like MEGA, which informs the likelihood calculations. An initial ML tree was then constructed based on the full dataset, evaluating various topologies to maximize likelihood. We performed bootstrap analysis by resampling the original alignment 100 times to create pseudo-replicate datasets, reconstructing an ML tree for each replicate to assess clade consistency **(Russo et al., 2018)**. Bootstrap support values were calculated for each node in the original tree, indicating the reliability of clades based on the proportion of supporting replicates. It is to note that standard bootstrap sampling was performed instead of adaptive. The standard method is slow but extremely accurate. The combination of ML methods and bootstrap sampling provides a statistically robust framework for accurately estimating evolutionary relationships while measuring uncertainty, resulting in phylogenetic trees that reflect evolutionary relationships with greater confidence. Where necessary, we reconstructed the tree using other algorithms as well, namely, NJ (neighbor joining) and ME (minimum evolution), MP (Maximum Parsimony), UPGMA (Unweight Pair Group Method using Arithmetic mean), alongside ML (maximum likelihood). Thus, instead of relying on a single evolutionary tree, insights were derived from different algorithms to enable efficiency and reliability of results.

### 2.6 Study of co-evolutionary dynamics

We assessed as to which interacting partners show similar evolutionary pattern with their respective BLs and related proteins. This was done by visually identifying how the genus and species divergence is occurring in the trees.

### 2.7 Study of closely and distantly related priority pathogens

We also assessed the phylogenetic relationships among priority pathogens for each protein in such a way so as to identify which two priority pathogens lie closer together and which one lays farther apart.

### 2.8 GC content analysis

GC content analysis was performed using Biopython for all sequences of a BL. This was done in order to account for stability of different BL types, and to compare the changes in nucleotide composition over the period of time within the group of a particular beta-lactamase.

### 2.9 Statistical analysis for protein features and physiochemical properties

The amino acid composition (AAC), atoms and bonds composition (ABC) and various advanced physiochemical properties (APP/PCP) of the proteins were studied using the Pfeature software. Pfeature is a renowned web server for computing various features of proteins from their amino acid sequence **(Pande et al., 2023)**. The values for AAC, ABC and APP descriptors for each of the proteins were subjected to PCA to cluster the sequences into different groups on the basis of their overall similarity. The contribution of each amino acid to AAC, ABC and APP was then calculated.

### 2.10 PFAM domain analysis

Protein family (PFAM) domains with different structural and functional roles were identified and studied across different BLs **(Finn et al., 2016)**. Comparisons were made both within (between different sequences of a particular BL type) and across BLs (between different MBLs, and between MBLs and AmpC). This was done to get an idea of how the protein structures of beta-lactamases have evolved over the period of time.

Further, PFAM domain conservation scores were calculated based on the number of bacterial strains containing each domain. Percentages were assigned based on conservation levels across different bacterial groups. Heatmaps were built using matrix representation of these conservation values, for visualization. Heatmaps were generated using seaborn.heatmap().

The conservation percentage of PFAMs was calculated as follows:

**Conservation Score = (Number of strains with the domain/Total strains in the group) × 100**

Example: If 90% of Pseudomonas strains contains the Beta-lactamase domain of AmpC, then the conservation score for AmpC-Pseudomonas will be 90.

### 2.11 Motif identification and comparison

Repeatedly occurring motifs were identified using the MEME software. TOMTOM comparative analysis was then performed to see how these differ across different BLs **(Nystrom and McKay, 2021)**. Comparisons were made both within and across different BLs (between different MBLs, and between MBLs and AmpC). This was done to find out how beta-lactamases have evolved over the period of time.

### 2.12 Defining Similarity Scores based on motifs analysis

We used approximate similarity values based on the p-values and motif conservation analysis from TOMTOM. The similarity score was derived from the motif alignment p-values in the TOMTOM results. A lower p-value means stronger similarity between motifs, while a higher p-value suggests weaker similarity.

We used the following formula to convert p-values into similarity scores:

**Similarity Score = 100 - (log10 (p-value) × 10)**

The above formula ensures that highly conserved motifs (lower p-values) result in higher similarity scores, whereas weakly conserved motifs (higher p-values) result in lower similarity scores. This approach provides normalized similarity scores. Scores close to 100 indicate high conservation, whereas lower scores (closer to 50-60) indicate moderate or weak conservation.

## 3. Results and discussion

### 3.1 NagZ as the topmost interacting partner for BL and related proteins

For *P. aeruginosa,* 87 entries of BLs and related proteins were found for *P. aeruginosa* on STRING database. String interactions for all the proteins were extracted in the form of nodes. Redundancy was removed to make sure that each node is taken into account only once for evaluation. A combined list containing STRING IDs of the interacting partners of all BLs and related proteins was formulated and was further annotated with the names of the BLs and related proteins with which they interact.

For *A. baumannii,* 18 entries of BLs and related proteins were found, whereas 350 entries of BLs and related proteins were found for Enterobacteriacea.

For *P. aeruginosa,* NagZ was found to be the topmost interacting protein, showing interactions with 19 BLs and related proteins. Gpmi, polA, and rpoB were found to interact with 13 proteins each. RpoZ was found to interact with 12 whereas guaA was found to interact with 11 such proteins. The topmost interacting partners and the associated BLs and related proteins for *P. aeruginosa* have been mentioned in **Table 1**.

**Table 1.**
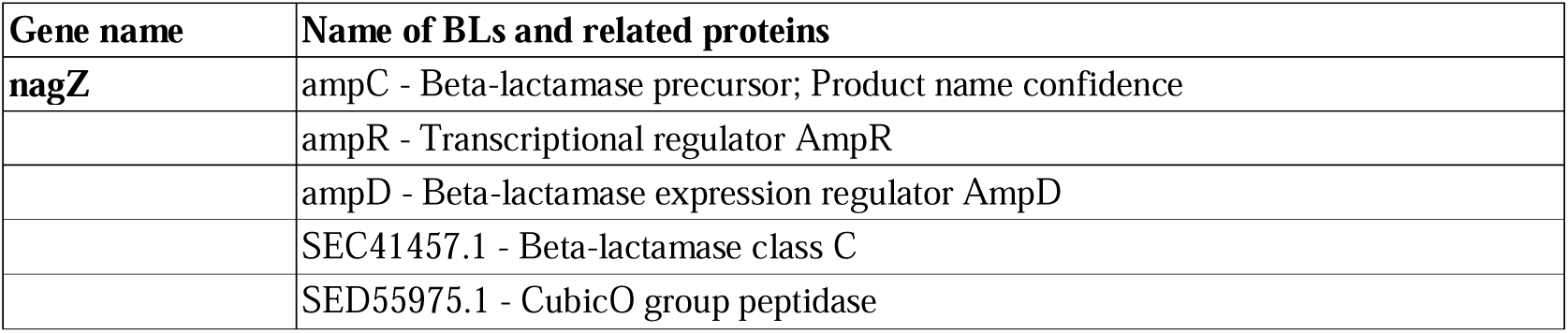
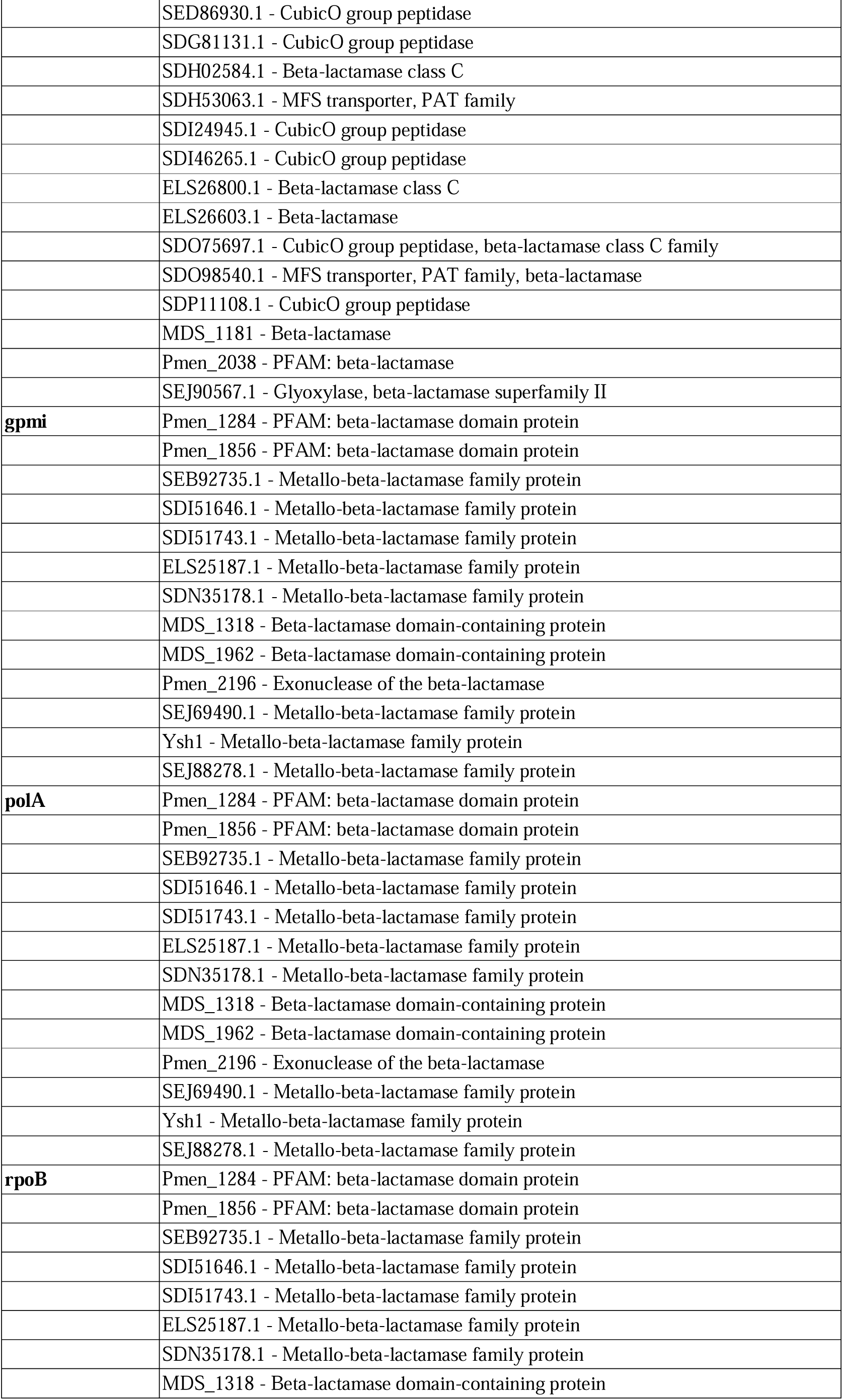
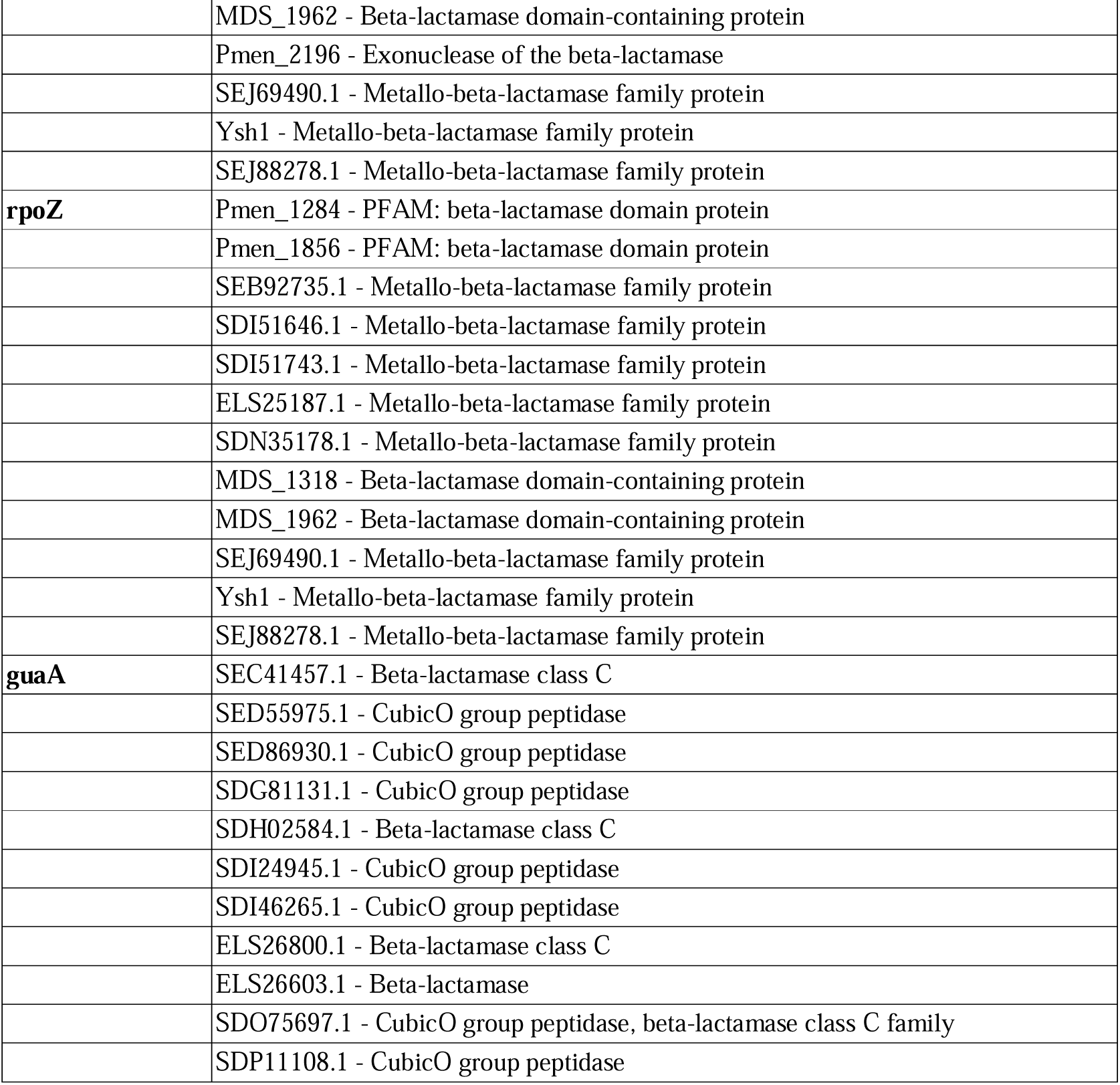
Topmost interacting partners and the associated BLs and related proteins for *P. aeruginosa*.

| Gene name | Name of BLs and related proteins |
| --- | --- |
| nagZ | ampC - Beta-lactamase precursor; Product name confidence |
|  | ampR - Transcriptional regulator AmpR |
|  | ampD - Beta-lactamase expression regulator AmpD |
|  | SEC41457.1 - Beta-lactamase class C |
|  | SED55975.1 - CubicO group peptidase |
|  | SED86930.1 - CubicO group peptidase |
|  | SDG81131.1 - CubicO group peptidase |
|  | SDH02584.1 - Beta-lactamase class C |
|  | SDH53063.1 - MFS transporter, PAT family |
|  | SDI24945.1 - CubicO group peptidase |
|  | SDI46265.1 - CubicO group peptidase |
|  | ELS26800.1 - Beta-lactamase class C |
|  | ELS26603.1 - Beta-lactamase |
|  | SDO75697.1 - CubicO group peptidase, beta-lactamase class C family |
|  | SDO98540.1 - MFS transporter, PAT family, beta-lactamase |
|  | SDP11108.1 - CubicO group peptidase |
|  | MDS_1181 - Beta-lactamase |
|  | Pmen_2038 - PFAM: beta-lactamase |
|  | SEJ90567.1 - Glyoxylase, beta-lactamase superfamily II |
| **gpmi** | Pmen_1284 - PFAM: beta-lactamase domain protein |
|  | Pmen_1856 - PFAM: beta-lactamase domain protein |
|  | SEB92735.1 - Metallo-beta-lactamase family protein |
|  | SDI51646.1 - Metallo-beta-lactamase family protein |
|  | SDI51743.1 - Metallo-beta-lactamase family protein |
|  | ELS25187.1 - Metallo-beta-lactamase family protein |
|  | SDN35178.1 - Metallo-beta-lactamase family protein |
|  | MDS_1318 - Beta-lactamase domain-containing protein |
|  | MDS_1962 - Beta-lactamase domain-containing protein |
|  | Pmen_2196 - Exonuclease of the beta-lactamase |
|  | SEJ69490.1 - Metallo-beta-lactamase family protein |
|  | Ysh1 - Metallo-beta-lactamase family protein |
|  | SEJ88278.1 - Metallo-beta-lactamase family protein |
| **polA** | Pmen_1284 - PFAM: beta-lactamase domain protein |
|  | Pmen_1856 - PFAM: beta-lactamase domain protein |
|  | SEB92735.1 - Metallo-beta-lactamase family protein |
|  | SDI51646.1 - Metallo-beta-lactamase family protein |
|  | SDI51743.1 - Metallo-beta-lactamase family protein |
|  | ELS25187.1 - Metallo-beta-lactamase family protein |
|  | SDN35178.1 - Metallo-beta-lactamase family protein |
|  | MDS_1318 - Beta-lactamase domain-containing protein |
|  | MDS_1962 - Beta-lactamase domain-containing protein |
|  | Pmen_2196 - Exonuclease of the beta-lactamase |
|  | SEJ69490.1 - Metallo-beta-lactamase family protein |
|  | Ysh1 - Metallo-beta-lactamase family protein |
|  | SEJ88278.1 - Metallo-beta-lactamase family protein |
| **rpoB** | Pmen_1284 - PFAM: beta-lactamase domain protein |
|  | Pmen_1856 - PFAM: beta-lactamase domain protein |
|  | SEB92735.1 - Metallo-beta-lactamase family protein |
|  | SDI51646.1 - Metallo-beta-lactamase family protein |
|  | SDI51743.1 - Metallo-beta-lactamase family protein |
|  | ELS25187.1 - Metallo-beta-lactamase family protein |
|  | SDN35178.1 - Metallo-beta-lactamase family protein |
|  | MDS_1318 - Beta-lactamase domain-containing protein |
|  | MDS_1962 - Beta-lactamase domain-containing protein |
|  | Pmen_2196 - Exonuclease of the beta-lactamase |
|  | SEJ69490.1 - Metallo-beta-lactamase family protein |
|  | Ysh1 - Metallo-beta-lactamase family protein |
|  | SEJ88278.1 - Metallo-beta-lactamase family protein |
| **rpoZ** | Pmen_1284 - PFAM: beta-lactamase domain protein |
|  | Pmen_1856 - PFAM: beta-lactamase domain protein |
|  | SEB92735.1 - Metallo-beta-lactamase family protein |
|  | SDI51646.1 - Metallo-beta-lactamase family protein |
|  | SDI51743.1 - Metallo-beta-lactamase family protein |
|  | ELS25187.1 - Metallo-beta-lactamase family protein |
|  | SDN35178.1 - Metallo-beta-lactamase family protein |
|  | MDS_1318 - Beta-lactamase domain-containing protein |
|  | MDS_1962 - Beta-lactamase domain-containing protein |
|  | SEJ69490.1 - Metallo-beta-lactamase family protein |
|  | Ysh1 - Metallo-beta-lactamase family protein |
|  | SEJ88278.1 - Metallo-beta-lactamase family protein |
| **guaA** | SEC41457.1 - Beta-lactamase class C |
|  | SED55975.1 - CubicO group peptidase |
|  | SED86930.1 - CubicO group peptidase |
|  | SDG81131.1 - CubicO group peptidase |
|  | SDH02584.1 - Beta-lactamase class C |
|  | SDI24945.1 - CubicO group peptidase |
|  | SDI46265.1 - CubicO group peptidase |
|  | ELS26800.1 - Beta-lactamase class C |
|  | ELS26603.1 - Beta-lactamase |
|  | SDO75697.1 - CubicO group peptidase, beta-lactamase class C family |
|  | SDP11108.1 - CubicO group peptidase |

For *A. baumannii,* no such top contender was concluded as the topmost interacting partner, because the maximum number of BLs and related proteins with which any protein was interacting was found to be 2. This indicated high variation in diversity or a large variety of proteins present in *A. baumannii*, such that there is minimal similarity of interactions across the BLs.

For Enterobacteriaceae, NagZ was again found to be the topmost interacting protein, showing interactions with 42 BLs and related proteins, followed by guaA, ybeY, ampD and BssS.

Overall, nagZ has the ability to act as a potential drug target in order to combat drug-resistance in *P. aeruginosa* and Enterbacteriaceae, due to its higher interaction ability with BLs and related proteins.

The details of all the interacting partners and the associated BLs and related proteins for *P. aeruginosa, A. baumannii* and Enterobacteriaceae can be found in the **Supplementary Files 1, 2 and 3 respectively.**

### 3.2 Phylogenetic analysis of NagZ

The results for phylogenetic analysis of all the genes, including NagZ, in the form of maximum likelihood tree, can be found in **Supplementary File 4.** Phylogenetic analysis of NagZ revealed a distinct evolutionary separation within the studied bacterial taxa. The root node of the tree bifurcates into two primary branches: the first branch groups the Enterobacteriaceae family, forming a cohesive clade. The second branch further splits into two distinct clades, representing *Pseudomonas* and *Acinetobacter*.

The distinct clustering of Enterobacteriaceae within a singular clade suggests a conserved evolutionary path for NagZ within this family, potentially linked to their specific metabolic requirements or environmental adaptations. The separation of *Pseudomonas* and *Acinetobacter* into individual clades under the second branch may indicate unique selective pressures or functional specializations in NagZ across these genera.

### 3.3 Co-evolutionary dynamics of NagZ with associated BLs and related proteins

Our analysis of the co-evolutionary dynamics between NagZ and associated beta-lactamase-related proteins reveals notable phylogenetic correlations (**Table 2**). The branching patterns observed in the phylogenetic trees of NagZ and AmpC exhibit a strong similarity, suggesting potential co-evolution between these two proteins. This co-evolutionary trend was also evident for AmpR, meaning that these proteins share a similar evolutionary trajectory with NagZ. However, no such pattern was detected for glyoxylase, MFS transporter, or AmpD, indicating a lack of phylogenetic congruence between these proteins and NagZ.

**Table 2.** Phylogenetic relationship of genes/proteins among different priority pathogens.

| Gene/Protein | Closely related | Comparatively distantly related |
| --- | --- | --- |
| NagZ | *Acinetobacter baumannii*,<br>*Pseudomonas aeruginosa* | *Escherichia coli* |
| AmpC | *Acinetobacter baumannii*,<br>*Pseudomonas aeruginosa* | *Escherichia coli* |
| AmpR | *Acinetobacter baumannii*,<br>*Pseudomonas aeruginosa* | *Escherichia coli* |
| AmpD | *Acinetobacter baumannii*, *Escherichia coli*;<br>*Acinetobacter baumannii*,<br>*Pseudomonas aeruginosa* | *Escherichia coli* from *Pseudomonas aeruginosa* |
| MDS 1318 | *Acinetobacter baumannii*,<br>*Pseudomonas aeruginosa* | *Escherichia coli* |
| polA | *Acinetobacter baumannii*,<br>*Pseudomonas aeruginosa* | *Escherichia coli* |
| Cubico | *Acinetobacter baumannii*, *Escherichia coli* | *Pseudomonas aeruginosa* |
| Gpmi | *Acinetobacter baumannii*, *Escherichia coli* | *Pseudomonas aeruginosa* |
| MFS | *Acinetobacter baumannii*, *Escherichia coli* | *Pseudomonas aeruginosa* |
| Glyoxylase | *Acinetobacter baumannii*, *Escherichia coli* | *Pseudomonas aeruginosa* |
| MDS 1962 | *Acinetobacter baumannii*, *Escherichia coli* | *Pseudomonas aeruginosa* |
| Pmen 1856 | *Acinetobacter baumannii*, *Escherichia coli* | *Pseudomonas aeruginosa* |
| Pmen 2196 | *Acinetobacter baumannii*, *Escherichia coli* | *Pseudomonas aeruginosa* |
| Pmen 1284 | *Acinetobacter baumannii*, *Escherichia coli* | *Pseudomonas aeruginosa* |
| ELS25187.1 | *Escherichia coli*, *Pseudomonas aeruginosa* | *Acinetobacter baumannii* |
| guaA | *Pseudomonas aeruginosa*, *Escherichia coli* | *Acinetobacter baumannii* |
| SDI51646 | *Pseudomonas aeruginosa*, *Escherichia coli* | None |
| SDI51743 | *Pseudomonas aeruginosa*, *Escherichia coli* | None |
| SDN35178 | *Escherichia coli*, *Pseudomonas aeruginosa* | *Acinetobacter baumannii* |
| SEB92735 | random | random |
| SEJ69490.1 | *Acinetobacter baumannii*,<br>*Pseudomonas aeruginosa* | None |
| Ysh.1 | random | random |
| rpoz | random | random |
| rpob | random | random |

The parallel branching patterns of NagZ with AmpC and AmpR not just suggests a co-evolutionary relationship, but also reflects that functionally-interdependent enzymes or proteins might undergo similar evolutionary changes. The same can be termed as ‘complementary adaptation of functionally-interdependent proteins’ **(Table 2).** The phylogenetic trees showing evolutionary patterns of all these proteins can be seen in **Supplementary File 4.**

### 3.4 Evolutionary dynamics of phosphoglycerate mutase (gpmi)

The phylogenetic analysis of gpmi reveals a distinct divergence at the root, where the first branch consists of a mixed clade of Enterobacteriaceae and *A. baumannii* while the second branch contains *Pseudomonas* species. The mixed clade of Enterobacteriaceae and *Acinetobacter baumannii* may indicate shared functional pressures or evolutionary constraints acting on gpmi within these groups. The distinct *Pseudomonas* clade suggests unique adaptations potentially linked to its ecological niche or metabolic requirements. The results for phylogenetic analysis of all the genes, including gpmi, in the form of maximum likelihood tree, can be found in **Supplementary File 4.**

### 3.5 Co-evolutionary dynamics of gpmi with associated BLs and related proteins

The phylogenetic analysis of gpmi reveals an evolutionary pattern indicative of co-evolution with several betalactamase-domain containing proteins. Gpmi displays a similar branching pattern with MDS_1962 (MBL fold metallohydrolase; PFAM: beta-lactamase domain protein, RNA-metabolising metallo-beta-lactamase), Pmen_1856 (MBL fold metallo-hydrolase; PFAM: beta-lactamase domain protein, RNA-metabolising metallo-beta-lactamase), Pmen_1284 (MBL fold metallo-hydrolase; PFAM: beta-lactamase domain protein; RNA-metabolising metallo-beta-lactamase), and Pmen_2196 (ligase-associated DNA damage response exonuclease; an exonuclease with a beta-lactamase domain). These co-clustering patterns suggest an evolutionary link between gpmi and the studied beta-lactamase domain proteins. It is of utmost importance to note that all enzymes containing the MBL fold are not strictly MBLs, as these differ in their overall function. Additionally, while true MBLs are plasmid-encoded, the MBL-fold containing proteins mentioned from STRING above are chromosomally-encoded. However, this parallel evolution may indicate shared functional or adaptive pressures, possibly linked to metabolic pathways or cellular processes in which both phosphoglycerate mutase (gpmi) and beta-lactamase domain-containing proteins play complementary roles. The coevolutionary signal in these proteins suggests that gpmi may have evolved in concert with MBL-fold containing enzymes, possibly enhancing or coordinating functions crucial for bacterial survival or resistance mechanisms. The phylogenetic trees showing evolutionary patterns of all these proteins can be seen in **Supplementary File 4.**

### 3.6 Evolution of rpoB, rpoZ, polA & guaA

Both *rpoB* and *rpoZ* exhibit a random distribution across the taxa studied, suggesting a lack of clear phylogenetic association or selective pattern. In contrast, *guaA* shows a closer phylogenetic association between *Pseudomonas* and Enterobacteriaceae, whereas *polA* shows a closer phylogenetic association between *Pseudomonas* and Acinetobacter, indicating potential shared evolutionary pressures or functional links between these groups for these specific genes. The random distribution of *rpoB* and *rpoZ* suggests that these genes may have evolved independently across taxa, potentially reflecting a functional independence from taxonomic constraints or ecological specialization.

The results for phylogenetic analysis of all the genes, in the form of maximum likelihood tree, can be found **Supplementary File 4.** However, for a clearer picture, the representative models for evolutionary relationships between *Pseudomonas, Acinetobacter* and *Enterobacteriaceae* have been shown in **Figure 1a-c**.

**Figure 1a.**
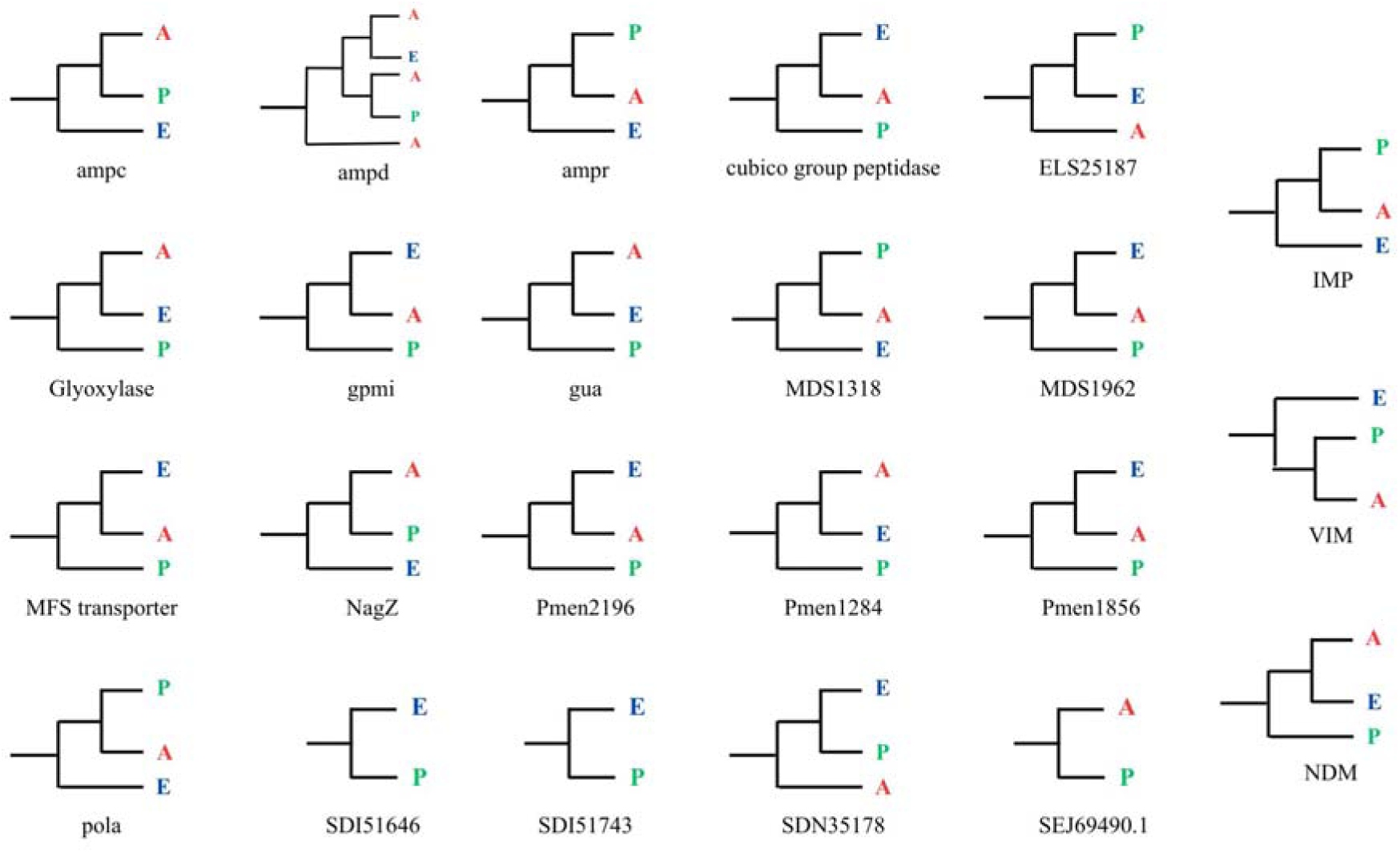
Representative models of proteins for studying their evolutionary relationships between Pseudomonas, Acinetobacter and Enterobacteriaceae.

Further, the phylogenetic relationship of proteins across *Acinetobacter baumannii*, *Pseudomonas aeruginosa*, and *Escherichia coli* has been summarized via **Table 2**.

### 3.7 Chromosomally-encoded BLs vs plasmid-encoded BLs

We found that chromosomally-encoded BLs exhibit a closer evolutionary relationship between *Acinetobacter baumannii* and *Pseudomonas aeruginosa* (as indicated by similar evolutionary patterns of AmpC, NagZ and AmpR) (**Figure 2. (a) correlated to Figure 1b**). On the other hand, plasmid-encoded MBLs, which have supposedly originated from their MBL-fold containing chromosomal ancestors were studied in two ways. First, the evolutionary history of the MBL-fold proteins (chromosomally-encoded) identified from the STRING database was traced. Later, phylogenetic trees were constructed for IMP, VIM and NDM (plasmid-encoded true MBLs), each separately. Relationships between *A. baumannii*, *P. aeruginosa* and Enterobacteriaceae were studied.

**Figure 1b.**
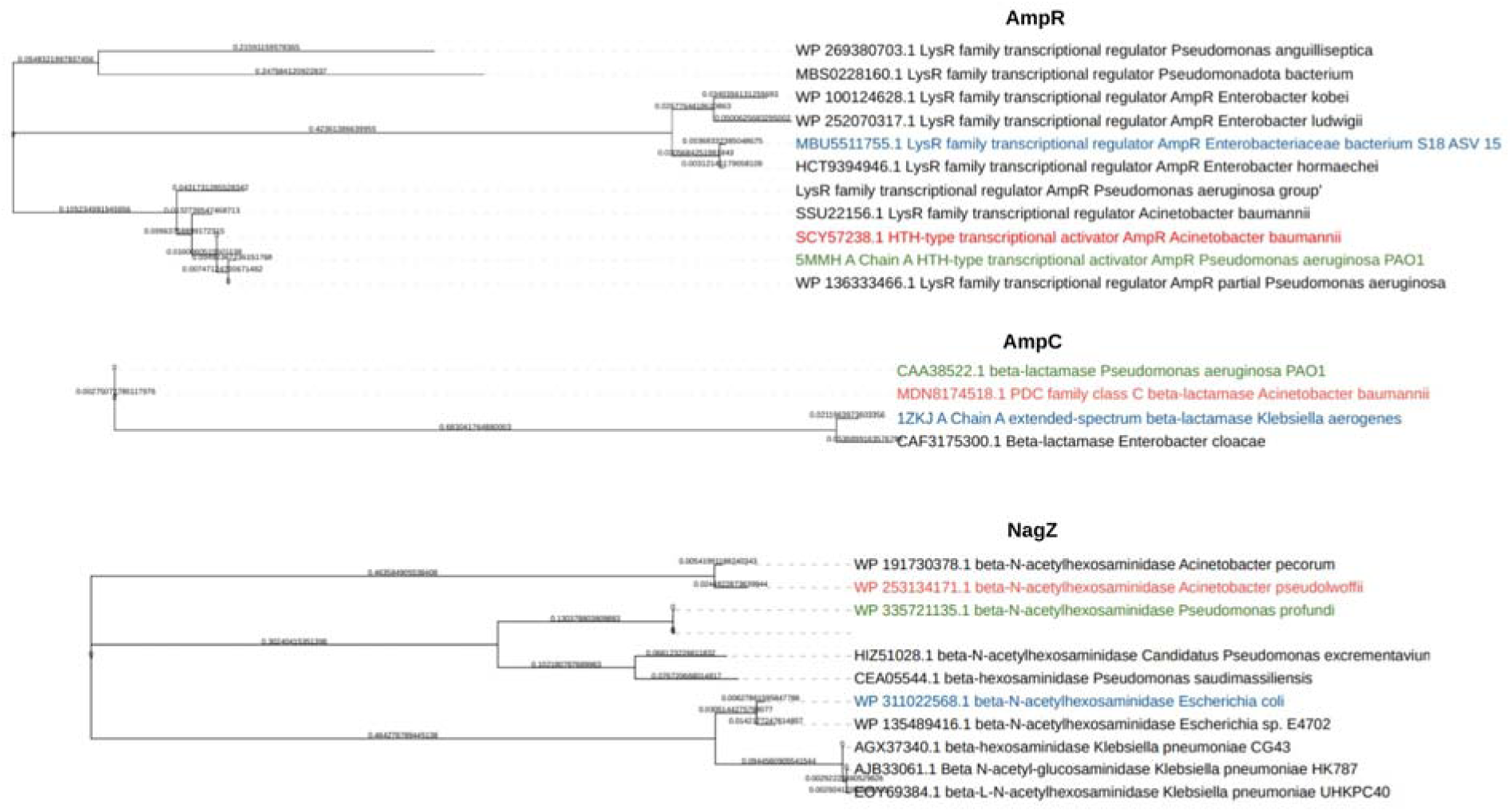
Maximum likelihood phylogenetic trees showcasing evolutionary relationships between Pseudomonas, Acinetobacter and Enterobacteriaceae, for the proteins AmpR, AmpC and NagZ. From the large tree of more than 100 sequences, the representatives were chosen based on the nearest distance to the root node of a clade.

As a result, the MBL-fold containing sequences demonstrated a higher similarity between *Acinetobacter* species and Enterobacteriaceae (**Figure 2. (b) correlated to Figure 1c**). When BL-specific study of the plasmid-encoded MBLs was performed, interesting results were found. **Figure 1a** shows the representative models for evolution of MBLs across different bacteria. The full-length trees for the same can be studied in **Supplementary File 5**. When BLASTP was performed using the clustered nr database, no cluster showed representative as *A. baumannii* in case of VIM. The only representative of clusters containing *A. baumannii* sequences was *E. coli*, indicating that *A. baumannii* and *E. coli* lie closer together for VIM. However, several clusters were found with representatives as *P. aeruginosa* and *E. coli.* On the other hand, in the maximum likelihood trees of VIM, *A. baumannii* and *P. aeruginosa* were found to lie closer together.

**Figure 1c.**
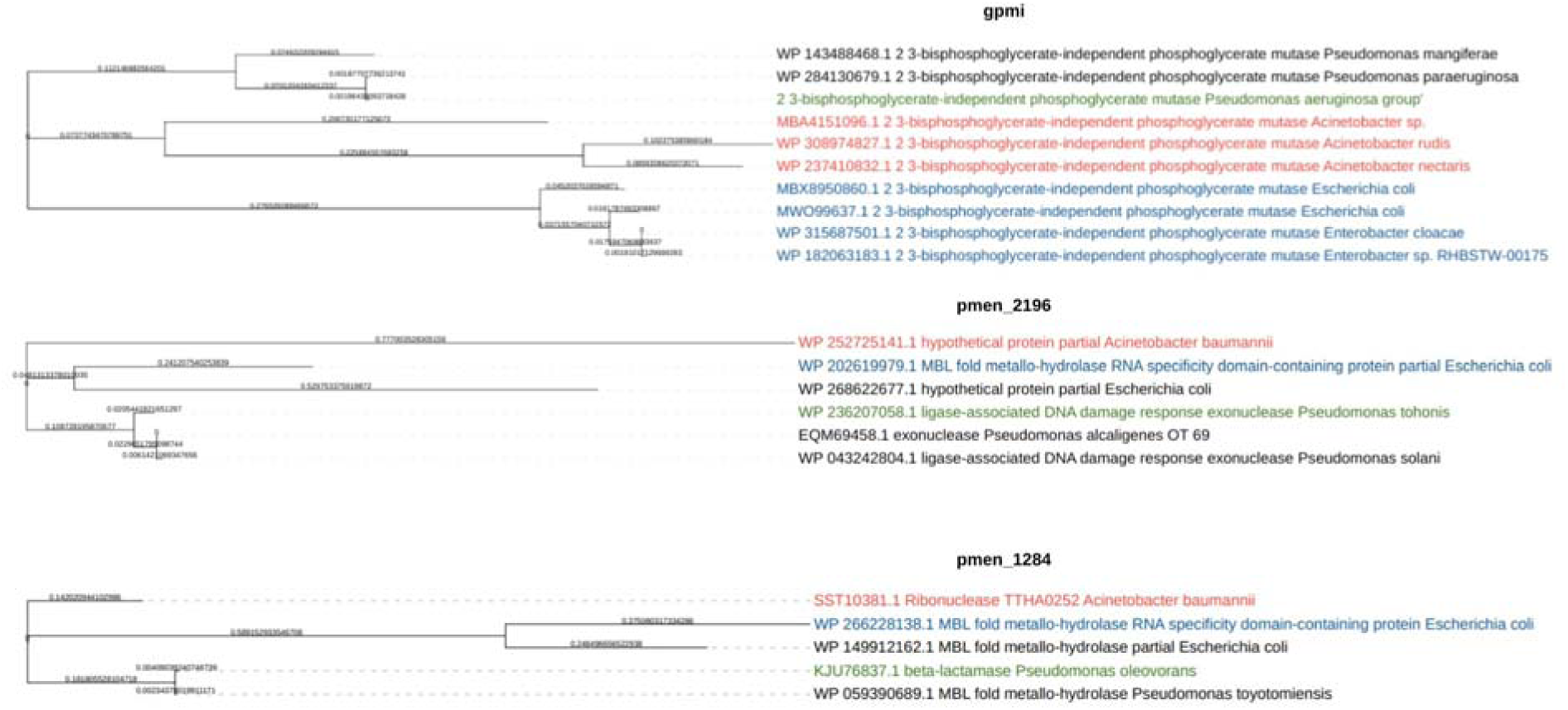
Representative Maximum likelihood phylogenetic trees showcasing evolutionary relationships between Pseudomonas, Acinetobacter and Enterobacteriaceae, for the proteins gpmi and MBL-fold containing proteins: pmen_2196 and pmen_1284. The representatives were chosen based on the nearest distance to the root node of a clade.

**Figure 2.**
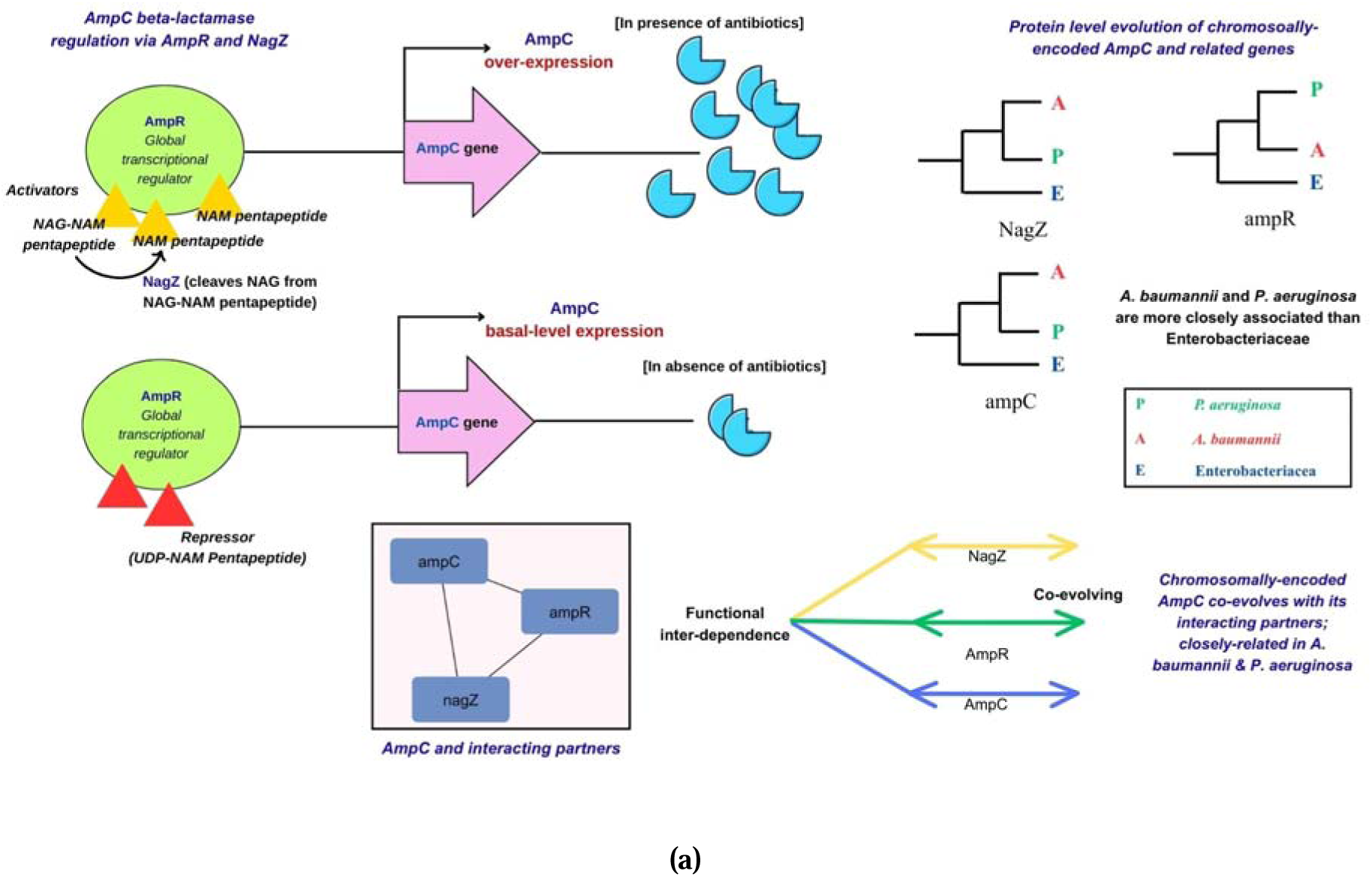
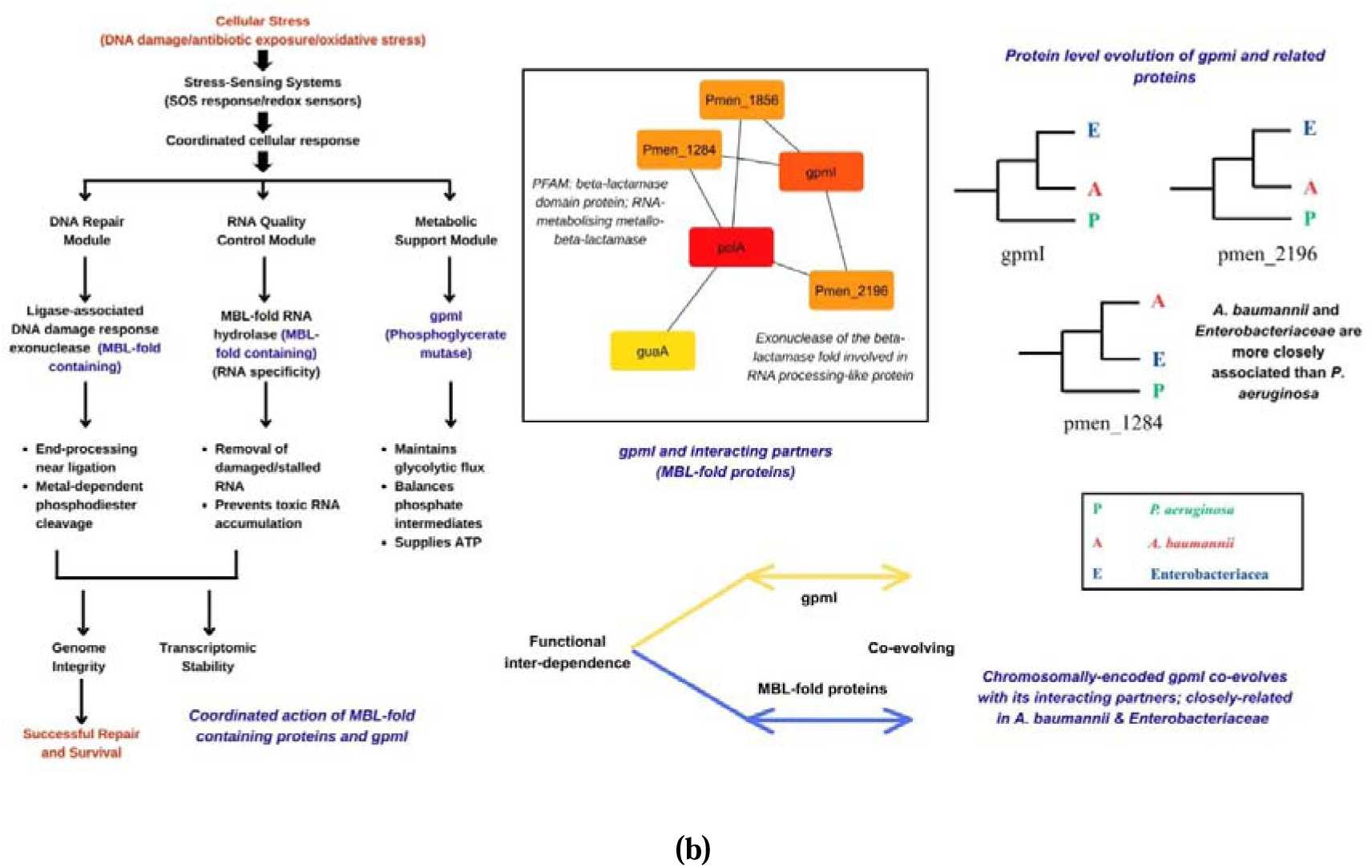
**(a)** Interaction and evolution of proteins where Acinetobacter baumannii and Pseudomanas aeruginosa are closely related. These include AmpC and regulators of AmpC expression-AmpR and NagZ **(b)** Interaction and evolution of proteins where Acinetobacter baumannii and Enterobacteriaceae are closely related. These include MBL-fold containing proteins alongside functionally related proteins gpmI, polA and guaA.

For IMP and NDM, all three pathogens showed considerable representation in BLASTP search. In the maximum likelihood trees of IMP and VIM, *A. baumannii* and *P. aeruginosa* were found to lie closer together. However, for NDM, *A. baumannii* and Enterobactericeae are closer than *P. aeruginosa,* showing that the recent MBLs (NDM type) show evolutionary patters similar to chromosomally-encoded MBL-fold proteins. A possible reason could be that lineages that possess similar chromosomally encoded MBL-fold families tend to exchange plasmid-borne MBLs more readily, because the successful acquisition and persistence of horizontally transferred MBLs is strongly influenced by compatibility with the host’s ancestral fold architecture. Such compatibility includes conservation of metal-binding residues, structural stability of the fold in the host cytosolic environment, compatibility with the host’s folding and chaperone systems, regulatory context, and similarities in codon usage or GC content. Consequently, bacterial groups that share a similar chromosomal MBL-fold background are inherently more permissive to each other’s plasmid-encoded MBLs. This explains why *A. baumannii* and Enterobacteriaceae, whose chromosomal MBL-fold proteins exhibit highly similar evolutionary patterns, show a preferential and more efficient exchange of NDM, leading to its stable integration and a stronger phylogenetic affinity of their NDM sequences. Thus, the observed clustering of NDM within the *A. baumannii*– Enterobacteriaceae clade reflects not a coincidence, but the underlying evolutionary compatibility shaped by their shared MBL-fold ancestry. VIM and IMP do not show this because they are old and have undergone decades of diversification and ecological reshuffling. It is possible that during their earliest evolutionary phases, VIM and IMP followed compatibility patterns similar to NDM, but these signals have been obscured over time by extensive diversification, recombination, and ecological redistribution. Thus, the divergence in patterns between NDM and VIM/IMP reflects differences in age, dissemination pathways, and the evolutionary reshaping of these MBL families.

### 3.8 GC Content analysis

The GC content analysis of the three MBLs, that is, NDM, VIM, and IMP, revealed significant differences, indicating distinct evolutionary origins (**Figure 3**.). Each MBL exhibited nearly consistent GC content across different species, suggesting that these genes retain their nucleotide composition when transferred to new bacterial hosts. However, NDM and VIM were more conserved across different species unlike IMP (**Supplementary File 6**). However, GC content of IMP was found to vary in several cases, indicating its lesser conserved nature as compared to VIM and NDM. **Figure 3** (a) and (b) show that NDM has a thin range of GC content percentage over which the sequences have been distributed, showing its highly conserved nature and lower variability as compared to the other MBLs taken into study. This indicates lower rates of HGT for NDM. On the other hand, IMP shows maximum variability out of the three MBLs, indicative of higher rates of HGT.

**Figure 3.**
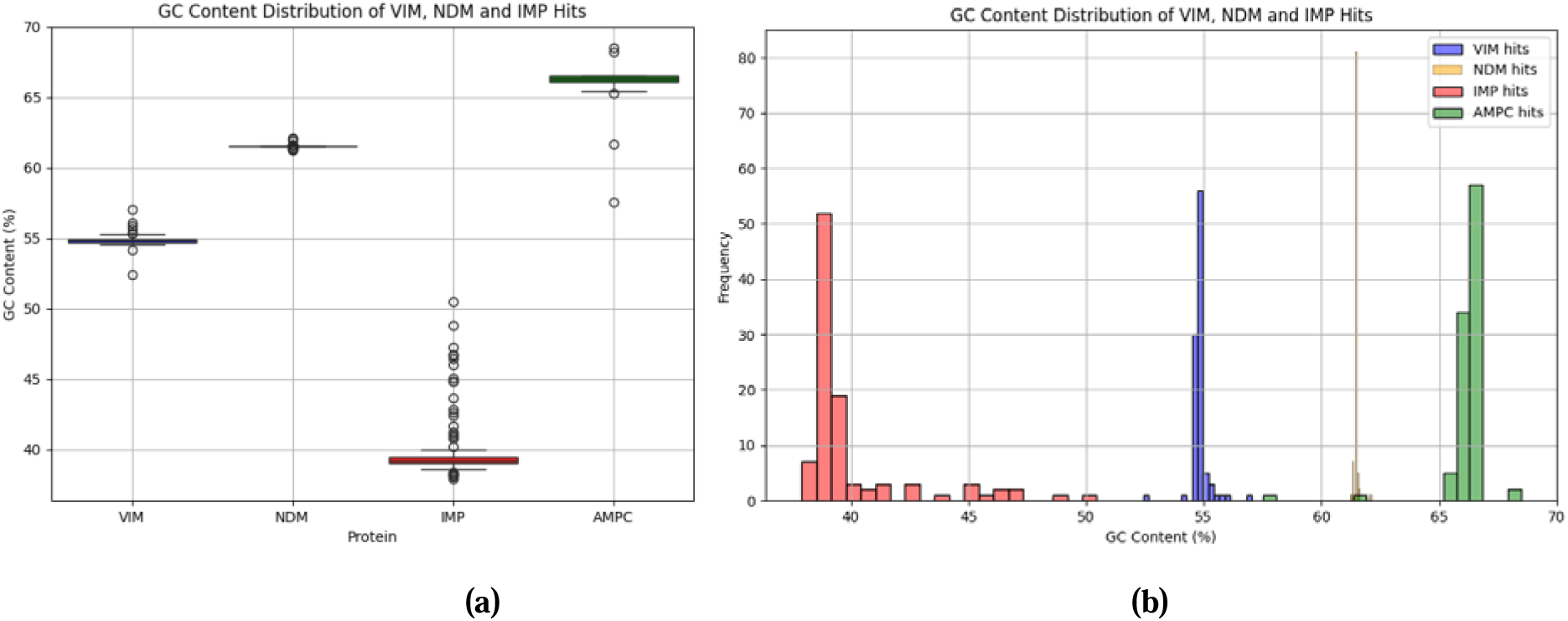
GC Content comparison of different BLs, and clustering of MBLs. (a) Distribution of GC content per MBL, indicating mean and standard deviation alongside highlighting outliers. (b) Frequency distribution of MBLs based on GC content

IMP was shown to possess lowest GC content out of all BLs taken for study. The average GC content for IMP was 40.07%, for VIM was 54.85%, for NDM was 61.51% and for AmpC was found to be 66.18%. The GC content analysis of AmpC can be seen in **Figure 3** and in **Supplementary File 6.** A comparison of the same with MBLs can be seen in **Figure 3a-b**. An interesting observation found was that lower GC content was shown to promote variability of the enzyme, such as in IMP and VIM, as we can see in **Figure 3b** that the range of GC percent is very big for IMP as different sequences vary a lot in their percent GC content alongside IMP having lowest GC content out of the three MBLs studied. This is in line with the fact that lower GC content promotes variability (IMP and VIM) whereas a higher GC content promotes stability (NDM and AmpC). **Figure 3** (b) clarifies NDM to be less evolved due to less spread of sequences (showing less variability) and a sharp peak for GC content.

The fact that the GC distribution of VIM lies between IMP and NDM, forced us to questionis VIM acting as an evolutionary link between IMP and NDM? To answer this, we ran the alignment for all MBLs together, followed by Pairwise sequence similarity mapping (PaSiMap). In PaSiMap results, we found VIM to lie between IMP and NDM, further supporting our hypothesis that VIM could act as an evolutionary link between the two (**Figure 4a**).

**Figure 4.**
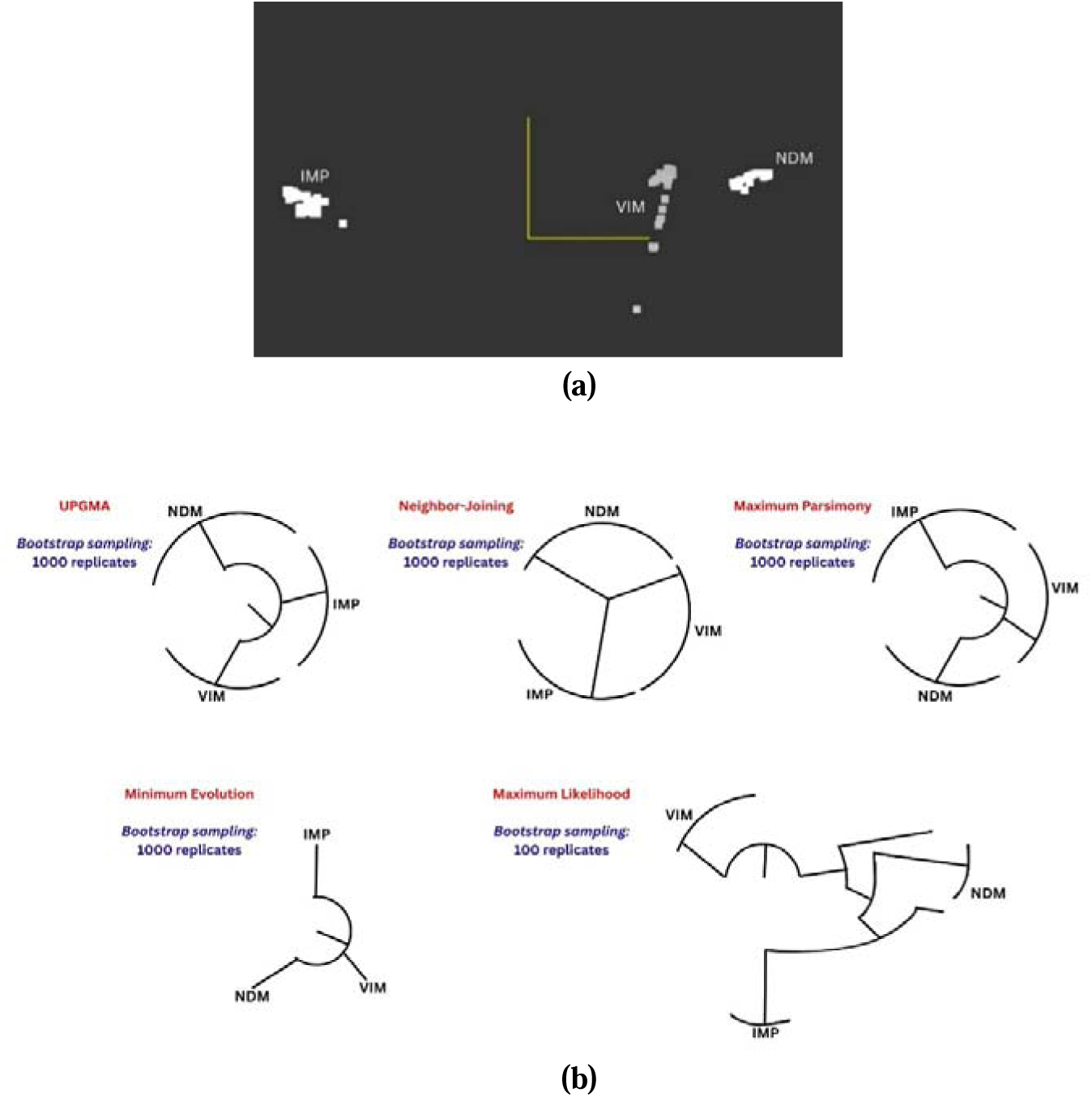
Distribution of the three MBLs (IMP, VIM and NDM) showing that sequences of the three MBLs are distinctly clustered, with sequences of VIM lying between that of NDM and IMP. **(a)** PaSiMap (Pairwise Sequence Similarity Mapping) analysis of IMP, VIM and NDM showing presence of VIM between the other two MBLs or proximity of VIM to both NDM and IMP. This further indicates that VIM might be the connecting link between IMP and NDM which lie farther apart from each other. **(b)** Representative models of trees showcasing the evolutionary relationship between the Metallo-beta-lactamases-IMP, VIM and NDM.

To further test our hypothesis that VIM could be an evolutionary link between IMP and NDM, several algorithms were employed for building phylogenetic trees of MBLs. **Figure 4b** shows the representative trees for explainability of relationship between NDM, VIM and IMP. The full-length trees for the same can be visualized in **Supplementary File 7.** In NJ, ME and MP trees constructed using bootstrap sampling method of 1000 replicates, we found that VIM lies between IMP and NDM. This indicates that VIM might be the evolutionary bridge between IMP and NDM. However, in UPGMA (bootstrap 1000), IMP was found to be located between NDM and VIM. Since UPGMA s weaker in identifying relations than MP, ME and NJ, and since majority of the bootstrap trees show VIM as an intermediate clade, there are high chances of VIM being the evolutionary link between NDM and IMP.

In ML tree (bootstrap 100), it was found that VIM further branched into NDM, followed by NDM branching into IMP, but since the bootstrap value of this branching was 0.53, it was a very low confidence prediction. On the other hand, the previous algorithms showed bootstrap values as 1 while branching from the root node to IMP, VIM and NDM clades. This suggests that these three beta-lactamase families are closely related, and that VIM might be the evolutionary bridge between NDM and IMP. This does not necessarily mean that NDM (most recent) came from VIM (intermediate) and that VIM came from IMP (older). Rather, if a sequence occupies an intermediate phylogenetic position, it may do so because it exhibits features derived from both neighboring lineages, functioning as a mosaic enzyme **(Truty et al., 2023)**. Thus, VIM could be a mosaic sequence having acquired characteristics from both NDM and IMP via HGT, recombination or other mechanisms. To further test our hypothesis, we studied if VIM shares protein features, motifs and domains from both IMP and NDM or not. Therefore, pfeature analysis, PFAM domain analysis and motif analysis were performed.

### 3.9 Protein features: carbapenem resistant vs non-carbapenem resistant BLs

The clustering of bacterial species based on contribution of each of the residues, atoms and bonds, and physiochemical properties to the principal components can be visualized in PCA scatter plots contained in **Figure 5a-c**.

**Figure 5a.**
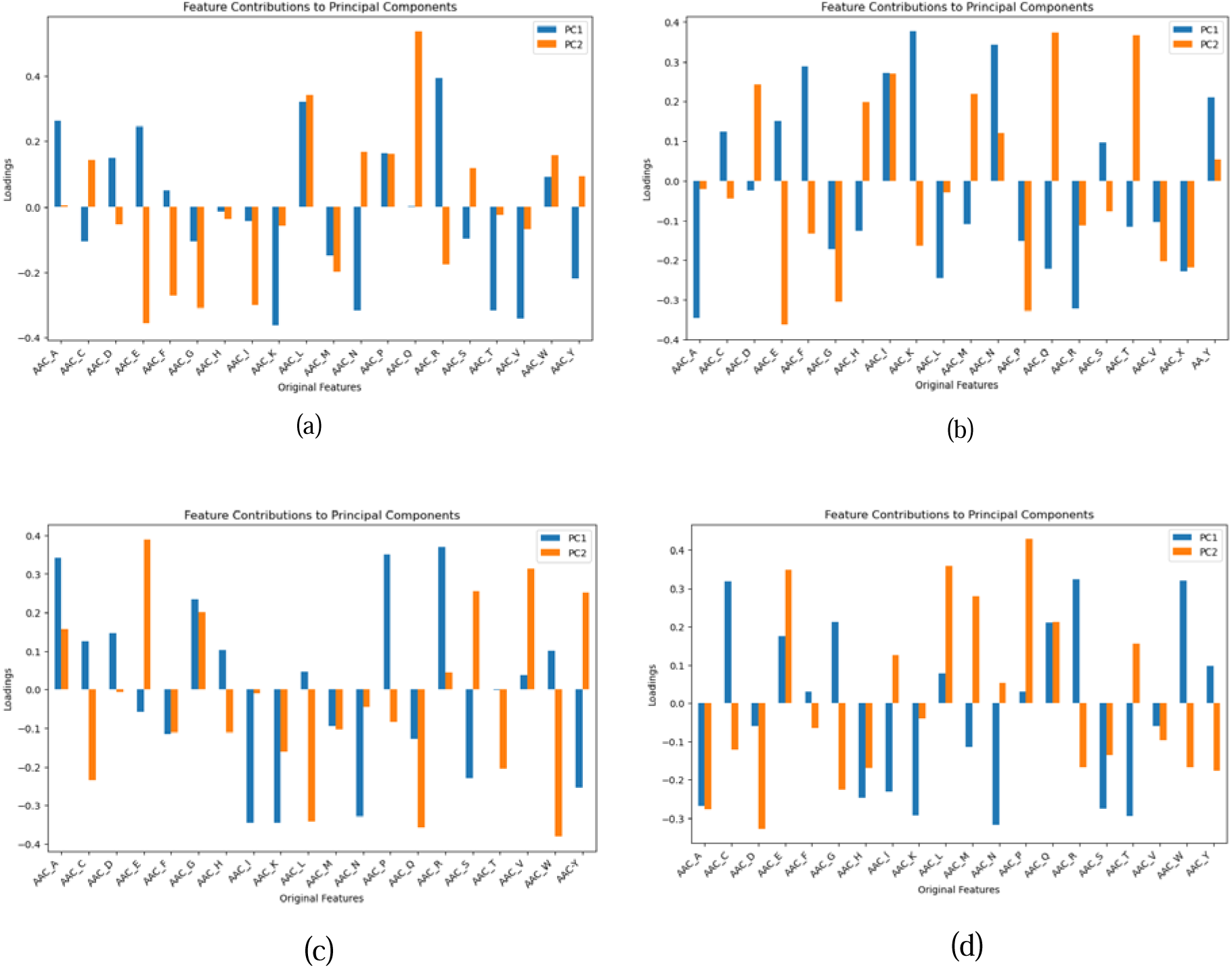
Amino acid composition (AAC) of Beta-lactamases. (a) AmpC (b) IMP (c) VIM (d) NDM

#### 3.9.1 Amino Acid Composition (AAC)

Principal component analysis (PCA) revealed distinct patterns in amino acid contributions across AmpC, NDM, VIM, and IMP **(Table 5 and Figure 5a).** PC1 +ve contributions reflect that AmpC, VIM and NDM sequences are clustered based on similar amino acids. Similarly, PC2 +ve contributions reflect that AmpC and NDM sequences are clustered based on similar residues. This emphasizes that there could be a link in evolvability of BLs from AmpC to VIM and NDM.

**Table 5.**
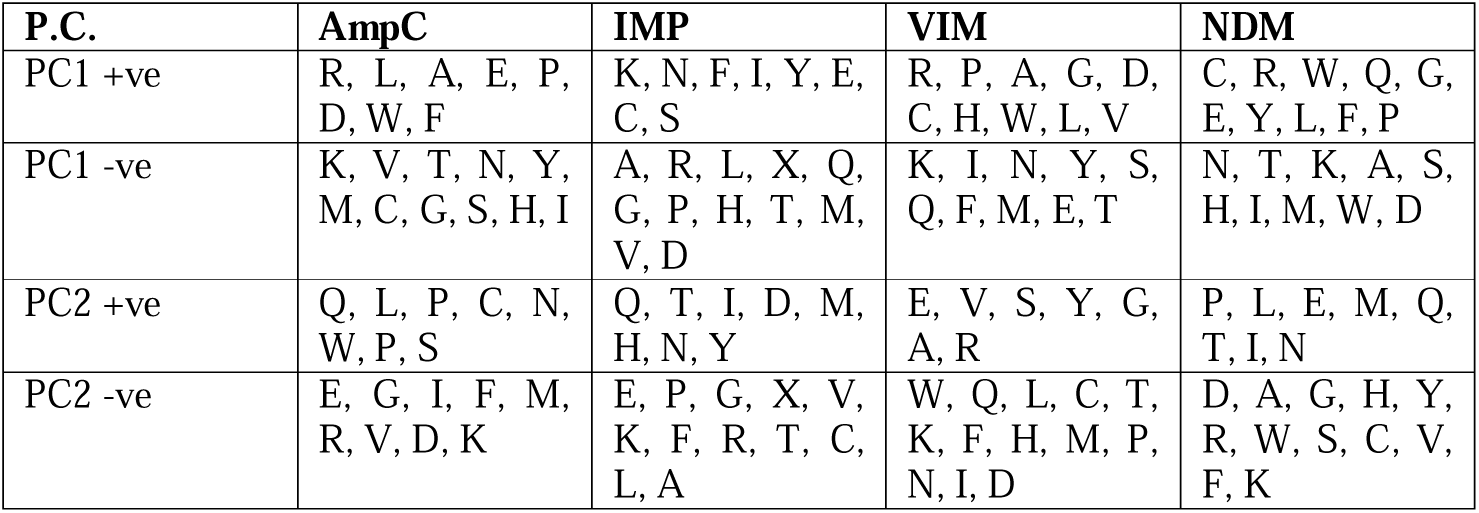
Amino Acid Composition.

| P.C. | AmpC | IMP | VIM | NDM |
| --- | --- | --- | --- | --- |
| PC1 +ve | R, L, A, E, P, D, W, F | K, N, F, I, Y, E, C, S | R, P, A, G, D, C, H, W, L, V | C, R, W, Q, G, E, Y, L, F, P |
| PC1 -ve | K, V, T, N, Y, M, C, G, S, H, I | A, R, L, X, Q, G, P, H, T, M, V, D | K, I, N, Y, S, Q, F, M, E, T | N, T, K, A, S, H, I, M, W, D |
| PC2 +ve | Q, L, P, C, N, W, P, S | Q, T, I, D, M, H, N, Y | E, V, S, Y, G, A, R | P, L, E, M, Q, T, I, N |
| PC2 -ve | E, G, I, F, M, R, V, D, K | E, P, G, X, V, K, F, R, T, C, L, A | W, Q, L, C, T, K, F, H, M, P, N, I, D | D, A, G, H, Y, R, W, S, C, V, F, K |

#### 3.9.2 Atom and Bond Composition (ABC)

PCA was further utilized to reveal distinctive atomic and bond compositions that contribute to clustering of bacterial species **(Table 6 and Figure 5b).** PC2 +ve contributions show that NDM shares features from AmpC and VIM as seen in previous findings as well. It shared the double bond of AmpC and C-atom of VIM. PC1 +ve contributions showed high variability or diverse set of bonds for IMP, indicating that each could be affected leading to distinct clustering of different bacterial species, further re-emphasizing its least stable nature.

**Table 6.**
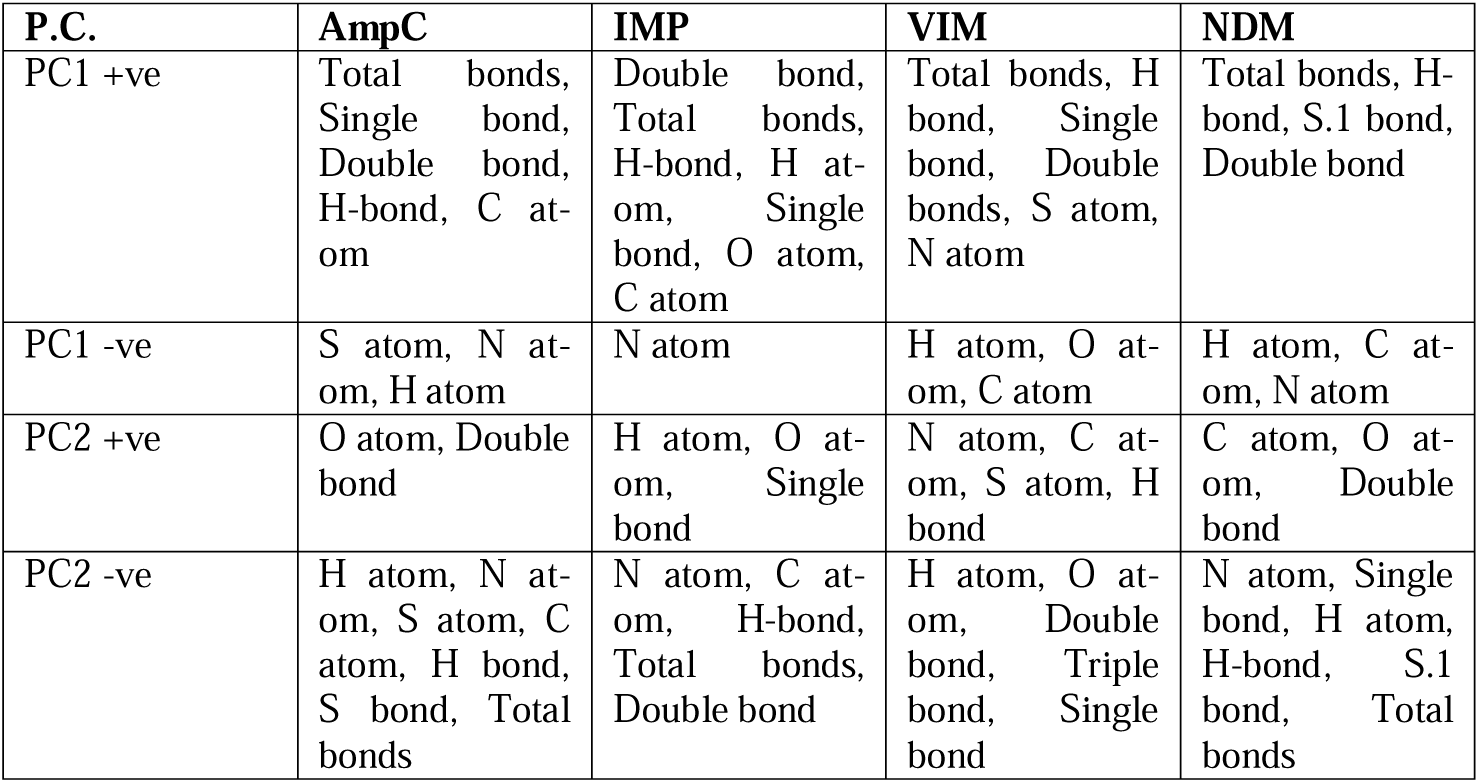
Atoms and Bonds Composition.

**Figure 5b.**
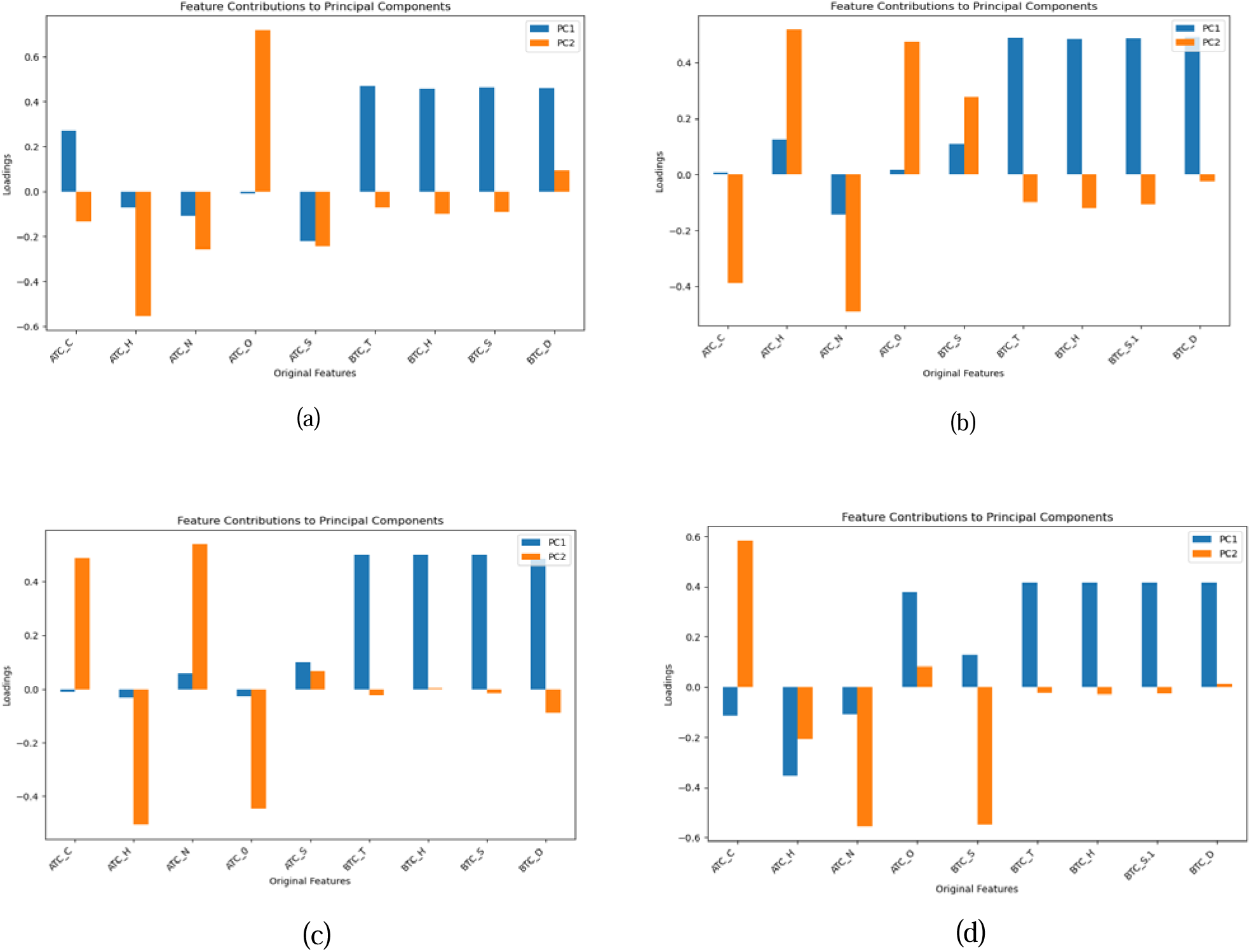
Atoms and bonds composition (ABC) of Beta-lactamases. (a) AmpC (b) IMP (c) VIM (d) NDM

#### 3.9.3 Advanced Physicochemical Properties (APP)

PCA of advanced physicochemical properties (Z-descriptors) helped us identify the distribution of unique attributes across enzymes **(Table 7 and Figure 5c)**. Here, lipophilicity (Z1), bulk (Z2), polarity / charge (Z3) and electronic effects (Z4 and Z5) all were taken into account.

**Table 7.** Advanced physiochemical properties.

| **P.C.** | **AmpC** | **IMP** | **VIM** | **NDM** |
| --- | --- | --- | --- | --- |
| PC1 +ve | Z2, Z5, Z1, Z4 | Z1, Z2, Z4, Z5 | Z5, Z3, Z1, Z4 | Z5, Z3 |
| PC1 -ve | Z3 | Z3 | Z2 | Z2, Z4, Z1 |
| PC2 +ve | Z1 | Z1 | Z4, Z2, Z5 | Z1, Z3, Z4, Z5 |
| PC2 -ve | Z5, Z3, Z4, Z2 | Z4, Z2, Z3, Z5 | Z1, Z3 | Z2 |

**Figure 5c.**
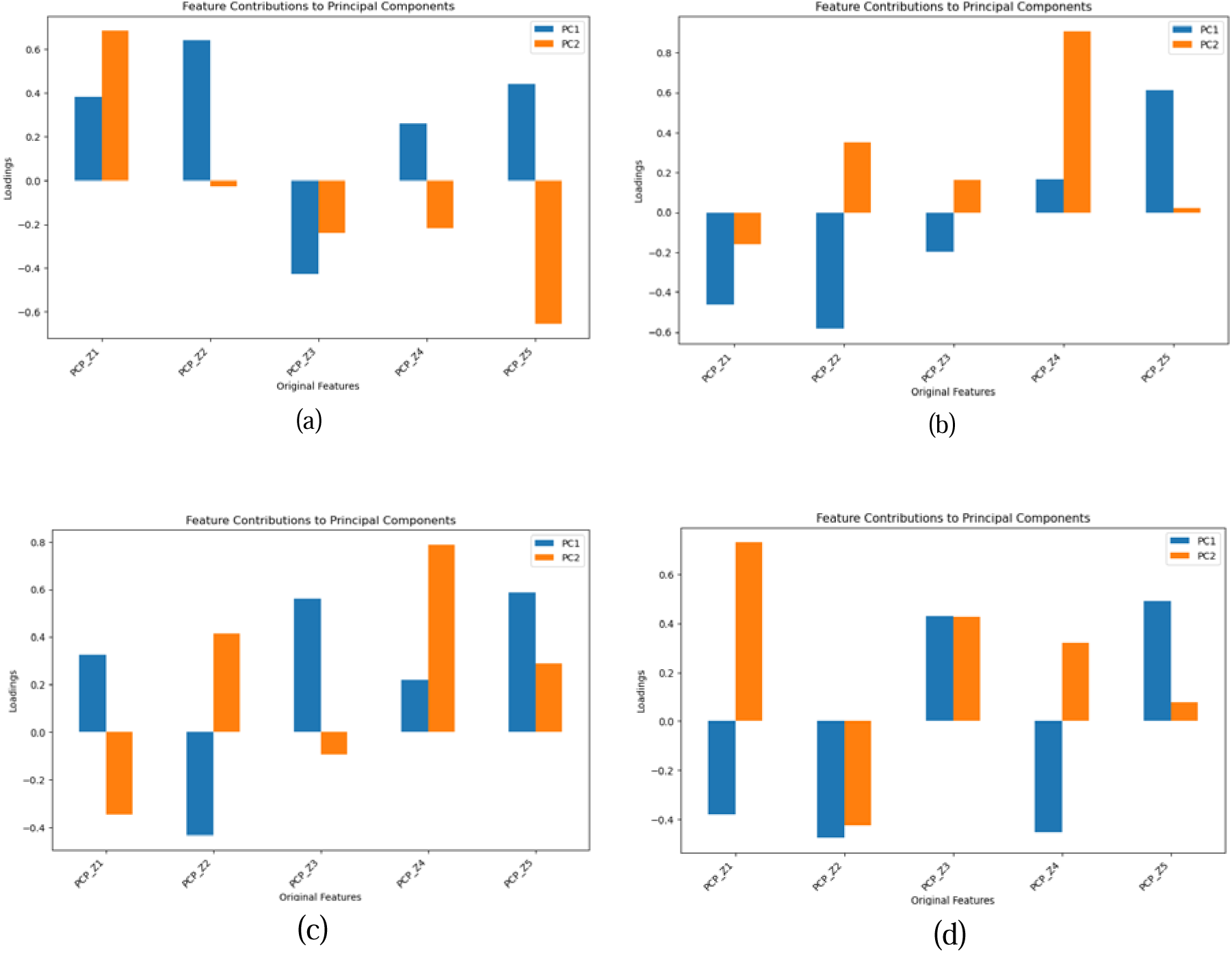
Advanced Physicochemical properties (APP/PCP) of Beta-lactamases. (a) AmpC (b) IMP (c) VIM (d) NDM

Positive contributors to PC1 included Z5 (electronegativity) for all enzymes and Z1 (lipophilicity) for AmpC, IMP and VIM. However, only VIM and NDM show Z3 and Z5 as large contributors in PC1. This suggests that VIM shares features from both NDM and IMP. Reliance of NDM on Z5 and Z3 (electronic properties) suggest specialization for electronic interactions. For PC2, AmpC relied on Z1, emphasizing hydrophobic interactions. NDM displays versatility with multiple descriptors (Z1, Z3, Z4, Z5), enabling adaptation to diverse environments.

Additionally, Z-descriptors reveal differences in resistance strategies. NDM’s reliance on Z5 and Z3 indicates specialized electronic interactions, potentially linked to its broad-spectrum resistance. AmpC’s focus on Z1 suggests a hydrophobic interaction-driven mechanism, while VIM and IMP’s use of Z2 and Z5 points to both steric/bulk and electronic adaptations for inhibitor resistance.

Finally, these carbapenem-resistant BLs (VIM, NDM, IMP) demonstrate unique structural features compared to the carbapenem-non-resistant beta-lactamase AmpC. The presence of sulfur and oxygen atoms in VIM and NDM highlights their capability for diverse catalytic interactions, enabling resistance to carbapenems. IMP’s nitrogen-centric composition further supports its adaptation for specific antibiotic challenges. In contrast, AmpC’s reliance on hydrophobic interactions, as reflected by its Z1-positive contributions, aligns with its role in resisting cephalosporins rather than carbapenems. These differences underscore the specialized evolutionary paths that equip resistant enzymes with broader substrate adaptability and advanced catalytic mechanisms.

Moreover, plasmid-encoded beta-lactamases (VIM, NDM, IMP) exhibit enhanced versatility due to their ability to spread horizontally across bacterial populations. This is reflected in their broader physicochemical adaptability, including reliance on Z5 (electronegativity) and Z3 (electronic properties). These traits enable them to confer resistance to a wide range of beta-lactam antibiotics, including carbapenems, in diverse environments. On the other hand, the chromosomally-encoded AmpC demonstrates a narrower focus, with adaptations emphasizing hydrophobic interactions and structural stability, as seen in its reliance on Z1 and fewer versatile residues. These distinctions highlight the evolutionary and functional advantages plasmid-encoded enzymes have in promoting antibiotic resistance.

### 3.10 PFAM Domain Analysis of Beta-Lactamases (BLs)

The PFAM domain analysis helped us identify conserved protein domains. These domains define the functional and structural characteristics of BLs. **Figure 6** shows the visual representation of PFAM domain distribution for different beta-lactamases.

**Figure 6.**
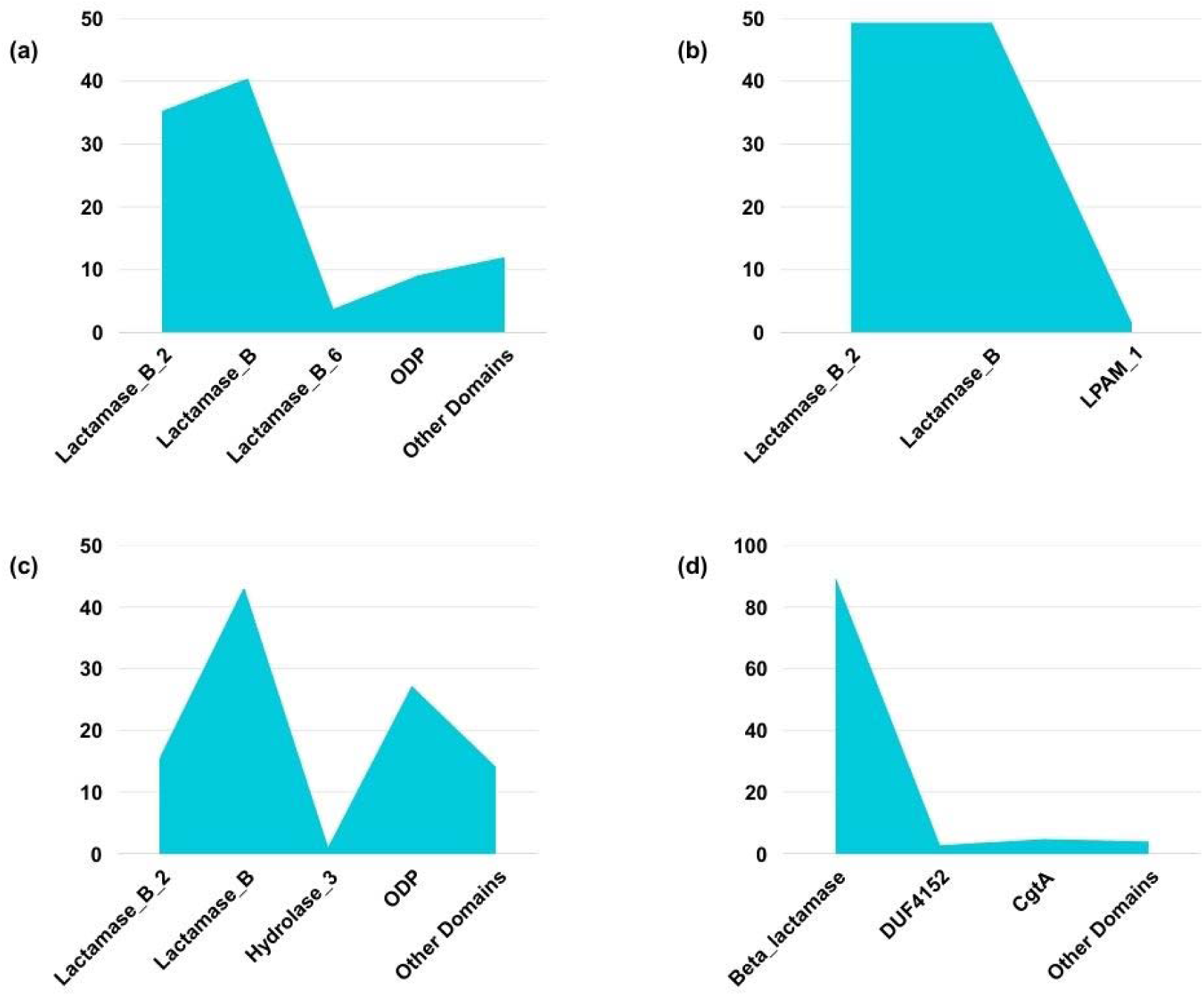
Distribution of major PFAM domains for different beta-lactamases. (a)**VIM** Beta-Lactamases: Dominated by Lactamase_B (40.31%) and Lactamase_B_2 (35.14%), with ODP (9.04%) playing a notable role. (b) **NDM** Beta-Lactamases: Highly conserved, with Lactamase_B and Lactamase_B_2 each comprising 49.3%. (c) **IMP** Beta-Lactamases: Shows more diversity, with Lactamase_B (42.93%), ODP (27.02%), and other minor domains. (d)**AmpC** Beta-Lactamases: Mostly Beta-lactamase domain (89.17%), with CgtA (4.46%) and DUF4152 (2.55%) suggesting functional links to glycosylation. It is to note that the figure here highlights the major domains in BLs. However, few other domains are also present which have been discussed in detail further.

**Figure 6** once again depicts NDM to be less evolved or more conserved to due fewer accessory domains than other MBLs.

**Table 8**. summarizes the PFAM domain analysis for different BLs.

**Table 8.** Comparative PFAM Domain Analysis of Beta-Lactamases.

| Feature | NDM | VIM | IMP | AmpC |
| --- | --- | --- | --- | --- |
| **Core Domains** | Lactamase_B, Lactamase_B_2 | Lactamase_B, Lactamase_B_2 | Lactamase_B, Lactamase_B_2, ODP | Beta-lactamase, Beta-lactamase_2 |
| **Structural Variability** | Low (Highly conserved) | High | Moderate | Low |
| **Unique Domains** | No | DUF6491, RelB_transactiv | Hydrolase_3, DUF1138, DUF1846 | CgtA, DUF4152 |
| **Presence of ODP Domain** | No | Yes | Yes | No |
| **Presence of DUF Domains** | No | Yes (Higher variability) | Yes (Moderate variability) | Yes (Lower variability) |
| **Association with Cell Wall Metabolism** | Indirect | Indirect | Potential (via Hydrolase_3) | Direct (CgtA association) |

We found that NDM is the most conserved beta-lactamase, containing only Lactamase_B and Lactamase_B_2 without additional domains, indicating structural optimization for function. VIM and IMP show greater domain diversity, with VIM exhibiting DUF domains and regulatory elements (RelB_transactiv), while IMP includes Hydrolase_3 and additional DUFs, suggesting functional adaptability and evolutionary divergence. Additionally, the ODP domain is exclusive to metallo-beta-lactamases (VIM, IMP), indicating a potential role in substrate specificity or structural stabilization. AmpC is structurally distinct from MBLs, lacking the ODP domain and instead containing CgtA (linked to glycosylation and cell wall interactions), emphasizing a different evolutionary pathway and functional role.

Overall, plasmid-encoded beta-lactamases (VIM, NDM, IMP) exhibit more domain variability, possibly enhancing adaptability to diverse antibiotics, while chromosomally encoded AmpC remains more conserved, relying on hydrophobic interactions rather than diverse catalytic elements.

### 3.11 Cluster-wise PFAM Analysis of Beta-Lactamases

We further correlated our results of PFAM analysis with clusters that were previously derived using pfeature. **Table 9** shows summarized results for cluster-wise PFAM analysis of BLs. **Figure 7** shows the distribution of PFAM domains across different clusters of the same BL.

**Figure 7.**
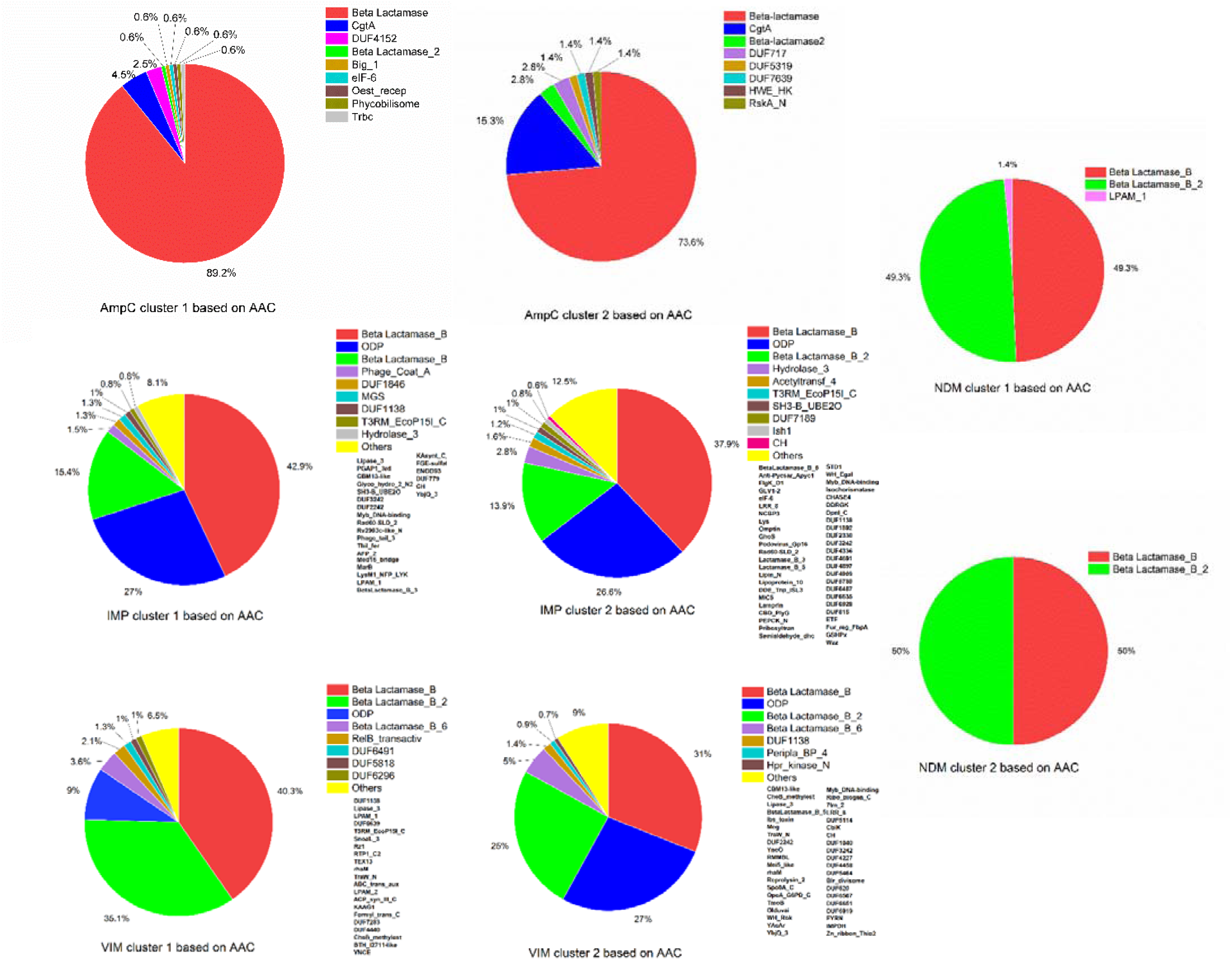
PFAM domain distribution across AmpC, VIM, NDM and IMP.

**Table 9.**
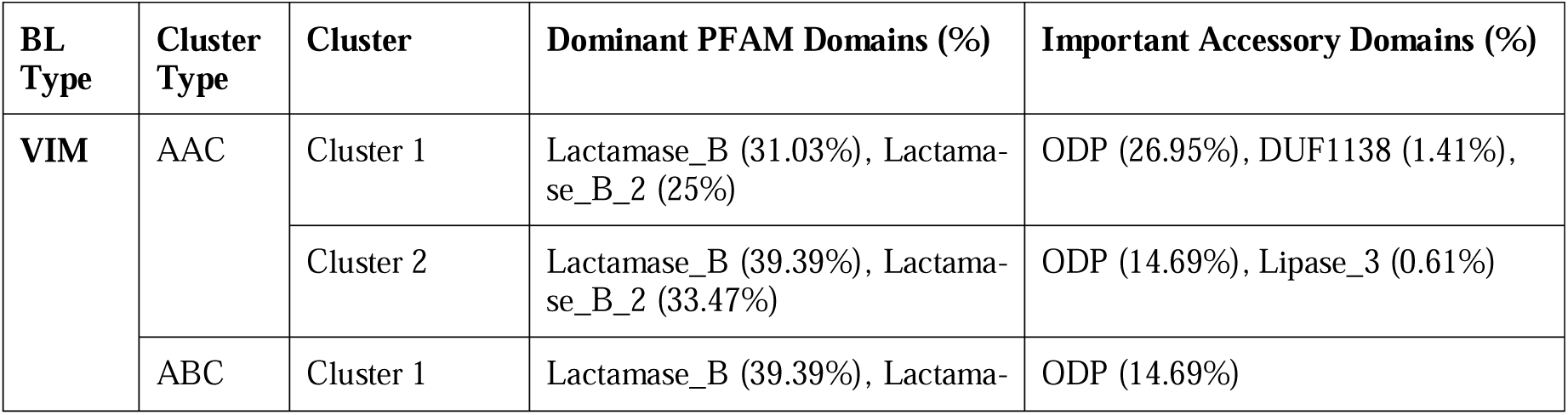
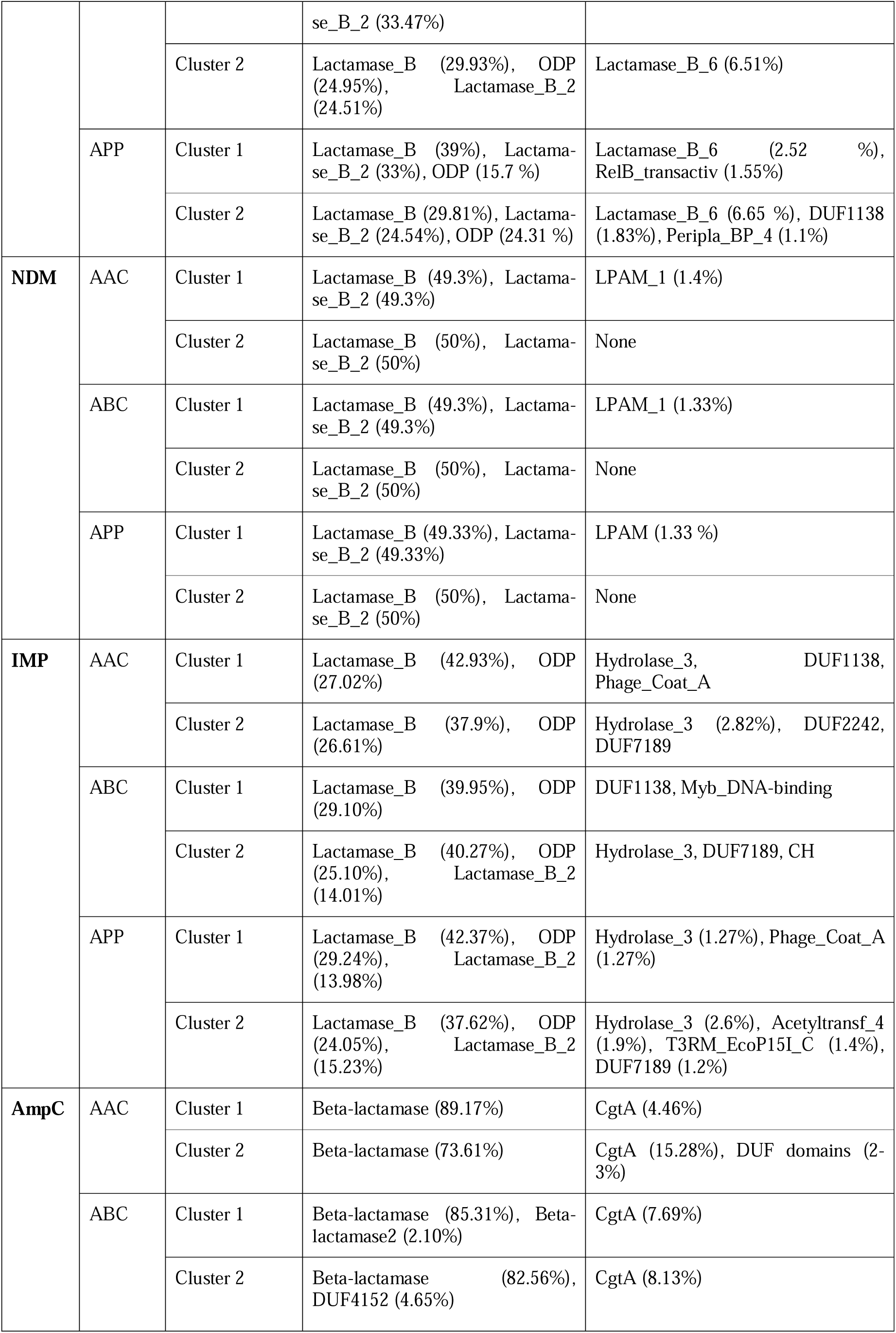
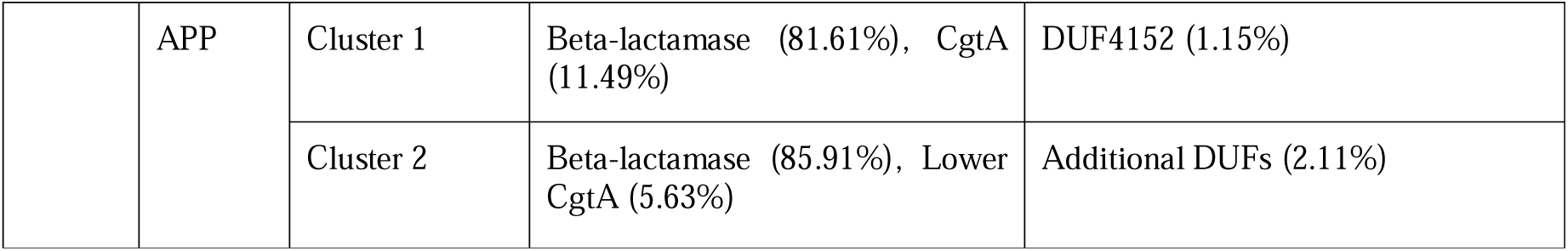
Cluster-wise PFAM Analysis of Beta-Lactamases Based on Amino Acid Composition (AAC), Atoms and Bonds Composition (ABC), and Advanced Physicochemical Properties (APP)

The cluster-wise PFAM analysis as summarized in the **Table 9 and Figure 7**. shows that two clusters of the same BL vary significantly in both the core Beta-lactamase domains as well as accessory domains, except for NDM. In case of NDM, the Beta-lactamase domain distribution is same across different clusters, even though the accessory domains vary in their percentage. Thus, NDM is the most conserved across all cluster types, with minimal structural or physicochemical variability. VIM and IMP display greater diversity, particularly in ODP domains, suggesting higher adaptability to different environments. AmpC remains structurally conserved but still exhibits some considerable variability.

### 3.12 Final interpretation of PFAM domain distribution across MBLs and AmpC

A final conservation matrix for PFAMs across all priority pathogen was obtained for all enzymes and can be studied in **Figure 8a**, showing a heatmap for the same. The heatmap was generated using the PFAM domain conservation scores, which were assigned based on the presence of each domain in different bacterial groups.

**Figure 8.**
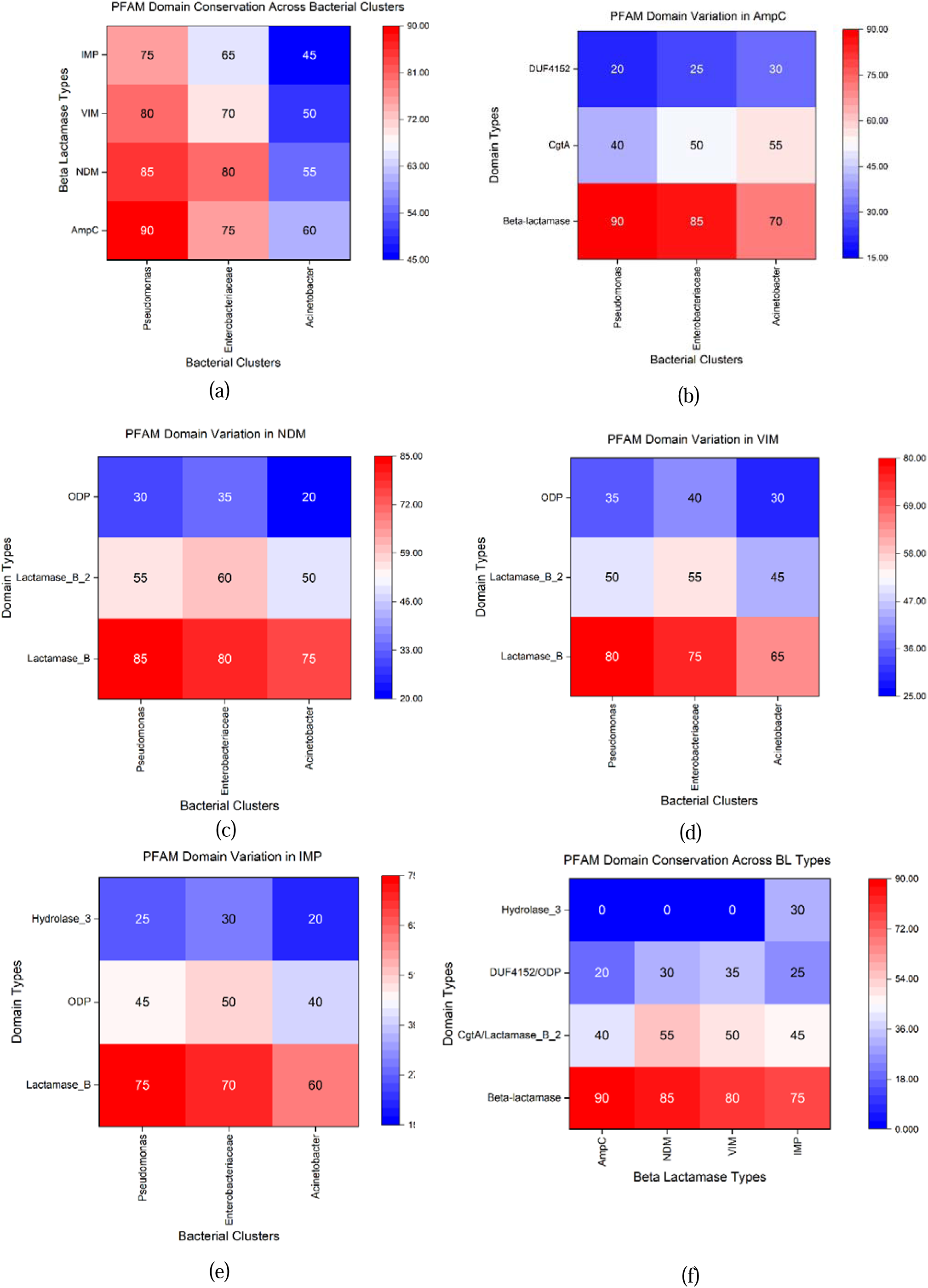
PFAM domain distribution of beta-lactamases across different bacteria. (a) PFAM domains conservation matrix for all BLs across different bacteria. (b) PFAM domains conservation matrix for AmpC across different bacteria. (c) PFAM domains conservation matrix for NDM across different bacteria. (d) PFAM domains conservation matrix for VIM across different bacteria. (e) PFAM domains conservation matrix for IMP across different bacteria. (f) Conservation matrix of key domains across all BLs.

AmpC domain variations across bacterial clusters were studied by calculating conservation scores and drawing heatmap. The Beta-lactamase, CgtA, and DUF4152 domains in AmpC were checked across different bacterial clusters. The final conservation matrix for AmpC can be seen in **Figure 8b** which shows the heatmap for the same. Similarly, NDM domain variations across bacterial clusters were studied. Lactamase_B, Lactamase_B_2, and ODP domains of NDM beta-lactamases were checked across bacterial groups. The final conservation matrix heatmap for NDM can be seen in **Figure 8c**. Further, a heatmap of VIM domain variations across bacterial clusters was generated. Lactamase_B, Lactamase_B_2, and ODP domains in VIM beta-lactamases were analyzed similarly. The final conservation matrix for VIM can be seen in **Figure 8d**. A heatmap of IMP domain variations across bacterial clusters was also generated. Lactamase_B, ODP, and Hydrolase_3 domains in IMP beta-lactamases were analyzed. The final conservation matrix for IMP can be seen in **Figure 8e**.

**Figure 8a** clearly shows that PFAM domains lying within IMP are least conserved across different bacteria, followed by VIM, NDM and finally AmpC. A closer look at **Figure 8b-f** shows that the core lactamase domains are most conserved in *P. aeruginosa*, followed by Enterobacteriaceae. On the other hand, *Acinetobacter* shows high domain variability.

Finally, a comparative study of PFAM domain conservation across all BL types was performed. The conservation percentage of key PFAM domains across all beta-lactamases (AmpC, NDM, VIM, IMP) was recorded. The final conservation matrix of key domains across all BLs can be seen in **Figure 8f**. The results highlight that the core lactamase domains are more conserved within the BLs as compared to accessory domains.

### 3.13 Motif Analysis of Beta-Lactamases: Insights from MEME and TOMTOM Discovery

The motif analysis conducted using MEME and TOMTOM provided significant insights into the structural and functional conservation of BLs. The analysis revealed that while some motifs were highly conserved across different betalactamase classes, others displayed notable sequence variations that could reflect functional divergence.

#### 3.13.1 AmpC vs. IMP: Conservation and Divergence in Functional Regions

Comparison of AmpC and IMP motifs highlighted multiple regions of sequence conservation, with significant statistical relevance indicated by low p-values and E-values. Several motifs were found in the same orientation (+) in both enzymes, suggesting conserved structural or functional regions that may contribute to similar enzymatic activities. One of the strongest motif matches was FAACTC in AmpC with consensus FAAGTT, with a q-value of 0.0133, indicating a conserved functional region. A particularly strong match was identified between DVTFATDTTFTVGKQ (IMP) and FAACTC (AmpC) (p-value = 0.0003), indicating moderate similarity in sequence elements. Another highly significant alignment was observed between MLSGVRNLLISALLLGAGNCM (AmpC) and MKKLFLLLLFL (IMP), which showed strong similarity with an E-value of 0.018, suggesting evolutionary conservation in sequence structure. Moreover, a highly conserved region was observed in MRIAIRGCAALMIGAIANAQA (IMP) and MRFTITALALLLIGQ (AmpC), with an extremely low p-value of 0.00016, suggesting evolutionary conservation in functionally significant regions.

Despite these similarities, various motifs displayed differences in specific amino acid residues, indicating functional divergence. Such variations may account for the mechanistic differences between AmpC, which is a serine betalactamase, and IMP, which functions as a metallo-beta-lactamase (MBL) requiring metal ion coordination for hydrolysis.

#### 3.13.2 AmpC vs. NDM: Structural and Functional Conservation in Active Site Motifs

The comparison between AmpC and NDM motifs revealed strong motif conservation, particularly in regions associated with enzymatic activity. One of the most significant matches was DRAASMLS (AmpC) and KLSTALAAALMLSGC (NDM), with an E-value of 0.012, suggesting conservation in a functionally important domain. Another notable match was observed between DRAASMLS (AmpC) and MELPNIMHPVAKLSTALAAALMLSGCMPG (NDM), which exhibited an E-value of 0.038, further supporting evolutionary conservation of sequence elements.

Several functionally conserved domains were identified, including a strong similarity between LSLSDKASQHLPALKGSAFDHISVLQLGTYTAGGLPLQFPDEADSADKML (AmpC) and WVEPATAPNFGPLKVFYPGPGHTSDNITVGIDGTDIAFGGCLIKDSKAKS (NDM) (p-value = 0.001). This alignment suggested that these enzymes share common catalytic or structural features, potentially influencing substrate binding and hydrolysis mechanisms. Interestingly, many motifs aligned with minimal offset, reinforcing the idea that AmpC and NDM may have evolved from a common ancestral beta-lactamase. However, regions of sequence variability were also present, suggesting adaptations in enzymatic activity and substrate specificity that differentiate AmpC from NDM.

#### 3.13.3 AmpC vs. VIM: Evolutionary Links and Divergent Functional Adaptations

The motif comparison between AmpC and VIM beta-lactamases further highlighted conserved sequence regions, supporting a possible evolutionary link between these enzymes. A highly significant match was observed between AAAQLRAVVDAAVKPLMQQQG (AmpC) and AEDTLGLPVRAAVVT (VIM), with a p-value of 0.00079, indicating strong conservation in an enzymatic region. Another motif pair, AADRLEALVDAAVQPVMQQQD (AmpC) and AEDTLGLPVRAAVVT (VIM), also displayed considerable sequence conservation (p-value = 0.0056).

Structural and functional conservation was further supported by motif alignments in substrate-binding regions, with sequences such as IPGLAVAVTVNGKAHYFNYGVASKETGQPVSENTLFEIGSVSKTFTATLA (AmpC) and KLAKEQGYEVPNPILDELTTL (VIM) (E-value = 0.031) showing similarities in enzymatic function. Another motif alignment, IPGMAVAITHKGQRHYFNYGVASKETGQAVTEDTLFEIGSVSKTFTATLG (AmpC) vs. KLAKEQGYEVPNPILDELTTL (VIM), further confirmed structural similarity (E-value = 0.010). However, sequence variability in certain motifs, such as CALALLLIGQSAC (AmpC) and MSLLRGAVFALLVGVAGCRVSSAAPQPTA.1 (VIM) (E-value = 0.112), suggested some degree of functional divergence, potentially contributing to differences in enzymatic activity and antibiotic resistance mechanisms.

#### 3.13.4 NDM vs. IMP: High Motif Conservation and Shared Functional Features

A notable alignment between ASNGLI (NDM) and ASNKSIQPTAEASAD (IMP) (p-value = 0.004) suggested considerable conservation. However, some motif matches exhibited significantly lower p-values (e.g., 6.5E-17), highlighting regions of functional similarity that may contribute to beta-lactam hydrolysis.

#### 3.13.5 NDM vs. VIM: High Motif Conservation and Shared Functional Features

NDM and VIM exhibited some of the highest motif conservation, suggesting a strong evolutionary relationship among metallo-beta-lactamases. One of the most conserved motifs was EVFYPGAGHTMDNIVVWLPQQKILFGGCLVKSLQAKDLGNTADADLNSWP (VIM) and WVEPATAPNFGPLKVFYPGPGHTSDNITVGIDGTDIAFGGCLIKDSKAKS (NDM), with a p-value of 2.6E-15, indicating a highly conserved functional region. Similarly, a sequence alignment between ASNGLI (NDM) and FPSNGLIVETGKGLVLIDTAWGEEQTEEL (VIM) (p-value = 4.7E-5) suggested strong conservation in this domain. The strong conservation of motifs in NDM and VIM indicates that these enzymes likely share a common structural framework, which may explain their similar catalytic mechanisms. However, sequence variations in certain motifs suggest subtle functional adaptations, potentially contributing to differences in substrate affinity and antibiotic resistance profiles.

Interestingly, the ASNGLI motif of NDM was found to be conserved across both IMP (**ASN**KS**I**QPTAEASAD; pvalue = 0.004) and VIM (FP**SNGLI**VETGKGLVLIDTAWGEEQTEEL; p-value = 4.7E-5). Therefore, targeting the ASNGLI motif could help in targeting multiple MBLs, that is, NDM, VIM and IMP-all three.

Similarly, DALHAAGI, MSIQHFRVALI, FGNTKV and several other motifs were also found conserved across all three MBLs. Finally, overall analysis revealed the presence of 90 such motifs to be conserved across all three MBLs, that is, IMP, VIM and NDM. The motif sequences found to be conserved across all three MBLs have been listed in **Table S1 of Supplementary File 8.**

Finally, the final motif conservation matrix for all BLs was generated, which can be seen in **Table 10**.

**Table 10.** Similarity matrix based on motif analysis (higher values mean more conservation)

| Query ↓ /<br>Target → | Motifs<br>of<br>AmpC | Motifs<br>of IMP | Motifs<br>of<br>NDM | Motifs<br>of<br>VIM |
| --- | --- | --- | --- | --- |
| Similarity<br>to AmpC<br>motifs | 100 | 85 | 80 | 88 |
| Similarity<br>to IMP<br>motifs | 85 | 100 | 90 | 92 |
| Similarity<br>to NDM<br>motifs | 80 | 90 | 100 | 96 |
| Similarity<br>to VIM<br>motifs | 88 | 92 | 96 | 100 |

A closer look at **Table 10** reveals that motifs of AmpC, IMP and NDM are most similar to motifs of VIM than other enzymes. This further re-emphasize the fact that VIM shares maximum features from different BLs, especially the MBLs and therefore could be the evolutionary link between NDM and IMP.

### 3.14 Finding mutations of critical importance

The alignment of protein sequences as analyzed using Jalview software, can be seen in **Figure S1a of Supplementary File 9.** In the VIM protein, positions 175 to 440 are largely conserved across all species, with only minimal mutations observed within this range. Before position 175, no consistent alignment was found, with gaps occurring frequently. Similarly, gaps were also observed between positions 440 and 443. Mutations involving one or two amino acids within conserved columns are detailed in Table **S1 of Supplementary File 9**. Sequence logos generated using the WebLogo software to visualize residue conservation across all aligned blocks in our MSA can be seen in **Figure S1b of Supplementary File 9.** These logos illustrate a highly conserved domain spanning positions 175 to 440, with minimal mutations observed within this conserved region. Thus, mutations are clustered in both central and terminal regions, often affecting polar/non-polar or charged/uncharged transitions, indicative of selective pressure on functionally critical residues in the active site and structure stabilization.

Similarly, the alignment of NDM protein sequences as analyzed using Jalview software, can be seen in **Figure S2a of Supplementary File 9.** In the NDM protein, positions 30 to 271 are largely conserved across all species, with only minimal mutations observed within this range. Before position 30, no consistent alignment was found, with gaps occurring frequently. Similarly, gaps were also observed between positions 272 and 292. Mutations involving one or two amino acids within conserved columns are detailed in **Table S2 of Supplementary File 9.** Sequence logos for the same can be seen in **Figure S2b of Supplementary File 9.** These logos illustrate a highly conserved domain spanning positions 30 to 271, with minimal mutations observed within this conserved region. Thus, mutational hotspots are distributed throughout, showing frequent polar/non-polar, charged/uncharged exchanges, likely driven by selective forces for broad substrate adaptation and resistance phenotype maintenance.

The alignment of IMP protein sequences as analyzed using Jalview software, can be seen in **Figure S3a of Supplementary File 9.** In the IMP protein, positions 65 to 285 are largely conserved across all species, with only minimal mutations observed within this range. Before position 65, no consistent alignment was found, with gaps occurring frequently. Similarly, gaps were also observed between positions 286 and 300. Mutations involving one or two amino acids within conserved columns are detailed in **Table S3 of Supplementary File 9**. Sequence logos for the same have been depicted in **Figure S3b of Supplementary File 9.** These logos illustrate a highly conserved domain spanning positions 65 to 285, with minimal mutations observed within this conserved region. Thus, we can see extensive spread from early to late sequence positions with multiple changes in charge, polarity, or aromaticity, reflecting adaptation across environmental niches and possibly compensatory mutations to maintain enzyme efficacy.

Similar to the above, the alignment of AmpC protein sequences can be studied from **Figure S4a of Supplementary File 9.** Key mutated positions include 75, 82, 85, 105, 111, 135, 137, 193, 197, 243, 255, 263, 329, 359, 373, 380, 392. Thus, several conserved positions undergo substitutions in AmpC sequences, notably at aromatic, polar, and charged sites, indicating hotspots in regions related to substrate recognition or enzyme structural integrity. Mutations in AmpC involving one or two amino acids within conserved regions are detailed in **Table S4 of Supplementary File 9.** Sequence logos generated to visualize residue conservation across all aligned blocks in our MSA can be seen in **Figure S4b of Supplementary File 9.**

Upon closer examination of the mutated residues, it was found that in VIM and NDM, several non-polar and uncharged polar residues were substituted by positively charged residues like Arginine, Lysine and Histidine. On the other hand, charged residues were substituted by non-polar and uncharged polar residues in IMP and AmpC. Additionally, in VIM and NDM, the negatively charged residues like glutamate and aspartate were substituted by non-polar or positively charged residues. In contrast, in IMP, it was observed at few instances that positively charged residues were replaced by negatively charged residues. Thus, VIM and NDM undergo similar types of mutation, further indicating that VIM is closer to NDM.

### 3.15 The dual mechanism of HGT and evolution

Overall, our findings suggest a dual mechanism of HGT and evolution (both intrinsic and extrinsic). While intrinsic evolution occurs as a result of chromosomal adaptations and co-evolution with regulatory proteins (e.g., NagZ with AmpC and AmpR), extrinsic evolution encompasses plasmid-mediated HGT, allowing rapid dissemination of resistance genes across diverse bacterial genera. The dual mechanism of HGT and evolution of BLs has been visually represented in **Figure 9a-c**.

**Figure 9(a).**
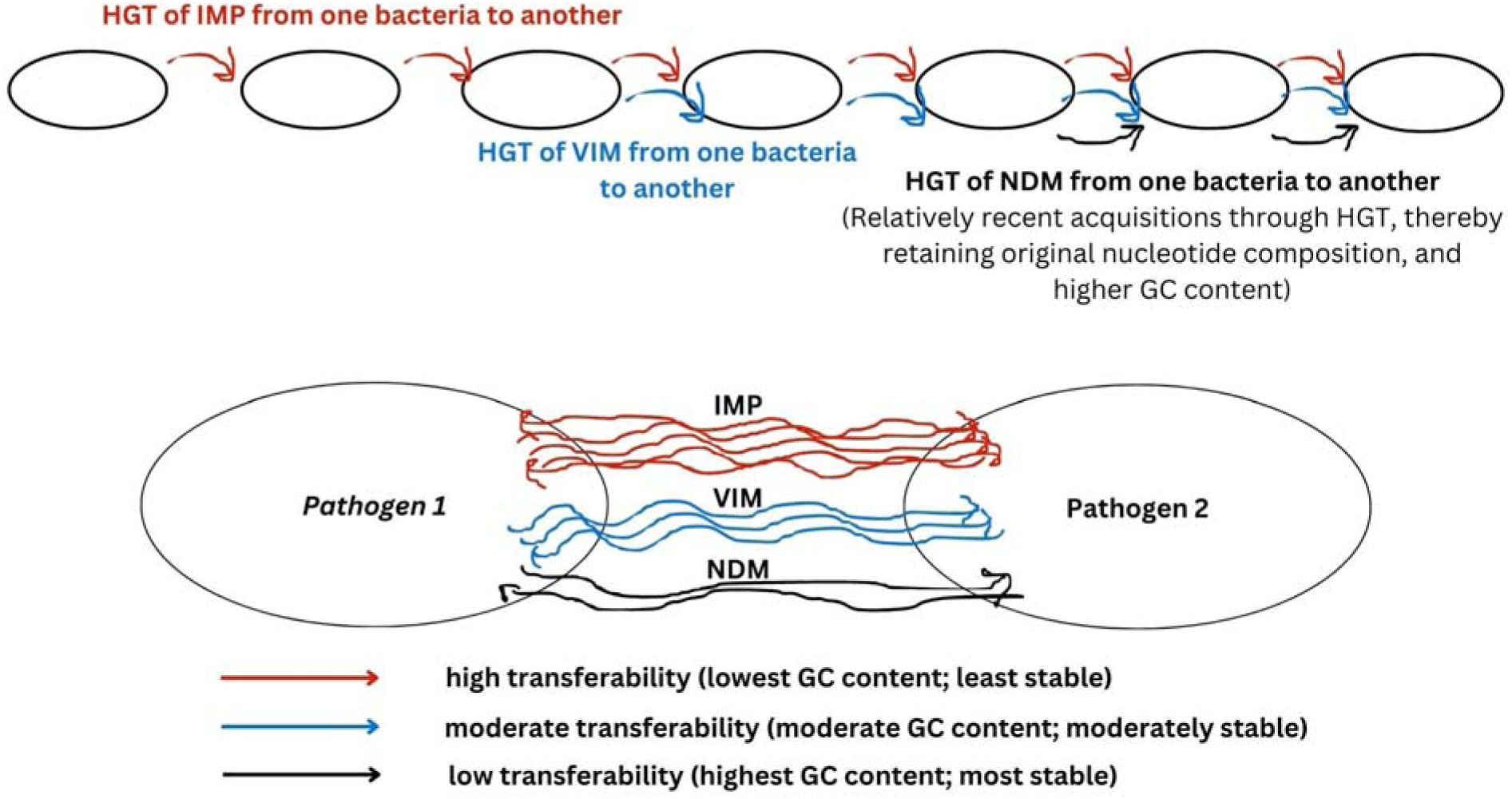
Horizontal gene transfer (HGT) of plasmid-encoded beta-lactamases from one bacteria to another.

Our findings suggest higher transferability for IMP, followed by VIM and NDM due to their increasing GC content and thereby stability, respectively (**Figure 9a).**

The **Figure 9b** shows that *Acinetobacter* and Enterobacteriaceae lie closer together in phylogenetic analysis of recent plasmid-encoded MBLs (such as NDM), similar to the chromosomally-encoded MBL-fold. On the other hand, the older MBLs like VIM and NDM have evolved such that they possess close similarity in *P. aeruginosa* and *A. baumannii*. We hereby propose a mechanism for the adaptive evolution of MBLs which has been visually presented in **Figure 9b**. Additionally, the chromosomally encoded BLs like AmpC and related proteins (AmpR and NagZ) show similar evolutionary patterns across *A. baumannii* and *P. aeruginosa* (**Figure 9c**). This clearly indicates that *Acinetobacter* is involved in both HGT (with Enterobacteria, in case of recent plasmid encoded-BLs) and ecological adaptation via mutation (similar to *Pseudomonas*, in case of chromosomally-encoded BLs), thereby highlighting the dual mechanism of HGT and evolution in *Acinteobacter*. We anticipate that similar mechanisms might be occurring in other bacteria as well. This also re-establishes the finding that there is more diversity in *Acinteobacter*, due to both HGT and chromosomal mutations, similar to the previous finding that *A. baumannii* showed highly diverse set of proteins interacting with beta-lactamases, with no common protein as a highly interacting partner for multiple BLs or related proteins.

**Figure 9(b).**
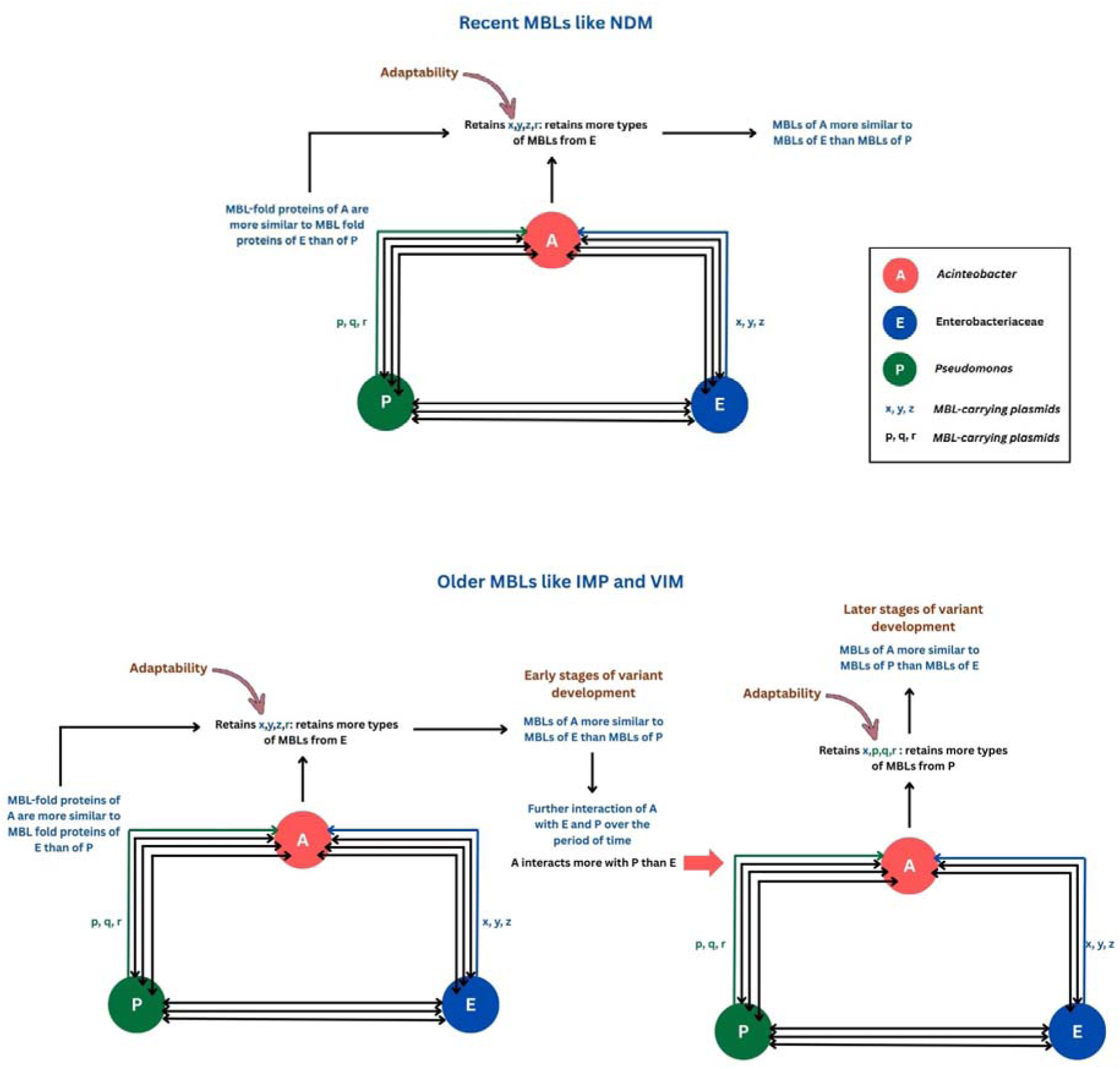
Proposed mechanism for adaptive evolution of Metallo-beta-lactamases. The figure shows adaptability and evolvability of recent MBLs vs older MBLs in Acinetobacter, Pseudomonas and Enterobacteriaceae. Phylogenetic analysis shows that recent plasmid-encoded MBLs (e.g., NDM variants) cluster more closely between Acinetobacter spp. and Enterobacteriaceae, reflecting a similarity to chromosomally encoded MBL-fold proteins. In contrast, older MBLs such as VIM and IMP lineages exhibit greater similarity between Pseudomonas aeruginosa and A. baumannii. Based on these patterns, we propose a mechanism for adaptive evolution in MBLs, highlighting shifts in host association and structural similarity over evolutionary time. The model visually depicts how horizontal gene transfer, environmental pressures (closer association of bacteria with each other), and structural adaptability (similarity of MBL-fold proteins) contribute to the observed phylogenetic relationships.

**Figure 9(c).**
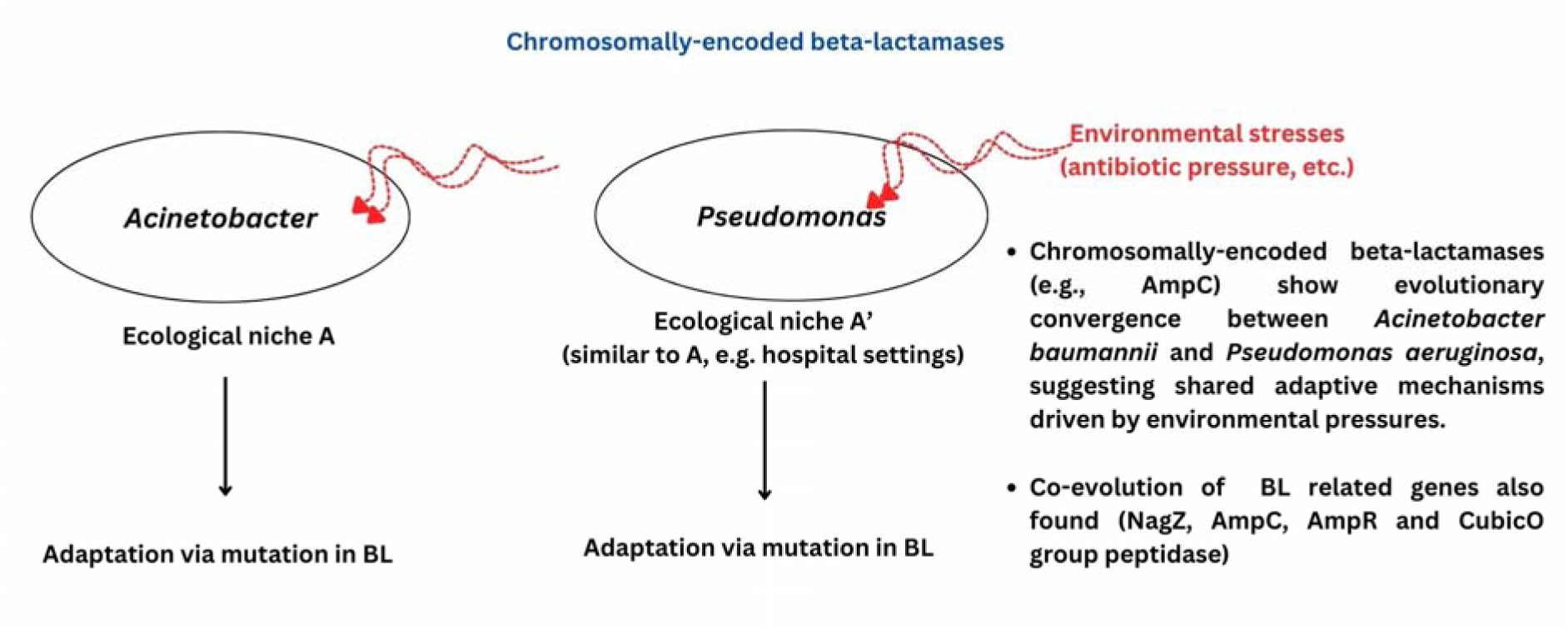
Plasmid-encoded BLs and chromosomally-encoded BLs show evolutionary trends different from each other.

#### 3.15.1 Extrinsic Evolution

##### 3.15.1.1 Plasmid-Mediated HGT

Thus, the study highlights the significant role of plasmid-encoded metallo-beta-lactamases (MBLs) such as NDM, VIM, and IMP in facilitating HGT. Plasmids seem to enable the rapid transfer of resistance genes between *Acinetobacter* and Enterobacteriaceae, enhancing the spread of carbapenem resistance. The increased similarity of plasmid-encoded MBLs between these genera indicates a higher rate of plasmid exchange, which supports the dissemination of antibiotic resistance traits through mobile genetic elements like integrons and transposons (**Noel et al., 2022; Yao et al., 2024**).

##### 3.15.1.2 GC content, transferability, and evolutionary flexibility of BLs

Genes with lower GC content are often more prone to mutations and recombination events. This genomic plasticity can facilitate integration into diverse bacterial genomes, enhancing the likelihood of horizontal transfer **(Hayek, 2013).** The differential GC content of MBLs (e.g., low GC in IMP and high GC in NDM) indicates distinct evolutionary origins and possibly varied pathways of HGT. The stability of GC content across different species in case of NDM suggests that these genes retain their original nucleotide composition when transferred to new hosts, supporting the notion of relatively recent acquisitions through HGT.

The study indicates that lower GC content promotes enzyme variability, as seen with IMP and VIM, while higher GC content like in NDM provides stability. This flexibility could mean that IMP is not only transferred more frequently but also adapts more readily to new host environments through mutations **(Michiels et al., 2016; Mori et al., 2021).** Additionally, in environments with strong antibiotic pressure, genes with lower GC content might accumulate mutations that help in evolving resistance, further promoting their transfer among bacteria **(Hayek, 2013).** Thus, the lower GC content of IMP it not just implies higher HGT rates (easier incorporation into diverse bacterial genomes), but also broader host range (facilitating the spread of carbapenem resistance across different genera), and adaptive potential (through increased mutational flexibility, IMP might evolve new resistance traits more readily than NDM or VIM). Finally, the lower GC content of IMP reflects its older evolutionary origin or a long period of time since when it has been spreading and adapting, and thus showing variability in GC content.

#### 3.15.2 Intrinsic Evolution

##### 3.15.2.1 Chromosomal vs. Plasmid Evolution

Chromosomally-encoded beta-lactamases (e.g., AmpC) show evolutionary convergence between *Acinetobacter baumannii* and *Pseudomonas aeruginosa*, suggesting shared adaptive mechanisms driven by environmental pressures **(Lister, 2002).** In contrast, plasmid-encoded MBLs like NDM show higher similarity between *Acinetobacter* and Enterobacteriaceae, highlighting the dynamic role of HGT in these bacteria **(Da Silva et al., 2016)**.

##### 3.15.2.2 Co-Evolutionary Dynamics

The analysis of NagZ and its associated beta-lactamase-related proteins (e.g., AmpC, AmpR) demonstrates coevolutionary trends. Phylogenetic congruence suggests functional interdependence, possibly reflecting concurrent adaptation to antibiotic pressures. The lack of phylogenetic congruence with proteins like glyoxylase and AmpD indicates independent evolutionary paths, hinting at selective pressures specific to certain proteins.

##### 3.15.2.3 Similar adaptive mechanisms in *Acinetobacter baumannii* and *Pseudomonas aeruginosa*, and biofilm formation

*Acinetobacter baumannii* and *Pseudomonas aeruginosa* are generally known to form more robust and mature biofilms compared to many members of the Enterobacteriaceae family **(Gedefie et al., 2021; Papa et al., 2018).** This might be linked to their close proximity in phylogenetic tree for NagZ, AmpC and AmpR, which are seen to co-evolve. *Pseudomonas aeruginosa* uses a complex regulatory network, including quorum sensing (las, rhl, and pqs systems) and cyclic-di-GMP signaling, to form thick, structured biofilms. It produces extracellular polymeric substances (EPS), including alginate, which contributes to the biofilm matrix **(De Kievit et al., 2001).** *Acinetobacter baumannii* forms biofilms on abiotic surfaces, often associated with hospital environments. The Bap (biofilm-associated protein) and Csu pili play critical roles in biofilm initiation and maturation **(Tomaras et al., 2003).** Many members of the Enterobacteriaceae family, like Escherichia coli, *Klebsiella pneumoniae*, and *Proteus mirabilis*, can also form biofilms. However, they typically do not produce biofilms as mature and resilient as *P. aeruginosa* and *A. baumannii*, especially in clinical and environmental contexts **(Ezzariga et al., 2025)**. E. coli can form biofilms, particularly in the case of uropathogenic strains (UPEC), but the structure and resistance of these biofilms vary widely among strains **(Ezzariga et al., 2025; Soto et al., 2007).** As compared to Enterobacteriaceae, both *P. aeruginosa* and *A. baumannii* excel on abiotic surfaces, giving them an advantage in healthcare settings (e.g., catheters, ventilators) **(Abdallah et al., 2014; Anurag Anand et al., 2025c).** The EPS produced by *P. aeruginosa* and *A. baumannii* often have stronger adhesive and protective properties. Biofilms of *P. aeruginosa* and *A. baumannii* are highly resistant to antibiotics and host immune responses, which is a major clinical challenge **(Gedefie et al., 2021; Papa et al., 2018)**. Thus, while Enterobacteriaceae can form biofilms, *P. aeruginosa* and *A. baumannii* generally form more robust and clinically challenging biofilms. The similar ecological niche or environmental stresses associated with both these bacteria may be the reason they show evolutionary convergence in case of chromosomally encoded BLs like AmpC and related genes (NagZ and AmpR). Overall, there is a possibility that *Pseudomonas and Acinetobacter* might be sharing other resistance patterns as well.

##### 3.15.2.4 Evidence from Motif Conservation and Mutational Hotspots

The presence of conserved motifs and mutational hotspots within beta-lactamases, as shown by MEME and TOMTOM analyses, suggests an evolutionary strategy where critical functional domains are maintained while allowing mutations at non-essential sites (**Table 10**). This could contribute to adaptive resistance while preserving enzymatic functionality.

**Figure 10(a)** highlights the mechanism of co-evolution as a result of functional inter-dependence, whereas **Figure 10(b)** highlights the adaptive nature of motifs and domains in response to antibiotic pressure.

**Figure 10(a).**
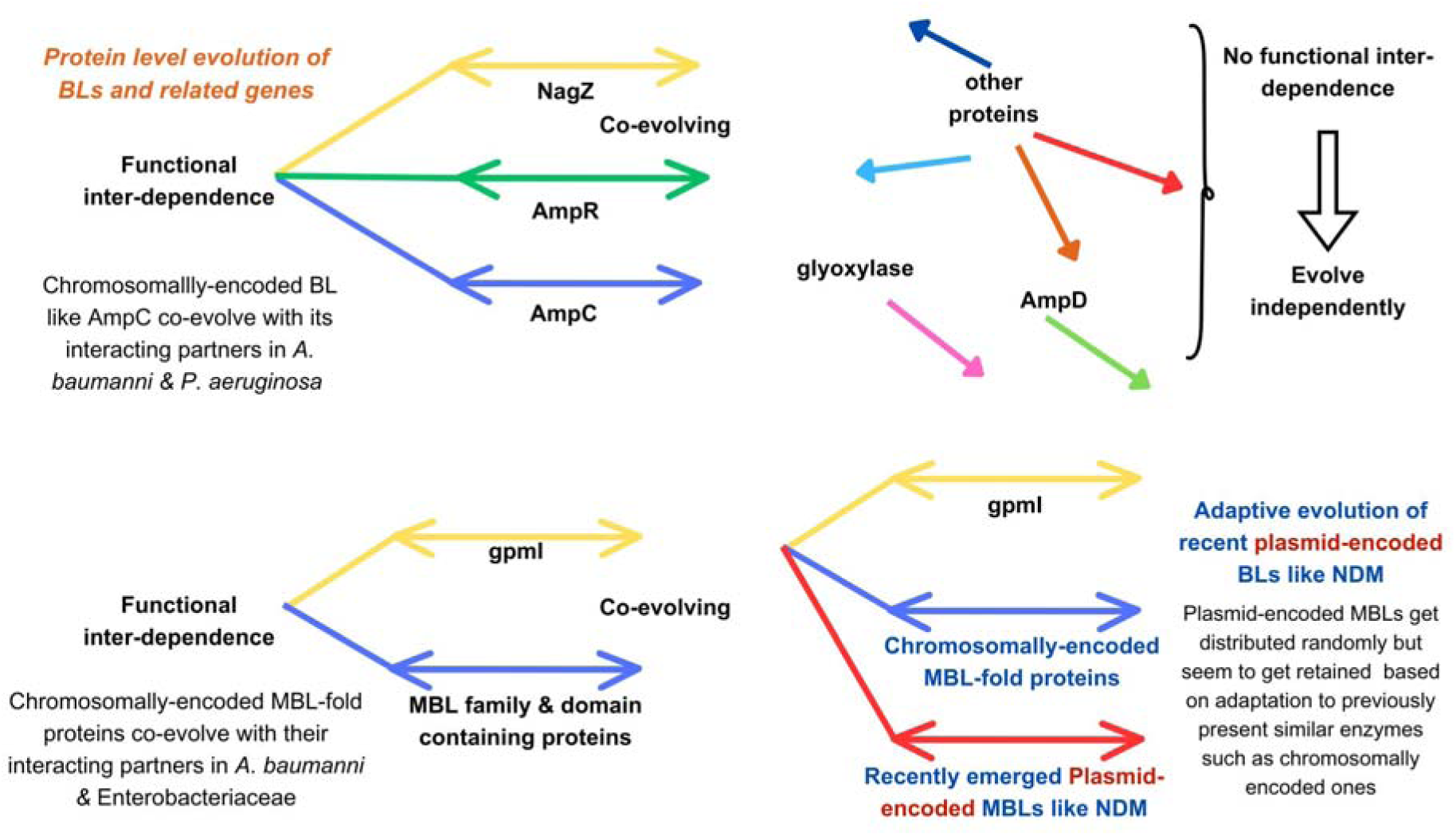
Emergence of co-evolution as a result of functional inter-dependence.

**Figure 10(b).**
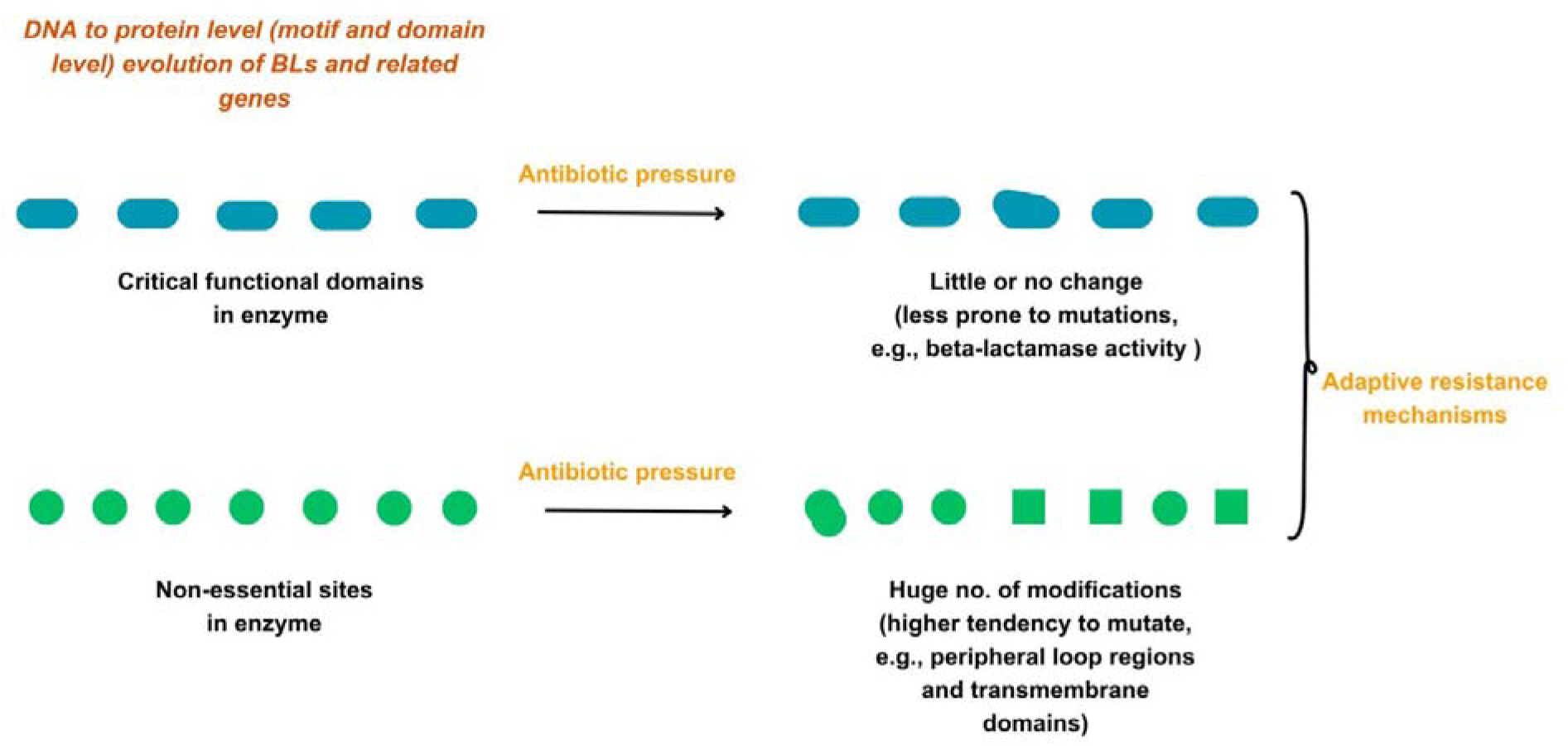
The adaptive nature of motifs and domains in response to antibiotic pressure.

### 3.16 VIM as an evolutionary link between IMP and NDM

The variety of shared characteristics between VIM and IMP and VIM and NDM is indicative of VIM acting as an evolutionary link between the other two. Phylogenetic trees and PaSiMap analysis show that VIM lies between NDM and IMP. GC content analysis showed VIM to have average GC content between NDM and IMP. Comparative analysis of AAC, ABC, and APP using Principal Component Analysis (PCA) reveals distinct clustering patterns among NDM, VIM, IMP, and AmpC beta-lactamases, but with similar positive contributors between VIM and NDM, and between VIM and IMP. Further, PFAM domains and motifs are conserved maximum in NDM but least in IMP, with VIM lying in between them, suggesting a strong evolutionary link. Thus, VIM seems to be acting as a mosaic, having acquired characteristics from both NDM and IMP via HGT, recombination or other mechanisms **(Queremel and Chauhan, 2025; Lister, 2002)**.

## 4 Conclusion and future perspectives

This study presents an integrated analysis of the co-evolutionary dynamics, structural stability, and functional divergence of beta-lactamases (BLs) and their associated proteins in priority pathogens, mainly, *Pseudomonas aeruginosa*, *Acinetobacter baumannii*, and Enterobacteriaceae. By combining phylogenetic, statistical, and sequence analyses with STRING-based protein-protein interaction (PPI) networks, we uncovered key insights into the molecular mechanisms driving beta-lactam resistance. NagZ emerges as the primary interacting partner for BLs in *P. aeruginosa* and Enterobacteriaceae, highlighting its critical role in the regulatory cascade leading to beta-lactamase expression. The strong co-evolutionary trends between NagZ, AmpC, and AmpR, emphasize functional interdependence and shared evolutionary pressures among these proteins. In contrast, *A. baumannii* exhibits a diverse range of interactions, indicating a more complex and varied resistance mechanism. Plasmid-encoded MBLs (e.g., NDM, VIM, IMP) display higher horizontal gene transfer (HGT) rates, with diversity of bacterial interactions. Study of recent MBLs NDM-type shows that high HGT occurs between *Acinetobacter* and Enterobacteriaceae, contributing to the widespread dissemination of carbapenem resistance across these organisms. However, older MBLs show closer association between *Pseudomonas* and *Acinetobacter.* Further, chromosomally encoded BLs (AmpC) exhibited closer evolutionary relationships between *Pseudomonas* and *Acinetobacter*. Lower GC content, as seen in IMP, correlates with higher mutational plasticity, while higher GC content, as in NDM, is associated with structural stability. VIM showed intermediate GC content. Principal Component Analysis (PCA) of amino acid composition and physicochemical properties demonstrate similar positive contributors between VIM and NDM, and between VIM and IMP. Finally, inter-BL analysis revealed that VIM can be understood as an evolutionary link between IMP and NDM due to presence of features shared from NDM as well as IMP, such as GC content, PFAM domains and structural and functional motifs.

In the light of these findings, we propose several therapeutic strategies. Given the strong co-evolutionary link between NagZ, AmpC, and AmpR, designing small-molecule inhibitors targeting NagZ could suppress the regulatory cascade leading to beta-lactamase expression. This offers a novel therapeutic strategy to combat resistance in *Pseudomonas aeruginosa* and Enterobacteriaceae. The high adaptability observed for recent plasmid-encoded MBLs like NDM across *Acinetobacter* and Enterobacteriaceae, highlight the need for targeted interventions to curb the spread of resistance genes across these pathogens. Enzymes with lower GC content, such as IMP, exhibit higher mutational plasticity, making them more susceptible to structural disruptions. This insight can guide the development of drugs targeting these vulnerabilities, particularly in highly mutable BLs. Given the distinct evolutionary bifurcation patterns observed in BLs, CRISPR-based strategies could selectively disable plasmid-encoded MBLs while preserving essential chromosomally encoded functions. Since *Acinetobacter baumannii* exhibits high genetic plasticity, targeted genomic surveillance can track emerging mutations and guide adaptive treatment strategies. The finding that VIM serves as a mosaic linking IMP and NDM, can help to better understand the evolution and emergence of new beta-lactamase variants. Moreover, there is a high possibility that inhibitors targeting VIM can also inhibit IMP and NDM as VIM shares features of both NDM and IMP, while IMP and NDM lie farther apart and share fewer common features. Additionally, targeting components conserved across VIM, NDM and IMP (such as the ASNGLI motif) can help in specific targeting of multiple betalactamases at a time. Lastly, findings such as the selective conservation of VIM in *Pseudomonas* and Enterobacteriaceae, with its strikingly low prevalence in *A. baumannii*, suggesting a differential distribution and reliance on distinct carbapenemase types across these critical pathogens indicate that there is much more to explore in BL evolution.

## 5. Data availability

Related data can be found in the supplementary files. Any additional data shall be made available on request to the corresponding author.

## 7. Acknowledgements

Ananya Anurag Anand, Kshitiz K and Sreedevi PC are thankful to Ministry of Education (MoE), Govt. of India, for their fellowship. Sarfraz Anwar is thankful to CSIR-UGC (Council of Scientific and Industrial Research - University Grants Commission) for his fellowship. Vidushi Yadav is thankful to CST-UP (Council of Science and Technology, Uttar Pradesh) for her fellowship. All the authors are thankful to IIIT Allahabad for providing the research facility.

## 8. Author contributions

Ananya Anurag Anand: Conceptualization, Methodology, Writing - original draft, review and editing; Vidushi Yadav: Methodology; Kshitiz K: Methodology; Sreedevi PC: Methodology; Sarfraz Anwar: Methodology; Sintu Kumar Samanta: Supervision and Conceptualization, Investigation, Resources, Writing - review and editing.

## 9. Corresponding author

Correspondence to Dr. Sintu Kumar Samanta

## References

Abdallah M, Benoliel C, Drider D, et al (2014) Biofilm formation and persistence on abiotic surfaces in the context of food and medical environments. Arch Microbiol 196:453–472. 10.1007/s00203-014-0983-1

Amod A, Singh S, Lawrence R, et al (2025) NCCD: A unique combination of MOF-scaffolded stable luminescent copper nanoclusters (Cu NCs) and carbon dots (Cdots) as a potent antimicrobial agent against Pseudomonas aeruginosa. Nanotechnology. 10.1088/1361-6528/addacd

Anand AA, Anwar S, Mondal RK, Samanta SK (2025a) Influence of F repeats and terminal residue polarity on peptide– VIM-2 metallo-β-lactamase interactions. bioRxiv

Anand AA, Yadav V, Anwar S, et al (2025b) The importance of phylogenetic studies in the evolution of betalactamases: Tracing antimicrobial resistance in priority pathogens. Gene Protein Dis 4:8278. 10.36922/gpd.8278

Anurag Anand A, Anwar S, Sahoo AK, Samanta SK (2025) In silico identification of Corylifol C as a potential natural inhibitor of BfrB-Bfd interaction in Pseudomonas aeruginosa. J Biomol Struct Dyn 1–15. 10.1080/07391102.2025.2472171

Anurag Anand A, Sahoo AK, Samanta SK (2024) Exploring the potential of designed peptides containing lysine and arginine repeats against VIM-2 metallo-beta-lactamases. Int J Pept Res Ther 30:. 10.1007/s10989-024-10619-5

Arer V, Anurag Anand A, Samanta SK, Kar D (2024) Deeper insights into the action of class C β-lactamases against cephalosporins through molecular docking and MD simulation studies. Biologia (Bratisl) 79:3695–3709. 10.1007/s11756-024-01802-6

Arer V, Kar D (2022) Biochemical exploration of β-lactamase inhibitors. Front Genet 13:1060736. 10.3389/fgene.2022.1060736

Bedhomme S, Perez Pantoja D, Bravo IG (2017) Plasmid and clonal interference during post horizontal gene transfer evolution. Mol Ecol 26:1832–1847. 10.1111/mec.14056

Bersani M, Failla M, Vascon F, et al (2023) Structure-based optimization of 1,2,4-triazole-3-thione derivatives: Improving inhibition of NDM-/VIM-type metallo-β-lactamases and synergistic activity on resistant bacteria. Pharmaceuticals (Basel) 16:1682. 10.3390/ph16121682

Brandt K, Barrangou R (2016) Phylogenetic analysis of the Bifidobacterium genus using glycolysis enzyme sequences. Front Microbiol 7:657. 10.3389/fmicb.2016.00657

Bush K, Bradford PA (2016) Β-lactams and β-lactamase inhibitors: An overview. Cold Spring Harb Perspect Med 6:. 10.1101/cshperspect.a025247

Bush K, Jacoby GA (2010) Updated functional classification of beta-lactamases. Antimicrob Agents Chemother 54:969–976. 10.1128/AAC.01009-09

Camacho C, Coulouris G, Avagyan V, et al (2009) BLAST+: architecture and applications. BMC Bioinformatics 10:421. 10.1186/1471-2105-10-421

Crooks GE, Hon G, Chandonia J-M, Brenner SE (2004) WebLogo: a sequence logo generator. Genome Res 14:1188– 1190. 10.1101/gr.849004

Da Silva GJ, Domingues S (2016) Insights on the horizontal gene transfer of carbapenemase determinants in the opportunistic pathogen Acinetobacter baumannii. Microorganisms 4:29. 10.3390/microorganisms4030029

De Kievit TR, Gillis R, Marx S, et al (2001) Quorum-sensing genes in Pseudomonas aeruginosa biofilms: their role and expression patterns. Appl Environ Microbiol 67:1865–1873. 10.1128/AEM.67.4.1865-1873.2001

Edgar RC (2004) MUSCLE: multiple sequence alignment with high accuracy and high throughput. Nucleic Acids Res 32:1792–1797. 10.1093/nar/gkh340

Ezzariga N, Zouhari O, Rhars A, et al (2025) Biofilm and antibiotic resistance study of bacteria involved in nosocomial infections. Cureus 17:e78673. 10.7759/cureus.78673

Felsenstein J (1981) Evolutionary trees from DNA sequences: a maximum likelihood approach. J Mol Evol 17:368–376. 10.1007/bf01734359

Finn RD, Coggill P, Eberhardt RY, et al (2016) The Pfam protein families database: towards a more sustainable future. Nucleic Acids Res 44:D279–85. 10.1093/nar/gkv1344

Gedefie A, Demsis W, Ashagrie M, et al (2021) Acinetobacter baumannii biofilm formation and its role in disease pathogenesis: A review. Infect Drug Resist 14:3711–3719. 10.2147/IDR.S332051

Hayek N (2013) Lateral transfer and GC content of bacterial resistant genes. Front Microbiol 4:41. 10.3389/fmicb.2013.00041

Imkamp F, Kolesnik-Goldmann N, Bodendoerfer E, et al (2022) Detection of extended-spectrum β-lactamases (ESBLs) and AmpC in class A and class B carbapenemase-producing Enterobacterales. Microbiol Spectr 10:e0213722. 10.1128/spectrum.02137-22

Kumar S, Nei M, Dudley J, Tamura K (2008) MEGA: a biologist-centric software for evolutionary analysis of DNA and protein sequences. Brief Bioinform 9:299–306. 10.1093/bib/bbn017

Lister PD (2002) Chromosomally-encoded resistance mechanisms of Pseudomonas aeruginosa: therapeutic implications. Am J Pharmacogenomics 2:235–243. 10.2165/00129785-200202040-00003

Liu G, Thomsen LE, Olsen JE (2022) Antimicrobial-induced horizontal transfer of antimicrobial resistance genes in bacteria: a mini-review. J Antimicrob Chemother 77:556–567. 10.1093/jac/dkab450

Mei H, Arbeithuber B, Cremona MA, et al (2019) A high-resolution view of adaptive event dynamics in a Plasmid. Genome Biol Evol 11:3022–3034. 10.1093/gbe/evz197

Michiels JE, Van den Bergh B, Fauvart M, Michiels J (2016) Draft genome sequence of Acinetobacter baumannii strain NCTC 13423, a multidrug-resistant clinical isolate. Stand Genomic Sci 11:57. 10.1186/s40793-016-0181-7

Mori N, Tada T, Oshiro S, et al (2021) A transferrable IncL/M plasmid harboring a gene encoding IMP-1 metallo-β-lactamase in clinical isolates of Enterobacteriaceae. BMC Infect Dis 21:1061. 10.1186/s12879-021-06758-5

Noel HR, Petrey JR, Palmer LD (2022) Mobile genetic elements in Acinetobacter antibiotic-resistance acquisition and dissemination. Ann N Y Acad Sci 1518:166–182. 10.1111/nyas.14918

Nystrom SL, McKay DJ (2021) Memes: A motif analysis environment in R using tools from the MEME Suite. PLoS Comput Biol 17:e1008991. 10.1371/journal.pcbi.1008991

Pande A, Patiyal S, Lathwal A, et al (2023) Pfeature: A tool for computing wide range of protein features and building prediction models. J Comput Biol 30:204–222. 10.1089/cmb.2022.0241

Papa R, Bado I, Iribarnegaray V, et al (2018) Biofilm formation in carbapenemase-producing Pseudomonas spp. and Acinetobacter baumannii clinical isolates. Int J Infect Dis 73:119–120. 10.1016/j.ijid.2018.04.3688

Queremel Milani DA, Chauhan PR (2025) Genetics, mosaicism. In: StatPearls. StatPearls Publishing, Treasure Island (FL)

Russo CA de M, Selvatti AP (2018) Bootstrap and rogue identification tests for phylogenetic analyses. Mol Biol Evol 35:2327–2333. 10.1093/molbev/msy118

Salahuddin P, Khan AU (2014) Studies on structure-based sequence alignment and phylogenies of beta-lactamases. Bioinformation 10:308–313. 10.6026/97320630010308

Schotte U, Ehlers J, Nieter J, et al (2024) ESBL-type and AmpC-type beta-lactamases in third generation cephalosporin-resistant Enterobacterales isolated from animal feces in Madagascar. Animals (Basel) 14:741. 10.3390/ani14050741

Sokal R, Michener C (1958) A statistical method for evaluating systematic relationships. University of Kansas science bulletin 38:1409–1438

Soto SM, Smithson A, Martinez JA, et al (2007) Biofilm formation in uropathogenic Escherichia coli strains: relationship with prostatitis, urovirulence factors and antimicrobial resistance. J Urol 177:365–368. 10.1016/j.juro.2006.08.081

Szklarczyk D, Nastou K, Koutrouli M, et al (2025) The STRING database in 2025: protein networks with directionality of regulation. Nucleic Acids Res 53:D730–D737. 10.1093/nar/gkae1113

Tamura K, Stecher G, Kumar S (2021) MEGA11: Molecular Evolutionary Genetics Analysis version 11. Mol Biol Evol 38:3022–3027. 10.1093/molbev/msab120

Tomaras AP, Dorsey CW, Edelmann RE, Actis LA (2003) Attachment to and biofilm formation on abiotic surfaces by Acinetobacter baumannii: involvement of a novel chaperone-usher pili assembly system. Microbiology 149:3473–3484. 10.1099/mic.0.26541-0

Truty R, Rojahn S, Ouyang K, et al (2023) Patterns of mosaicism for sequence and copy-number variants discovered through clinical deep sequencing of disease-related genes in one million individuals. Am J Hum Genet 110:551–564. 10.1016/j.ajhg.2023.02.013

Waterhouse AM, Procter JB, Martin DMA, et al (2009) Jalview Version 2--a multiple sequence alignment editor and analysis workbench. Bioinformatics 25:1189–1191. 10.1093/bioinformatics/btp033

Yao Y, Imirzalioglu C, Falgenhauer L, et al (2024) Plasmid-mediated spread of carbapenem resistance in Enterobacterales: A three-year genome-based survey. Antibiotics (Basel) 13:682. 10.3390/antibiotics13080682

WHO publishes list of bacteria for which new antibiotics are urgently needed. In: Who.int. https://www.who.int/news/item/27-02-2017-who-publishes-list-of-bacteria-for-which-new-antibiotics-are-urgently-needed. Accessed 16 Dec 2025a

(2024) WHO bacterial priority pathogens list, 2024: Bacterial pathogens of public health importance to guide research, development and strategies to prevent and control antimicrobial resistance. In: Who.int. https://www.who.int/publications/i/item/9789240093461. Accessed 16 Dec 2025

