## Supplementary File 4 for "Phylogenetic Analysis of Beta-Lactamases Reveals Distinct Evolutionary Patterns of Chromosomal and Plasmid-Encoded BLs and the Mosaic Role of VIM Linking NDM and IMP"

### Phylogenetic tree of AmpC showing proximity between Acinetobacter, Pseudomonas and Enterobacteriaceae

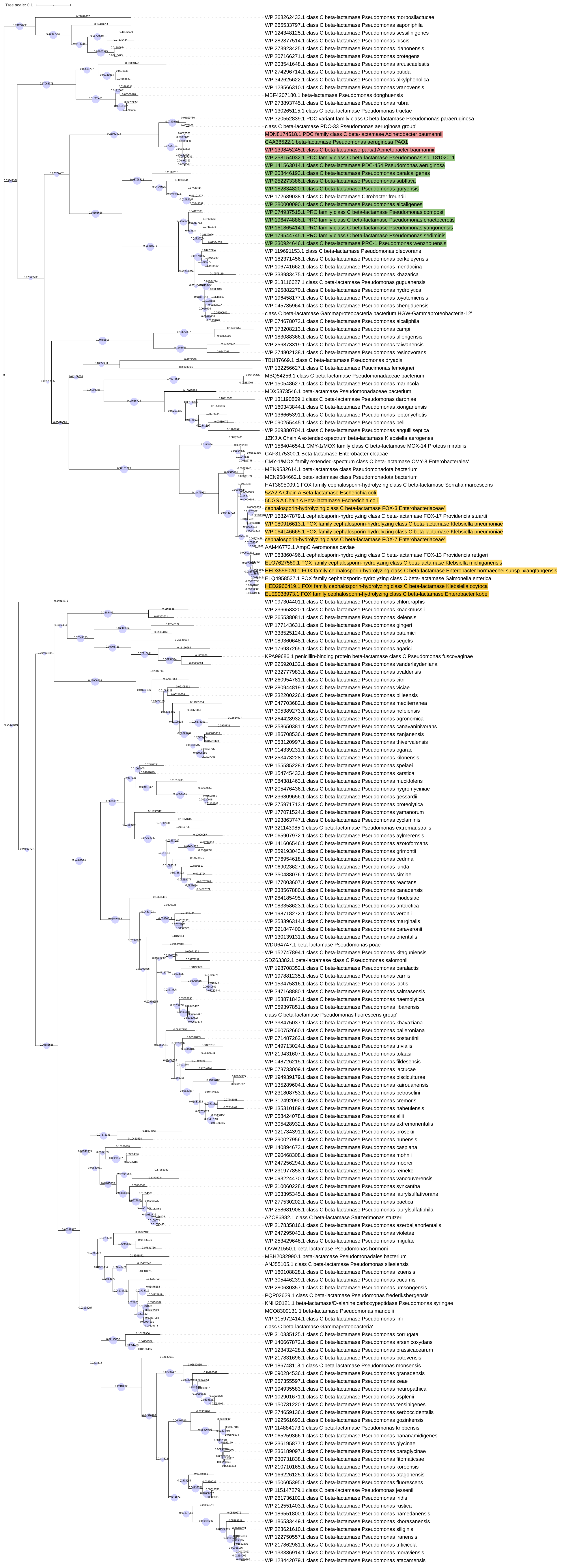

Phylogenetic tree of AmpR

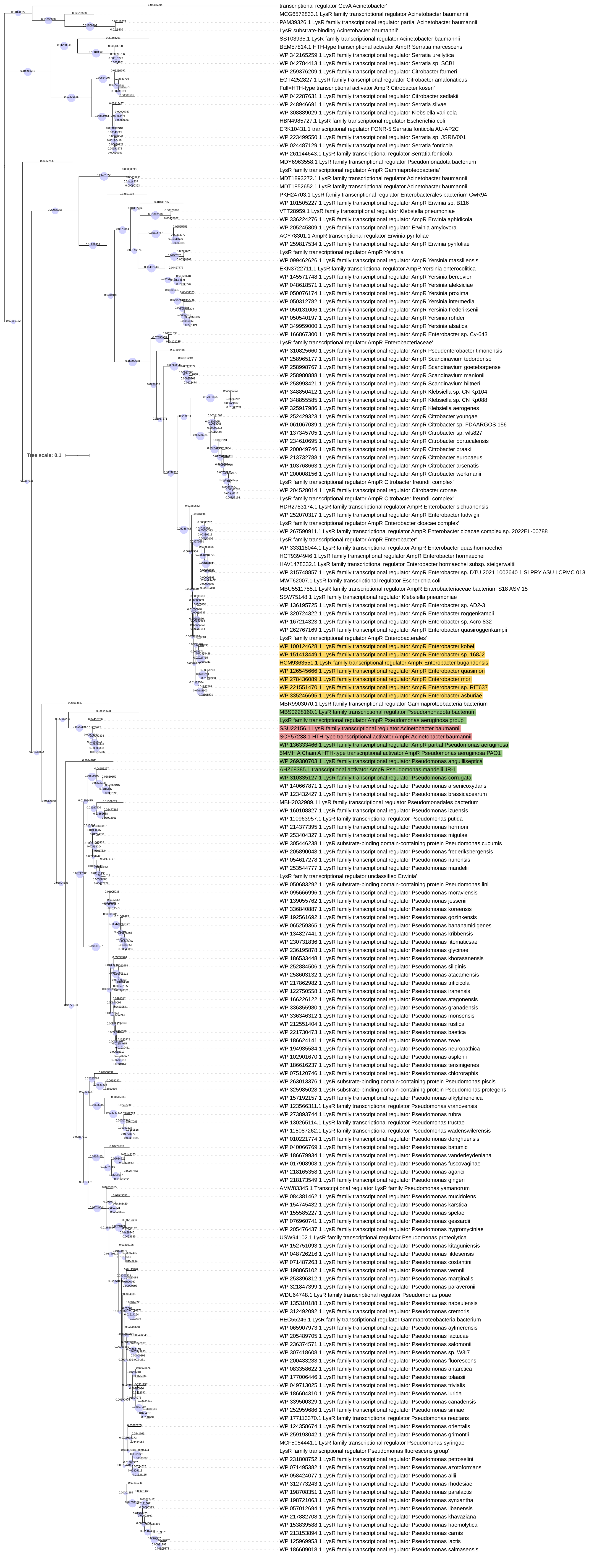

Phylogenetic tree of NagZ

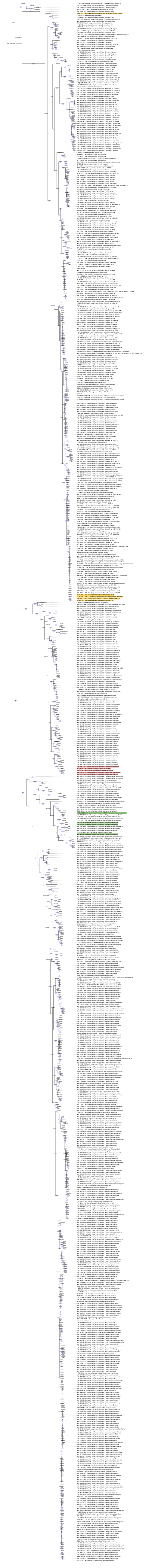

Phylogenetic tree of glyoxylase showing proximity between Acinetobacter, Pseudomonas and Enterobacteriaceae

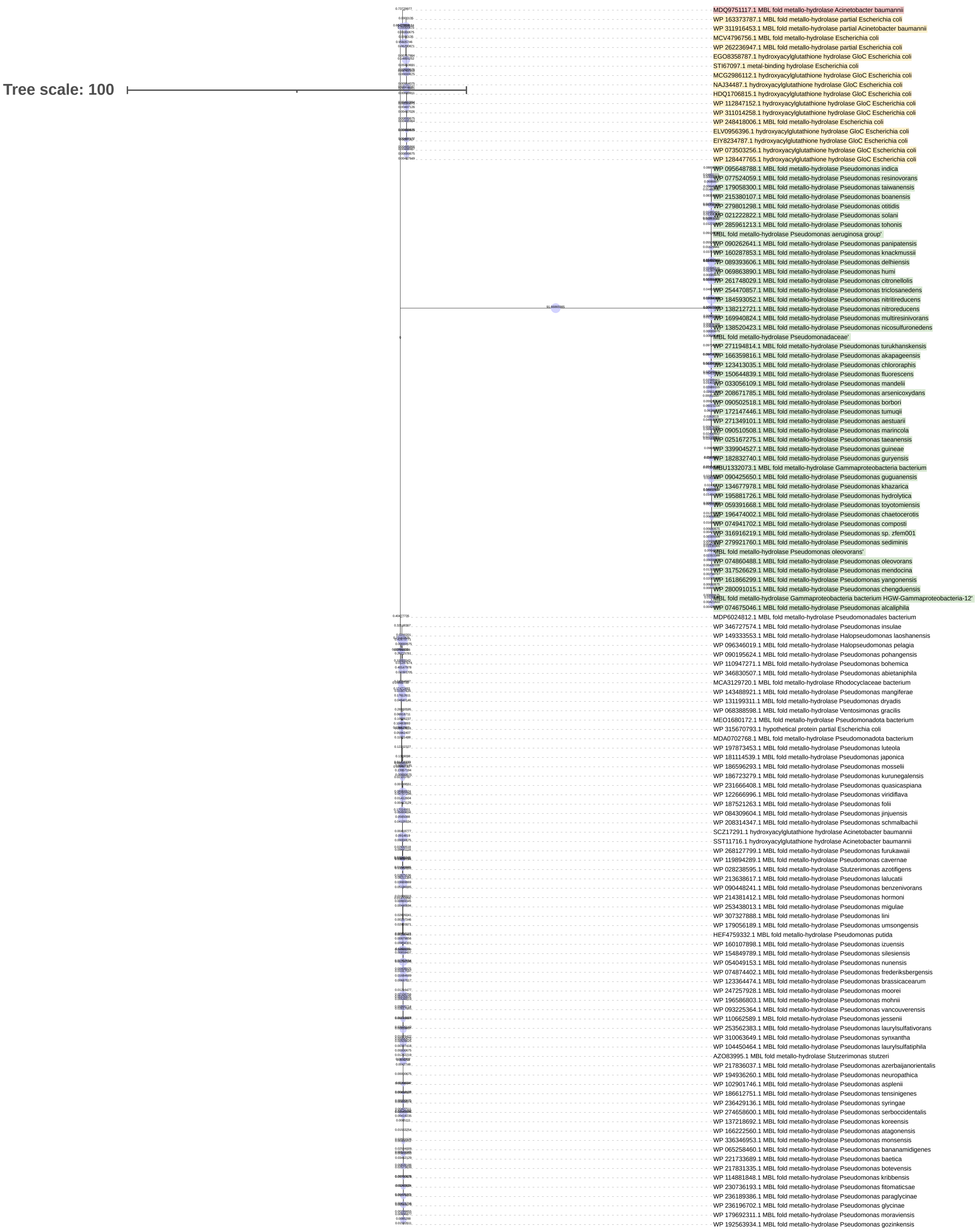

Phylogenetic tree of gua showing proximity between Acinetobacter, Pseudomonas and Enterobacteriaceae

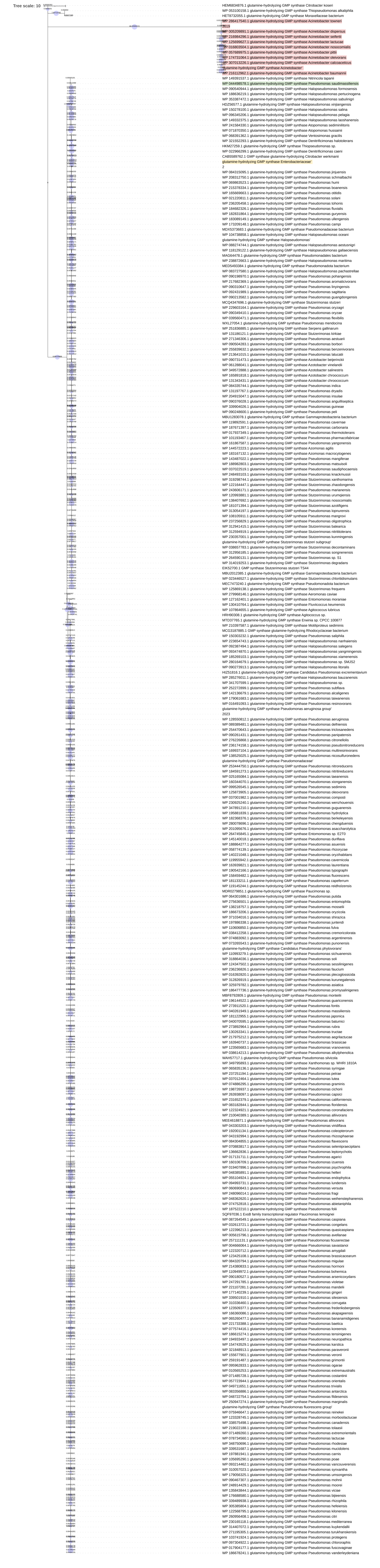

Phylogenetic tree of MDS_1318 showing proximity between Acinetobacter, Pseudomonas and Enterobacteriaceae

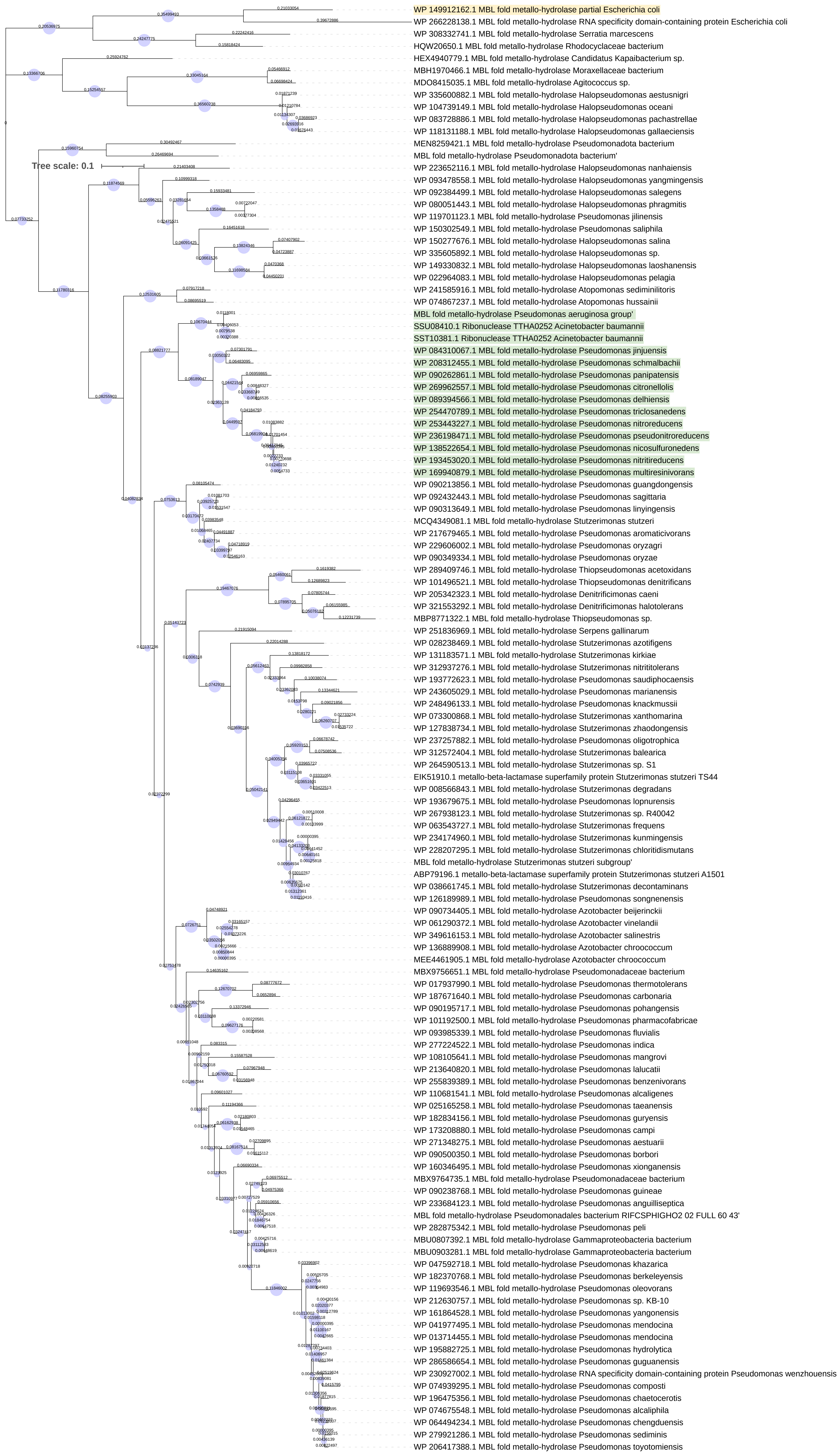

Tree scale: 0.1

Phylogenetic tree of MFS-transporter showing proximity between Acinetobacter, Pseudomonas and Enterobacteriaceae

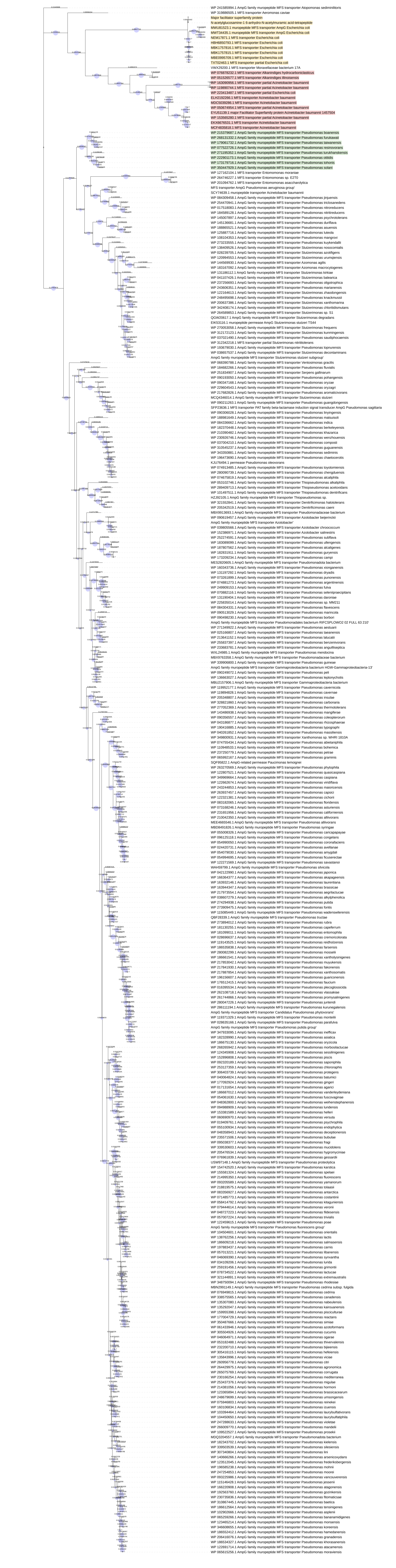

Phylogenetic tree of polA showing proximity between Acinetobacter, Pseudomonas and Enterobacteriaceae

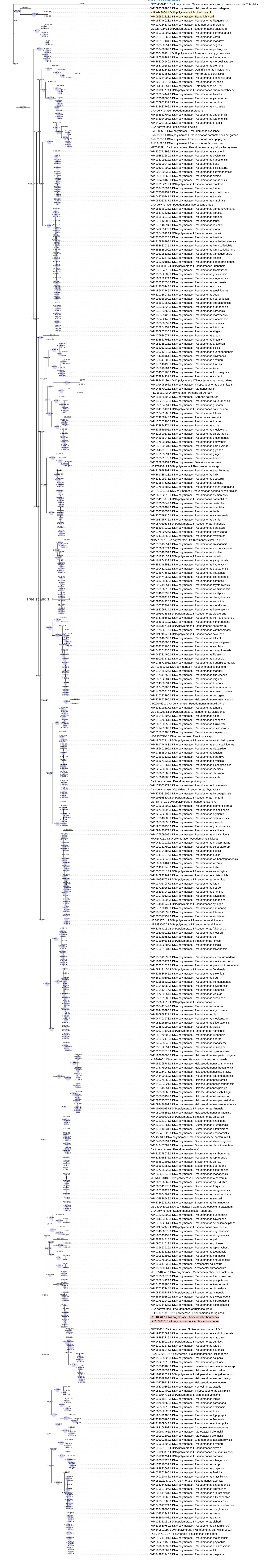

Phylogenetic tree of rpoB showing proximity between Acinetobacter, Pseudomonas and Enterobacteriaceae

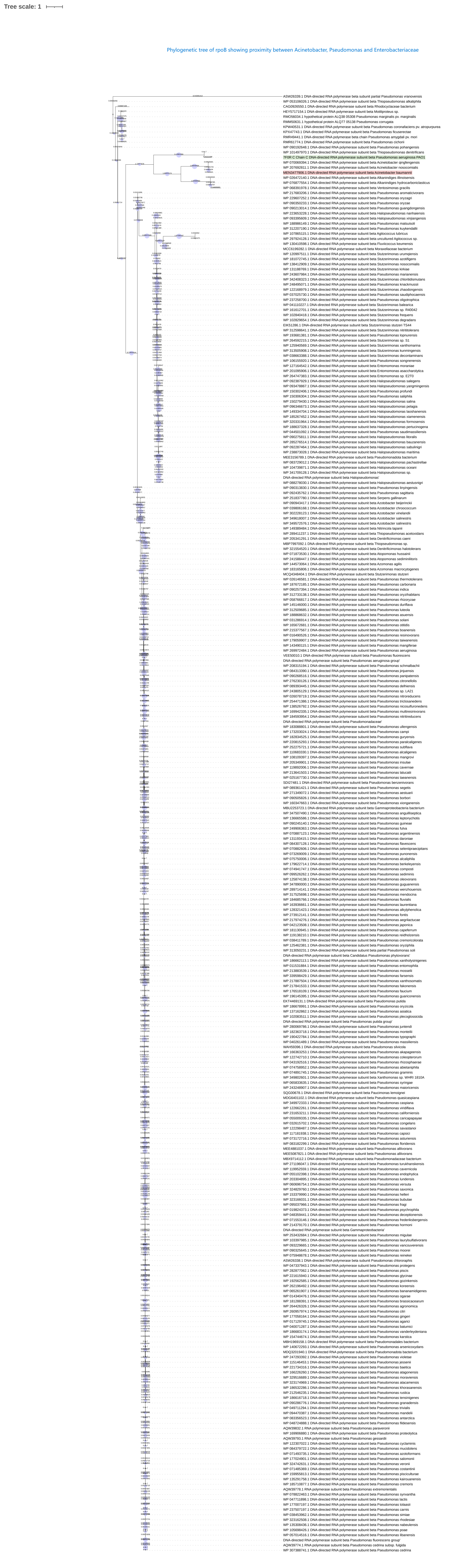

### Phylogenetic tree of SDI51743

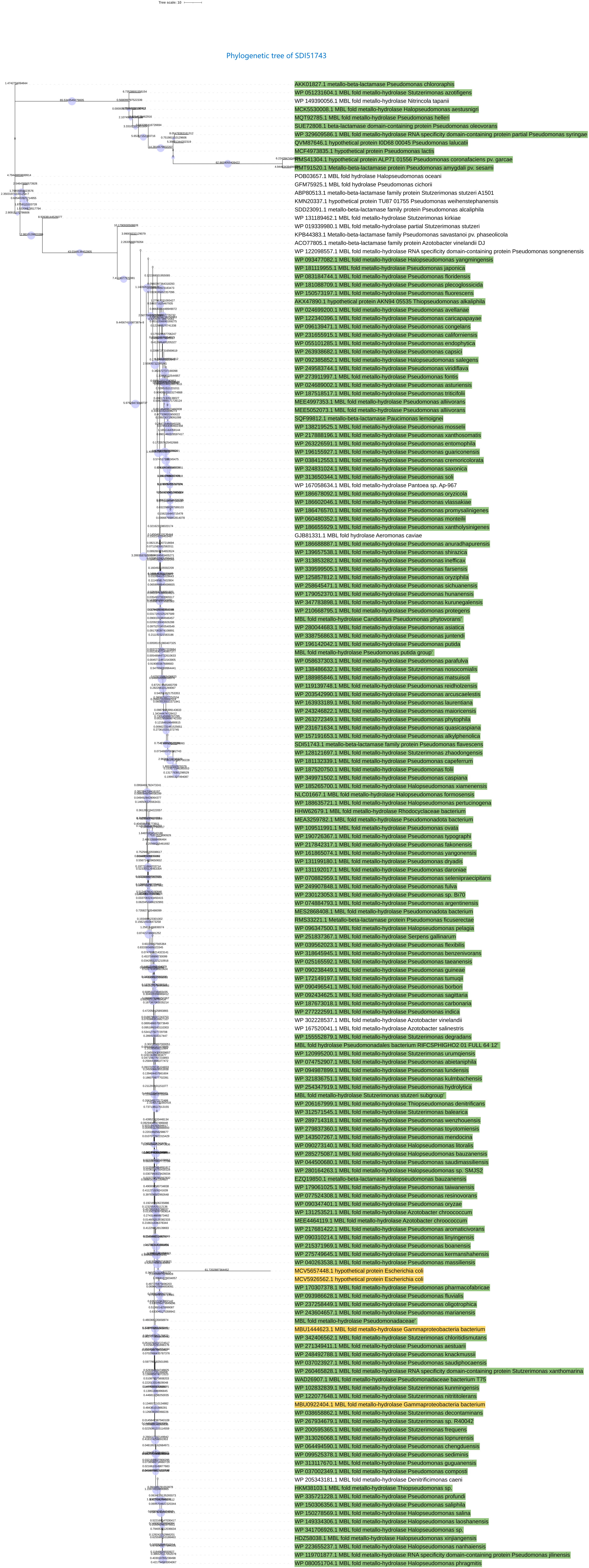

Phylogenetic tree of SEJ69490.1

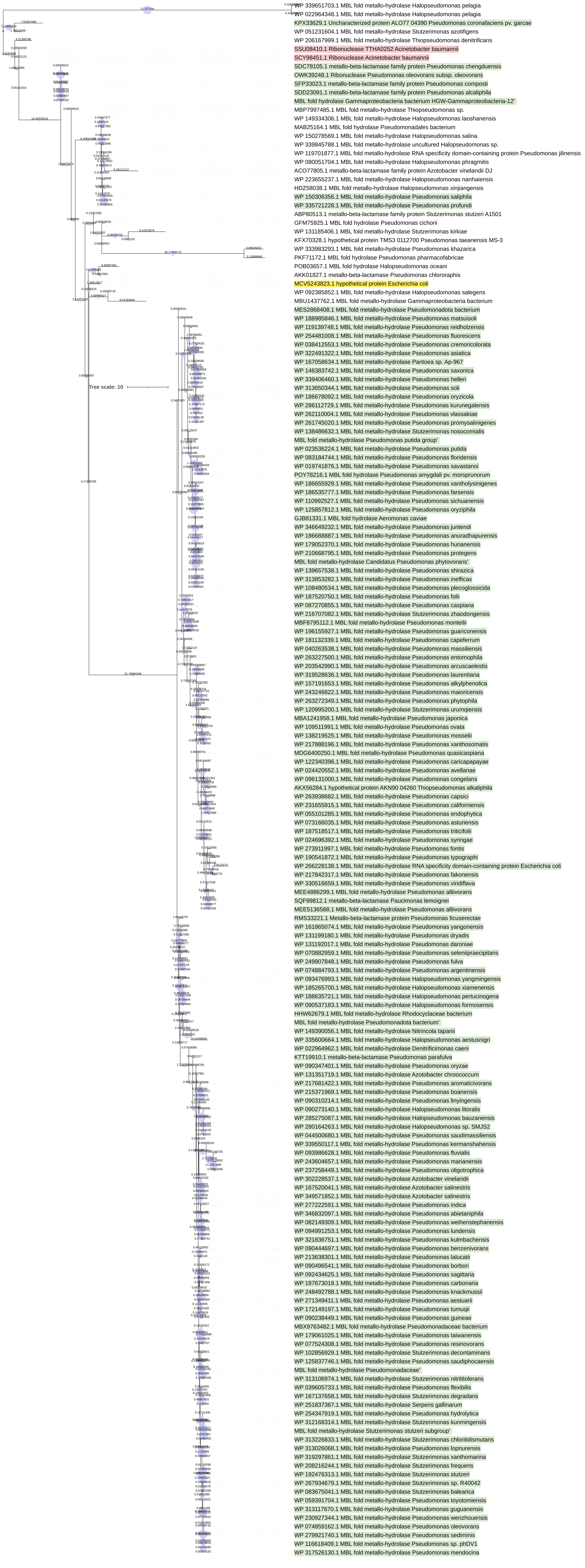

### Phylogenetic tree of AmpD

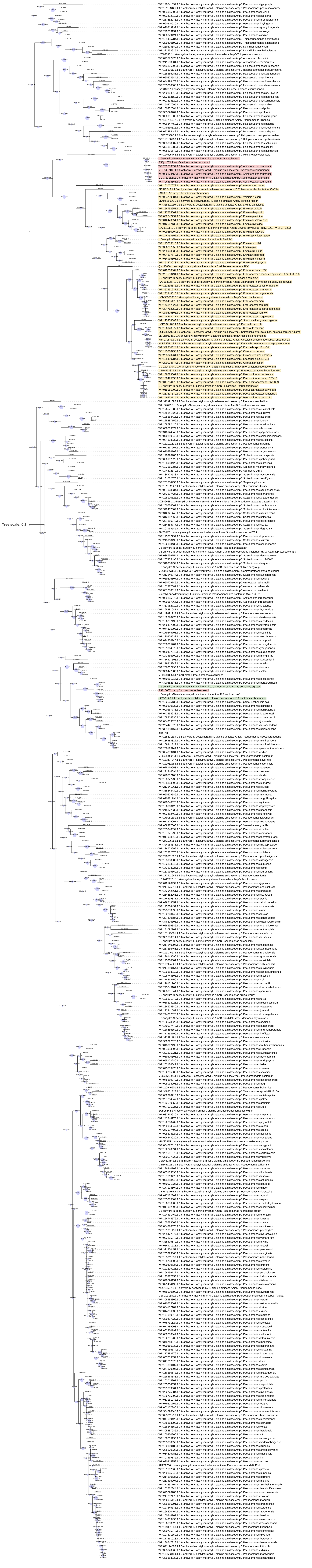

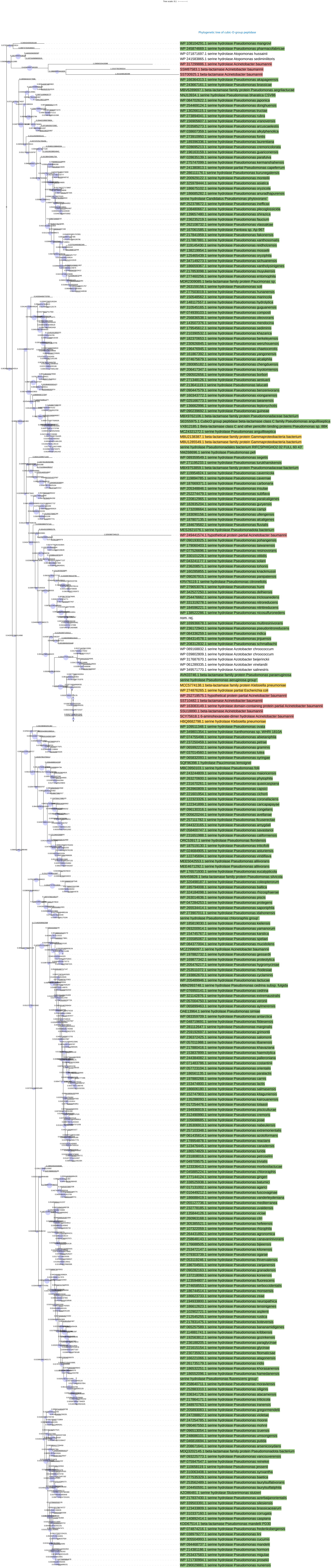

Tree scale: 0.1
Phylogenetic tree of cubic-O group peptidase
WP 108104291.1 serine hydrolase Pseudomonas mangrovi
WP 245874669.1 serine hydrolase Pseudomonas pharmacofabricae
WP 071871697.1 serine hydrolase Atopomonas hussainii
WP 241583865.1 serine hydrolase Atopomonas sedimenilitoris
WP 317299886.1 serine hydrolase Acinetobacter baumannii
SSM87583.1 beta-lactamase Acinetobacter baumannii
SST00925.1 beta-lactamase Acinetobacter baumannii
WP 166364313.1 serine hydrolase Pseudomonas akapageensis
WP 243667161.1 serine hydrolase Pseudomonas brassicae
MBV6289697.1 beta-lactamase family protein Pseudomonas aegrilactucae
NNJ13934.1 serine hydrolase Pseudomonas bharatica CSV86
WP 084702822.1 serine hydrolase Pseudomonas japonica
WP 254469124.1 serine hydrolase Pseudomonas donghuensis
WP 130266115.1 serine hydrolase Pseudomonas fructue
WP 273894041.1 serine hydrolase Pseudomonas rubra
WP 156905607.1 serine hydrolase Pseudomonas vranovensis
WP 205885273.1 serine hydrolase Pseudomonas arcuscaelestis
WP 038607359.1 serine hydrolase Pseudomonas alkylphenolica
WP 273910950.1 serine hydrolase Pseudomonas fontis
WP 189396336.1 serine hydrolase Pseudomonas laurentiana
WP 028695213.1 serine hydrolase Pseudomonas cremoricolorata
WP 196163159.1 serine hydrolase Pseudomonas guariconensis
WP 028635139.1 serine hydrolase Pseudomonas parafulva
WP 275747099.1 serine hydrolase Pseudomonas kermanshahensis
WP 241380813.1 serine hydrolase partial Pseudomonas capeferrum
WP 286111176.1 serine hydrolase Pseudomonas kurunegalensis
WP 200929122.1 serine hydrolase Pseudomonas monteilii
WP 325978444.1 serine hydrolase Pseudomonas asiatica
WP 186875102.1 serine hydrolase Pseudomonas oryzicola
WP 186685282.1 serine hydrolase Pseudomonas anuradhapurensis
serine hydrolase Candidatus Pseudomonas phytovorans
WP 252378672.1 serine hydrolase Pseudomonas inefficax
WP 108480667.1 serine hydrolase Pseudomonas plecoglossicida
WP 139657480.1 serine hydrolase Pseudomonas shirazica
WP 236235219.1 serine hydrolase Pseudomonas faucium
WP 262108732.1 serine hydrolase Pseudomonas vlassakiae
WP 167061585.1 serine hydrolase Pantoea sp. Ap-967
WP 217841959.1 serine hydrolase Pseudomonas fakonensis
WP 217887881.1 serine hydrolase Pseudomonas xanthosomatis
WP 219145430.1 serine hydrolase Pseudomonas reidholzensis
WP 138219954.1 serine hydrolase Pseudomonas mosselii
WP 125465439.1 serine hydrolase Pseudomonas oryzihabitans
WP 347149273.1 serine hydrolase Pseudomonas sichuanensis
WP 186659367.1 serine hydrolase Pseudomonas xanthosyringes
WP 217853098.1 serine hydrolase Pseudomonas muyukensis
WP 227460256.1 serine hydrolase Pseudomonas entomophila
MDR2309085.1 beta-lactamase family protein Pseudomonas sp.
WP 263159156.1 serine hydrolase Pseudomonas soli
WP 277503019.1 serine hydrolase Pseudomonas hunanensis
WP 150548562.1 serine hydrolase Pseudomonas maniocola
WP 148117557.1 serine hydrolase Pseudomonas hydrolytica
WP 310545165.1 serine hydrolase Pseudomonas guguanensis
WP 074939103.1 serine hydrolase Pseudomonas composti
WP 256835538.1 serine hydrolase Pseudomonas oleovorans
WP 143507376.1 serine hydrolase Pseudomonas mendocina
WP 179545612.1 serine hydrolase Pseudomonas sediminis
WP 210390532.1 serine hydrolase Pseudomonas khazarica
WP 182370653.1 serine hydrolase Pseudomonas berkeleyensis
WP 230929845.1 serine hydrolase Pseudomonas wenzhouensis
WP 190476945.1 serine hydrolase Pseudomonas chaetoceroris
WP 161867392.1 serine hydrolase Pseudomonas yangonensis
WP 074675679.1 serine hydrolase Pseudomonas alcaliphila
WP 280085182.1 serine hydrolase Pseudomonas chengduensis
WP 206417347.1 serine hydrolase Pseudomonas toyotomiensis
WP 090502656.1 serine hydrolase Pseudomonas borbori
WP 271348126.1 serine hydrolase Pseudomonas aestuarii
WP 213643119.1 serine hydrolase Pseudomonas taiwanensis
WP 090447579.1 serine hydrolase Pseudomonas benzenivorans
WP 160343772.1 serine hydrolase Pseudomonas xionganensis
WP 025166773.1 serine hydrolase Pseudomonas taiwanensis
WP 136662996.1 serine hydrolase Pseudomonas leptonychotis
WP 090239692.1 serine hydrolase Pseudomonas guineae
MBX9762106.1 beta-lactamase family protein Pseudomonadaceae bacterium
SED55975.1 CubicO group peptidase beta-lactamase class C family Pseudomonas anguilliseptica
VXB32185.1 Beta-lactamase class C and other penicillin binding proteins Pseudomonas sp. 8BK
MCZ4321222.1 serine hydrolase Pseudomonas anguilliseptica
MBU2138387.1 beta-lactamase family protein Gammaproteobacteria bacterium
MBU1285549.1 beta-lactamase family protein Gammaproteobacteria bacterium
serine hydrolase Pseudomonadales bacterium RIFCSPHIGHO2 02 FULL 60 43
NMZ68696.1 serine hydrolase Pseudomonas pelli
WP 089359549.1 serine hydrolase Pseudomonas segetis
WP 271195233.1 serine hydrolase Pseudomonas turukhanskensis
MBX9753859.1 beta-lactamase family protein Pseudomonadaceae bacterium
WP 119954824.1 serine hydrolase Pseudomonas cavernae
WP 119894785.1 serine hydrolase Pseudomonas cavernicola
WP 187669371.1 serine hydrolase Pseudomonas carboxidia
WP 205348846.1 serine hydrolase Pseudomonas insulae
WP 252274479.1 serine hydrolase Pseudomonas sulfuyia
WP 220812965.1 serine hydrolase Pseudomonas paralcaligenes
WP 182835204.1 serine hydrolase Pseudomonas guryensis
WP 173208664.1 serine hydrolase Pseudomonas campi
WP 183090156.1 serine hydrolase Pseudomonas ullengensis
WP 187807135.1 serine hydrolase Pseudomonas alcaligenes
WP 184679582.1 serine hydrolase Pseudomonas fluvialis
MES2821076.1 serine hydrolase Pseudomonadota bacterium
WP 249441574.1 hypothetical protein partial Acinetobacter baumannii
WP 090193025.1 serine hydrolase Pseudomonas pohangensis
WP 179060403.1 serine hydrolase Pseudomonas taiwanensis
WP 077526696.1 serine hydrolase Pseudomonas resinovorans
WP 330101228.1 serine hydrolase Pseudomonas otitidis
WP 043241177.1 serine hydrolase Pseudomonas solani
WP 236208571.1 serine hydrolase Pseudomonas tohonis
WP 160285855.1 serine hydrolase Pseudomonas knackmussii
WP 090267915.1 serine hydrolase Pseudomonas panipatensis
KRV76119.1 serine hydrolase Pseudomonas citronellolis
WP 279653076.1 serine hydrolase Pseudomonas humi
WP 342527250.1 serine hydrolase Pseudomonas delhiensis
WP 254476662.1 serine hydrolase Pseudomonas triclosanidens
WP 331319278.1 serine hydrolase Pseudomonas nitroreducens
WP 184596221.1 serine hydrolase Pseudomonas nitritireducens
WP 138522286.1 serine hydrolase Pseudomonas nicosulfuronedens
nom. rej.
WP 169936676.1 serine hydrolase Pseudomonas multiresinivorans
WP 236172943.1 serine hydrolase Pseudomonas pseudonitroreducens
WP 084336259.1 serine hydrolase Pseudomonas indica
WP 084314578.1 serine hydrolase Pseudomonas jinjuensis
WP 208312832.1 serine hydrolase Pseudomonas schmalbachii
WP 089168832.1 serine hydrolase Azotobacter chroococcum
WP 039802809.1 serine hydrolase Azotobacter chroococcum
WP 317687670.1 serine hydrolase Azotobacter beijerinckii
WP 061289335.1 serine hydrolase Azotobacter vinelandii
WP 349571770.1 serine hydrolase Azotobacter salinestris
AVK03746.1 beta-lactamase family protein Pseudomonas paraeruginosa
serine hydrolase Pseudomonas aeruginosa group
MCC5774138.1 beta-lactamase family protein Klebsiella pneumoniae
WP 274876265.1 serine hydrolase partial Escherichia coli
WP 252719575.1 hypothetical protein partial Acinetobacter baumannii
SST10482.1 beta-lactamase Acinetobacter baumannii
WP 163083149.1 serine hydrolase domain-containing protein partial Acinetobacter baumannii
SSU18890.1 beta-lactamase Acinetobacter baumannii
SCY75618.1 6-aminohexanoate-dimer hydrolase Acinetobacter baumannii
HBQ868798.1 serine hydrolase Klebsiella pneumoniae
WP 109513348.1 serine hydrolase Pseudomonas ovata
WP 349801354.1 serine hydrolase Xanthomonas sp. WHRI 1810A
WP 074755498.1 serine hydrolase Pseudomonas abietaniphila
WP 237260459.1 serine hydrolase Pseudomonas petrae
WP 065992232.1 serine hydrolase Pseudomonas graminis
WP 037014560.1 serine hydrolase Pseudomonas lutea
WP 065832093.1 serine hydrolase Pseudomonas syringae
SQF96398.1 hydrolase Pseudomonas lemoignei
MBC3950103.1 serine hydrolase Pseudomonas folii
WP 243244809.1 serine hydrolase Pseudomonas maioricensis
WP 263270600.1 serine hydrolase Pseudomonas phylophila
WP 231670261.1 serine hydrolase Pseudomonas quasicaspiana
WP 263960809.1 serine hydrolase Pseudomonas caspiana
WP 221601954.1 serine hydrolase Pseudomonas cichorii
WP 122323326.1 serine hydrolase Pseudomonas coronafaciens
WP 122341899.1 serine hydrolase Pseudomonas cancapapayae
WP 086130316.1 serine hydrolase Pseudomonas congelans
WP 005620244.1 serine hydrolase Pseudomonas avellanae
WP 257117821.1 serine hydrolase Pseudomonas ficuserectae
WP 044323165.1 serine hydrolase Pseudomonas amygdali
WP 058400747.1 serine hydrolase Pseudomonas savastanoi
WP 231651988.1 serine hydrolase Pseudomonas californiensis
CRC53917.1 serine hydrolase Pseudomonas floridensis
WP 187519130.1 serine hydrolase Pseudomonas triticifolii
WP 024684905.1 serine hydrolase Pseudomonas asturiensis
WP 122745694.1 serine hydrolase Pseudomonas viridiflava
MEE5042553.1 serine hydrolase Pseudomonas allivorans
MEE4671292.1 serine hydrolase Pseudomonas allivorans
WP 176571930.1 serine hydrolase Pseudomonas eucalypticola
WAH59828.1 beta-lactamase family protein Pseudomonas silvicola
WP 320498187.1 serine hydrolase Pseudomonas coleopterorum
WP 185794008.1 serine hydrolase Pseudomonas baltica
WP 324184098.1 serine hydrolase Pseudomonas rhizosphaerae
WP 263014636.1 serine hydrolase Pseudomonas piscis
WP 247284253.1 serine hydrolase Pseudomonas protegens
WP 265534414.1 serine hydrolase Pseudomonas saponiphila
WP 273907011.1 serine hydrolase Pseudomonas idahonensis
serine hydrolase Pseudomonas chlororaphis group
WP 185819030.1 serine hydrolase Pseudomonas kielensis
WP 269200914.1 serine hydrolase Pseudomonas yamanorum
WP 154745767.1 serine hydrolase Pseudomonas karstica
WP 155580067.1 serine hydrolase Pseudomonas spelaei
WP 084377004.1 serine hydrolase Pseudomonas mucidolens
MCZ2996097.1 serine hydrolase Acinetobacter baumannii
WP 197882236.1 serine hydrolase Pseudomonas gessardii
WP 169877342.1 serine hydrolase Pseudomonas proteolytica
WP 205476217.1 serine hydrolase Pseudomonas hygromyciniae
WP 253511072.1 serine hydrolase Pseudomonas rhodesiae
WP 193662670.1 serine hydrolase Pseudomonas cyclaminis
WP 205489946.1 serine hydrolase Pseudomonas lactucae
MBN2993748.1 serine hydrolase Pseudomonas cedrina subsp. fulgida
WP 076950541.1 serine hydrolase Pseudomonas cedrina
WP 321142876.1 serine hydrolase Pseudomonas extremaustralis
WP 057004755.1 serine hydrolase Pseudomonas veronii
WP 085899463.1 serine hydrolase Pseudomonas azotoformans
OAE13964.1 serine hydrolase Pseudomonas simiae
WP 083399709.1 serine hydrolase Pseudomonas antarctica
WP 048719691.1 serine hydrolase Pseudomonas fildesensis
WP 281112647.1 serine hydrolase Pseudomonas marginalis
WP 259192697.1 serine hydrolase Pseudomonas grimontii
WP 236372425.1 serine hydrolase Pseudomonas salomonii
WP 057011988.1 serine hydrolase Pseudomonas libanensis
WP 217680416.1 serine hydrolase Pseudomonas khavaziana
WP 153837899.1 serine hydrolase Pseudomonas haemolytica
WP 244384082.1 serine hydrolase Pseudomonas palleroniana
WP 071483786.1 serine hydrolase Pseudomonas costantinii
WP 057723104.1 serine hydrolase Pseudomonas orientalis
WP 186561135.1 serine hydrolase Pseudomonas paracarnis
WP 197880268.1 serine hydrolase Pseudomonas carnis
WP 153474800.1 serine hydrolase Pseudomonas lactis
WP 186608180.1 serine hydrolase Pseudomonas salmasensis
WP 152747903.1 serine hydrolase Pseudomonas kitaguniensis
WP 135289063.1 serine hydrolase Pseudomonas kairouanensis
WP 017254476.1 serine hydrolase Pseudomonas tolaasii
WP 194936919.1 serine hydrolase Pseudomonas glycinae
WP 312490990.1 serine hydrolase Pseudomonas cremoris
WP 105698685.1 serine hydrolase Pseudomonas poae
WP 135309613.1 serine hydrolase Pseudomonas nabeulensis
WP 257103348.1 serine hydrolase Pseudomonas extremorientalis
WP 061435614.1 serine hydrolase Pseudomonas azotoformans
WP 178954878.1 serine hydrolase Pseudomonas reactans
WP 123478445.1 serine hydrolase Pseudomonas canadensis
WP 186574829.1 serine hydrolase Pseudomonas lundia
WP 231808016.1 serine hydrolase Pseudomonas petroselini
WP 049709575.1 serine hydrolase Pseudomonas trivialis
WP 123336413.1 serine hydrolase Pseudomonas mohnii/glaciei
WP 045885224.1 serine hydrolase Pseudomonas chlororaphis
WP 177144124.1 serine hydrolase Pseudomonas gingeri
WP 138525938.1 serine hydrolase Pseudomonas vranovensis
WP 017130022.1 serine hydrolase Pseudomonas agarici
WP 101449212.1 serine hydrolase Pseudomonas fuscovaginae
WP 186886419.1 serine hydrolase Pseudomonas vanderleydeniana
WP 055127736.1 serine hydrolase Pseudomonas mediterranea
WP 232778185.1 serine hydrolase Pseudomonas uvaldensis
WP 135844126.1 serine hydrolase Pseudomonas viciae
WP 260963169.1 serine hydrolase Pseudomonas cilri
WP 305380521.1 serine hydrolase Pseudomonas hefeiensis
WP 107320559.1 serine hydrolase Pseudomonas rhizophila
WP 264431892.1 serine hydrolase Pseudomonas agronomica
WP 258648143.1 serine hydrolase Pseudomonas canavaninivorans
WP 176688505.1 serine hydrolase Pseudomonas bijieensis
WP 253472147.1 serine hydrolase Pseudomonas kilonensis
WP 079303739.1 serine hydrolase Pseudomonas ogarae
WP 253119246.1 serine hydrolase Pseudomonas thivervalensis
WP 186704501.1 serine hydrolase Pseudomonas zarqanensis
WP 060282163.1 serine hydrolase Pseudomonas granadensis
WP 137218060.1 serine hydrolase Pseudomonas koreensis
WP 123594807.1 serine hydrolase Pseudomonas fluorescens
WP 274658553.1 serine hydrolase Pseudomonas serbocciidentalis
WP 186744514.1 serine hydrolase Pseudomonas monsensis
WP 186623733.1 serine hydrolase Pseudomonas zeae
WP 184933800.1 serine hydrolase Pseudomonas neuropathica
WP 186612603.1 serine hydrolase Pseudomonas tensinigenes
WP 102927721.1 serine hydrolase Pseudomonas asplenii
WP 212545255.1 serine hydrolase Pseudomonas rustica
WP 217831475.1 serine hydrolase Pseudomonas botevensis
WP 060257599.1 serine hydrolase Pseudomonas bananamidigenes
WP 114881741.1 serine hydrolase Pseudomonas kribbensis
WP 192562811.1 serine hydrolase Pseudomonas gozinkensis
WP 236180255.1 serine hydrolase Pseudomonas paraglycinae
WP 221615154.1 serine hydrolase Pseudomonas glycinae
WP 230730599.1 serine hydrolase Pseudomonas fitomaticsae
WP 166210066.1 serine hydrolase Pseudomonas atagonensis
WP 181735379.1 serine hydrolase Pseudomonas laurylsulfatiphila
WP 186532251.1 serine hydrolase Pseudomonas khorasanensis
WP 186552096.1 serine hydrolase Pseudomonas hamedanensis
serine hydrolase Pseudomonas fluorescens group
WP 258540711.1 serine hydrolase Pseudomonas moraviensis
WP 253051315.1 serine hydrolase Pseudomonas siliginis
WP 336341726.1 serine hydrolase Pseudomonas atacamensis
WP 217864171.1 serine hydrolase Pseudomonas triticicola
WP 349970763.1 serine hydrolase Pseudomonas iranensis
WP 205893083.1 serine hydrolase Pseudomonas gregormendelii
WP 247288827.1 serine hydrolase Pseudomonas violetae
WP 247254785.1 serine hydrolase Pseudomonas moorei
WP 090497500.1 serine hydrolase Pseudomonas mohnii
WP 098513054.1 serine hydrolase Pseudomonas izuensis
WP 248698101.1 serine hydrolase Pseudomonas umsongensis
WP 246816694.1 serine hydrolase Pseudomonas putida
WP 208671641.1 serine hydrolase Pseudomonas arsenicoxydans
MCQ0202145.1 beta-lactamase family protein Pseudomonadota bacterium
WP 093225573.1 serine hydrolase Pseudomonas vancouverensis
WP 075947547.1 serine hydrolase Pseudomonas reinekei
WP 110658119.1 serine hydrolase Pseudomonas jessenii
WP 110063408.1 serine hydrolase Pseudomonas synxantha
WP 277535329.1 serine hydrolase Pseudomonas baetica
WP 255662489.1 serine hydrolase Pseudomonas laurylsulfativorans
WP 104450591.1 serine hydrolase Pseudomonas laurylsulfatovorans
AZO86481.1 serine hydrolase Stutzerimonas stutzeri
WP 217837430.1 serine hydrolase Pseudomonas azerbaijanorientalis
WP 339603291.1 serine hydrolase Pseudomonas silesiensis
WP 123433809.1 serine hydrolase Pseudomonas brassicacearum
WP 310337160.1 serine hydrolase Pseudomonas corrugata
WP 140892414.1 serine hydrolase Pseudomonas caspiana
KDD67514.1 beta-lactamase Pseudomonas mandelii PD30
WP 074874216.1 serine hydrolase Pseudomonas frederiksbergensis
WP 038977977.1 serine hydrolase Pseudomonas lini
WP 080950040.1 serine hydrolase Pseudomonas cucumis
WP 094488737.1 serine hydrolase Pseudomonas mandelii
WP 214381186.1 serine hydrolase Pseudomonas hormoni
WP 253427583.1 serine hydrolase Pseudomonas migulae
WP 121730994.1 serine hydrolase Pseudomonas prosekii
WP 290028881.1 serine hydrolase Pseudomonas sivasensis

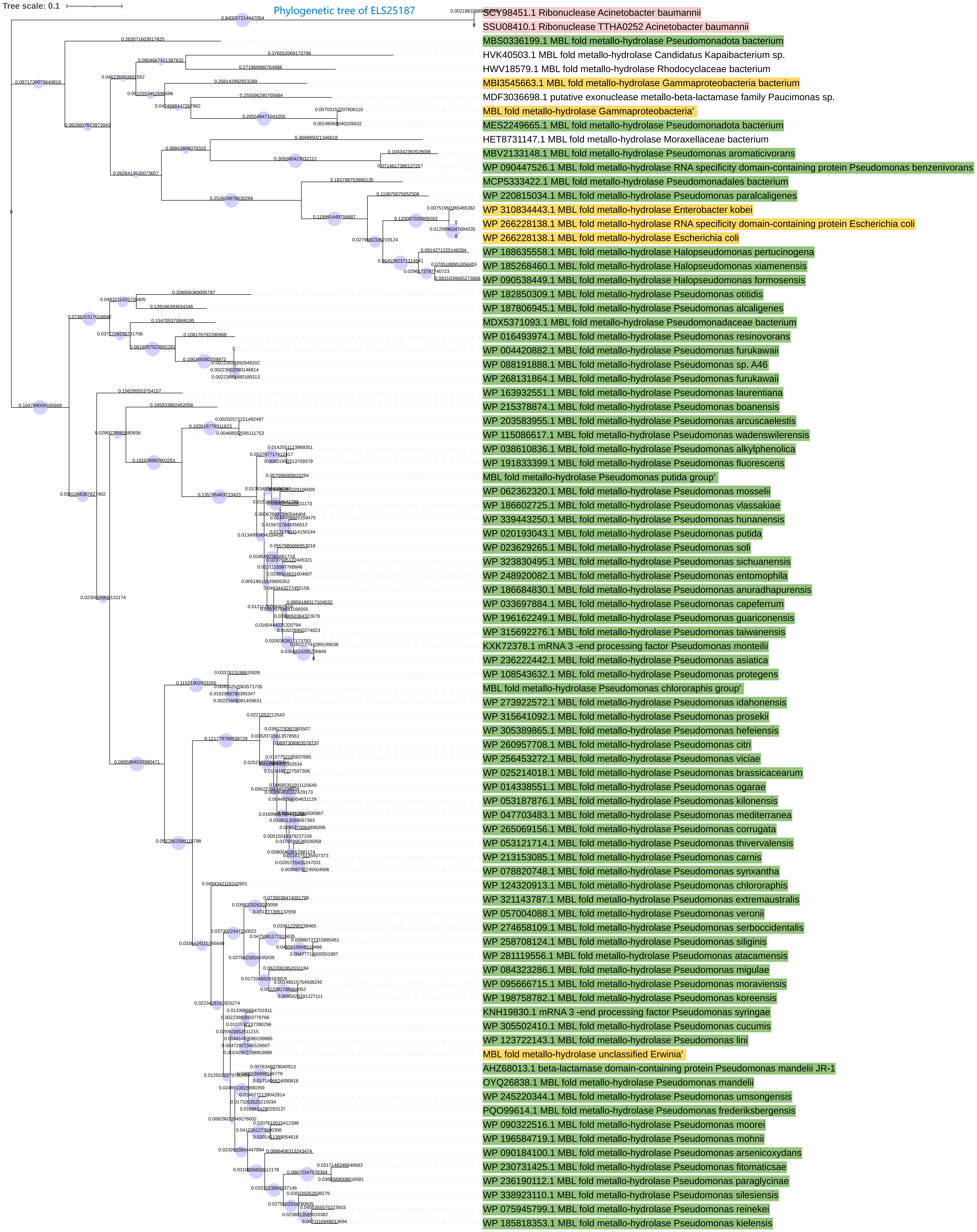

Tree scale: 0.1
Phylogenetic tree of ELS25187
SCY98451.1 Ribonuclease Acinetobacter baumannii
SSU08410.1 Ribonuclease TTHA0252 Acinetobacter baumannii
MBS0336199.1 MBL fold metallo-hydrolase Pseudomonadota bacterium
HVK40503.1 MBL fold metallo-hydrolase Candidatus Kapaibacterium sp.
HWV18579.1 MBL fold metallo-hydrolase Rhodocyclaceae bacterium
MBI3545663.1 MBL fold metallo-hydrolase Gammaproteobacteria bacterium
MDF3036698.1 putative exonuclease metallo-beta-lactamase family Paucimonas sp.
MBL fold metallo-hydrolase Gammaproteobacteria'
MES2249665.1 MBL fold metallo-hydrolase Pseudomonadota bacterium
HET8731147.1 MBL fold metallo-hydrolase Moraxellaceae bacterium
MBV2133148.1 MBL fold metallo-hydrolase Pseudomonas aromaticivorans
WP 090447526.1 MBL fold metallo-hydrolase RNA specificity domain-containing protein Pseudomonas benzenivorans
MCP5333422.1 MBL fold metallo-hydrolase Pseudomonadales bacterium
WP 220815034.1 MBL fold metallo-hydrolase Pseudomonas paralcaligenes
WP 310834443.1 MBL fold metallo-hydrolase Enterobacter kobei
WP 266228138.1 MBL fold metallo-hydrolase RNA specificity domain-containing protein Escherichia coli
WP 266228138.1 MBL fold metallo-hydrolase Escherichia coli
WP 188635558.1 MBL fold metallo-hydrolase Halopseudomonas pertucinogena
WP 185268460.1 MBL fold metallo-hydrolase Halopseudomonas xiamenensis
WP 090538449.1 MBL fold metallo-hydrolase Halopseudomonas formosensis
WP 182850309.1 MBL fold metallo-hydrolase Pseudomonas otitidis
WP 187806945.1 MBL fold metallo-hydrolase Pseudomonas alcaligenes
MDX5371093.1 MBL fold metallo-hydrolase Pseudomonadaceae bacterium
WP 016493974.1 MBL fold metallo-hydrolase Pseudomonas resinovorans
WP 004420882.1 MBL fold metallo-hydrolase Pseudomonas furukawaii
WP 088191888.1 MBL fold metallo-hydrolase Pseudomonas sp. A46
WP 268131864.1 MBL fold metallo-hydrolase Pseudomonas furukawaii
WP 163932551.1 MBL fold metallo-hydrolase Pseudomonas laurentiana
WP 215378874.1 MBL fold metallo-hydrolase Pseudomonas boanensis
WP 203583955.1 MBL fold metallo-hydrolase Pseudomonas arcuscaelestis
WP 115086617.1 MBL fold metallo-hydrolase Pseudomonas wadenswilerensis
WP 038610836.1 MBL fold metallo-hydrolase Pseudomonas alkylphenolica
WP 191833399.1 MBL fold metallo-hydrolase Pseudomonas fluorescens
MBL fold metallo-hydrolase Pseudomonas putida group'
WP 062362320.1 MBL fold metallo-hydrolase Pseudomonas mosselii
WP 186602725.1 MBL fold metallo-hydrolase Pseudomonas vlassakiae
WP 339443250.1 MBL fold metallo-hydrolase Pseudomonas hunanensis
WP 020193043.1 MBL fold metallo-hydrolase Pseudomonas putida
WP 023629265.1 MBL fold metallo-hydrolase Pseudomonas soli
WP 323830495.1 MBL fold metallo-hydrolase Pseudomonas sichuanensis
WP 248920082.1 MBL fold metallo-hydrolase Pseudomonas entomophila
WP 186684830.1 MBL fold metallo-hydrolase Pseudomonas anuradhapurensis
WP 033697884.1 MBL fold metallo-hydrolase Pseudomonas capeferrum
WP 196162249.1 MBL fold metallo-hydrolase Pseudomonas guariconensis
WP 315692276.1 MBL fold metallo-hydrolase Pseudomonas taiwanensis
KXK72378.1 mRNA 3 -end processing factor Pseudomonas monteilii
WP 236222442.1 MBL fold metallo-hydrolase Pseudomonas asiatica
WP 108543632.1 MBL fold metallo-hydrolase Pseudomonas protegens
MBL fold metallo-hydrolase Pseudomonas chlororaphis group'
WP 273922572.1 MBL fold metallo-hydrolase Pseudomonas idahonensis
WP 315641092.1 MBL fold metallo-hydrolase Pseudomonas prosekii
WP 305389865.1 MBL fold metallo-hydrolase Pseudomonas hefeiensis
WP 260957708.1 MBL fold metallo-hydrolase Pseudomonas citri
WP 256453272.1 MBL fold metallo-hydrolase Pseudomonas viciae
WP 025214018.1 MBL fold metallo-hydrolase Pseudomonas brassicacearum
WP 014338551.1 MBL fold metallo-hydrolase Pseudomonas ogarae
WP 053187876.1 MBL fold metallo-hydrolase Pseudomonas kilonensis
WP 047703483.1 MBL fold metallo-hydrolase Pseudomonas mediterranea
WP 265069156.1 MBL fold metallo-hydrolase Pseudomonas corrugata
WP 053121714.1 MBL fold metallo-hydrolase Pseudomonas thivervalensis
WP 213153085.1 MBL fold metallo-hydrolase Pseudomonas carnis
WP 078820748.1 MBL fold metallo-hydrolase Pseudomonas synxantha
WP 124320913.1 MBL fold metallo-hydrolase Pseudomonas chlororaphis
WP 321143787.1 MBL fold metallo-hydrolase Pseudomonas extremaustralis
WP 057004088.1 MBL fold metallo-hydrolase Pseudomonas veronii
WP 274658109.1 MBL fold metallo-hydrolase Pseudomonas serboccidentalis
WP 258708124.1 MBL fold metallo-hydrolase Pseudomonas siliginis
WP 281119556.1 MBL fold metallo-hydrolase Pseudomonas atacamensis
WP 084323286.1 MBL fold metallo-hydrolase Pseudomonas migulae
WP 095666715.1 MBL fold metallo-hydrolase Pseudomonas moraviensis
WP 198758782.1 MBL fold metallo-hydrolase Pseudomonas koreensis
KNH19830.1 mRNA 3 -end processing factor Pseudomonas syringae
WP 305502410.1 MBL fold metallo-hydrolase Pseudomonas cucumis
WP 123722143.1 MBL fold metallo-hydrolase Pseudomonas lini
MBL fold metallo-hydrolase unclassified Erwinia'
AHZ68013.1 beta-lactamase domain-containing protein Pseudomonas mandelii JR-1
OYQ26838.1 MBL fold metallo-hydrolase Pseudomonas mandelii
WP 245220344.1 MBL fold metallo-hydrolase Pseudomonas umsongensis
PQO99614.1 MBL fold metallo-hydrolase Pseudomonas frederiksbergensis
WP 090322516.1 MBL fold metallo-hydrolase Pseudomonas moorei
WP 196584719.1 MBL fold metallo-hydrolase Pseudomonas mohnii
WP 090184100.1 MBL fold metallo-hydrolase Pseudomonas arsenicoxydans
WP 230731425.1 MBL fold metallo-hydrolase Pseudomonas fitomaticsae
WP 236190112.1 MBL fold metallo-hydrolase Pseudomonas paraglycinae
WP 338923110.1 MBL fold metallo-hydrolase Pseudomonas silesiensis
WP 075945799.1 MBL fold metallo-hydrolase Pseudomonas reinekei
WP 185818353.1 MBL fold metallo-hydrolase Pseudomonas kielensis

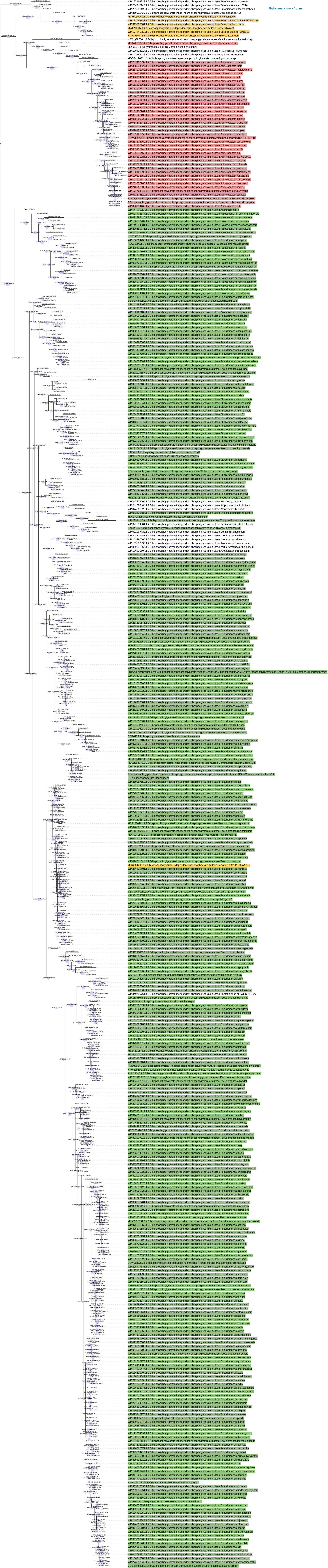
Phylogenetic tree of gpmI

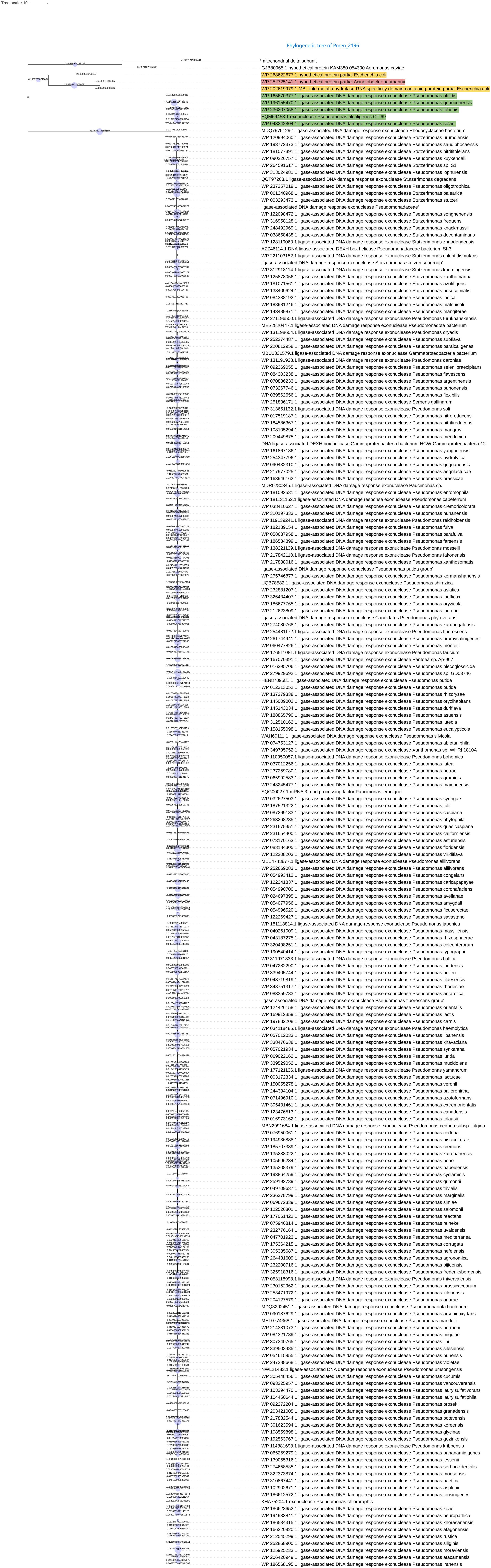
Tree scale: 10
Phylogenetic tree of Pmen_2196
mitochondrial delta subunit
GJB80965.1 hypothetical protein KAM380 054300 Aeromonas caviae
WP 268622677.1 hypothetical protein partial Escherichia coli
WP 252725141.1 hypothetical protein partial Acinetobacter baumannii
WP 202619979.1 MBL fold metallo-hydrolase RNA specificity domain-containing protein partial Escherichia coli
WP 165670377.1 ligase-associated DNA damage response exonuclease Pseudomonas otitidis
WP 196155470.1 ligase-associated DNA damage response exonuclease Pseudomonas guariconensis
WP 236207058.1 ligase-associated DNA damage response exonuclease Pseudomonas tohonis
EQM69458.1 exonuclease Pseudomonas alcaligenes OT 69
WP 043242804.1 ligase-associated DNA damage response exonuclease Pseudomonas solani
MDQ7975129.1 ligase-associated DNA damage response exonuclease Rhodocyclaceae bacterium
WP 120994060.1 ligase-associated DNA damage response exonuclease Stutzerimonas urumqiensis
WP 193772373.1 ligase-associated DNA damage response exonuclease Pseudomonas saudiphocaensis
WP 181077391.1 ligase-associated DNA damage response exonuclease Stutzerimonas nitrititolerans
WP 090226757.1 ligase-associated DNA damage response exonuclease Pseudomonas kuykendallii
WP 264591617.1 ligase-associated DNA damage response exonuclease Stutzerimonas sp. S1
WP 313024981.1 ligase-associated DNA damage response exonuclease Pseudomonas lopnurensis
QCT97263.1 ligase-associated DNA damage response exonuclease Stutzerimonas degradans
WP 237257019.1 ligase-associated DNA damage response exonuclease Pseudomonas oligotrophica
WP 061340968.1 ligase-associated DNA damage response exonuclease Stutzerimonas balearica
WP 003293473.1 ligase-associated DNA damage response exonuclease Stutzerimonas stutzeri
ligase-associated DNA damage response exonuclease Pseudomonadaceae'
WP 122098472.1 ligase-associated DNA damage response exonuclease Pseudomonas songnenensis
WP 316958128.1 ligase-associated DNA damage response exonuclease Stutzerimonas frequens
WP 248492969.1 ligase-associated DNA damage response exonuclease Pseudomonas knackmussii
WP 038658438.1 ligase-associated DNA damage response exonuclease Stutzerimonas decontaminans
WP 128119063.1 ligase-associated DNA damage response exonuclease Stutzerimonas zhaodongensis
AZZ46114.1 DNA ligase-associated DEXH box helicase Pseudomonadaceae bacterium SI-3
WP 221103152.1 ligase-associated DNA damage response exonuclease Stutzerimonas chloritidismutans
ligase-associated DNA damage response exonuclease Stutzerimonas stutzeri subgroup'
WP 312918114.1 ligase-associated DNA damage response exonuclease Stutzerimonas kunmingensis
WP 125878056.1 ligase-associated DNA damage response exonuclease Stutzerimonas xanthomarina
WP 181071561.1 ligase-associated DNA damage response exonuclease Stutzerimonas azotifigens
WP 138409624.1 ligase-associated DNA damage response exonuclease Stutzerimonas nosocomialis
WP 084338192.1 ligase-associated DNA damage response exonuclease Pseudomonas indica
WP 188981246.1 ligase-associated DNA damage response exonuclease Pseudomonas matsuisoli
WP 143489871.1 ligase-associated DNA damage response exonuclease Pseudomonas mangiferae
WP 271196500.1 ligase-associated DNA damage response exonuclease Pseudomonas turukhanskensis
MES2820447.1 ligase-associated DNA damage response exonuclease Pseudomonadota bacterium
WP 131198604.1 ligase-associated DNA damage response exonuclease Pseudomonas dryadis
WP 252274487.1 ligase-associated DNA damage response exonuclease Pseudomonas subflava
WP 220812958.1 ligase-associated DNA damage response exonuclease Pseudomonas paralcaligenes
MBU1331579.1 ligase-associated DNA damage response exonuclease Gammaproteobacteria bacterium
WP 131191928.1 ligase-associated DNA damage response exonuclease Pseudomonas daroniae
WP 092369055.1 ligase-associated DNA damage response exonuclease Pseudomonas seleniipraecipitans
WP 084303238.1 ligase-associated DNA damage response exonuclease Pseudomonas flavescens
WP 070886233.1 ligase-associated DNA damage response exonuclease Pseudomonas argentinensis
WP 073267746.1 ligase-associated DNA damage response exonuclease Pseudomonas punonensis
WP 039562656.1 ligase-associated DNA damage response exonuclease Pseudomonas flexibilis
WP 251836171.1 ligase-associated DNA damage response exonuclease Serpens gallinarum
WP 313651132.1 ligase-associated DNA damage response exonuclease Pseudomonas soli
WP 017519187.1 ligase-associated DNA damage response exonuclease Pseudomonas nitroreducens
WP 184586367.1 ligase-associated DNA damage response exonuclease Pseudomonas nitritireducens
WP 108105294.1 ligase-associated DNA damage response exonuclease Pseudomonas mangrovi
WP 209449875.1 ligase-associated DNA damage response exonuclease Pseudomonas mendocina
DNA ligase-associated DEXH box helicase Gammaproteobacteria bacterium HGW-Gammaproteobacteria-12'
WP 161867136.1 ligase-associated DNA damage response exonuclease Pseudomonas yangonensis
WP 254347796.1 ligase-associated DNA damage response exonuclease Pseudomonas hydrolytica
WP 090432310.1 ligase-associated DNA damage response exonuclease Pseudomonas guguanensis
WP 217977025.1 ligase-associated DNA damage response exonuclease Pseudomonas aegrilactucae
WP 163946162.1 ligase-associated DNA damage response exonuclease Pseudomonas brassicae
MDR0280345.1 ligase-associated DNA damage response exonuclease Paucimonas sp.
WP 181092531.1 ligase-associated DNA damage response exonuclease Pseudomonas entomophila
WP 181131152.1 ligase-associated DNA damage response exonuclease Pseudomonas capeferrum
WP 038410627.1 ligase-associated DNA damage response exonuclease Pseudomonas cremoricolorata
WP 310197333.1 ligase-associated DNA damage response exonuclease Pseudomonas hunanensis
WP 119139241.1 ligase-associated DNA damage response exonuclease Pseudomonas reidholzensis
WP 182139154.1 ligase-associated DNA damage response exonuclease Pseudomonas fulva
WP 058637958.1 ligase-associated DNA damage response exonuclease Pseudomonas parafulva
WP 186534899.1 ligase-associated DNA damage response exonuclease Pseudomonas farsensis
WP 138221139.1 ligase-associated DNA damage response exonuclease Pseudomonas mosselii
WP 217842110.1 ligase-associated DNA damage response exonuclease Pseudomonas fakonensis
WP 217888016.1 ligase-associated DNA damage response exonuclease Pseudomonas xanthosomatis
ligase-associated DNA damage response exonuclease Pseudomonas putida group'
WP 275746877.1 ligase-associated DNA damage response exonuclease Pseudomonas kermanshahensis
UQB78582.1 ligase-associated DNA damage response exonuclease Pseudomonas shirazica
WP 232881207.1 ligase-associated DNA damage response exonuclease Pseudomonas asiatica
WP 326434407.1 ligase-associated DNA damage response exonuclease Pseudomonas inefficax
WP 186677765.1 ligase-associated DNA damage response exonuclease Pseudomonas oryzicola
WP 212623809.1 ligase-associated DNA damage response exonuclease Pseudomonas juntendi
ligase-associated DNA damage response exonuclease Candidatus Pseudomonas phytovorans'
WP 274080768.1 ligase-associated DNA damage response exonuclease Pseudomonas kurunegalensis
WP 254481172.1 ligase-associated DNA damage response exonuclease Pseudomonas fluorescens
WP 261744941.1 ligase-associated DNA damage response exonuclease Pseudomonas promysalinigenes
WP 060477826.1 ligase-associated DNA damage response exonuclease Pseudomonas monteilii
WP 176511081.1 ligase-associated DNA damage response exonuclease Pseudomonas faucium
WP 167070391.1 ligase-associated DNA damage response exonuclease Pantoea sp. Ap-967
WP 016395706.1 ligase-associated DNA damage response exonuclease Pseudomonas plecoglossicida
WP 279929692.1 ligase-associated DNA damage response exonuclease Pseudomonas sp. GD03746
HEN8709581.1 ligase-associated DNA damage response exonuclease Pseudomonas putida
WP 012313052.1 ligase-associated DNA damage response exonuclease Pseudomonas putida
WP 137279338.1 ligase-associated DNA damage response exonuclease Pseudomonas rhizoryzae
WP 145009002.1 ligase-associated DNA damage response exonuclease Pseudomonas oryzihabitans
WP 145143034.1 ligase-associated DNA damage response exonuclease Pseudomonas duriflava
WP 188865790.1 ligase-associated DNA damage response exonuclease Pseudomonas asuensis
WP 312510162.1 ligase-associated DNA damage response exonuclease Pseudomonas luteola
WP 158155098.1 ligase-associated DNA damage response exonuclease Pseudomonas eucalypticola
WAH60111.1 ligase-associated DNA damage response exonuclease Pseudomonas silvicola
WP 074753127.1 ligase-associated DNA damage response exonuclease Pseudomonas abietaniphila
WP 349795752.1 ligase-associated DNA damage response exonuclease Xanthomonas sp. WHRI 1810A
WP 110950057.1 ligase-associated DNA damage response exonuclease Pseudomonas bohemica
WP 037012256.1 ligase-associated DNA damage response exonuclease Pseudomonas lutea
WP 237259780.1 ligase-associated DNA damage response exonuclease Pseudomonas petrae
WP 065992583.1 ligase-associated DNA damage response exonuclease Pseudomonas graminis
WP 243245477.1 ligase-associated DNA damage response exonuclease Pseudomonas maioricensis
SQG00027.1 mRNA 3 -end processing factor Paucimonas lemoignei
WP 032627503.1 ligase-associated DNA damage response exonuclease Pseudomonas syringae
WP 187521322.1 ligase-associated DNA damage response exonuclease Pseudomonas folii
WP 087269183.1 ligase-associated DNA damage response exonuclease Pseudomonas caspiana
WP 263268235.1 ligase-associated DNA damage response exonuclease Pseudomonas phytophila
WP 231675451.1 ligase-associated DNA damage response exonuclease Pseudomonas quasicaspiana
WP 231654400.1 ligase-associated DNA damage response exonuclease Pseudomonas californiensis
WP 073170163.1 ligase-associated DNA damage response exonuclease Pseudomonas asturiensis
WP 083184305.1 ligase-associated DNA damage response exonuclease Pseudomonas floridensis
WP 122208203.1 ligase-associated DNA damage response exonuclease Pseudomonas viridiflava
MEE4743877.1 ligase-associated DNA damage response exonuclease Pseudomonas alliivorans
WP 252669083.1 ligase-associated DNA damage response exonuclease Pseudomonas alliivorans
WP 054993412.1 ligase-associated DNA damage response exonuclease Pseudomonas congelans
WP 122341837.1 ligase-associated DNA damage response exonuclease Pseudomonas caricapapayae
WP 054990700.1 ligase-associated DNA damage response exonuclease Pseudomonas coronafaciens
WP 024697395.1 ligase-associated DNA damage response exonuclease Pseudomonas avellanae
WP 054077956.1 ligase-associated DNA damage response exonuclease Pseudomonas amygdali
WP 054996520.1 ligase-associated DNA damage response exonuclease Pseudomonas ficuserectae
WP 122269427.1 ligase-associated DNA damage response exonuclease Pseudomonas savastanoi
WP 181118814.1 ligase-associated DNA damage response exonuclease Pseudomonas japonica
WP 040261009.1 ligase-associated DNA damage response exonuclease Pseudomonas massiliensis
WP 043187275.1 ligase-associated DNA damage response exonuclease Pseudomonas rhizosphaerae
WP 320498251.1 ligase-associated DNA damage response exonuclease Pseudomonas coleopterorum
WP 190540414.1 ligase-associated DNA damage response exonuclease Pseudomonas typographi
WP 311971333.1 ligase-associated DNA damage response exonuclease Pseudomonas baltica
WP 047282290.1 ligase-associated DNA damage response exonuclease Pseudomonas lundensis
WP 339405744.1 ligase-associated DNA damage response exonuclease Pseudomonas helleri
WP 048719819.1 ligase-associated DNA damage response exonuclease Pseudomonas fildesensis
WP 348751317.1 ligase-associated DNA damage response exonuclease Pseudomonas rhodesiae
WP 083359783.1 ligase-associated DNA damage response exonuclease Pseudomonas antarctica
ligase-associated DNA damage response exonuclease Pseudomonas fluorescens group'
WP 124426158.1 ligase-associated DNA damage response exonuclease Pseudomonas orientalis
WP 169912359.1 ligase-associated DNA damage response exonuclease Pseudomonas lactis
WP 197882208.1 ligase-associated DNA damage response exonuclease Pseudomonas carnis
WP 034118485.1 ligase-associated DNA damage response exonuclease Pseudomonas haemolytica
WP 057012033.1 ligase-associated DNA damage response exonuclease Pseudomonas libanensis
WP 338476638.1 ligase-associated DNA damage response exonuclease Pseudomonas khavaziana
WP 057021934.1 ligase-associated DNA damage response exonuclease Pseudomonas synxantha
WP 069022162.1 ligase-associated DNA damage response exonuclease Pseudomonas lurida
WP 339529052.1 ligase-associated DNA damage response exonuclease Pseudomonas mucidolens
WP 177121136.1 ligase-associated DNA damage response exonuclease Pseudomonas yamanorum
WP 003172334.1 ligase-associated DNA damage response exonuclease Pseudomonas lactucae
WP 150055278.1 ligase-associated DNA damage response exonuclease Pseudomonas veronii
WP 244384104.1 ligase-associated DNA damage response exonuclease Pseudomonas palleroniana
WP 071496910.1 ligase-associated DNA damage response exonuclease Pseudomonas azotoformans
WP 305431461.1 ligase-associated DNA damage response exonuclease Pseudomonas extremorientalis
WP 123476513.1 ligase-associated DNA damage response exonuclease Pseudomonas canadensis
WP 016973162.1 ligase-associated DNA damage response exonuclease Pseudomonas tolaasii
MBN2991684.1 ligase-associated DNA damage response exonuclease Pseudomonas cedrina subsp. fulgida
WP 076950061.1 ligase-associated DNA damage response exonuclease Pseudomonas cedrina
WP 194936888.1 ligase-associated DNA damage response exonuclease Pseudomonas pisciculturae
WP 185707339.1 ligase-associated DNA damage response exonuclease Pseudomonas cremoris
WP 135288022.1 ligase-associated DNA damage response exonuclease Pseudomonas kairouanensis
WP 105696234.1 ligase-associated DNA damage response exonuclease Pseudomonas poae
WP 135308379.1 ligase-associated DNA damage response exonuclease Pseudomonas nabeulensis
WP 193864259.1 ligase-associated DNA damage response exonuclease Pseudomonas cyclaminis
WP 259192739.1 ligase-associated DNA damage response exonuclease Pseudomonas grimontii
WP 049709637.1 ligase-associated DNA damage response exonuclease Pseudomonas trivialis
WP 236378799.1 ligase-associated DNA damage response exonuclease Pseudomonas marginalis
WP 069672339.1 ligase-associated DNA damage response exonuclease Pseudomonas simiae
WP 122526801.1 ligase-associated DNA damage response exonuclease Pseudomonas salomonii
WP 177061422.1 ligase-associated DNA damage response exonuclease Pseudomonas reactans
WP 075946814.1 ligase-associated DNA damage response exonuclease Pseudomonas reinekei
WP 232776164.1 ligase-associated DNA damage response exonuclease Pseudomonas uvaldensis
WP 047701923.1 ligase-associated DNA damage response exonuclease Pseudomonas mediterranea
WP 175364215.1 ligase-associated DNA damage response exonuclease Pseudomonas corrugata
WP 305385687.1 ligase-associated DNA damage response exonuclease Pseudomonas hefeiensis
WP 264431609.1 ligase-associated DNA damage response exonuclease Pseudomonas agronomica
WP 232200716.1 ligase-associated DNA damage response exonuclease Pseudomonas bijieensis
WP 325918316.1 ligase-associated DNA damage response exonuclease Pseudomonas frederiksbergensis
WP 053118998.1 ligase-associated DNA damage response exonuclease Pseudomonas thivervalensis
WP 230152962.1 ligase-associated DNA damage response exonuclease Pseudomonas brassicacearum
WP 253471972.1 ligase-associated DNA damage response exonuclease Pseudomonas kilonensis
WP 204127579.1 ligase-associated DNA damage response exonuclease Pseudomonas ogarae
MDQ3202451.1 ligase-associated DNA damage response exonuclease Pseudomonadota bacterium
WP 090187629.1 ligase-associated DNA damage response exonuclease Pseudomonas arsenicoxydans
MET0774368.1 ligase-associated DNA damage response exonuclease Pseudomonas mandelii
WP 214381073.1 ligase-associated DNA damage response exonuclease Pseudomonas hormoni
WP 084321789.1 ligase-associated DNA damage response exonuclease Pseudomonas migulae
WP 307340765.1 ligase-associated DNA damage response exonuclease Pseudomonas lini
WP 339503485.1 ligase-associated DNA damage response exonuclease Pseudomonas silesiensis
WP 054615955.1 ligase-associated DNA damage response exonuclease Pseudomonas nunensis
WP 247288668.1 ligase-associated DNA damage response exonuclease Pseudomonas violetae
NWL21483.1 ligase-associated DNA damage response exonuclease Pseudomonas umsongensis
WP 305448456.1 ligase-associated DNA damage response exonuclease Pseudomonas cucumis
WP 093225957.1 ligase-associated DNA damage response exonuclease Pseudomonas vancouverensis
WP 103394470.1 ligase-associated DNA damage response exonuclease Pseudomonas laurylsulfativorans
WP 104450644.1 ligase-associated DNA damage response exonuclease Pseudomonas laurylsulfatiphila
WP 092272204.1 ligase-associated DNA damage response exonuclease Pseudomonas prosekii
WP 203421005.1 ligase-associated DNA damage response exonuclease Pseudomonas granadensis
WP 217832544.1 ligase-associated DNA damage response exonuclease Pseudomonas botevensis
WP 301623594.1 ligase-associated DNA damage response exonuclease Pseudomonas koreensis
WP 108559898.1 ligase-associated DNA damage response exonuclease Pseudomonas glycinae
WP 192563767.1 ligase-associated DNA damage response exonuclease Pseudomonas gozinkensis
WP 114881698.1 ligase-associated DNA damage response exonuclease Pseudomonas kribbensis
WP 065259279.1 ligase-associated DNA damage response exonuclease Pseudomonas bananamidigenes
WP 139055316.1 ligase-associated DNA damage response exonuclease Pseudomonas jessenii
WP 274658535.1 ligase-associated DNA damage response exonuclease Pseudomonas serboccidentalis
WP 322373874.1 ligase-associated DNA damage response exonuclease Pseudomonas monsensis
WP 310867441.1 ligase-associated DNA damage response exonuclease Pseudomonas baetica
WP 102902671.1 ligase-associated DNA damage response exonuclease Pseudomonas asplenii
WP 186612572.1 ligase-associated DNA damage response exonuclease Pseudomonas tensinigenes
KHA75204.1 exonuclease Pseudomonas chlororaphis
WP 186623652.1 ligase-associated DNA damage response exonuclease Pseudomonas zeae
WP 194933841.1 ligase-associated DNA damage response exonuclease Pseudomonas neuropathica
WP 186534315.1 ligase-associated DNA damage response exonuclease Pseudomonas khorasanensis
WP 166220920.1 ligase-associated DNA damage response exonuclease Pseudomonas atagonensis
WP 212545299.1 ligase-associated DNA damage response exonuclease Pseudomonas rustica
WP 252868900.1 ligase-associated DNA damage response exonuclease Pseudomonas siliginis
WP 125925233.1 ligase-associated DNA damage response exonuclease Pseudomonas moraviensis
WP 206420949.1 ligase-associated DNA damage response exonuclease Pseudomonas atacamensis
WP 186568195.1 ligase-associated DNA damage response exonuclease Pseudomonas iranensis

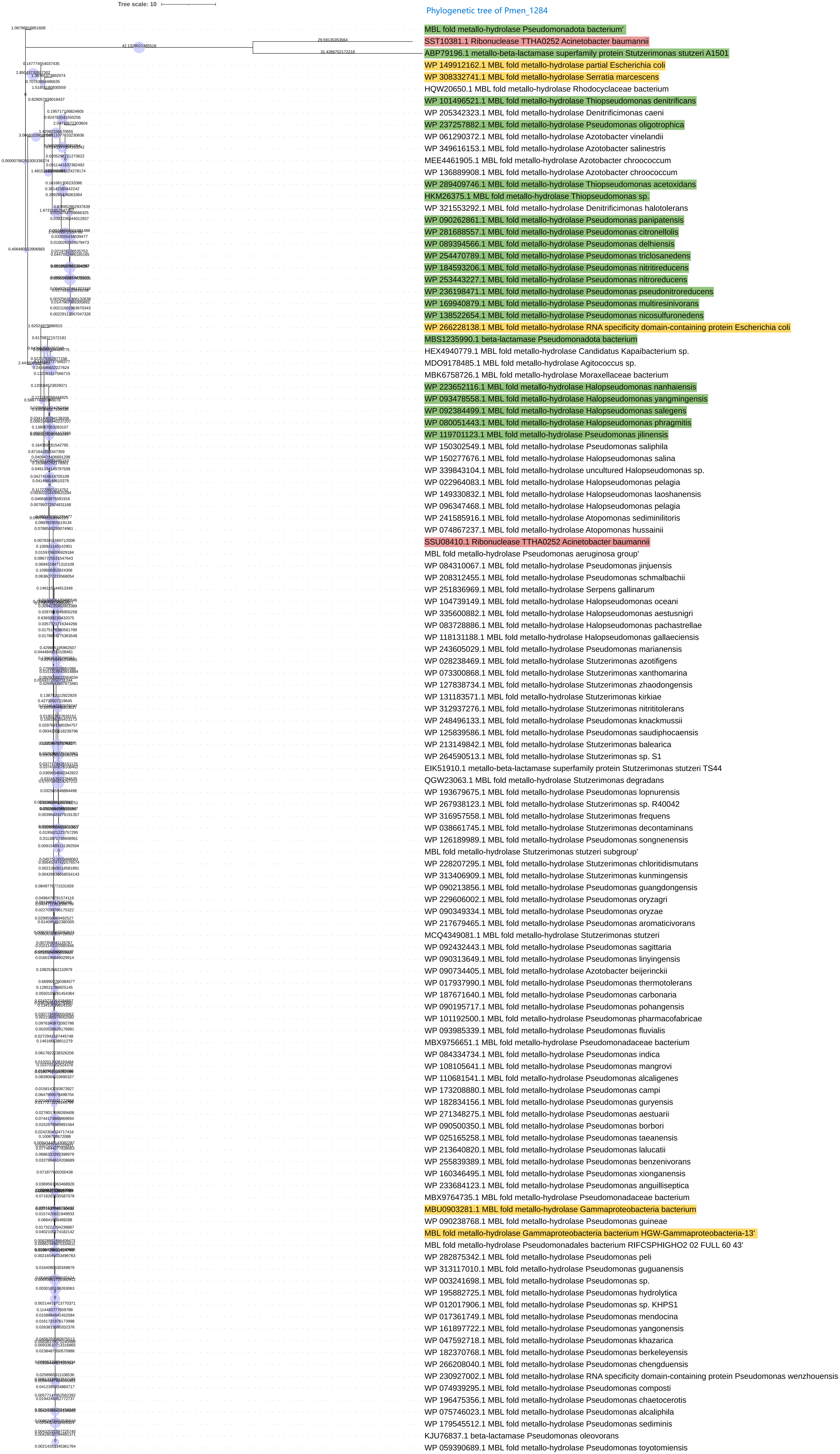
Tree scale: 10
Phylogenetic tree of Pmen_1284
MBL fold metallo-hydrolase Pseudomonadota bacterium'
SST10381.1 Ribonuclease TTHA0252 Acinetobacter baumannii
ABP79196.1 metallo-beta-lactamase superfamily protein Stutzerimonas stutzeri A1501
WP 149912162.1 MBL fold metallo-hydrolase partial Escherichia coli
WP 308332741.1 MBL fold metallo-hydrolase Serratia marcescens
HQW20650.1 MBL fold metallo-hydrolase Rhodocyclaceae bacterium
WP 101496521.1 MBL fold metallo-hydrolase Thiopseudomonas denitrificans
WP 205342323.1 MBL fold metallo-hydrolase Denitrificimonas caeni
WP 237257882.1 MBL fold metallo-hydrolase Pseudomonas oligotrophica
WP 061290372.1 MBL fold metallo-hydrolase Azotobacter vinelandii
WP 349616153.1 MBL fold metallo-hydrolase Azotobacter salinestris
MEE4461905.1 MBL fold metallo-hydrolase Azotobacter chroococcum
WP 136889908.1 MBL fold metallo-hydrolase Azotobacter chroococcum
WP 289409746.1 MBL fold metallo-hydrolase Thiopseudomonas acetoxidans
HKM26375.1 MBL fold metallo-hydrolase Thiopseudomonas sp.
WP 321553292.1 MBL fold metallo-hydrolase Denitrificimonas halotolerans
WP 090262861.1 MBL fold metallo-hydrolase Pseudomonas panipatensis
WP 281688557.1 MBL fold metallo-hydrolase Pseudomonas citronellolis
WP 089394566.1 MBL fold metallo-hydrolase Pseudomonas delhiensis
WP 254470789.1 MBL fold metallo-hydrolase Pseudomonas triclosanedens
WP 184593206.1 MBL fold metallo-hydrolase Pseudomonas nitritireducens
WP 253443227.1 MBL fold metallo-hydrolase Pseudomonas nitroreducens
WP 236198471.1 MBL fold metallo-hydrolase Pseudomonas pseudonitroreducens
WP 169940879.1 MBL fold metallo-hydrolase Pseudomonas multiresinivorans
WP 138522654.1 MBL fold metallo-hydrolase Pseudomonas nicosulfuronedens
WP 266228138.1 MBL fold metallo-hydrolase RNA specificity domain-containing protein Escherichia coli
MBS1235990.1 beta-lactamase Pseudomonadota bacterium
HEX4940779.1 MBL fold metallo-hydrolase Candidatus Kapaibacterium sp.
MDO9178485.1 MBL fold metallo-hydrolase Agitococcus sp.
MBK6758726.1 MBL fold metallo-hydrolase Moraxellaceae bacterium
WP 223652116.1 MBL fold metallo-hydrolase Halopseudomonas nanhaiensis
WP 093478558.1 MBL fold metallo-hydrolase Halopseudomonas yangmingensis
WP 092384499.1 MBL fold metallo-hydrolase Halopseudomonas salegens
WP 080051443.1 MBL fold metallo-hydrolase Halopseudomonas phragmitis
WP 119701123.1 MBL fold metallo-hydrolase Pseudomonas jilinensis
WP 150302549.1 MBL fold metallo-hydrolase Pseudomonas saliphila
WP 150277676.1 MBL fold metallo-hydrolase Halopseudomonas salina
WP 339843104.1 MBL fold metallo-hydrolase uncultured Halopseudomonas sp.
WP 022964083.1 MBL fold metallo-hydrolase Halopseudomonas pelagia
WP 149330832.1 MBL fold metallo-hydrolase Halopseudomonas laoshanensis
WP 096347468.1 MBL fold metallo-hydrolase Halopseudomonas pelagia
WP 241585916.1 MBL fold metallo-hydrolase Atopomonas sediminilitoris
WP 074867237.1 MBL fold metallo-hydrolase Atopomonas hussainii
SSU08410.1 Ribonuclease TTHA0252 Acinetobacter baumannii
MBL fold metallo-hydrolase Pseudomonas aeruginosa group'
WP 084310067.1 MBL fold metallo-hydrolase Pseudomonas jinjuensis
WP 208312455.1 MBL fold metallo-hydrolase Pseudomonas schmalbachii
WP 251836969.1 MBL fold metallo-hydrolase Serpens gallinarum
WP 104739149.1 MBL fold metallo-hydrolase Halopseudomonas oceani
WP 335600882.1 MBL fold metallo-hydrolase Halopseudomonas aestusnigri
WP 083728886.1 MBL fold metallo-hydrolase Halopseudomonas pachastrellae
WP 118131188.1 MBL fold metallo-hydrolase Halopseudomonas gallaeciensis
WP 243605029.1 MBL fold metallo-hydrolase Pseudomonas marianensis
WP 028238469.1 MBL fold metallo-hydrolase Stutzerimonas azotifigens
WP 073300868.1 MBL fold metallo-hydrolase Stutzerimonas xanthomarina
WP 127838734.1 MBL fold metallo-hydrolase Stutzerimonas zhaodongensis
WP 131183571.1 MBL fold metallo-hydrolase Stutzerimonas kirkiae
WP 312937276.1 MBL fold metallo-hydrolase Stutzerimonas nitrititolerans
WP 248496133.1 MBL fold metallo-hydrolase Pseudomonas knackmussii
WP 125839586.1 MBL fold metallo-hydrolase Pseudomonas saudiphocaensis
WP 213149842.1 MBL fold metallo-hydrolase Stutzerimonas balearica
WP 264590513.1 MBL fold metallo-hydrolase Stutzerimonas sp. S1
EIK51910.1 metallo-beta-lactamase superfamily protein Stutzerimonas stutzeri TS44
QGW23063.1 MBL fold metallo-hydrolase Stutzerimonas degradans
WP 193679675.1 MBL fold metallo-hydrolase Pseudomonas lopnurensis
WP 267938123.1 MBL fold metallo-hydrolase Stutzerimonas sp. R40042
WP 316957558.1 MBL fold metallo-hydrolase Stutzerimonas frequens
WP 038661745.1 MBL fold metallo-hydrolase Stutzerimonas decontaminans
WP 126189989.1 MBL fold metallo-hydrolase Pseudomonas songnenensis
MBL fold metallo-hydrolase Stutzerimonas stutzeri subgroup'
WP 228207295.1 MBL fold metallo-hydrolase Stutzerimonas chloritidismutans
WP 313406909.1 MBL fold metallo-hydrolase Stutzerimonas kunmingensis
WP 090213856.1 MBL fold metallo-hydrolase Pseudomonas guangdongensis
WP 229606002.1 MBL fold metallo-hydrolase Pseudomonas oryzagri
WP 090349334.1 MBL fold metallo-hydrolase Pseudomonas oryzae
WP 217679465.1 MBL fold metallo-hydrolase Pseudomonas aromaticivorans
MCQ4349081.1 MBL fold metallo-hydrolase Stutzerimonas stutzeri
WP 092432443.1 MBL fold metallo-hydrolase Pseudomonas sagittaria
WP 090313649.1 MBL fold metallo-hydrolase Pseudomonas linyingensis
WP 090734405.1 MBL fold metallo-hydrolase Azotobacter beijerinckii
WP 017937990.1 MBL fold metallo-hydrolase Pseudomonas thermotolerans
WP 187671640.1 MBL fold metallo-hydrolase Pseudomonas carbonaria
WP 090195717.1 MBL fold metallo-hydrolase Pseudomonas pohangensis
WP 101192500.1 MBL fold metallo-hydrolase Pseudomonas pharmacofabricae
WP 093985339.1 MBL fold metallo-hydrolase Pseudomonas fluvialis
MBX9756651.1 MBL fold metallo-hydrolase Pseudomonadaceae bacterium
WP 084334734.1 MBL fold metallo-hydrolase Pseudomonas indica
WP 108105641.1 MBL fold metallo-hydrolase Pseudomonas mangrovi
WP 110681541.1 MBL fold metallo-hydrolase Pseudomonas alcaligenes
WP 173208880.1 MBL fold metallo-hydrolase Pseudomonas campi
WP 182834156.1 MBL fold metallo-hydrolase Pseudomonas guryensis
WP 271348275.1 MBL fold metallo-hydrolase Pseudomonas aestuarii
WP 090500350.1 MBL fold metallo-hydrolase Pseudomonas borbori
WP 025165258.1 MBL fold metallo-hydrolase Pseudomonas taeanensis
WP 213640820.1 MBL fold metallo-hydrolase Pseudomonas lalucatii
WP 255839389.1 MBL fold metallo-hydrolase Pseudomonas benzenivorans
WP 160346495.1 MBL fold metallo-hydrolase Pseudomonas xionganensis
WP 233684123.1 MBL fold metallo-hydrolase Pseudomonas anguilliseptica
MBX9764735.1 MBL fold metallo-hydrolase Pseudomonadaceae bacterium
MBU0903281.1 MBL fold metallo-hydrolase Gammaproteobacteria bacterium
WP 090238768.1 MBL fold metallo-hydrolase Pseudomonas guineae
MBL fold metallo-hydrolase Gammaproteobacteria bacterium HGW-Gammaproteobacteria-13'
MBL fold metallo-hydrolase Pseudomonadales bacterium RIFCSPHIGHO2 02 FULL 60 43'
WP 282875342.1 MBL fold metallo-hydrolase Pseudomonas peli
WP 313117010.1 MBL fold metallo-hydrolase Pseudomonas guguanensis
WP 003241698.1 MBL fold metallo-hydrolase Pseudomonas sp.
WP 195882725.1 MBL fold metallo-hydrolase Pseudomonas hydrolytica
WP 012017906.1 MBL fold metallo-hydrolase Pseudomonas sp. KHPS1
WP 017361749.1 MBL fold metallo-hydrolase Pseudomonas mendocina
WP 161897722.1 MBL fold metallo-hydrolase Pseudomonas yangonensis
WP 047592718.1 MBL fold metallo-hydrolase Pseudomonas khazarica
WP 182370768.1 MBL fold metallo-hydrolase Pseudomonas berkeleyensis
WP 266208040.1 MBL fold metallo-hydrolase Pseudomonas chengduensis
WP 230927002.1 MBL fold metallo-hydrolase RNA specificity domain-containing protein Pseudomonas wenzhouensis
WP 074939295.1 MBL fold metallo-hydrolase Pseudomonas composti
WP 196475356.1 MBL fold metallo-hydrolase Pseudomonas chaetocerotis
WP 075746023.1 MBL fold metallo-hydrolase Pseudomonas alcaliphila
WP 179545512.1 MBL fold metallo-hydrolase Pseudomonas sediminis
KJU76837.1 beta-lactamase Pseudomonas oleovorans
WP 059390689.1 MBL fold metallo-hydrolase Pseudomonas toyotomiensis

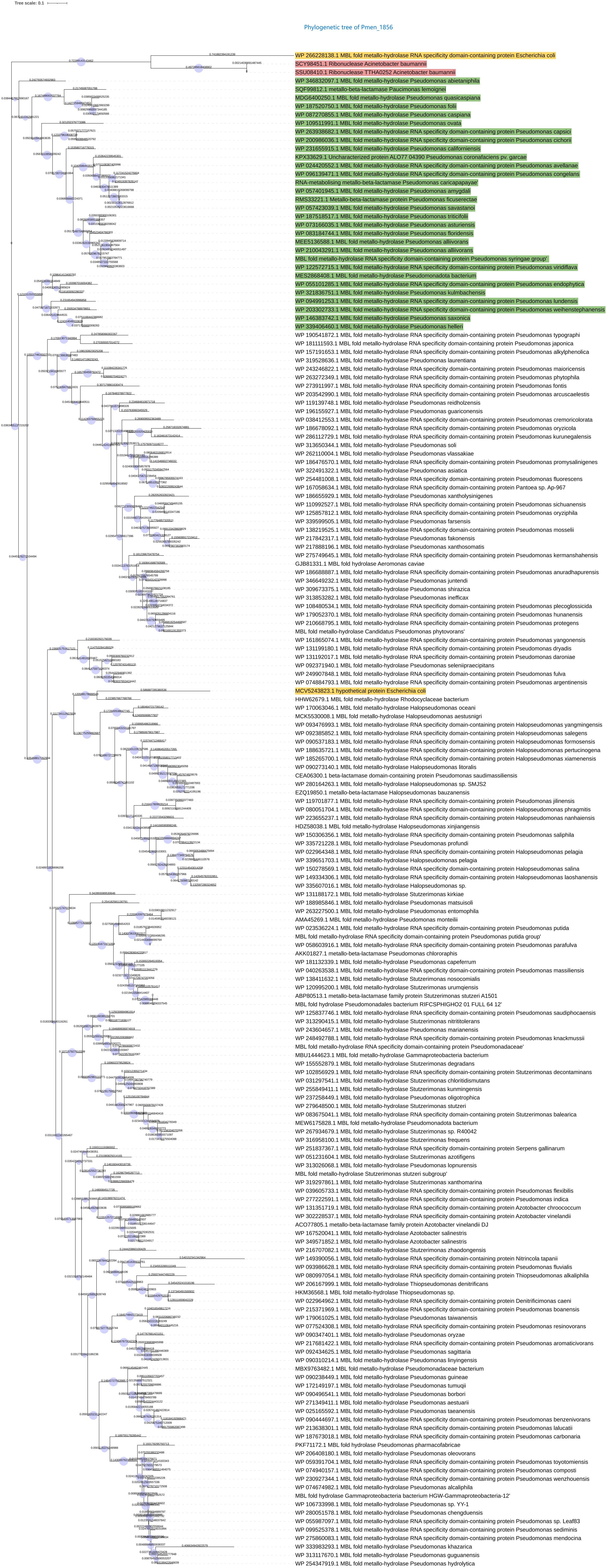
Tree scale: 0.1
Phylogenetic tree of Pmen_1856
WP 266228138.1 MBL fold metallo-hydrolase RNA specificity domain-containing protein Escherichia coli
SCY98451.1 Ribonuclease Acinetobacter baumannii
SSU08410.1 Ribonuclease TTHA0252 Acinetobacter baumannii
WP 346832097.1 MBL fold metallo-hydrolase Pseudomonas abietaniphila
SQF99812.1 metallo-beta-lactamase Paucimonas lemoignei
MDG6400250.1 MBL fold metallo-hydrolase Pseudomonas quasicaspiana
WP 187520750.1 MBL fold metallo-hydrolase Pseudomonas folii
WP 087270855.1 MBL fold metallo-hydrolase Pseudomonas caspiana
WP 109511991.1 MBL fold metallo-hydrolase Pseudomonas ovata
WP 263938682.1 MBL fold metallo-hydrolase RNA specificity domain-containing protein Pseudomonas capsici
WP 200986036.1 MBL fold metallo-hydrolase RNA specificity domain-containing protein Pseudomonas cichorii
WP 231655915.1 MBL fold metallo-hydrolase Pseudomonas californiensis
KPX33629.1 Uncharacterized protein ALO77 04390 Pseudomonas coronafaciens pv. garcae
WP 024420552.1 MBL fold metallo-hydrolase RNA specificity domain-containing protein Pseudomonas avellanae
WP 096139471.1 MBL fold metallo-hydrolase RNA specificity domain-containing protein Pseudomonas congelans
RNA-metabolising metallo-beta-lactamase Pseudomonas caricapapayae'
WP 057401945.1 MBL fold metallo-hydrolase Pseudomonas amygdali
RMS33221.1 Metallo-beta-lactamase protein Pseudomonas ficuserectae
WP 057423039.1 MBL fold metallo-hydrolase Pseudomonas savastanoi
WP 187518517.1 MBL fold metallo-hydrolase Pseudomonas triticifolii
WP 073166035.1 MBL fold metallo-hydrolase Pseudomonas asturiensis
WP 083184744.1 MBL fold metallo-hydrolase Pseudomonas floridensis
MEE5136588.1 MBL fold metallo-hydrolase Pseudomonas alliivorans
WP 210043291.1 MBL fold metallo-hydrolase Pseudomonas alliivorans
MBL fold metallo-hydrolase RNA specificity domain-containing protein Pseudomonas syringae group'
WP 122572715.1 MBL fold metallo-hydrolase Pseudomonas viridiflava
MES2868408.1 MBL fold metallo-hydrolase Pseudomonadota bacterium
WP 055101285.1 MBL fold metallo-hydrolase RNA specificity domain-containing protein Pseudomonas endophytica
WP 321836751.1 MBL fold metallo-hydrolase Pseudomonas kulmbachensis
WP 094991253.1 MBL fold metallo-hydrolase RNA specificity domain-containing protein Pseudomonas lundensis
WP 203302733.1 MBL fold metallo-hydrolase RNA specificity domain-containing protein Pseudomonas weihenstephanensis
WP 146383742.1 MBL fold metallo-hydrolase Pseudomonas saxonica
WP 339406460.1 MBL fold metallo-hydrolase Pseudomonas helleri
WP 190541872.1 MBL fold metallo-hydrolase RNA specificity domain-containing protein Pseudomonas typographi
WP 181111593.1 MBL fold metallo-hydrolase RNA specificity domain-containing protein Pseudomonas japonica
WP 157191653.1 MBL fold metallo-hydrolase RNA specificity domain-containing protein Pseudomonas alkylphenolica
WP 319528636.1 MBL fold metallo-hydrolase Pseudomonas laurentiana
WP 243246822.1 MBL fold metallo-hydrolase RNA specificity domain-containing protein Pseudomonas maioricensis
WP 263272349.1 MBL fold metallo-hydrolase RNA specificity domain-containing protein Pseudomonas phytophila
WP 273911997.1 MBL fold metallo-hydrolase RNA specificity domain-containing protein Pseudomonas fontis
WP 203542990.1 MBL fold metallo-hydrolase RNA specificity domain-containing protein Pseudomonas arcuscaelestis
WP 119139748.1 MBL fold metallo-hydrolase Pseudomonas reidholzensis
WP 196155927.1 MBL fold metallo-hydrolase Pseudomonas guariconensis
WP 038412553.1 MBL fold metallo-hydrolase RNA specificity domain-containing protein Pseudomonas cremoricolorata
WP 186678092.1 MBL fold metallo-hydrolase RNA specificity domain-containing protein Pseudomonas oryzicola
WP 286112729.1 MBL fold metallo-hydrolase RNA specificity domain-containing protein Pseudomonas kurunegalensis
WP 313650344.1 MBL fold metallo-hydrolase Pseudomonas soli
WP 262110004.1 MBL fold metallo-hydrolase Pseudomonas vlassakiae
WP 186476570.1 MBL fold metallo-hydrolase RNA specificity domain-containing protein Pseudomonas promysalinigenes
WP 322491322.1 MBL fold metallo-hydrolase Pseudomonas asiatica
WP 254481008.1 MBL fold metallo-hydrolase RNA specificity domain-containing protein Pseudomonas fluorescens
WP 167058634.1 MBL fold metallo-hydrolase RNA specificity domain-containing protein Pantoea sp. Ap-967
WP 186655929.1 MBL fold metallo-hydrolase Pseudomonas xantholysinigenes
WP 110992527.1 MBL fold metallo-hydrolase RNA specificity domain-containing protein Pseudomonas sichuanensis
WP 325857812.1 MBL fold metallo-hydrolase RNA specificity domain-containing protein Pseudomonas oryziphila
WP 339599505.1 MBL fold metallo-hydrolase Pseudomonas farsensis
WP 138219525.1 MBL fold metallo-hydrolase RNA specificity domain-containing protein Pseudomonas mosselii
WP 217842317.1 MBL fold metallo-hydrolase Pseudomonas fakonensis
WP 217888196.1 MBL fold metallo-hydrolase Pseudomonas xanthosomatis
WP 275749645.1 MBL fold metallo-hydrolase RNA specificity domain-containing protein Pseudomonas kermanshahensis
GJB81331.1 MBL fold hydrolase Aeromonas caviae
WP 186688887.1 MBL fold metallo-hydrolase RNA specificity domain-containing protein Pseudomonas anuradhapurensis
WP 346649232.1 MBL fold metallo-hydrolase Pseudomonas juntendi
WP 309673375.1 MBL fold metallo-hydrolase Pseudomonas shirazica
WP 313853282.1 MBL fold metallo-hydrolase Pseudomonas inefficax
WP 108480534.1 MBL fold metallo-hydrolase RNA specificity domain-containing protein Pseudomonas plecoglossicida
WP 179052370.1 MBL fold metallo-hydrolase RNA specificity domain-containing protein Pseudomonas hunanensis
WP 210668795.1 MBL fold metallo-hydrolase RNA specificity domain-containing protein Pseudomonas protegens
MBL fold metallo-hydrolase Candidatus Pseudomonas phytovorans'
WP 161865074.1 MBL fold metallo-hydrolase RNA specificity domain-containing protein Pseudomonas yangonensis
WP 131199180.1 MBL fold metallo-hydrolase RNA specificity domain-containing protein Pseudomonas dryadis
WP 131192017.1 MBL fold metallo-hydrolase RNA specificity domain-containing protein Pseudomonas daroniae
WP 092371940.1 MBL fold metallo-hydrolase Pseudomonas seleniipraecipitans
WP 249907848.1 MBL fold metallo-hydrolase RNA specificity domain-containing protein Pseudomonas fulva
WP 074884793.1 MBL fold metallo-hydrolase RNA specificity domain-containing protein Pseudomonas argentinensis
MCV5243823.1 hypothetical protein Escherichia coli
HHW62679.1 MBL fold metallo-hydrolase Rhodocyclaceae bacterium
WP 170063046.1 MBL fold metallo-hydrolase Halopseudomonas oceani
MCK5530008.1 MBL fold metallo-hydrolase Halopseudomonas aestusnigri
WP 093476993.1 MBL fold metallo-hydrolase RNA specificity domain-containing protein Halopseudomonas yangmingensis
WP 092385852.1 MBL fold metallo-hydrolase RNA specificity domain-containing protein Halopseudomonas salegens
WP 090537183.1 MBL fold metallo-hydrolase RNA specificity domain-containing protein Halopseudomonas formosensis
WP 188635721.1 MBL fold metallo-hydrolase RNA specificity domain-containing protein Halopseudomonas pertucinogena
WP 185265700.1 MBL fold metallo-hydrolase RNA specificity domain-containing protein Halopseudomonas xiamenensis
WP 090273140.1 MBL fold metallo-hydrolase Halopseudomonas litoralis
CEA06300.1 beta-lactamase domain-containing protein Pseudomonas saudimassiliensis
WP 280164263.1 MBL fold metallo-hydrolase Halopseudomonas sp. SMJS2
EZQ19850.1 beta-lactamase Halopseudomonas bauzanensis
WP 119701877.1 MBL fold metallo-hydrolase RNA specificity domain-containing protein Pseudomonas jilinensis
WP 080051704.1 MBL fold metallo-hydrolase RNA specificity domain-containing protein Halopseudomonas phragmitis
WP 223655237.1 MBL fold metallo-hydrolase RNA specificity domain-containing protein Halopseudomonas nanhaiensis
HDZ58038.1 MBL fold metallo-hydrolase Halopseudomonas xinjiangensis
WP 150306356.1 MBL fold metallo-hydrolase RNA specificity domain-containing protein Pseudomonas saliphila
WP 335721228.1 MBL fold metallo-hydrolase Pseudomonas profundi
WP 022964348.1 MBL fold metallo-hydrolase RNA specificity domain-containing protein Pseudomonas pelagia
WP 339651703.1 MBL fold metallo-hydrolase Halopseudomonas pelagia
WP 150278569.1 MBL fold metallo-hydrolase RNA specificity domain-containing protein Pseudomonas salina
WP 149334306.1 MBL fold metallo-hydrolase RNA specificity domain-containing protein Pseudomonas laoshanensis
WP 335607016.1 MBL fold metallo-hydrolase Halopseudomonas sp.
WP 131188172.1 MBL fold metallo-hydrolase Stutzerimonas nirkuisei
WP 188985846.1 MBL fold metallo-hydrolase Pseudomonas mediterranea
WP 263227500.1 MBL fold metallo-hydrolase Pseudomonas entomophila
AMA45269.1 MBL fold metallo-hydrolase Pseudomonas monteilii
WP 023536224.1 MBL fold metallo-hydrolase RNA specificity domain-containing protein Pseudomonas putida
MBL fold metallo-hydrolase RNA specificity domain-containing protein Pseudomonas putida group'
WP 058603916.1 MBL fold metallo-hydrolase RNA specificity domain-containing protein Pseudomonas parafulva
AKK01827.1 metallo-beta-lactamase Pseudomonas chlororaphis
WP 181132339.1 MBL fold metallo-hydrolase Pseudomonas capeferrum
WP 040263538.1 MBL fold metallo-hydrolase RNA specificity domain-containing protein Pseudomonas massiliensis
WP 138411632.1 MBL fold metallo-hydrolase Stutzerimonas nosocomialis
WP 120995200.1 MBL fold metallo-hydrolase Stutzerimonas urumqiensis
ABP80513.1 metallo-beta-lactamase family protein Stutzerimonas stutzeri A1501
MBL fold hydrolase Pseudomonadales bacterium RIFCSPHIGHO2 01 FULL 64 12'
WP 125837746.1 MBL fold metallo-hydrolase RNA specificity domain-containing protein Pseudomonas saudiphocaensis
WP 313290415.1 MBL fold metallo-hydrolase Stutzerimonas nitrititolerans
WP 243604657.1 MBL fold metallo-hydrolase Pseudomonas marianensis
WP 248492788.1 MBL fold metallo-hydrolase RNA specificity domain-containing protein Pseudomonas knackmussii
MBL fold metallo-hydrolase RNA specificity domain-containing protein Pseudomonadaceae'
MBU1444623.1 MBL fold metallo-hydrolase Gammaproteobacteria bacterium
WP 155552879.1 MBL fold metallo-hydrolase Stutzerimonas degradans
WP 102856929.1 MBL fold metallo-hydrolase RNA specificity domain-containing protein Stutzerimonas decontaminans
WP 031297541.1 MBL fold metallo-hydrolase Stutzerimonas chloritidismutans
WP 255849411.1 MBL fold metallo-hydrolase Stutzerimonas kunmingensis
WP 237258449.1 MBL fold metallo-hydrolase Stutzerimonas oligotrophica
WP 279648500.1 MBL fold metallo-hydrolase Stutzerimonas stutzeri
WP 083675041.1 MBL fold metallo-hydrolase RNA specificity domain-containing protein Stutzerimonas balearica
MEW6175828.1 MBL fold metallo-hydrolase Pseudomonadota bacterium
WP 267934679.1 MBL fold metallo-hydrolase Pseudomonas sp. R40042
WP 316958100.1 MBL fold metallo-hydrolase Stutzerimonas frequens
WP 251837367.1 MBL fold metallo-hydrolase RNA specificity domain-containing protein Serpens gallinarum
WP 051231604.1 MBL fold metallo-hydrolase Stutzerimonas azotifigens
WP 313026068.1 MBL fold metallo-hydrolase Pseudomonas lopnurensis
MBL fold metallo-hydrolase Stutzerimonas stutzeri subgroup'
WP 319297861.1 MBL fold metallo-hydrolase Stutzerimonas xanthomarina
WP 039605733.1 MBL fold metallo-hydrolase RNA specificity domain-containing protein Pseudomonas flexibilis
WP 277222591.1 MBL fold metallo-hydrolase RNA specificity domain-containing protein Pseudomonas indica
WP 131351719.1 MBL fold metallo-hydrolase RNA specificity domain-containing protein Azotobacter chroococcum
WP 302228537.1 MBL fold metallo-hydrolase RNA specificity domain-containing protein Azotobacter vinelandii
ACO77805.1 metallo-beta-lactamase family protein Azotobacter vinelandii DJ
WP 167520041.1 MBL fold metallo-hydrolase Azotobacter salinestris
WP 349571852.1 MBL fold metallo-hydrolase Azotobacter salinestris
WP 216707082.1 MBL fold metallo-hydrolase Stutzerimonas zhaodongensis
WP 149390056.1 MBL fold metallo-hydrolase RNA specificity domain-containing protein Nitrincola tapanii
WP 093986628.1 MBL fold metallo-hydrolase RNA specificity domain-containing protein Pseudomonas fluvialis
WP 080997054.1 MBL fold metallo-hydrolase RNA specificity domain-containing protein Thiopseudomonas alkaliphila
WP 206167999.1 MBL fold metallo-hydrolase Thiopseudomonas denitrificans
HKM36568.1 MBL fold metallo-hydrolase Thiopseudomonas sp.
WP 022964962.1 MBL fold metallo-hydrolase RNA specificity domain-containing protein Denitrificimonas caeni
WP 215371969.1 MBL fold metallo-hydrolase RNA specificity domain-containing protein Pseudomonas boanensis
WP 179061025.1 MBL fold metallo-hydrolase Pseudomonas taiwanensis
WP 077524308.1 MBL fold metallo-hydrolase RNA specificity domain-containing protein Pseudomonas resinovorans
WP 090347401.1 MBL fold metallo-hydrolase Pseudomonas oryzae
WP 217681422.1 MBL fold metallo-hydrolase RNA specificity domain-containing protein Pseudomonas aromaticivorans
WP 092434625.1 MBL fold metallo-hydrolase Pseudomonas sagittaria
WP 090310214.1 MBL fold metallo-hydrolase Pseudomonas linyingensis
MBX9763482.1 MBL fold metallo-hydrolase Pseudomonadaceae bacterium
WP 090238449.1 MBL fold metallo-hydrolase Pseudomonas guineae
WP 172149197.1 MBL fold metallo-hydrolase Pseudomonas tumuqii
WP 090496541.1 MBL fold metallo-hydrolase Pseudomonas borbori
WP 271349411.1 MBL fold metallo-hydrolase Pseudomonas aestusnigri
WP 025165592.1 MBL fold metallo-hydrolase Pseudomonas taeanensis
WP 090444697.1 MBL fold metallo-hydrolase RNA specificity domain-containing protein Pseudomonas benzenivorans
WP 213638301.1 MBL fold metallo-hydrolase RNA specificity domain-containing protein Pseudomonas lalucatii
WP 187673018.1 MBL fold metallo-hydrolase RNA specificity domain-containing protein Pseudomonas carbonaria
PKF71172.1 MBL fold hydrolase Pseudomonas pharmacofabricae
WP 206408180.1 MBL fold metallo-hydrolase Pseudomonas oleovorans
WP 059391704.1 MBL fold metallo-hydrolase RNA specificity domain-containing protein Pseudomonas toyotomiensis
WP 074940157.1 MBL fold metallo-hydrolase RNA specificity domain-containing protein Pseudomonas composti
WP 230927344.1 MBL fold metallo-hydrolase RNA specificity domain-containing protein Pseudomonas wenzhouensis
WP 074674982.1 MBL fold metallo-hydrolase Pseudomonas alcaliphila
MBL fold hydrolase Gammaproteobacteria bacterium HGW-Gammaproteobacteria-12'
WP 106733998.1 MBL fold metallo-hydrolase Pseudomonas sp. YY-1
WP 280051578.1 MBL fold metallo-hydrolase Pseudomonas chengduensis
WP 055987097.1 MBL fold metallo-hydrolase RNA specificity domain-containing protein Pseudomonas sp. Leaf83
WP 099525378.1 MBL fold metallo-hydrolase RNA specificity domain-containing protein Pseudomonas sediminis
WP 275860083.1 MBL fold metallo-hydrolase RNA specificity domain-containing protein Pseudomonas mendocina
WP 333983293.1 MBL fold metallo-hydrolase Pseudomonas khazarica
WP 313176670.1 MBL fold metallo-hydrolase Pseudomonas guguanensis
WP 254347919.1 MBL fold metallo-hydrolase Pseudomonas hydrolytica

Phylogenetic tree of rpoZ

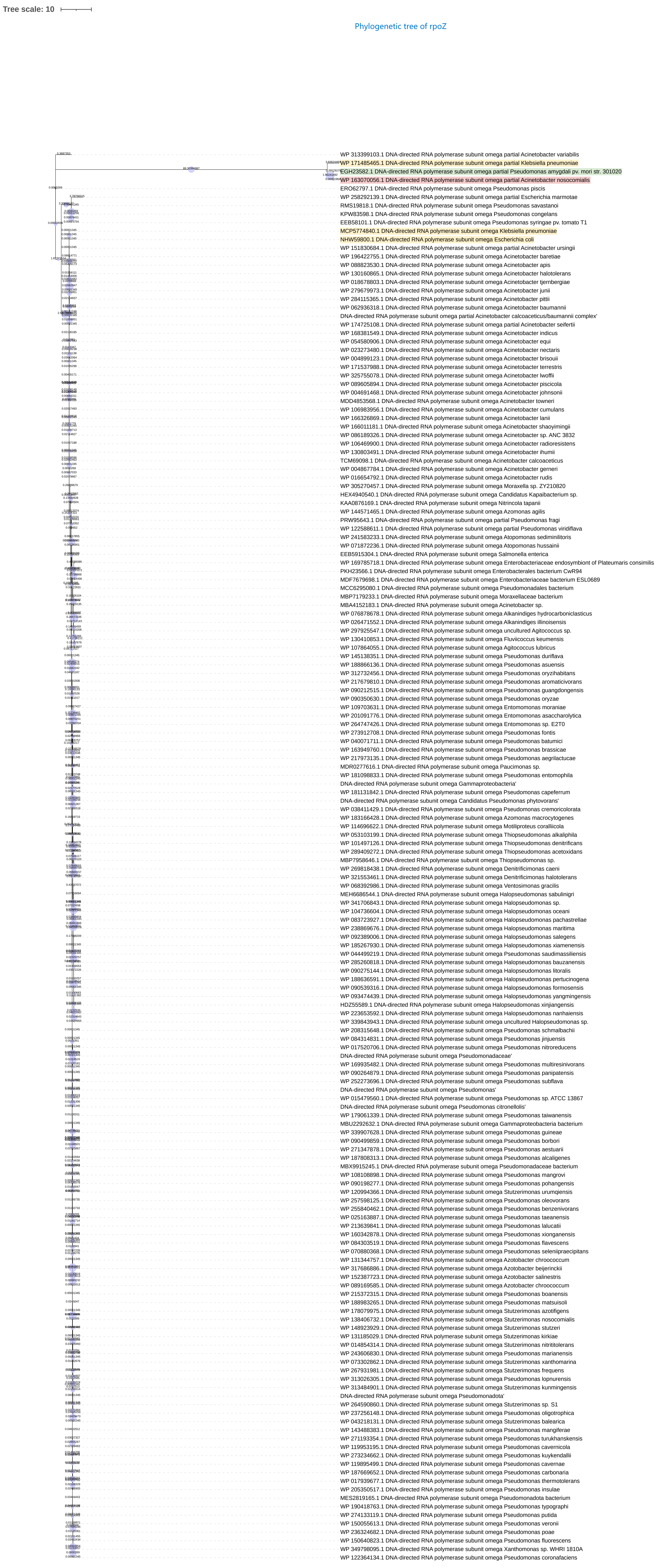

Phylogenetic tree of MDS_1962

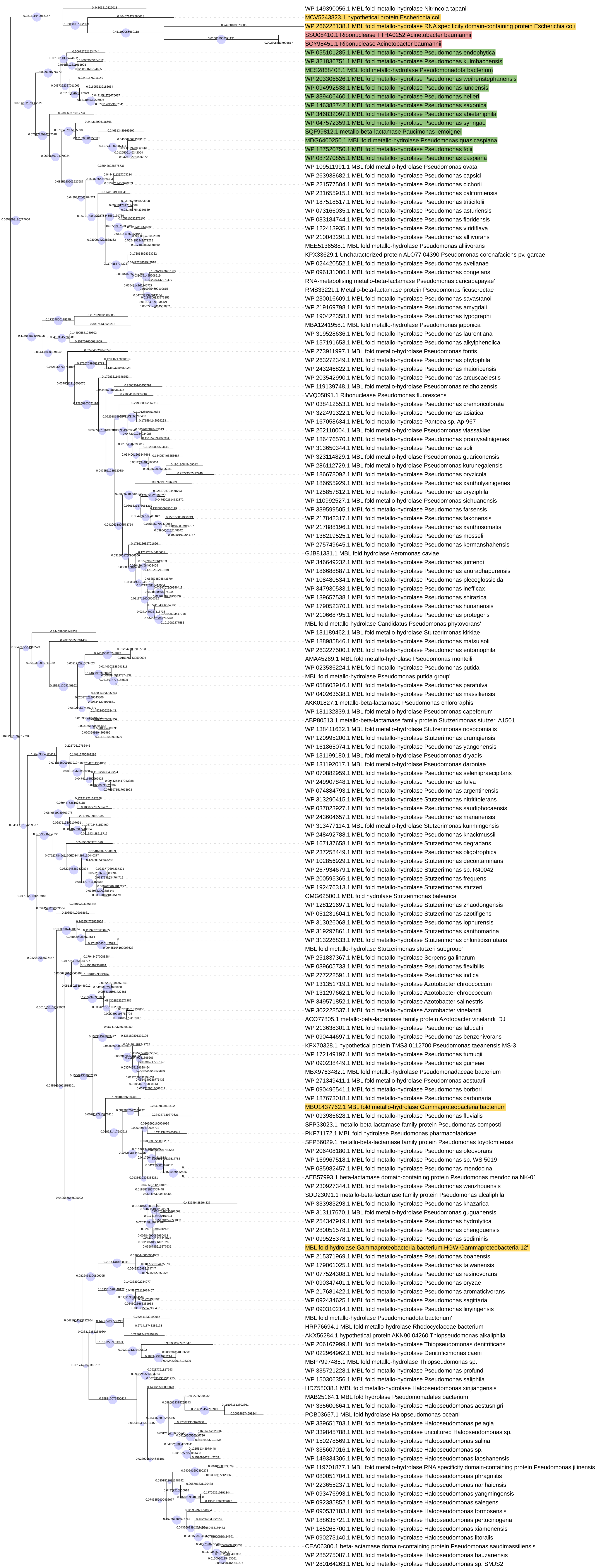

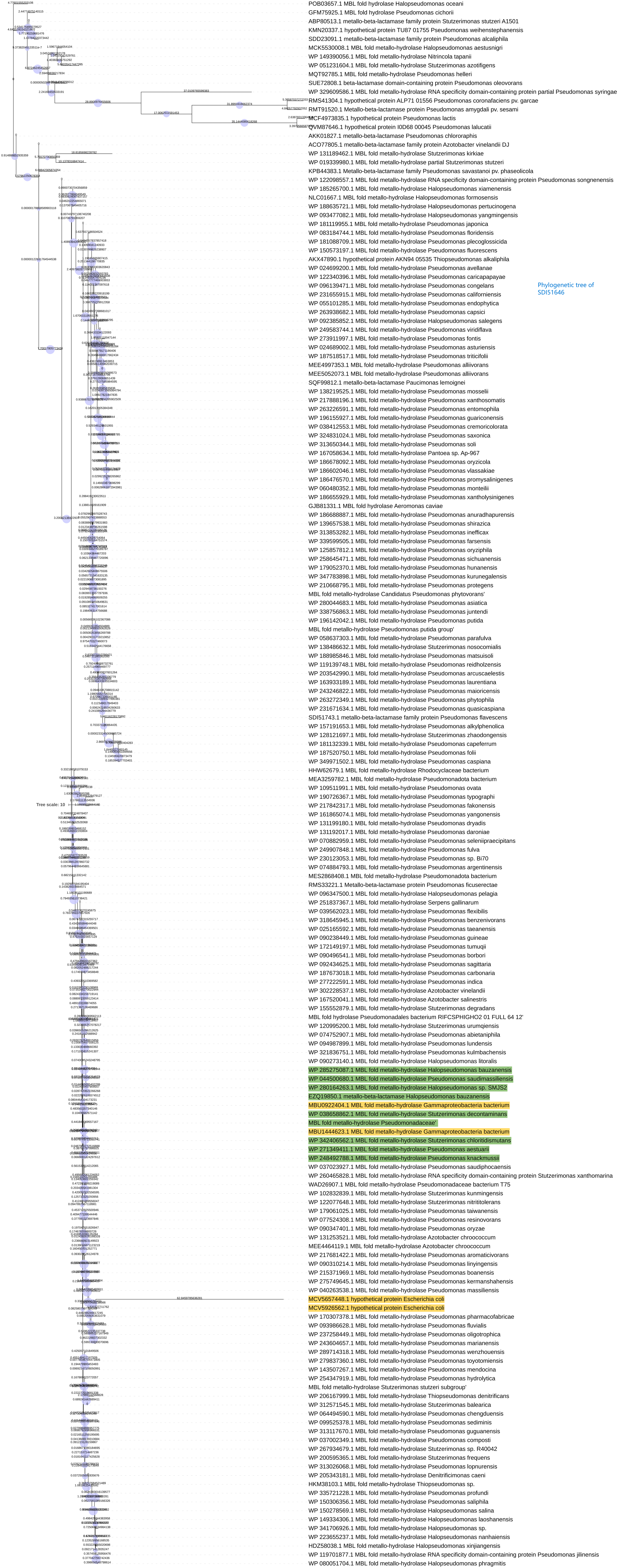


Phylogenetic tree of SDI51646

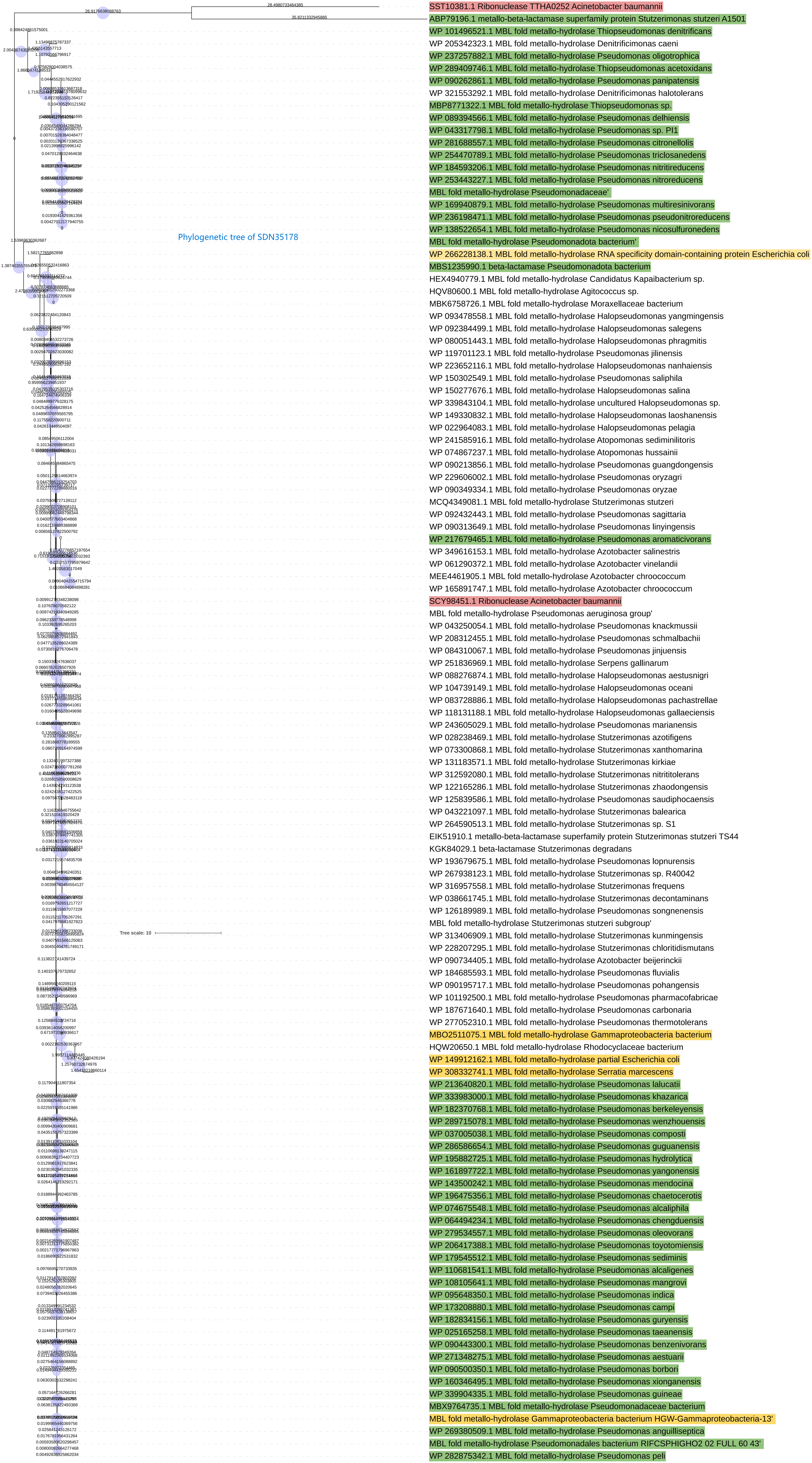
Phylogenetic tree of SDN35178
Tree scale: 10
SST10381.1 Ribonuclease TTHA0252 Acinetobacter baumannii
ABP79196.1 metallo-beta-lactamase superfamily protein Stutzerimonas stutzeri A1501
WP 101496521.1 MBL fold metallo-hydrolase Thiopseudomonas denitrificans
WP 205342323.1 MBL fold metallo-hydrolase Denitrificimonas caeni
WP 237257882.1 MBL fold metallo-hydrolase Pseudomonas oligotrophica
WP 289409746.1 MBL fold metallo-hydrolase Thiopseudomonas acetoxidans
WP 090262861.1 MBL fold metallo-hydrolase Pseudomonas panipatensis
WP 321553292.1 MBL fold metallo-hydrolase Denitrificimonas halotolerans
MBP8771322.1 MBL fold metallo-hydrolase Thiopseudomonas sp.
WP 089394566.1 MBL fold metallo-hydrolase Pseudomonas delhiensis
WP 043317798.1 MBL fold metallo-hydrolase Pseudomonas sp. PI1
WP 281688557.1 MBL fold metallo-hydrolase Pseudomonas citronellolis
WP 254470789.1 MBL fold metallo-hydrolase Pseudomonas triclosanedens
WP 184593206.1 MBL fold metallo-hydrolase Pseudomonas nitritireducens
WP 253443227.1 MBL fold metallo-hydrolase Pseudomonas nitroreducens
MBL fold metallo-hydrolase Pseudomonadaceae'
WP 169940879.1 MBL fold metallo-hydrolase Pseudomonas multiresinivorans
WP 236198471.1 MBL fold metallo-hydrolase Pseudomonas pseudonitroreducens
WP 138522654.1 MBL fold metallo-hydrolase Pseudomonas nicosulfuronedens
MBL fold metallo-hydrolase Pseudomonadota bacterium'
WP 266228138.1 MBL fold metallo-hydrolase RNA specificity domain-containing protein Escherichia coli
MBS1235990.1 beta-lactamase Pseudomonadota bacterium
HEX4940779.1 MBL fold metallo-hydrolase Candidatus Kapaibacterium sp.
HQV80600.1 MBL fold metallo-hydrolase Agitococcus sp.
MBK6758726.1 MBL fold metallo-hydrolase Moraxellaceae bacterium
WP 093478558.1 MBL fold metallo-hydrolase Halopseudomonas yangmingensis
WP 092384499.1 MBL fold metallo-hydrolase Halopseudomonas salegens
WP 080051443.1 MBL fold metallo-hydrolase Halopseudomonas phragmitis
WP 119701123.1 MBL fold metallo-hydrolase Pseudomonas jilinensis
WP 223652116.1 MBL fold metallo-hydrolase Halopseudomonas nanhaiensis
WP 150302549.1 MBL fold metallo-hydrolase Pseudomonas saliphila
WP 150277676.1 MBL fold metallo-hydrolase Halopseudomonas salina
WP 339843104.1 MBL fold metallo-hydrolase uncultured Halopseudomonas sp.
WP 149330832.1 MBL fold metallo-hydrolase Halopseudomonas laoshanensis
WP 022964083.1 MBL fold metallo-hydrolase Halopseudomonas pelagia
WP 241585916.1 MBL fold metallo-hydrolase Atopomonas sediminilitoris
WP 074867237.1 MBL fold metallo-hydrolase Atopomonas hussainii
WP 090213856.1 MBL fold metallo-hydrolase Pseudomonas guangdongensis
WP 229606002.1 MBL fold metallo-hydrolase Pseudomonas oryzagri
WP 090349334.1 MBL fold metallo-hydrolase Pseudomonas oryzae
MCQ4349081.1 MBL fold metallo-hydrolase Stutzerimonas stutzeri
WP 092432443.1 MBL fold metallo-hydrolase Pseudomonas sagittaria
WP 090313649.1 MBL fold metallo-hydrolase Pseudomonas linyingensis
WP 217679465.1 MBL fold metallo-hydrolase Pseudomonas aromaticivorans
WP 349616153.1 MBL fold metallo-hydrolase Azotobacter salinestris
WP 061290372.1 MBL fold metallo-hydrolase Azotobacter vinelandii
MEE4461905.1 MBL fold metallo-hydrolase Azotobacter chroococcum
WP 165891747.1 MBL fold metallo-hydrolase Azotobacter chroococcum
SCY98451.1 Ribonuclease Acinetobacter baumannii
MBL fold metallo-hydrolase Pseudomonas aeruginosa group'
WP 043250054.1 MBL fold metallo-hydrolase Pseudomonas knackmussii
WP 208312455.1 MBL fold metallo-hydrolase Pseudomonas schmalbachii
WP 084310067.1 MBL fold metallo-hydrolase Pseudomonas jinjuensis
WP 251836969.1 MBL fold metallo-hydrolase Serpens gallinarum
WP 088276874.1 MBL fold metallo-hydrolase Halopseudomonas aestusnigri
WP 104739149.1 MBL fold metallo-hydrolase Halopseudomonas oceani
WP 083728886.1 MBL fold metallo-hydrolase Halopseudomonas pachastrellae
WP 118131188.1 MBL fold metallo-hydrolase Halopseudomonas gallaeciensis
WP 243605029.1 MBL fold metallo-hydrolase Pseudomonas marianensis
WP 028238469.1 MBL fold metallo-hydrolase Stutzerimonas azotifigens
WP 073300868.1 MBL fold metallo-hydrolase Stutzerimonas xanthomarina
WP 131183571.1 MBL fold metallo-hydrolase Stutzerimonas kirkiae
WP 312592080.1 MBL fold metallo-hydrolase Stutzerimonas nitrititolerans
WP 122165286.1 MBL fold metallo-hydrolase Stutzerimonas zhaodongensis
WP 125839586.1 MBL fold metallo-hydrolase Pseudomonas saudiphocaensis
WP 043221097.1 MBL fold metallo-hydrolase Stutzerimonas balearica
WP 264590513.1 MBL fold metallo-hydrolase Stutzerimonas sp. S1
EIK51910.1 metallo-beta-lactamase superfamily protein Stutzerimonas stutzeri TS44
KGK84029.1 beta-lactamase Stutzerimonas degradans
WP 193679675.1 MBL fold metallo-hydrolase Pseudomonas lopnurensis
WP 267938123.1 MBL fold metallo-hydrolase Stutzerimonas sp. R40042
WP 316957558.1 MBL fold metallo-hydrolase Stutzerimonas frequens
WP 038661745.1 MBL fold metallo-hydrolase Stutzerimonas decontaminans
WP 126189989.1 MBL fold metallo-hydrolase Pseudomonas songnenensis
MBL fold metallo-hydrolase Stutzerimonas stutzeri subgroup'
WP 313406909.1 MBL fold metallo-hydrolase Stutzerimonas kunmingensis
WP 228207295.1 MBL fold metallo-hydrolase Stutzerimonas chloritidismutans
WP 090734405.1 MBL fold metallo-hydrolase Azotobacter beijerinckii
WP 184685593.1 MBL fold metallo-hydrolase Pseudomonas fluvialis
WP 090195717.1 MBL fold metallo-hydrolase Pseudomonas pohangensis
WP 101192500.1 MBL fold metallo-hydrolase Pseudomonas pharmacofabricae
WP 187671640.1 MBL fold metallo-hydrolase Pseudomonas carbonaria
WP 277052310.1 MBL fold metallo-hydrolase Pseudomonas thermotolerans
MBO2511075.1 MBL fold metallo-hydrolase Gammaproteobacteria bacterium
HQW20650.1 MBL fold metallo-hydrolase Rhodocyclaceae bacterium
WP 149912162.1 MBL fold metallo-hydrolase partial Escherichia coli
WP 308332741.1 MBL fold metallo-hydrolase Serratia marcescens
WP 213640820.1 MBL fold metallo-hydrolase Pseudomonas lalucatii
WP 333983000.1 MBL fold metallo-hydrolase Pseudomonas khazarica
WP 182370768.1 MBL fold metallo-hydrolase Pseudomonas berkeleyensis
WP 289715078.1 MBL fold metallo-hydrolase Pseudomonas wenzhouensis
WP 037005038.1 MBL fold metallo-hydrolase Pseudomonas composti
WP 286586654.1 MBL fold metallo-hydrolase Pseudomonas guguanensis
WP 195882725.1 MBL fold metallo-hydrolase Pseudomonas hydrolytica
WP 161897722.1 MBL fold metallo-hydrolase Pseudomonas yangonensis
WP 143500242.1 MBL fold metallo-hydrolase Pseudomonas mendocina
WP 196475356.1 MBL fold metallo-hydrolase Pseudomonas chaetocerotis
WP 074675548.1 MBL fold metallo-hydrolase Pseudomonas alcaliphila
WP 064494234.1 MBL fold metallo-hydrolase Pseudomonas chengduensis
WP 279534557.1 MBL fold metallo-hydrolase Pseudomonas oleovorans
WP 206417388.1 MBL fold metallo-hydrolase Pseudomonas toyotomiensis
WP 179545512.1 MBL fold metallo-hydrolase Pseudomonas sediminis
WP 110681541.1 MBL fold metallo-hydrolase Pseudomonas alcaligenes
WP 108105641.1 MBL fold metallo-hydrolase Pseudomonas mangrovi
WP 095648350.1 MBL fold metallo-hydrolase Pseudomonas indica
WP 173208880.1 MBL fold metallo-hydrolase Pseudomonas campi
WP 182834156.1 MBL fold metallo-hydrolase Pseudomonas guryensis
WP 025165258.1 MBL fold metallo-hydrolase Pseudomonas taeanensis
WP 090443300.1 MBL fold metallo-hydrolase Pseudomonas benzenivorans
WP 271348275.1 MBL fold metallo-hydrolase Pseudomonas aestuarii
WP 090500350.1 MBL fold metallo-hydrolase Pseudomonas borbori
WP 160346495.1 MBL fold metallo-hydrolase Pseudomonas xionganensis
WP 339904335.1 MBL fold metallo-hydrolase Pseudomonas guineae
MBX9764735.1 MBL fold metallo-hydrolase Pseudomonadaceae bacterium
MBL fold metallo-hydrolase Gammaproteobacteria bacterium HGW-Gammaproteobacteria-13'
WP 269380509.1 MBL fold metallo-hydrolase Pseudomonas anguilliseptica
MBL fold metallo-hydrolase Pseudomonadales bacterium RIFCSPHIGHO2 02 FULL 60 43'
WP 282875342.1 MBL fold metallo-hydrolase Pseudomonas peli

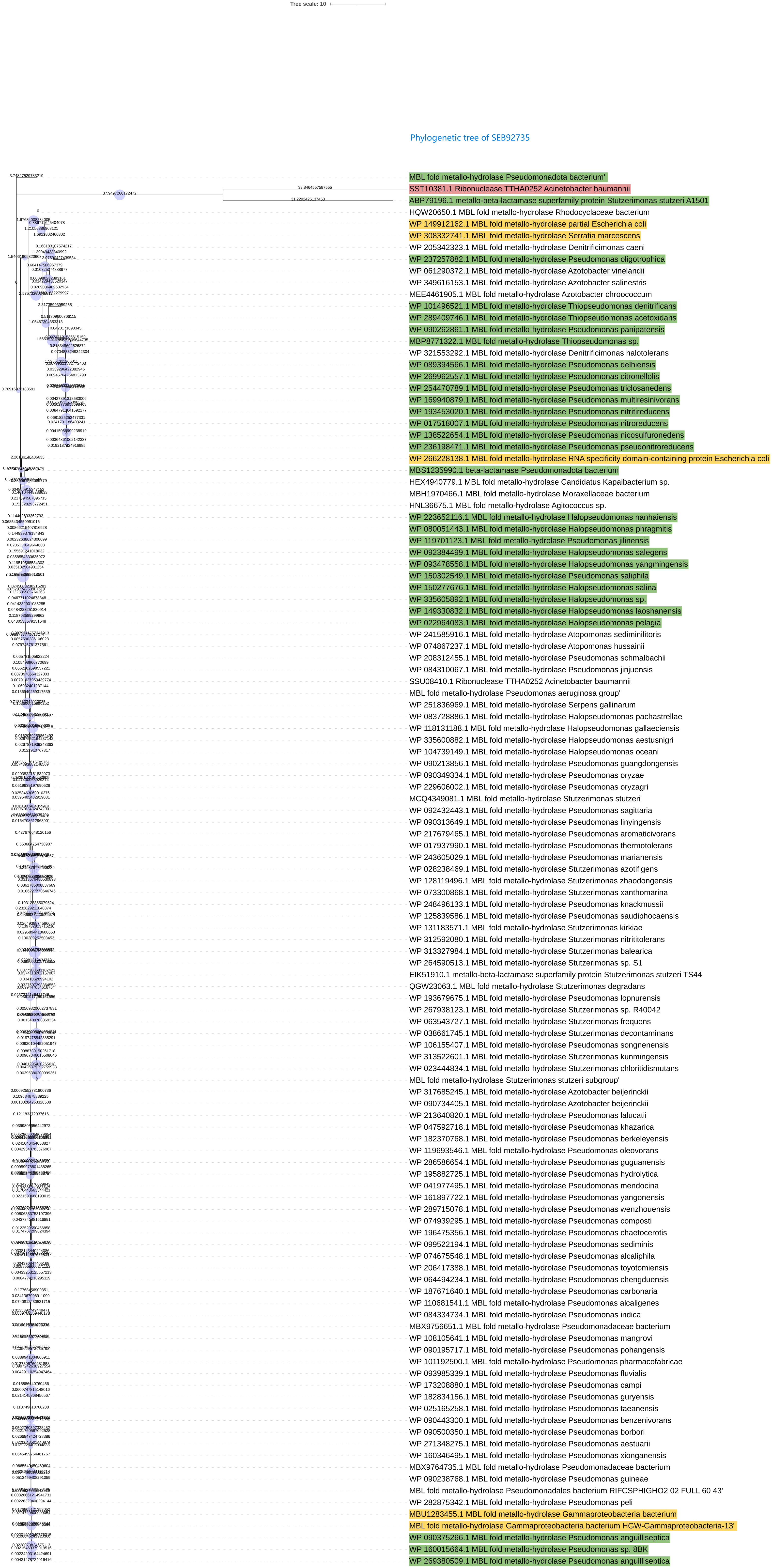
Tree scale: 10
Phylogenetic tree of SEB92735
MBL fold metallo-hydrolase Pseudomonadota bacterium'
SST10381.1 Ribonuclease TTHA0252 Acinetobacter baumannii
ABP79196.1 metallo-beta-lactamase superfamily protein Stutzerimonas stutzeri A1501
HQW20650.1 MBL fold metallo-hydrolase Rhodocyclaceae bacterium
WP 149912162.1 MBL fold metallo-hydrolase partial Escherichia coli
WP 308332741.1 MBL fold metallo-hydrolase Serratia marcescens
WP 205342323.1 MBL fold metallo-hydrolase Denitrificimonas caeni
WP 237257882.1 MBL fold metallo-hydrolase Pseudomonas oligotrophica
WP 061290372.1 MBL fold metallo-hydrolase Azotobacter vinelandii
WP 349616153.1 MBL fold metallo-hydrolase Azotobacter salinestris
MEE4461905.1 MBL fold metallo-hydrolase Azotobacter chroococcum
WP 101496521.1 MBL fold metallo-hydrolase Thiopseudomonas denitrificans
WP 289409746.1 MBL fold metallo-hydrolase Thiopseudomonas acetoxidans
WP 090262861.1 MBL fold metallo-hydrolase Pseudomonas panipatensis
MBP8771322.1 MBL fold metallo-hydrolase Thiopseudomonas sp.
WP 321553292.1 MBL fold metallo-hydrolase Denitrificimonas halotolerans
WP 089394566.1 MBL fold metallo-hydrolase Pseudomonas delhiensis
WP 269962557.1 MBL fold metallo-hydrolase Pseudomonas citronellolis
WP 254470789.1 MBL fold metallo-hydrolase Pseudomonas triclosanedens
WP 169940879.1 MBL fold metallo-hydrolase Pseudomonas multiresinivorans
WP 193453020.1 MBL fold metallo-hydrolase Pseudomonas nitritireducens
WP 017518007.1 MBL fold metallo-hydrolase Pseudomonas nitroreducens
WP 138522654.1 MBL fold metallo-hydrolase Pseudomonas nicosulfuronedens
WP 236198471.1 MBL fold metallo-hydrolase Pseudomonas pseudonitroreducens
WP 266228138.1 MBL fold metallo-hydrolase RNA specificity domain-containing protein Escherichia coli
MBS1235990.1 beta-lactamase Pseudomonadota bacterium
HEX4940779.1 MBL fold metallo-hydrolase Candidatus Kapaibacterium sp.
MBH1970466.1 MBL fold metallo-hydrolase Moraxellaceae bacterium
HNL36675.1 MBL fold metallo-hydrolase Agitococcus sp.
WP 223652116.1 MBL fold metallo-hydrolase Halopseudomonas nanhaiensis
WP 080051443.1 MBL fold metallo-hydrolase Halopseudomonas phragmitis
WP 119701123.1 MBL fold metallo-hydrolase Pseudomonas jilinensis
WP 092384499.1 MBL fold metallo-hydrolase Halopseudomonas salegens
WP 093478558.1 MBL fold metallo-hydrolase Halopseudomonas yangmingensis
WP 150302549.1 MBL fold metallo-hydrolase Pseudomonas saliphila
WP 150277676.1 MBL fold metallo-hydrolase Halopseudomonas salina
WP 335605892.1 MBL fold metallo-hydrolase Halopseudomonas sp.
WP 149330832.1 MBL fold metallo-hydrolase Halopseudomonas laoshanensis
WP 022964083.1 MBL fold metallo-hydrolase Halopseudomonas pelagia
WP 241585916.1 MBL fold metallo-hydrolase Atopomonas sediminilitoris
WP 074867237.1 MBL fold metallo-hydrolase Atopomonas hussainii
WP 208312455.1 MBL fold metallo-hydrolase Pseudomonas schmalbachii
WP 084310067.1 MBL fold metallo-hydrolase Pseudomonas jinjuensis
SSU08410.1 Ribonuclease TTHA0252 Acinetobacter baumannii
MBL fold metallo-hydrolase Pseudomonas aeruginosa group'
WP 251836969.1 MBL fold metallo-hydrolase Serpens gallinarum
WP 083728886.1 MBL fold metallo-hydrolase Halopseudomonas pachastrellae
WP 118131188.1 MBL fold metallo-hydrolase Halopseudomonas gallaeciensis
WP 335600882.1 MBL fold metallo-hydrolase Halopseudomonas aestusnigri
WP 104739149.1 MBL fold metallo-hydrolase Halopseudomonas oceani
WP 090213856.1 MBL fold metallo-hydrolase Pseudomonas guangdongensis
WP 090349334.1 MBL fold metallo-hydrolase Pseudomonas oryzae
WP 229606002.1 MBL fold metallo-hydrolase Pseudomonas oryzagri
MCQ4349081.1 MBL fold metallo-hydrolase Stutzerimonas stutzeri
WP 092432443.1 MBL fold metallo-hydrolase Pseudomonas sagittaria
WP 090313649.1 MBL fold metallo-hydrolase Pseudomonas linyingensis
WP 217679465.1 MBL fold metallo-hydrolase Pseudomonas aromaticivorans
WP 017937990.1 MBL fold metallo-hydrolase Pseudomonas thermotolerans
WP 243605029.1 MBL fold metallo-hydrolase Pseudomonas marianensis
WP 028238469.1 MBL fold metallo-hydrolase Stutzerimonas azotifigens
WP 128119496.1 MBL fold metallo-hydrolase Stutzerimonas zhaodongensis
WP 073300868.1 MBL fold metallo-hydrolase Stutzerimonas xanthomarina
WP 248496133.1 MBL fold metallo-hydrolase Pseudomonas knackmussii
WP 125839586.1 MBL fold metallo-hydrolase Pseudomonas saudiphocaensis
WP 131183571.1 MBL fold metallo-hydrolase Stutzerimonas kirkiae
WP 312592080.1 MBL fold metallo-hydrolase Stutzerimonas nitrititolerans
WP 313327984.1 MBL fold metallo-hydrolase Stutzerimonas balearica
WP 264590513.1 MBL fold metallo-hydrolase Stutzerimonas sp. S1
EIK51910.1 metallo-beta-lactamase superfamily protein Stutzerimonas stutzeri TS44
QGW23063.1 MBL fold metallo-hydrolase Stutzerimonas degradans
WP 193679675.1 MBL fold metallo-hydrolase Pseudomonas lopnurensis
WP 267938123.1 MBL fold metallo-hydrolase Stutzerimonas sp. R40042
WP 063543727.1 MBL fold metallo-hydrolase Stutzerimonas frequens
WP 038661745.1 MBL fold metallo-hydrolase Stutzerimonas decontaminans
WP 106155407.1 MBL fold metallo-hydrolase Pseudomonas songnenensis
WP 313522601.1 MBL fold metallo-hydrolase Stutzerimonas kunmingensis
WP 023444834.1 MBL fold metallo-hydrolase Stutzerimonas chloritidismutans
MBL fold metallo-hydrolase Stutzerimonas stutzeri subgroup'
WP 317685245.1 MBL fold metallo-hydrolase Azotobacter beijerinckii
WP 090734405.1 MBL fold metallo-hydrolase Azotobacter beijerinckii
WP 213640820.1 MBL fold metallo-hydrolase Pseudomonas lalucatii
WP 047592718.1 MBL fold metallo-hydrolase Pseudomonas khazarica
WP 182370768.1 MBL fold metallo-hydrolase Pseudomonas berkeleyensis
WP 119693546.1 MBL fold metallo-hydrolase Pseudomonas oleovorans
WP 286586654.1 MBL fold metallo-hydrolase Pseudomonas guguanensis
WP 195882725.1 MBL fold metallo-hydrolase Pseudomonas hydrolytica
WP 041977495.1 MBL fold metallo-hydrolase Pseudomonas mendocina
WP 161897722.1 MBL fold metallo-hydrolase Pseudomonas yangonensis
WP 289715078.1 MBL fold metallo-hydrolase Pseudomonas wenzhouensis
WP 074939295.1 MBL fold metallo-hydrolase Pseudomonas composti
WP 196475356.1 MBL fold metallo-hydrolase Pseudomonas chaetocerotis
WP 099522194.1 MBL fold metallo-hydrolase Pseudomonas sediminis
WP 074675548.1 MBL fold metallo-hydrolase Pseudomonas alcaliphila
WP 206417388.1 MBL fold metallo-hydrolase Pseudomonas toyotomiensis
WP 064494234.1 MBL fold metallo-hydrolase Pseudomonas chengduensis
WP 187671640.1 MBL fold metallo-hydrolase Pseudomonas carbonaria
WP 110681541.1 MBL fold metallo-hydrolase Pseudomonas alcaligenes
WP 084334734.1 MBL fold metallo-hydrolase Pseudomonas indica
MBX9756651.1 MBL fold metallo-hydrolase Pseudomonadaceae bacterium
WP 108105641.1 MBL fold metallo-hydrolase Pseudomonas mangrovi
WP 090195717.1 MBL fold metallo-hydrolase Pseudomonas pohangensis
WP 101192500.1 MBL fold metallo-hydrolase Pseudomonas pharmacofabricae
WP 093985339.1 MBL fold metallo-hydrolase Pseudomonas fluvialis
WP 173208880.1 MBL fold metallo-hydrolase Pseudomonas campi
WP 182834156.1 MBL fold metallo-hydrolase Pseudomonas guryensis
WP 025165258.1 MBL fold metallo-hydrolase Pseudomonas taeanensis
WP 090443300.1 MBL fold metallo-hydrolase Pseudomonas benzenivorans
WP 090500350.1 MBL fold metallo-hydrolase Pseudomonas borbori
WP 271348275.1 MBL fold metallo-hydrolase Pseudomonas aestuarii
WP 160346495.1 MBL fold metallo-hydrolase Pseudomonas xionganensis
MBX9764735.1 MBL fold metallo-hydrolase Pseudomonadaceae bacterium
WP 090238768.1 MBL fold metallo-hydrolase Pseudomonas guineae
MBL fold metallo-hydrolase Pseudomonadales bacterium RIFCSPHIGHO2 02 FULL 60 43'
WP 282875342.1 MBL fold metallo-hydrolase Pseudomonas peli
MBU1283455.1 MBL fold metallo-hydrolase Gammaproteobacteria bacterium
MBL fold metallo-hydrolase Gammaproteobacteria bacterium HGW-Gammaproteobacteria-13'
WP 090375266.1 MBL fold metallo-hydrolase Pseudomonas anguilliseptica
WP 160015664.1 MBL fold metallo-hydrolase Pseudomonas sp. 8BK
WP 269380509.1 MBL fold metallo-hydrolase Pseudomonas anguilliseptica

Phylogenetic tree of Ysh1

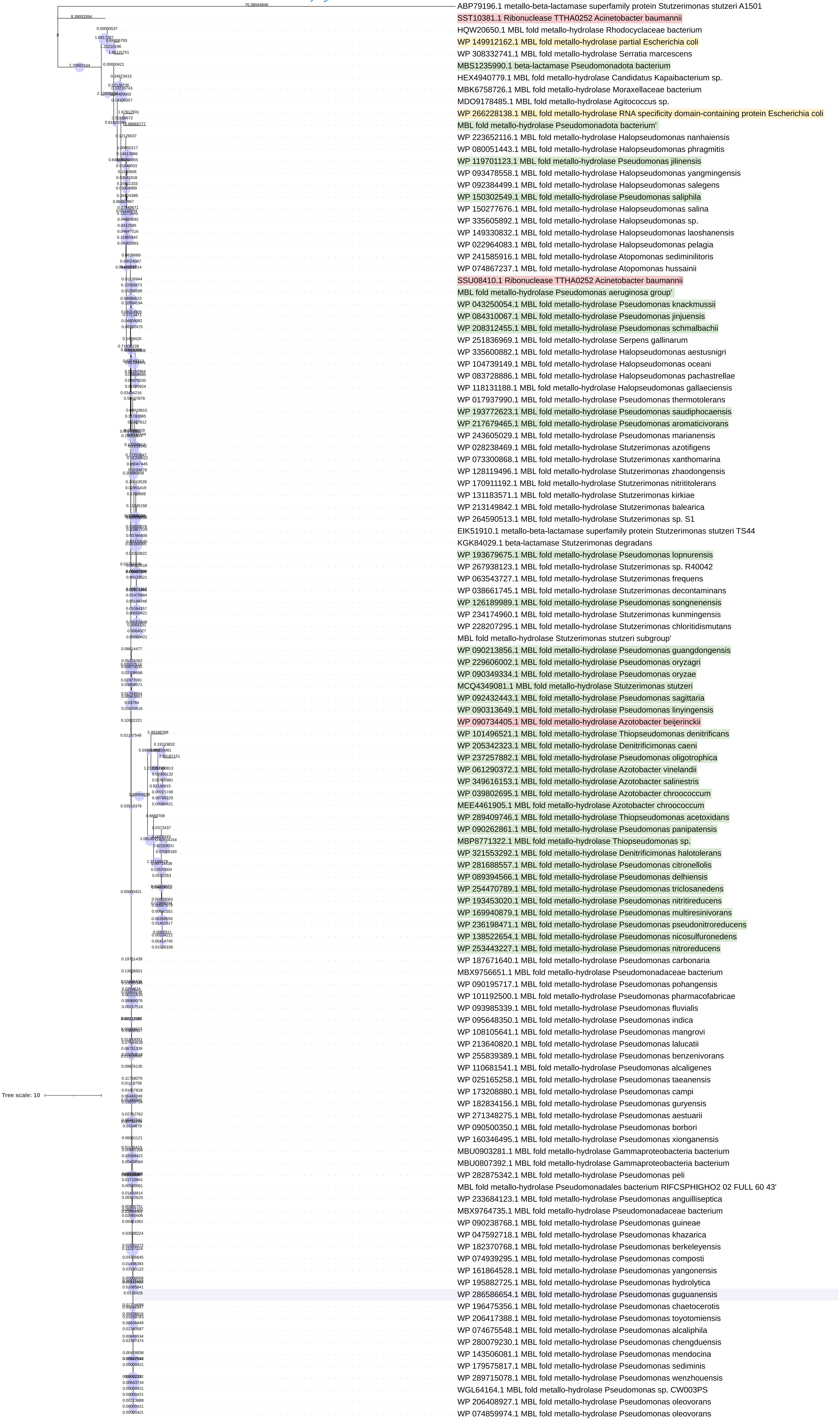
