## Supplementary figures and images for "Phylogenetic Analysis of Beta-Lactamases Reveals Distinct Evolutionary Patterns of Chromosomal and Plasmid-Encoded BLs and the Mosaic Role of VIM Linking NDM and IMP"

### Supplementary File 5

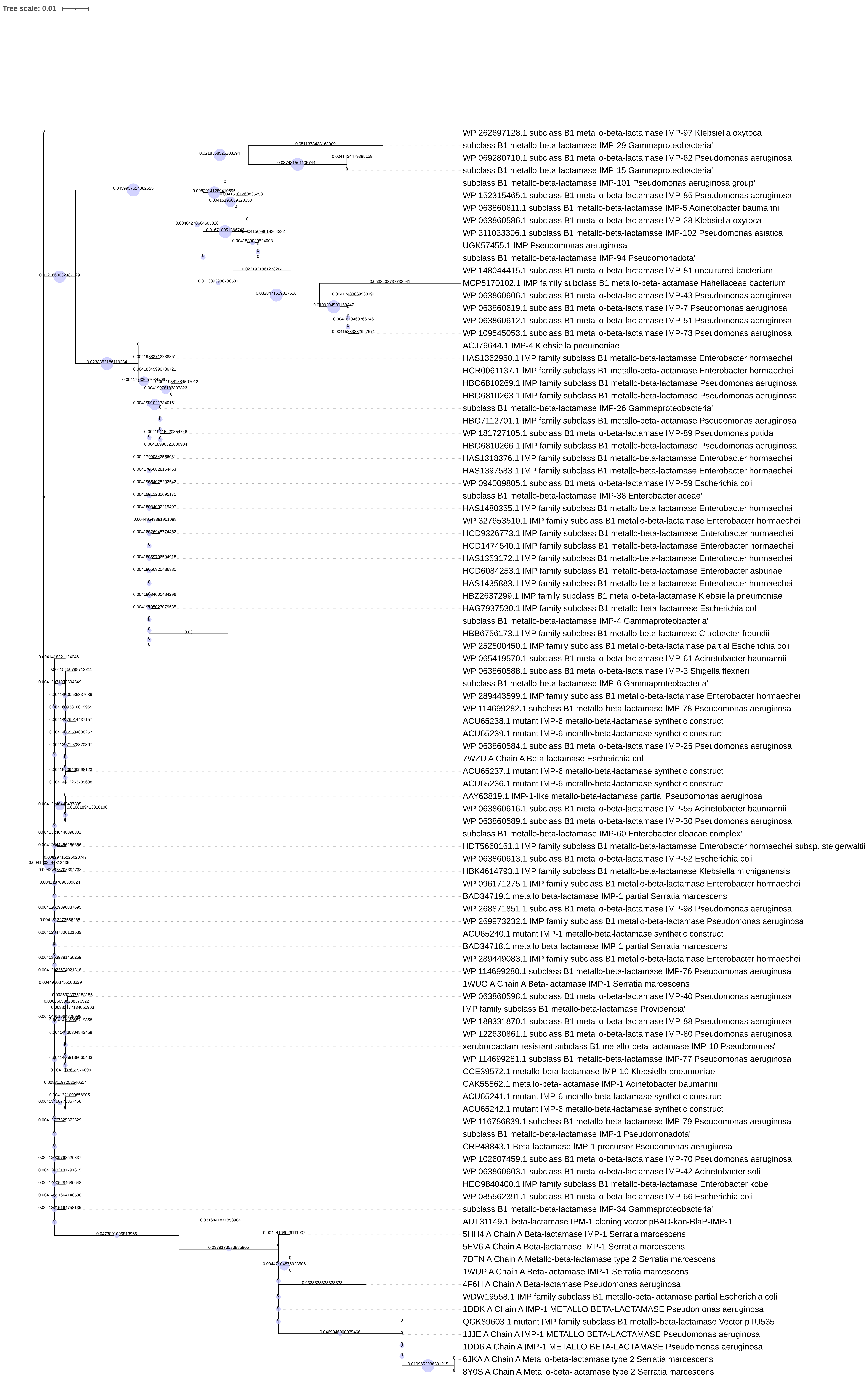

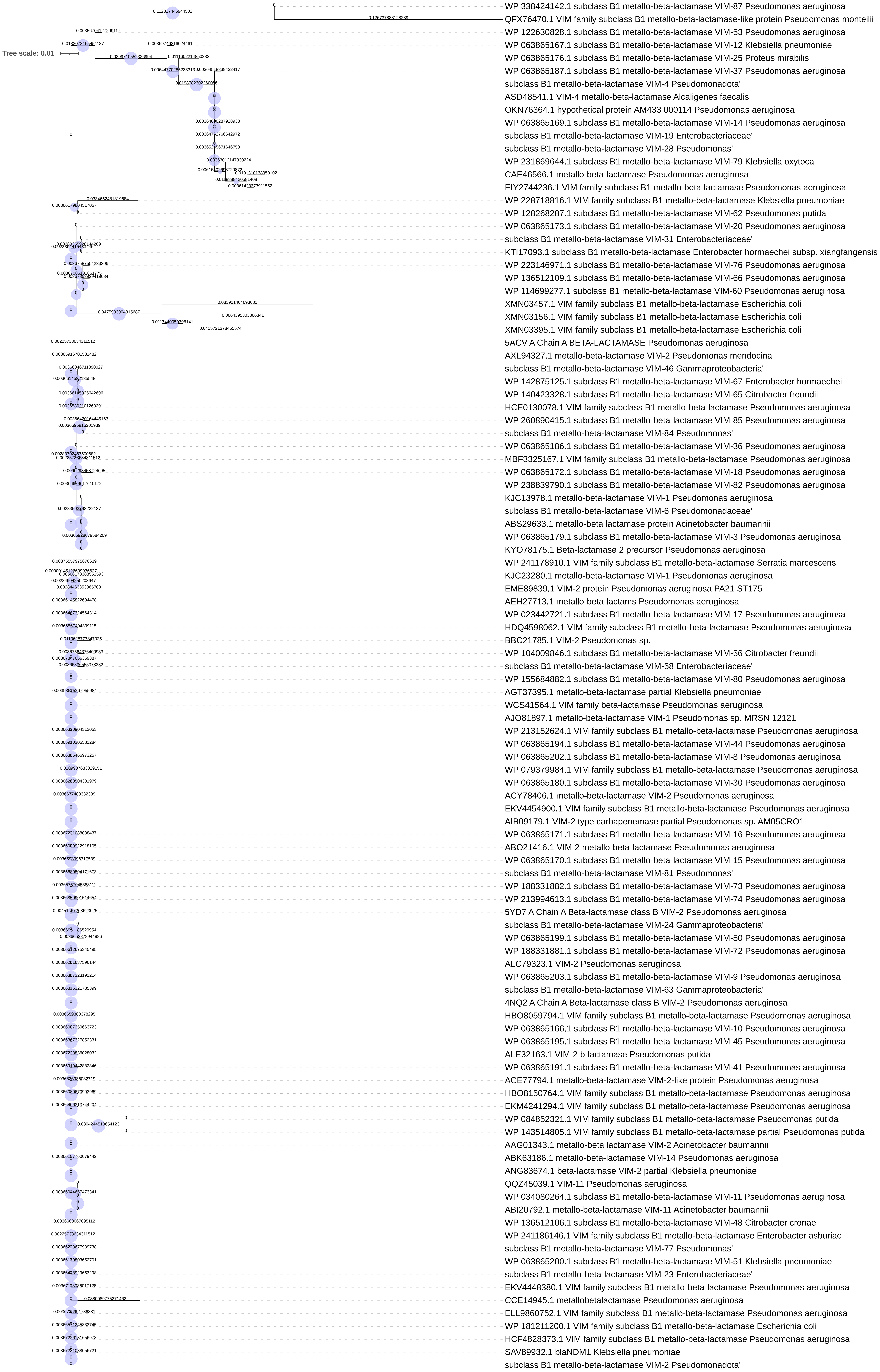

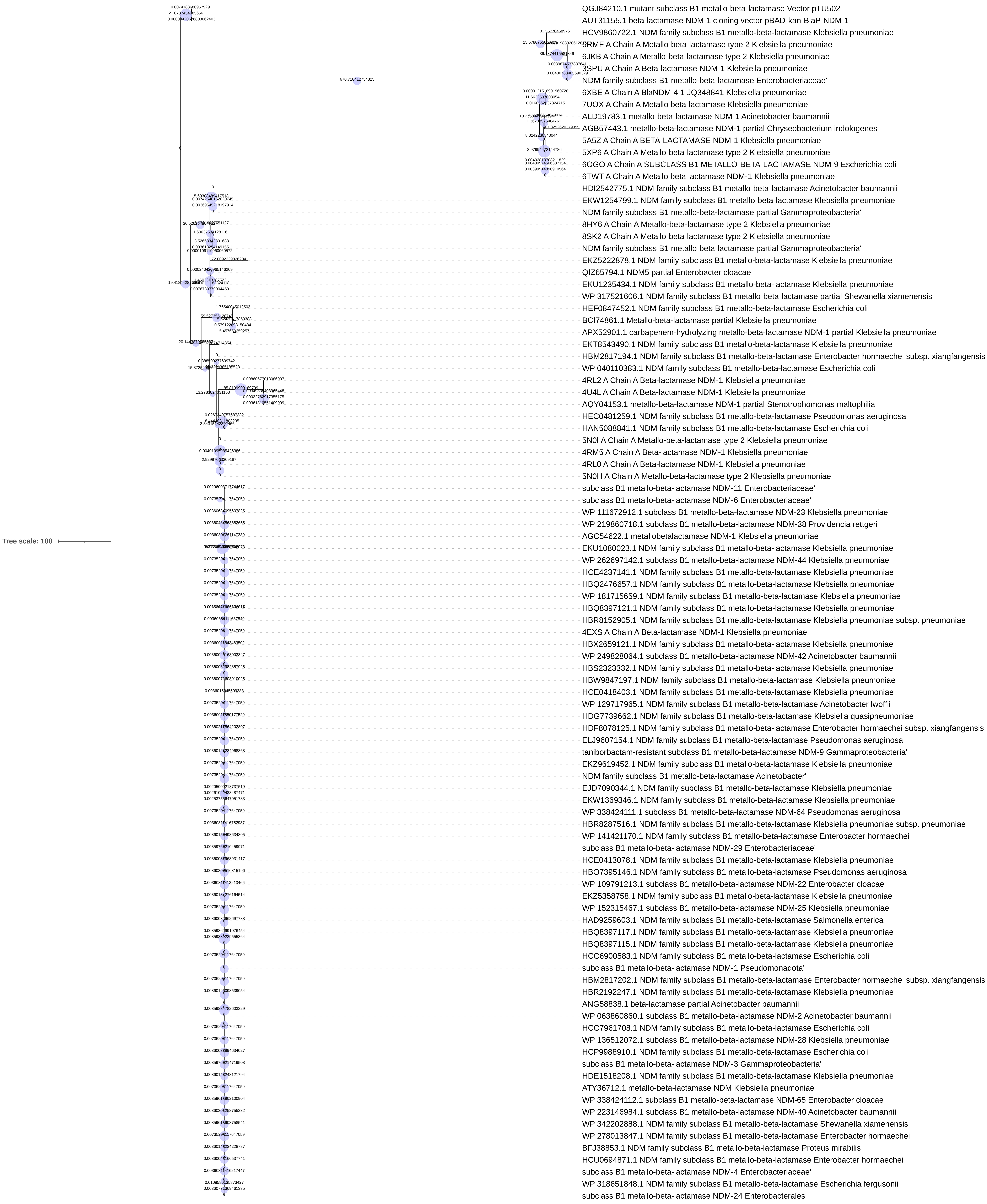

### Supplementary File 6

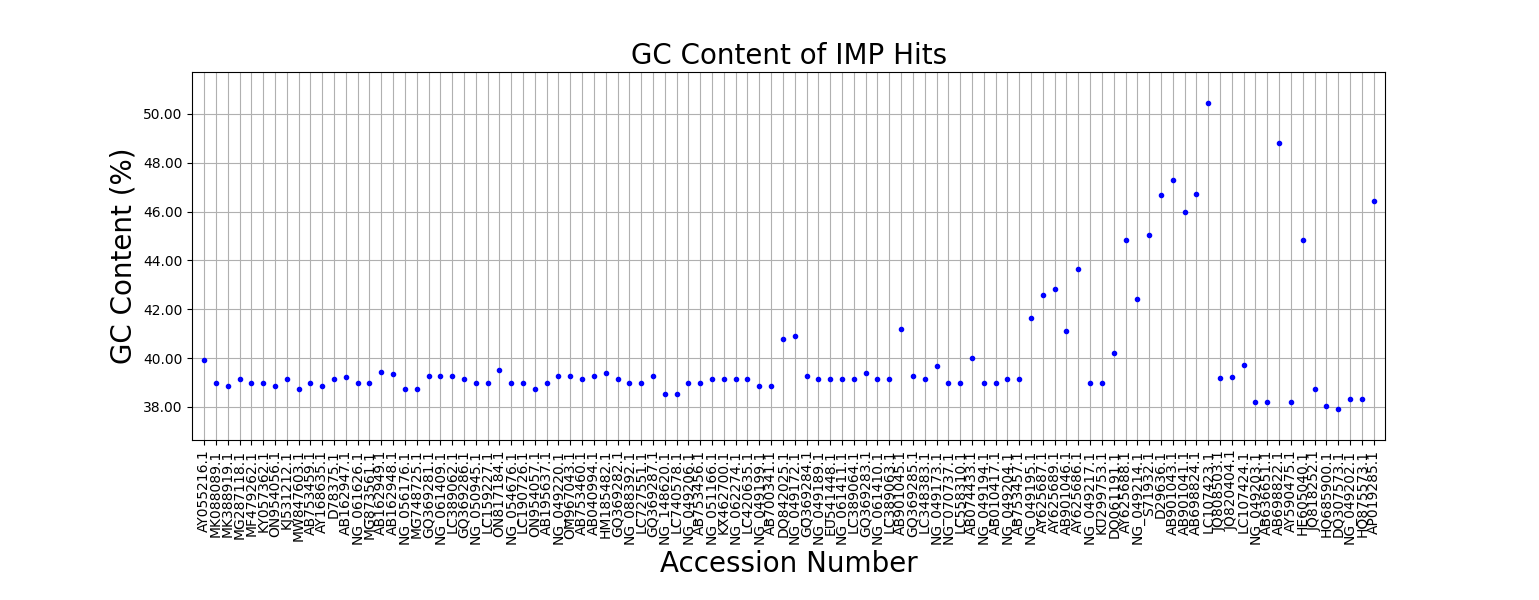

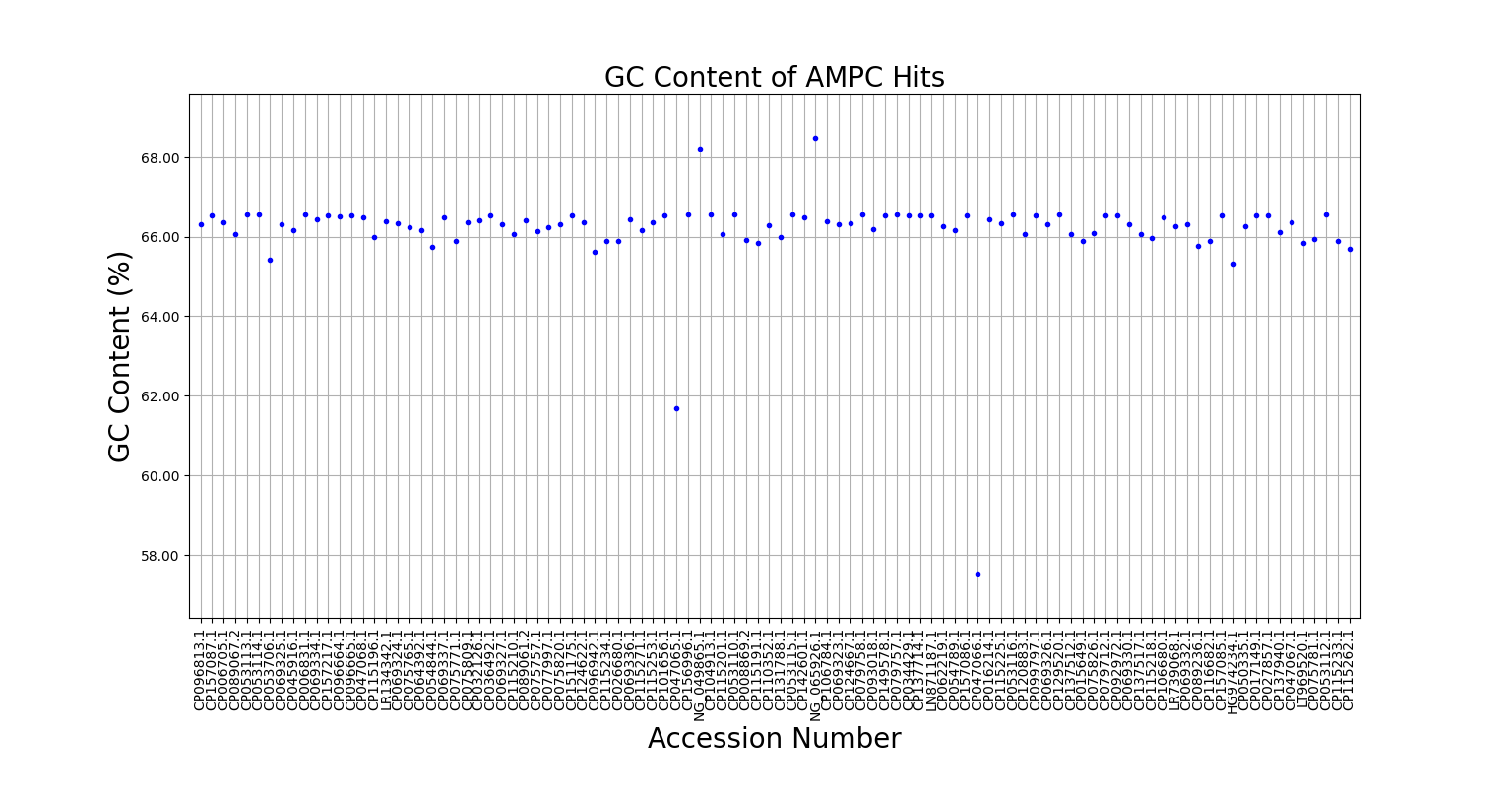

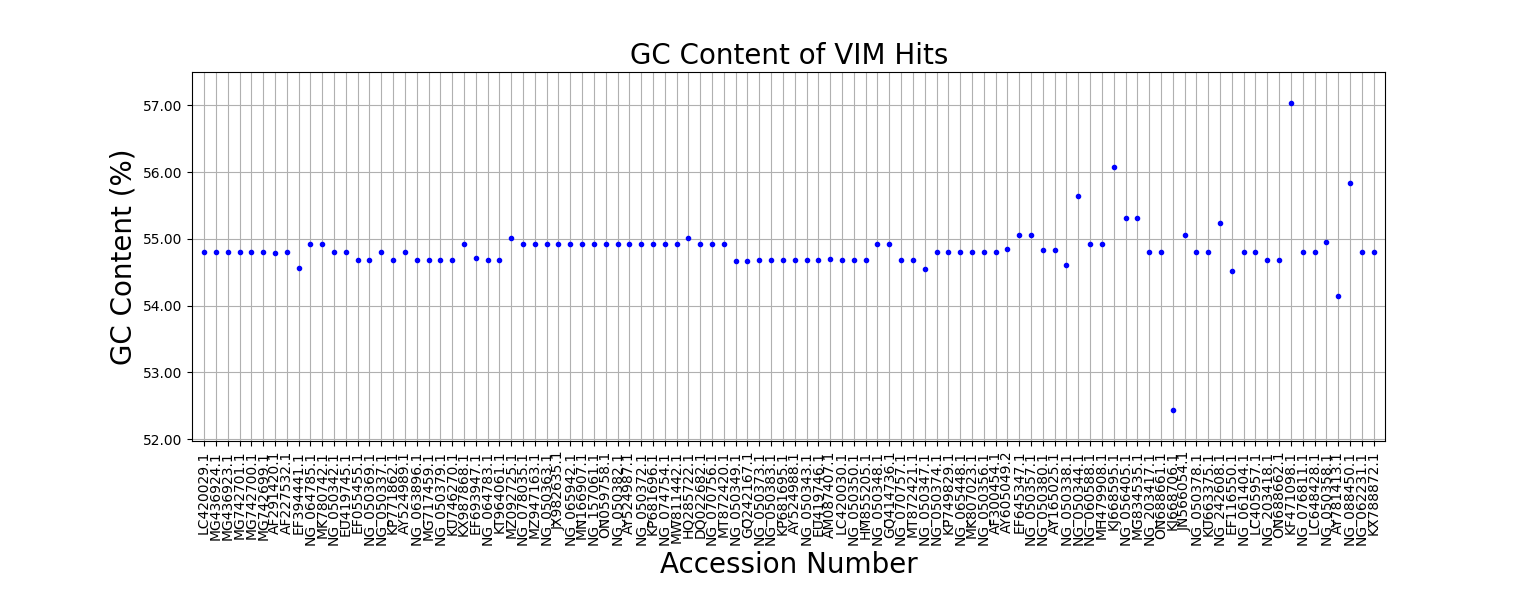

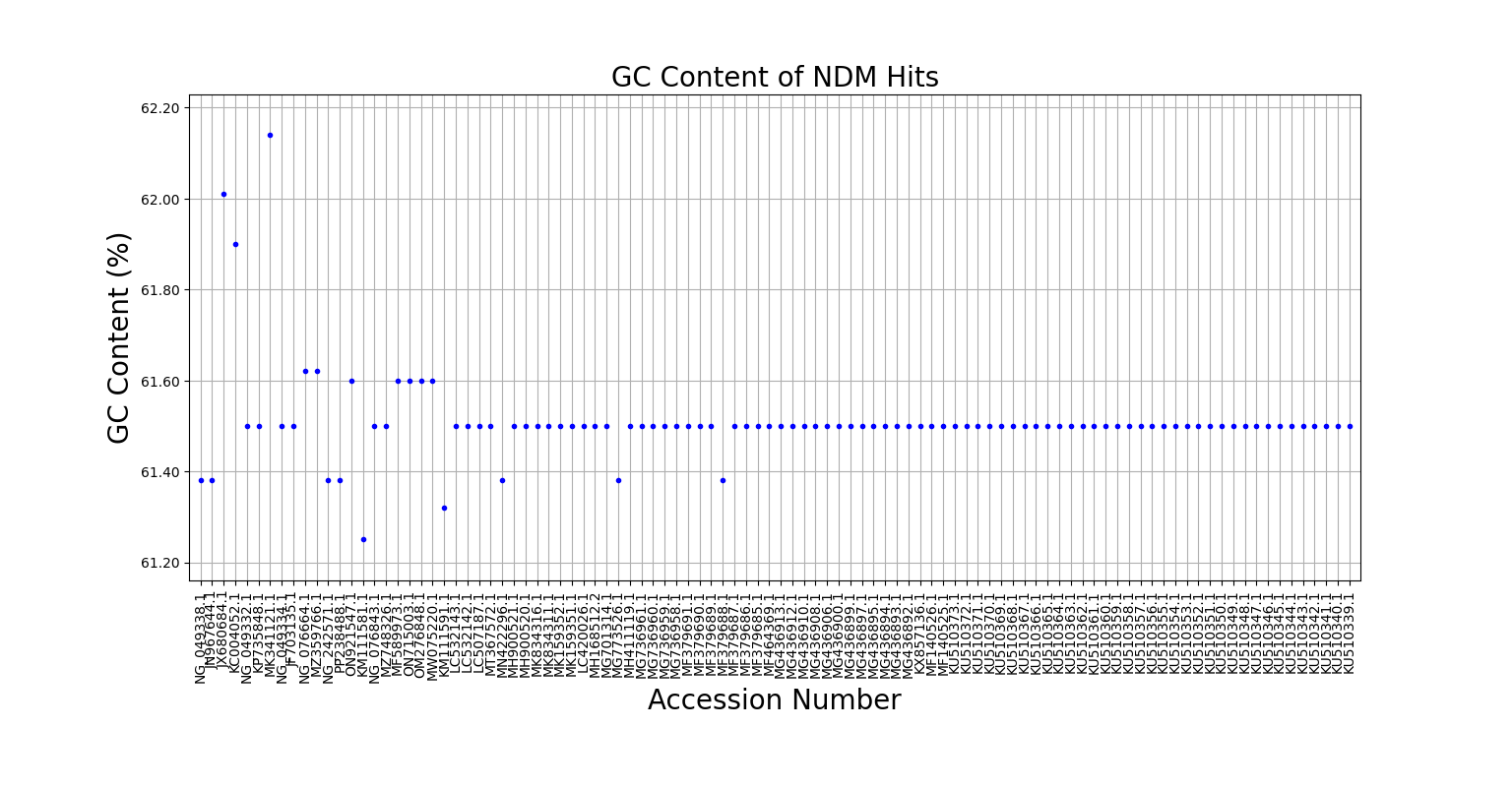
