## Supplementary File 7 for "Phylogenetic Analysis of Beta-Lactamases Reveals Distinct Evolutionary Patterns of Chromosomal and Plasmid-Encoded BLs and the Mosaic Role of VIM Linking NDM and IMP"

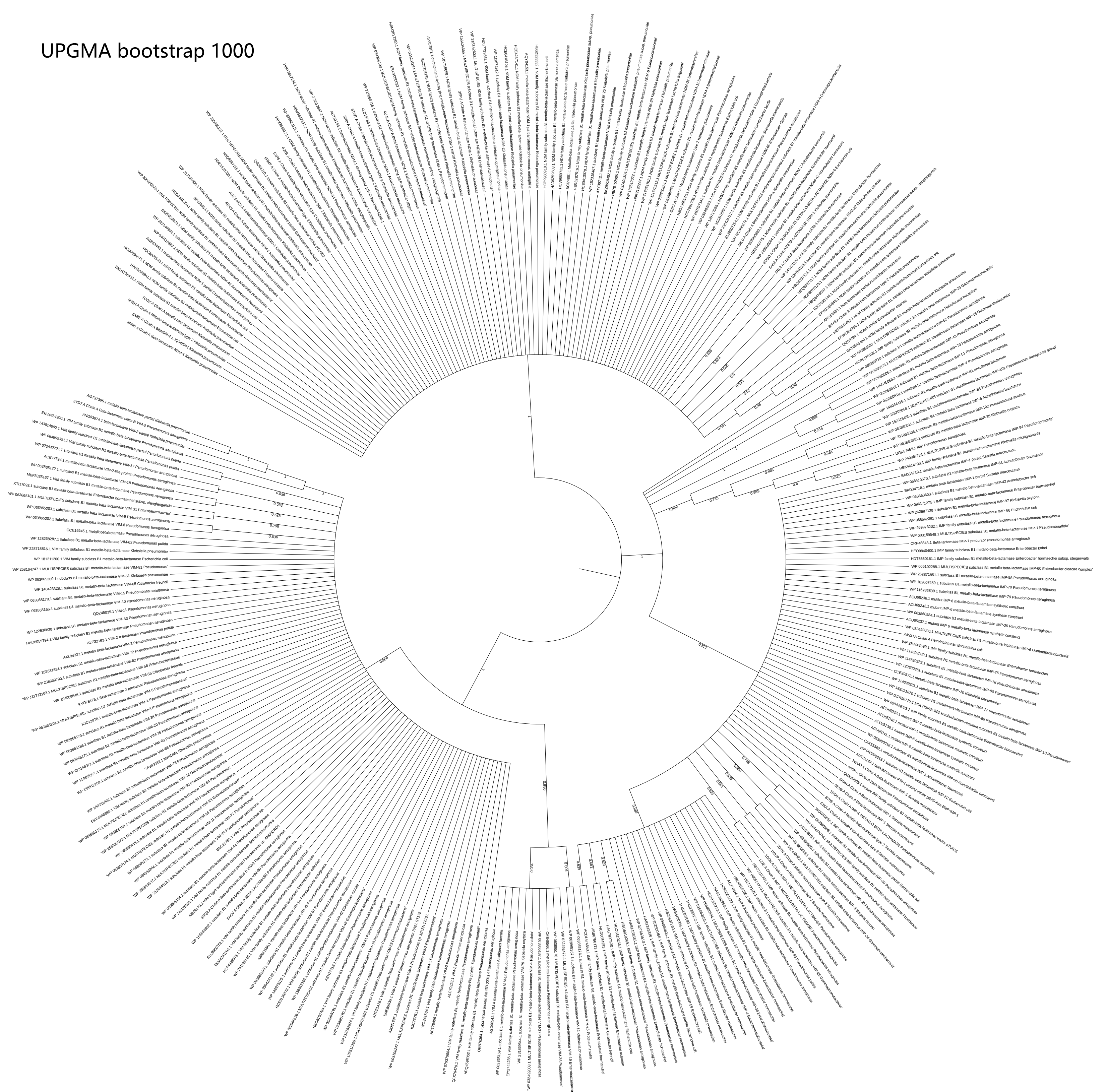


Evolutionary relationship between IMP, VIM and NDM as derived from UPGMA analysis

Neighbor-joining bootstrap 1000

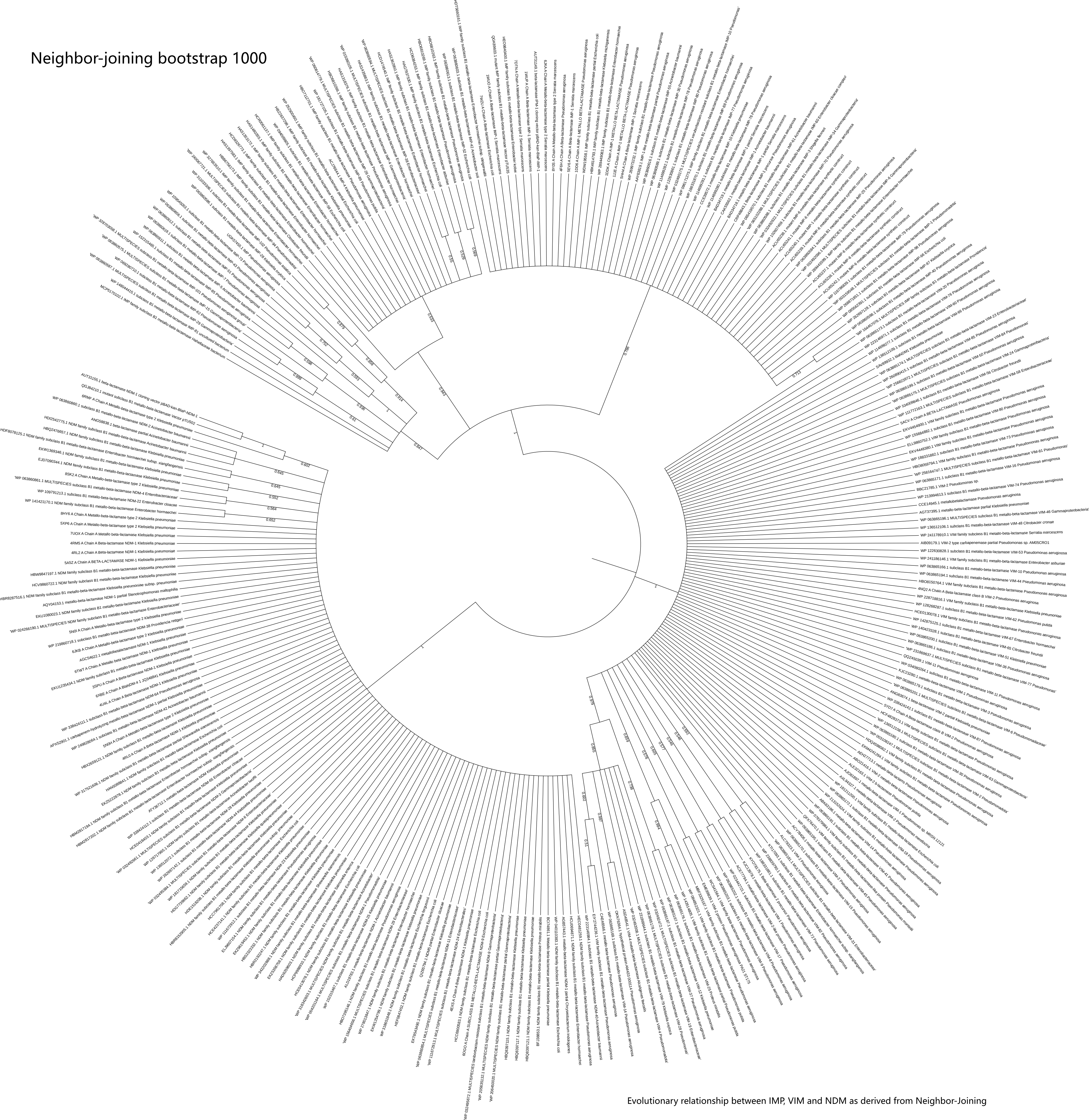

Evolutionary relationship between IMP, VIM and NDM as derived from Neighbor-Joining

Minimum Evolution
bootstrap 1000

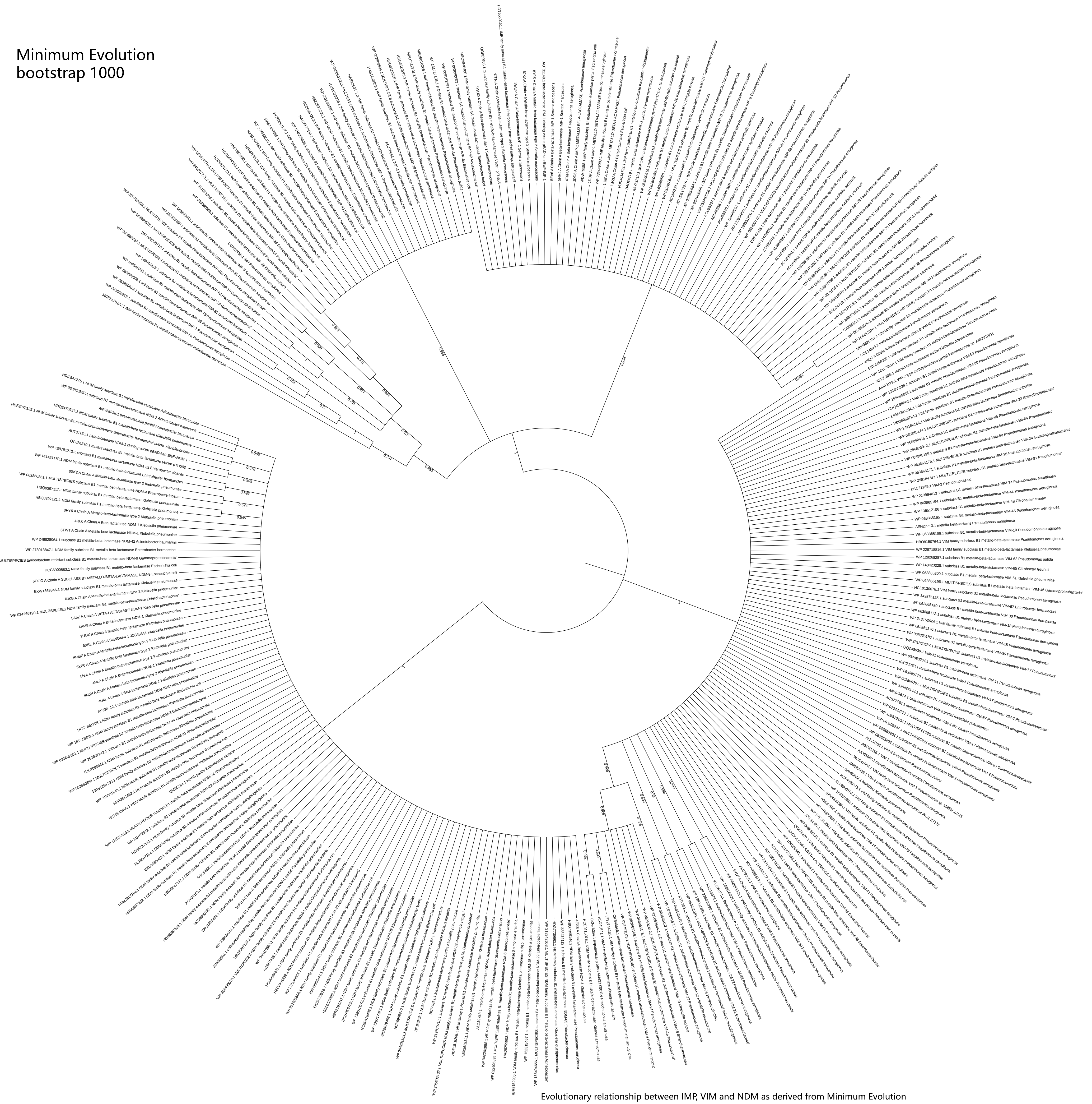

Evolutionary relationship between IMP, VIM and NDM as derived from Minimum Evolution

Maximum Parsimony
bootstrap 1000

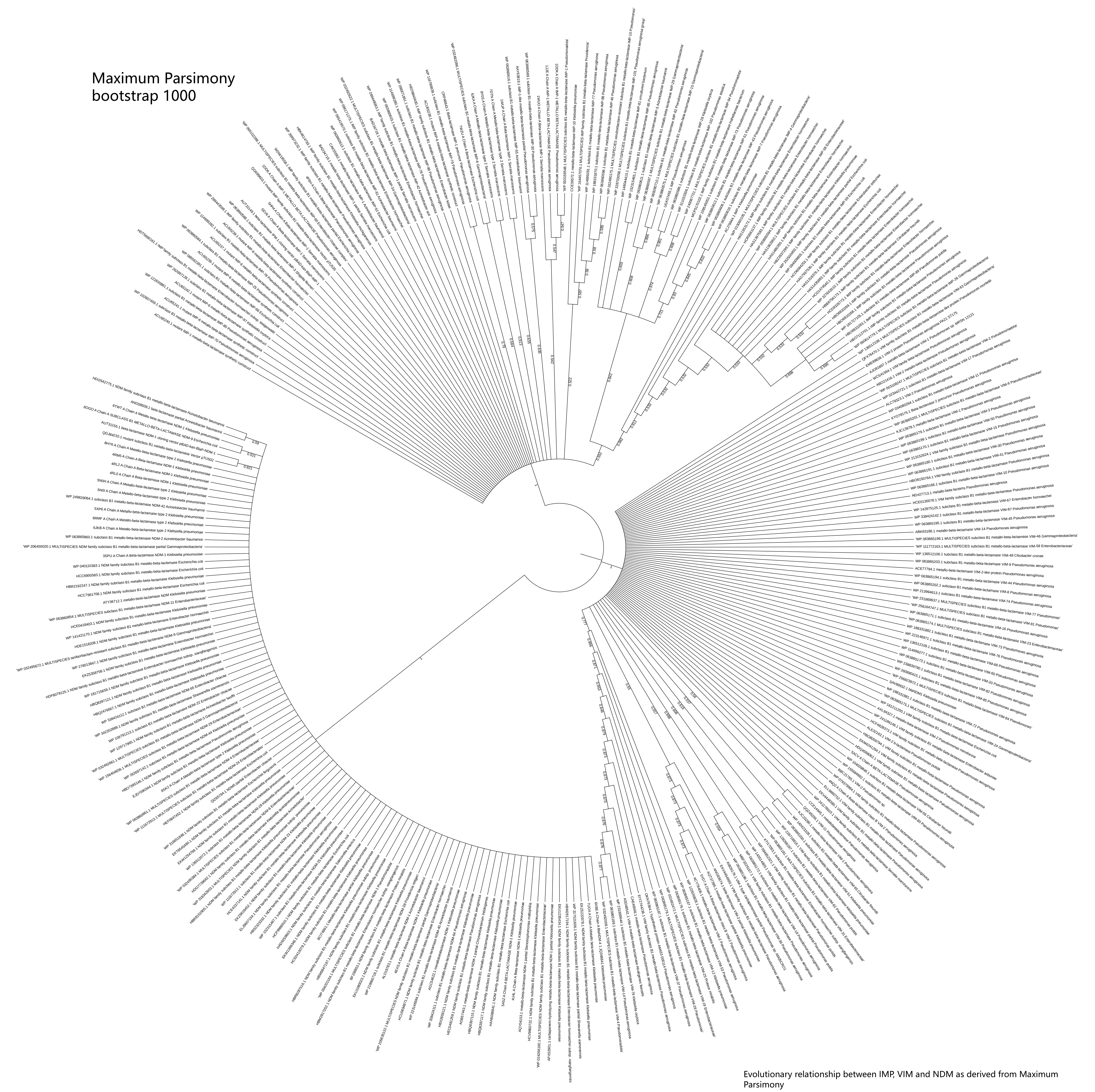

Evolutionary relationship between IMP, VIM and NDM as derived from Maximum Parsimony

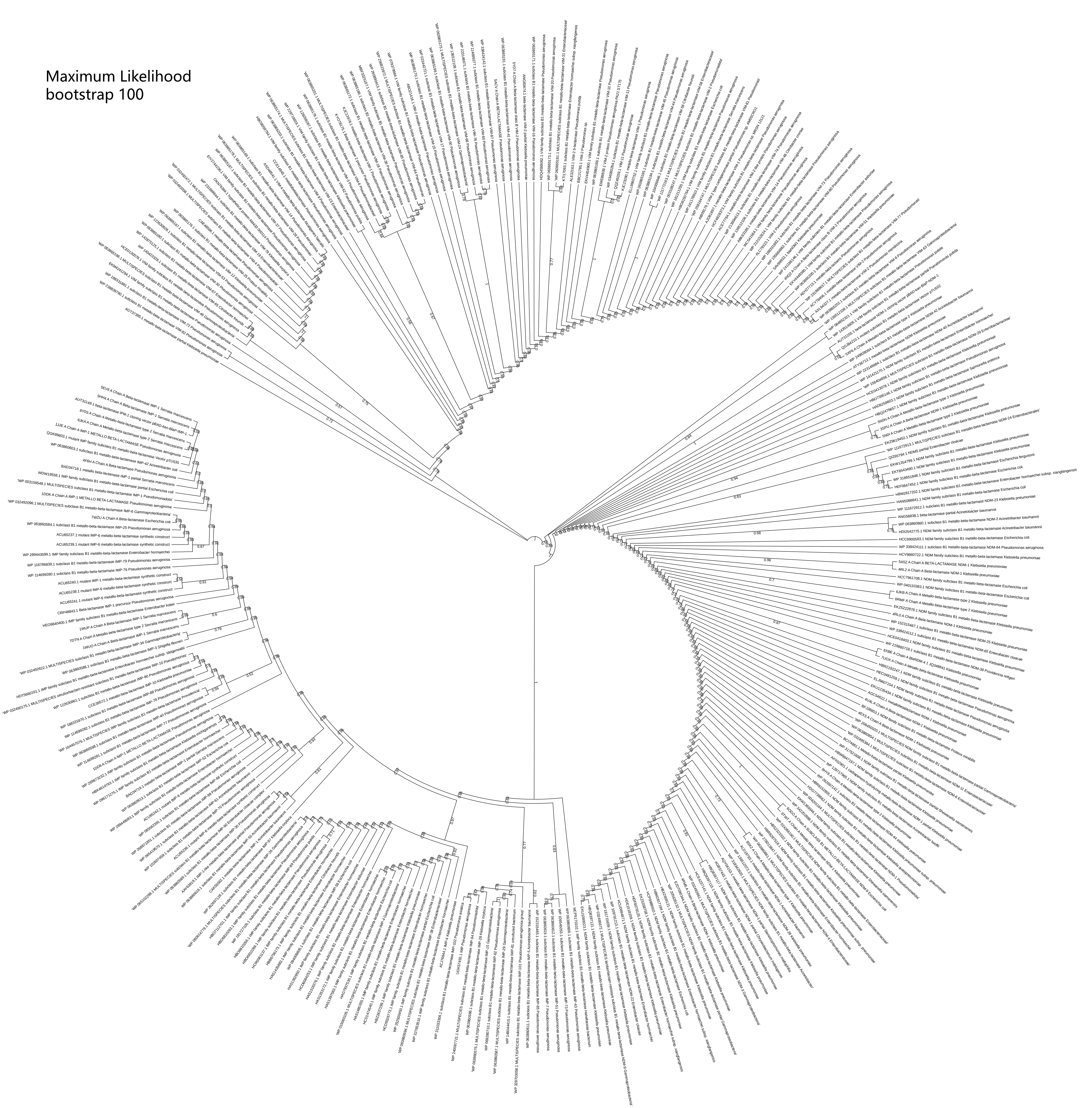


Evolutionary relationship between IMP, VIM and NDM as derived from Maximum Likelihood
