## Supplementary File 8 for "Phylogenetic Analysis of Beta-Lactamases Reveals Distinct Evolutionary Patterns of Chromosomal and Plasmid-Encoded BLs and the Mosaic Role of VIM Linking NDM and IMP"

**Table S1. Motifs conserved across all 3 MBLs- IMP, VIM and NDM**

ADADLDAWPASIAKVQARYPDAKIVVPGHGK

AEDTJGLPVRAAVVT

AEDTLGLPVRAAVVT

AGLGAD

AKDLGNLADADVAAWPASLER

ASNGLI

ASNKSIQPTAEASAD

AYIIDTPWTEKDTEKLVDWIEAQGLTLKASISTHSHEDRTGGIGYLNSIG

AYIIDTPWTEKDTEKLVDWIEAQGLTLKASISTHSHEDRTGGIGYLNSKG

AYJIDTPWTEKDTEKLVDWIEAQGLTLKASISTHSHZDRTGGIGYLNSKG

DALHAAGI

DAQTLGPLEVFFPGAGHAPDNJVVWHPASGVLFGGCFVKDA

DAQTLGPLEVFFPGAGHAPDNLVVWHPASGVLFGGCFVKDA

DQQEAIIFDTPATDQASEELI

EVFYPGAGHTKDNVVVWLPEEKILFGGCFVKS

EVFYPGAGHTMDNIVVWLPQQKILFGGCLVKSLQAKDLGNTADADLNSWP

EVFYPGPGHTMDNIVVWLPZQKILFGGCLVKSLQAKDLGNTADADLNEWP

EWIKTKLKKPVKKAI

FDTPWTBEPTEQLLAWVKDNLKAPVKAFVPTHWHDDCLGGL

FDTPWTDEPTEQLLAWVKDNLKAPVKAFVPTHWHDDCLGGL

FDTPWTDEQTEQLLAWVKDNLKAPVKAFVPTHWHDDCLGGL

FGNTKV

FPSNGLIVETGKGLVLIDTAWGEEQTEEL

FPSNGLIVETGKGLVLIDTAWGEEQTEZL

GCFGKD

GDABVSAWPNSVEKVKKK

GDANVSAWPNSVEKVKKK

GKDAYIIDTPWTEADTEKLVDWIEQQGLTLKASVSTHSHEDRTGGIGYLN

GKDAYJIDTPWTEADTEKLVDWIEQQGLTLKASVSTHSHEDRTGGIGYLN

GPGHTKDNVVVWLPKEKILFGGCLVKSLGAGNLGL

GVLFPSHGLVVSTKDGAVLVDTGWGNEPTEQLLA

GVLFPSHGLVVSTKGGAVLVDTGWGNEPTEQLLA

GYKIKGSISTHFHEDSTGGIEWLNSHSIPTYASELTNELLK

GYKJKGSISTHFHEDSTGGJEWLNSHSIPTYASELTNELLK

HFHDDRTGGVPVLRARGIPVYALPDTARL

HRPVRAAVVTHFHDDRTGGIPALVARGIPVHALED

HRPVRAAVVTHFHDDRTGGIPALVARGIPVYALED

IEKJSENVYLHTSFKZTNGWG

IEKLSENVYLHTSFKETNGWG

IEVFYPGAGHTKDNIVVWLPKQKILFGGCLVKS

IEVFYPGAGHTKDNLVVWLPKQKILFGGCLVKSLESKSLGY

IEVYYPGAGHTKDNLVVWLPKQKJLFGGCLVKSLESKSLGY

INLPVALA

IPTYASELTNALLAQNGKPLA

IPTYASELTNELLKKKGKPQA

IPTYASELTNELLKKNGKPQA

KGLGNLGDANIEAWPKSAKKVKSK

KJADGVYLHTSYKEVEGFGLVSSNGLVVV

KLADGVYLHTSYKEVEGFGLVSSNGLVVV

KSHIT

LGDADTEHYAASARAFGAAFPKASMIVMSHSAPDSRAAITHTARMADKLR

LVDTGWGPRQTEALLDWARDTL

LVPANGLVVLDNKEAYJIDTPWTAKDTEKLVTWIEER

LVPANGLVVLDNKEAYLIDTPWTAKDTEKLVTWIVER

MERGEDMEMEOUTPUT

MHPPLFKHPLFKSLGLALLALLLGAAGAR

MHPPLRPHPGWKRLGPALLALLLGVFGAR

MKKLJLLLLLLLILL

MKKLLLLLLLLLILL

MNT

MSIQHFRVALI

MSIQHFRVALIPFFAAFCLP

MSIQHFRVALIPFFAAFCLPV

MSIQHFRVALIPFFAAFCLPVFAGEQGH

MSIQHFRVALIPFFAAFCLPVFALSQGH

NLGDADT

NLGDPDC

PAYTLLAQEQELQVTKLAPGVWVHTSYSTYGGVLVPSHGLVVSTKGGAVL

PAYTLQAQEQEJQVTKJAPGVWVHTSYSTYNGVLVPSHGLVVSTKEGAVL

PCNGLVVIDNKEAYJIDTPWNAKDTE

PCNGLVVIDNKEAYLIDTPWNAKDTE

PEARSON

PJADGVYLHTSYKZVEGFGLVDSNGLVVV

PLADGVYLHTSYKQVEGFGLVDSNGLVVV

RARYPEARVVVPGHGAPGGPELLDHTEAL

RGYKIKASISTHFHEDSTGGLEYLNSHSIPTYASELTNELL

RGYKJKASISTHFHEDSTGGJEYLNSHSIPTYASELTNELL

SIQHFRVALIPFFAAFCLPVF

SLGNTGDADLEAWPASIAKVQAR

VRDGGRVLVVDTAWTDDQTAQILNWIKQEINLPVALAVVTHAHQDKMGGM

VRFGPVELFFPGAGHSPDNLV

VTHAHDDRIGGIDVLKKRGIPVYSTPLTA

VVTHAHQDKMGGMDALHAAGIATYANALSNQLAPQEGMVAAQHSLTFAAN

WQHTSYLDMPGFGAVASNGLIVRDGGRVLVVDTAWTDDQTAQILNWIKQE

WVEPATAPNFGPLKVFYPGPGHTSDNITVGIDGTDIAFGGCLIKDSKAKS

WVETNLKQPVKAVVATHFHDDCLGGLGAF

WVETNLKQPVKAVVATHFHEDCLGGLQAF

YPANGLIVEDGDELLLVDTAWGARQTAALL

YPANGLIVEDGDGSL

YPANGLJVEDGDGSJ

**Additional details:**

**AmpC vs IMP**

| **Query_ID (AmpC)** | **Target_ID (IMP)** | Optimal_offset | p-value | E-value | q-value | Overlap | **Query_consensus (AmpC)** | **Target_consensus**  **(IMP)** | Orientation |
| --- | --- | --- | --- | --- | --- | --- | --- | --- | --- |
| CALCFS | MFNRTTRKSINWLLISLCFSACLTTTLLSMSAKANPKAEAEKEAEKPSLQ | 14 | 0.000999502 | 0.0499751 | 0.0499751 | 6 | AALCSS | MFNRTTRKSINWLLISLCFSACLTTTLLSMSAKANPKAEAEKEAEKPSLQ | + |
| FAACTC | DVTFATDTTFTVGKQ | 3 | 0.000266228 | 0.0133114 | 0.0133114 | 6 | FAAGTT | DVTFATDTTFTVGGQ | + |
| MLSGVRNLLISALLLGAGNCM | MKKJFLLLJFL | -4 | 0.00036013 | 0.0180065 | 0.0180065 | 11 | MLSGVRNLLISALLLGAGNCM | MKKLFLLLLFL | + |
| MQQRRAFALLTLGSLLLAPCT | MKLAIALFLAYILLLVPSTHASQEQFKSATNQLEPTKGQLS | -2 | 0.00010469 | 0.00523452 | 0.00523452 | 19 | MQQRRAFALLTLGSLLLAPCT | MKLAIALFLAYILLLVPSTHASQEQFKSATNQLEPTKGQLS | + |
| MRFTITALALLLIGQ | MRIAIRGCAALMIGAIANAQA | 0 | 0.000177765 | 0.00888827 | 0.00888827 | 15 | MRLTITALALLLIGQ | MRIAISGCAALMIGAIANAQA | + |
| MSSINPSK | MFSSIN | 1 | 0.000219925 | 0.0109963 | 0.0109963 | 5 | MSSINLSK | MFSSIN | + |
| MSSINPSK | NESKKPSKPSN | 0 | 0.000969706 | 0.0484853 | 0.0242427 | 8 | MSSINLSK | NESKKPSKPSN | + |
| RNJLAGALLLAAGTC | MKKLJLLLLJL | 1 | 9.59632e-06 | 0.000479816 | 0.000479816 | 10 | RNLLAGALLLAAGTC | MKSLILLLLIL | + |
| RNJLAGALLLAAGTC | MKKLLVFFJFLFCSI | 1 | 0.000392483 | 0.0196242 | 0.00981209 | 14 | RNLLAGALLLAAGTC | MKKLLVFFIFLFCSI | + |
| RNJLAGALLLAAGTC | MKYIJLLLLLI | 1 | 0.000671791 | 0.0335896 | 0.0111965 | 10 | RNLLAGALLLAAGTC | MKYIILLLLLI | + |
| EANMGYQGDAA | KGLGNLGDANIEAWPKSAKKVKSK | -1 | 0.000274357 | 0.0137179 | 0.0137179 | 10 | EANMGYQGDAA | KGLGNLGDANIEAWPKSAKKVKSK | + |
| MQQRRAFA | KTKQRAAEASA | 1 | 0.00098068 | 0.049034 | 0.049034 | 8 | MQQRRAFA | KTKQRAAEASA | + |
| MQQRRAFALLTLGSLLLAPCT | MKKLJLLLLLLLILL | -2 | 0.00049853 | 0.0249265 | 0.0181307 | 15 | MQQRRAFALLTLGSLLLAPCT | MKKLLLLLLLLLILL | + |
| MQQRRAFALLTLGSLLLAPCT | MPNRALWQRIAILLJMPILSLPLLAANLT | 5 | 0.000725229 | 0.0362615 | 0.0181307 | 21 | MQQRRAFALLTLGSLLLAPCT | MPNRALWQRIAILLLMPILSLPALAANLT | + |
| PCTYAS | IPTYASELTNALLAQNGKPLA | 0 | 0.000307092 | 0.0153546 | 0.0153546 | 6 | PCTYAS | IPTYASELTNALLAQNGKPLA | + |
| PCTYAS | GYKJKGSISTHFHEDSTGGJEWLNSHSIPTYASELTNELLK | 27 | 0.000909859 | 0.045493 | 0.0227465 | 6 | PCTYAS | GYKIKGSISTHFHEDSTGGIEWLNSHSIPTYASELTNELLK | + |
| QKDQAQ | MJSAPSFAHETEQQPAQSNTDAAQKRQQQ | 23 | 0.000516303 | 0.0258152 | 0.0258152 | 6 | QKDQAQ | MLSAPSFAHETEQQTAQSNTDAAQKRQQQ | + |
| RPLLAALALLLASQS | MKKLJLLLLLLLILL | 1 | 1.07428e-05 | 0.000537138 | 0.000537138 | 14 | RPLLAALALLLASQS | MKKLLLLLLLLLILL | + |
| RPLLAALALLLASQS | MKFLAPFLFLLPCVTVATEAS | 1 | 0.000166392 | 0.00831962 | 0.00415981 | 15 | RPLLAALALLLASQS | MKFLAPFLFLLPCVTVATEAS | + |
| RPLLAALALLLASQS | MHPPLRPHPGWKRLGPALLALLLGVFGAR | 12 | 0.00028153 | 0.0140765 | 0.00469217 | 15 | RPLLAALALLLASQS | MHPPLFPHPLFKSLGLALLALLLGAAGAR | + |
| RPLLAALALLLASQS | MKYJJLJJLLL | 1 | 0.000800625 | 0.0400313 | 0.0100078 | 10 | RPLLAALALLLASQS | MKYILLLLLLL | + |
| RSYSAFALLLGAGTC | MHPPLRPHPGWKRLGPALLALLLGVFGAR | 13 | 8.78345e-05 | 0.00439173 | 0.00439173 | 15 | RSYSAFALLLGAGTC | MHPPLFPHPLFKSLGLALLALLLGAAGAR | + |
| RSYSAFALLLGAGTC | MKYJJLJJLLL | 1 | 0.000186565 | 0.00932825 | 0.00466412 | 10 | RSYSAFALLLGAGTC | MKYILLLLLLL | + |
| RSYSAFALLLGAGTC | MKFLAPFLFLLPCVTVATEAS | 1 | 0.000384849 | 0.0192425 | 0.00530658 | 15 | RSYSAFALLLGAGTC | MKFLAPFLFLLPCVTVATEAS | + |
| RSYSAFALLLGAGTC | MKKLJLLLLLLLILL | 1 | 0.000424526 | 0.0212263 | 0.00530658 | 14 | RSYSAFALLLGAGTC | MKKLLLLLLLLLILL | + |

**AmpC vs VIM**

| Query_ID (AmpC) | Target_ID (VIM) | Optimal_offset | p-value | E-value | q-value | Overlap | Query_consensus | Target_consensus | Orientation |
| --- | --- | --- | --- | --- | --- | --- | --- | --- | --- |
| CALALLLIGQSAC | MSLLRGAVFALLVGVAGCRVSSAAPQPTA.1 | 5 | 0.00225844 | 0.112922 | 0.0392759 | 13 | FALALLLSGASAM | MSLLRGATFALLVGVAACSVSSAAAQATA | + |
| CALALLLIGQSAC | MSLLRGAVFALLVGVAGCRVSSAAPQPTA.2 | 5 | 0.00225844 | 0.112922 | 0.0392759 | 13 | FALALLLSGASAM | MSLLRGATFALLVGVAACSVSSAAAQATA | + |
| CALALLLIGQSAC | CPLLLLLTACASTPS.1 | -1 | 0.00314207 | 0.157103 | 0.0392759 | 12 | FALALLLSGASAM | CPLLLLLTACASTPS | + |
| CALALLLIGQSAC | CPLLLLLTACASTPS.2 | -1 | 0.00314207 | 0.157103 | 0.0392759 | 12 | FALALLLSGASAM | CPLLLLLTACASTPS | + |
| MESNSI | MTSRSLRV.1 | 0 | 0.00143044 | 0.0715222 | 0.0357611 | 6 | MESNSI | MTSRSLRV | + |
| MESNSI | MTSRSLRV.2 | 0 | 0.00143044 | 0.0715222 | 0.0357611 | 6 | MESNSI | MTSRSLRV | + |
| MLDYYRNWQPVYPPGTQRLYSNPSIGLFGYLAARSLGQPFDQLMEQDLFP | TYPGGT.1 | -10 | 0.00153179 | 0.0765897 | 0.0382949 | 6 | MLDYYRNWQPVYPPGTQRLYSNPSIGLFGYLAARSLGQPFQQLMEQDLFP | TYPGGT | + |
| MLDYYRNWQPVYPPGTQRLYSNPSIGLFGYLAARSLGQPFDQLMEQDLFP | TYPGGT.2 | -10 | 0.00153179 | 0.0765897 | 0.0382949 | 6 | MLDYYRNWQPVYPPGTQRLYSNPSIGLFGYLAARSLGQPFQQLMEQDLFP | TYPGGT | + |
| MPKKNQKK | LKKASR.1 | -1 | 0.000566999 | 0.02835 | 0.014175 | 6 | MPKKNQKK | LKKASR | + |
| MPKKNQKK | LKKASR.2 | -1 | 0.000566999 | 0.02835 | 0.014175 | 6 | MPKKNQKK | LKKASR | + |
| MSSINPSK | MRSPFWS.1 | 0 | 0.0015292 | 0.0764602 | 0.0382301 | 7 | MSSINLSK | MRSPFWS | + |
| MSSINPSK | MRSPFWS.2 | 0 | 0.0015292 | 0.0764602 | 0.0382301 | 7 | MSSINLSK | MRSPFWS | + |
| MSSINPSK | MSLLRGAVFALLVGVAGCRVSSAAPQPTA.1 | 0 | 0.00379673 | 0.189836 | 0.0474591 | 8 | MSSINLSK | MSLLRGATFALLVGVAACSVSSAAAQATA | + |
| MSSINPSK | MSLLRGAVFALLVGVAGCRVSSAAPQPTA.2 | 0 | 0.00379673 | 0.189836 | 0.0474591 | 8 | MSSINLSK | MSLLRGATFALLVGVAACSVSSAAAQATA | + |
| RPLLAALALLLASQS | MSLLRGAVFALLVGVAGCRVSSAAPQPTA.1 | 2 | 0.00252446 | 0.126223 | 0.0487002 | 15 | RPLLAALALLLASQS | MSLLRGATFALLVGVAACSVSSAAAQATA | + |
| RPLLAALALLLASQS | MSLLRGAVFALLVGVAGCRVSSAAPQPTA.2 | 2 | 0.00252446 | 0.126223 | 0.0487002 | 15 | RPLLAALALLLASQS | MSLLRGATFALLVGVAACSVSSAAAQATA | + |
| RPLLAALALLLASQS | CPLLLLLTACASTPS.1 | 0 | 0.00389601 | 0.194801 | 0.0487002 | 15 | RPLLAALALLLASQS | CPLLLLLTACASTPS | + |
| RPLLAALALLLASQS | CPLLLLLTACASTPS.2 | 0 | 0.00389601 | 0.194801 | 0.0487002 | 15 | RPLLAALALLLASQS | CPLLLLLTACASTPS | + |
| RSYSAFALLLGAGTC | MSLLRGAVFALLVGVAGCRVSSAAPQPTA.1 | 3 | 9.52699e-05 | 0.00476349 | 0.00238175 | 15 | RSYSAFALLLGAGTC | MSLLRGATFALLVGVAACSVSSAAAQATA | + |
| RSYSAFALLLGAGTC | MSLLRGAVFALLVGVAGCRVSSAAPQPTA.2 | 3 | 9.52699e-05 | 0.00476349 | 0.00238175 | 15 | RSYSAFALLLGAGTC | MSLLRGATFALLVGVAACSVSSAAAQATA | + |
| YFQQWKPTYAPGTHRLYSNPSJGLFGYLAAQSLGQPFBQLMEKTLLPKLG | TYPGGT.1 | -7 | 0.00185503 | 0.0927515 | 0.0463758 | 6 | YFQQWKPTYAPGTHRLYSNPSIGLFGYLAAQSLGQPFDQLMEKTLLPKLG | TYPGGT | + |
| YFQQWKPTYAPGTHRLYSNPSJGLFGYLAAQSLGQPFBQLMEKTLLPKLG | TYPGGT.2 | -7 | 0.00185503 | 0.0927515 | 0.0463758 | 6 | YFQQWKPTYAPGTHRLYSNPSIGLFGYLAAQSLGQPFDQLMEKTLLPKLG | TYPGGT | + |
| IPGLAVAITVNGQPHYFNYGVASKDTGQAVSEBTLFEIGSVSKTFTATLA | AKLAKEQGYEVPBPTLDELTT | -19 | 0.000885923 | 0.0442962 | 0.0442962 | 21 | IPGLAVAITVNGQAHYFNYGVASKDTGQAVSENTLFEIGSVSKTFTATLA | AKLAKEQGYEVPNPSLDELTT | + |
| IPGMAVAVVHNGKAHYFNYGVASKETGQPVTEDTLFEIGSVSKTFTATLG | AKLAKEQGYEVPBPTLDELTT | -19 | 0.000936769 | 0.0468384 | 0.0468384 | 21 | IPGMAVAVVHNGKAHYFNYGVASKETGQPVTEDTLFEIGSVSKTFTATLG | AKLAKEQGYEVPNPSLDELTT | + |
| QQAIATTHTGYYTVGGMTQGLGWERYPYPITLQALLAGNSTPMAMZPHKV | LLAFLACAATASAAPPKTTST | -31 | 0.000777532 | 0.0388766 | 0.0388766 | 19 | QQAIATTHTGYYTVGGMTQGLGWERYPYPITLQALLAGNSTPMAMEPHKV | LLAFLACAATASAAPPKTTSS | + |
| RNJLAGALLLAAGTC | MKRJJLLLLJL | 1 | 0.00020824 | 0.010412 | 0.010412 | 10 | RNLLAGALLLAAGTC | MKRILLLLLLL | + |
| RNJLAGALLLAAGTC | CPLLLLLTACASTPS | -3 | 0.00143293 | 0.0716466 | 0.0358233 | 12 | RNLLAGALLLAAGTC | CPLLLLLTACASTPS | + |
| AAAQLRAVVDAAVKPLMQQQG | AEDTJGLPVRAAVVT | 0 | 0.00079097 | 0.0395485 | 0.0395485 | 15 | AAAQLRAVVDAAVKPLMQQQG | AEDTLGLPVRAAVVT | + |
| AADRLEALVDAAVQPVMQQQD | AEDTJGLPVRAAVVT | 0 | 0.000560223 | 0.0280111 | 0.0280111 | 15 | AADRLEALVDAAVQPVMQQQD | AEDTLGLPVRAAVVT | + |
| IPGLAVAVTVBGKAHYFNYGVASKETGQPVSEBTLFEIGSVSKTFTATLA | KLAKEQGYEVPBPILDELTTL | -20 | 0.000639191 | 0.0319595 | 0.0319595 | 21 | IPGLAVAVTVNGKAHYFNYGVASKETGQPVSENTLFEIGSVSKTFTATLA | KLAKEQGYEVPNPILDELTTL | + |
| IPGMAVAITHKGQRHYFNYGVASKETGQAVTEDTLFEIGSVSKTFTATLG | KLAKEQGYEVPBPILDELTTL | -20 | 0.000205886 | 0.0102943 | 0.0102943 | 21 | IPGMAVAITHKGQRHYFNYGVASKETGQAVTEDTLFEIGSVSKTFTATLG | KLAKEQGYEVPNPILDELTTL | + |
| MCRRRL | MTPLRL | 0 | 0.000102469 | 0.00512345 | 0.00512345 | 6 | MTRQRL | MTSLRL | + |
| MCRRRL | MTSRSLRV | 0 | 0.000419316 | 0.0209658 | 0.0104829 | 6 | MTRQRL | MTSRSLRV | + |
| VRVPADQMARYAQGY | MPRASKQQARYAVGRCLMLWSSNDVTQQGSRPKTKLCRTHP | 0 | 0.000838039 | 0.041902 | 0.041902 | 15 | VRVPADQMARYAQGY | MPRASKQQARYAVGRCLMLWSSNDVTQQGSRPKTKLCRTHP | + |

**AmpC vs NDM**

| Query_ID | Target_ID | Optimal_offset | p-value | E-value | q-value | Overlap | Query_consensus | Target_consensus | Orientation |
| --- | --- | --- | --- | --- | --- | --- | --- | --- | --- |
| MQQRRAFALLTLGSLLLAPCT | IGRCLA | -12 | 0.00126975 | 0.0419016 | 0.0419016 | 6 | MQQRRAFALLTLGSLLLAPCT | IGLLLA | + |
| QPVERLSPPQP | MELPNIMHPVAKLSTALAAALMLSGCMPG | 7 | 0.000644672 | 0.0212742 | 0.0212742 | 11 | QPVERLSPPQP | MELPNIMHPVAKLSTALAAALMLSGCMPG | + |
| THRGYYQVGDMTQGLGWERYAYPISLERLQAGNSAEMALZP | VVTHAHQDKMGGMDALHAAGIATYANALSNQLAPQEGMVAAQHSLTFAAN | 2 | 0.000901566 | 0.0297517 | 0.0297517 | 41 | THRGYYQVGDMTQGLGWERYAYPISLERLQAGNSAEMALQP | VVTHAHQDKMGGMDALHAAGIATYANALSNQLAPQEGMVAAQHSLTFAAN | + |
| DRAASMLS | KLSTALAAALMLSGC | 5 | 0.000369494 | 0.0121933 | 0.0121933 | 8 | DRAASMLS | KLSTALAAALMLSGC | + |
| DRAASMLS | MELPNIMHPVAKLSTALAAALMLSGCMPG | 16 | 0.00116658 | 0.0384972 | 0.0192486 | 8 | DRAASMLS | MELPNIMHPVAKLSTALAAALMLSGCMPG | + |
| DRAASMLS | RRALCR | -1 | 0.00263512 | 0.0869589 | 0.0289863 | 6 | DRAASMLS | AAALML | + |
| KTJAACMAJLAAGQC | DALHAAGI | -6 | 0.00187873 | 0.061998 | 0.0433283 | 8 | KTIAACMAILAAGQC | DALHAAGI | + |
| KTJAACMAJLAAGQC | KLSTALAAALMLSGC | 0 | 0.00262596 | 0.0866566 | 0.0433283 | 15 | KTIAACMAILAAGQC | KLSTALAAALMLSGC | + |
| LSLSDKASQHLPALKGSAFDHISVLQLGTYTAGGLPLQFPDEADSADKML | WVEPATAPNFGPLKVFYPGPGHTSDNITVGIDGTDIAFGGCLIKDSKAKS.2 | 5 | 0.00100378 | 0.0331247 | 0.0171303 | 45 | LSLSDKASQHLPALKGSAFDHISVLQLGTYTAGGLPLQFPDEADSADKML | WVEPATAPNFGPLKVFYPGPGHTSDNITVGIDGTDIAFGGCLIKDSKAKS | + |
| LSLSDKASQHLPALKGSAFDHISVLQLGTYTAGGLPLQFPDEADSADKML | WVEPATAPNFGPLKVFYPGPGHTSDNITVGIDGTDIAFGGCLIKDSKAKS.1 | 5 | 0.0010382 | 0.0342606 | 0.0171303 | 45 | LSLSDKASQHLPALKGSAFDHISVLQLGTYTAGGLPLQFPDEADSADKML | WVEPATAPNFGPLKVFYPGPGHTSDNITVGIDGTDIAFGGCLIKDSKAKS | + |
| MESNSIMP | MELPNIMHPVAKLSTALAAALMLSGCMPG | 0 | 0.000920237 | 0.0303678 | 0.0303678 | 8 | MESNSIMP | MELPNIMHPVAKLSTALAAALMLSGCMPG | + |
| MHQKTNLKPGILAAFTLSL | MELPNIMHPVAKLSTALAAALMLSGCMPG | 6 | 0.000280359 | 0.00925185 | 0.00925185 | 19 | MHQKTNLKPGILAAFTLSL | MELPNIMHPVAKLSTALAAALMLSGCMPG | + |
| MQQRRAFALLTLGSLLLAPCT | IGRCLA | -12 | 0.000371935 | 0.0122739 | 0.0122739 | 6 | MQQRRAFALLTLGSLLLAPCT | IGLLLA | + |
| MRFTITALALLLIGQ | KLSTALAAALMLSGC | 0 | 0.000752922 | 0.0248464 | 0.0248464 | 15 | MRLTITALALLLIGQ | KLSTALAAALMLSGC | + |
| MRFTITALALLLIGQ | MELPNIMHPVAKLSTALAAALMLSGCMPG | 11 | 0.00282166 | 0.0931149 | 0.0465575 | 15 | MRLTITALALLLIGQ | MELPNIMHPVAKLSTALAAALMLSGCMPG | + |
| MRPSLRPLLVALGLC | SIQHFRVALIPFFAAFCLPVF | 4 | 0.00226834 | 0.0748554 | 0.0419112 | 15 | MRPSLRPLLVALGLC | SIQHFRVALIPFFAAFCLPVF | + |
| MRPSLRPLLVALGLC | INLPVALA | -5 | 0.00254007 | 0.0838223 | 0.0419112 | 8 | MRPSLRPLLVALGLC | INLPVALA | + |

**IMP vs AmpC**

| Query_ID | Target_ID | Optimal_offset | p-value | E-value | q-value | Overlap | Query_consensus | Target_consensus | Orientation |
| --- | --- | --- | --- | --- | --- | --- | --- | --- | --- |
| KGLGNLGDANIEAWPKSAKKVKSK | EANMGYQGDAA | 1 | 0.000869434 | 0.0434717 | 0.0434717 | 10 | KGLGNLGDANIEAWPKSAKKVKSK | EANMGYQGDAA | + |
| LVPANGLVVLDNKEAYJIDTPWTAKDTEKLVTWIEER | IPGLAVAVTVBGKAHYFNYGVASKETGQPVSEBTLFEIGSVSKTFTATLA | -1 | 0.000485832 | 0.0242916 | 0.0242916 | 36 | LVPANGLVVLDNKEAYLIDTPWTAKDTEKLVTWIVER | IPGLAVAVTVNGKAHYFNYGVASKETGQPVSENTLFEIGSVSKTFTATLA | + |
| MHPPLRPHPGWKRLGPALLALLLGVFGAR | RPLLAALALLLASQS | -12 | 0.00015113 | 0.00755648 | 0.00377824 | 15 | MHPPLFKHPLFKSLGLALLALLLGAAGAR | RPLLAALALLLASQS | + |
| MHPPLRPHPGWKRLGPALLALLLGVFGAR | RSYSAFALLLGAGTC | -13 | 0.00015113 | 0.00755648 | 0.00377824 | 15 | MHPPLFKHPLFKSLGLALLALLLGAAGAR | RSYSAFALLLGAGTC | + |
| MKFLAPFLFLLPCVTVATEAS | RPLLAALALLLASQS | -1 | 5.44152e-05 | 0.00272076 | 0.00272076 | 15 | MKFLAPFLFLLPCVTVATEAS | RPLLAALALLLASQS | + |
| MKFLAPFLFLLPCVTVATEAS | RSYSAFALLLGAGTC | -1 | 0.000330397 | 0.0165198 | 0.00825992 | 15 | MKFLAPFLFLLPCVTVATEAS | RSYSAFALLLGAGTC | + |
| MKKLJLLLLLLLILL | RPLLAALALLLASQS | -1 | 7.1651e-06 | 0.000358255 | 0.000358255 | 14 | MKKLLLLLLLLLILL | RPLLAALALLLASQS | + |
| MKKLJLLLLLLLILL | MQQRRAFALLTLGSLLLAPCT | 2 | 0.000721803 | 0.0360902 | 0.0123735 | 15 | MKKLLLLLLLLLILL | MQQRRAFALLTLGSLLLAPCT | + |
| MKKLJLLLLLLLILL | RSYSAFALLLGAGTC | -1 | 0.000742408 | 0.0371204 | 0.0123735 | 14 | MKKLLLLLLLLLILL | RSYSAFALLLGAGTC | + |
| MKYJJLJJLLL | RSYSAFALLLGAGTC | -1 | 4.67226e-05 | 0.00233613 | 0.00233613 | 10 | MKYILLLLLLL | RSYSAFALLLGAGTC | + |
| MKYJJLJJLLL | RPLLAALALLLASQS | -1 | 0.000153943 | 0.00769714 | 0.00384857 | 10 | MKYILLLLLLL | RPLLAALALLLASQS | + |
| MPNRALWQRIAILLJMPILSLPLLAANLT | MQQRRAFALLTLGSLLLAPCT | -5 | 0.00095674 | 0.047837 | 0.047837 | 21 | MPNRALWQRIAILLLMPILSLPLLAANLT | MQQRRAFALLTLGSLLLAPCT | + |
| MFSSIN | MSSINPSK | -1 | 0.000139207 | 0.00696035 | 0.00696035 | 5 | MFSSIN | MSSINLSK | + |
| MKKLJLLLLJL | RNJLAGALLLAAGTC | -1 | 2.86127e-06 | 0.000143063 | 0.000143063 | 10 | MKKLILLLLIL | RNLLAGALLLAAGTC | + |
| MKKLLVFFJFLFCSI | RNJLAGALLLAAGTC | -1 | 3.3345e-05 | 0.00166725 | 0.00166725 | 14 | MKKLLVFFIFLFCSI | RNLLAGALLLAAGTC | + |
| MKYIJLLLLLI | RNJLAGALLLAAGTC | -1 | 0.000139194 | 0.00695969 | 0.00695969 | 10 | MKYIILLLLLI | RNLLAGALLLAAGTC | + |
| DVTFATDTTFTVGKQ | FAACTC | -3 | 0.000609888 | 0.0304944 | 0.0304944 | 6 | DVTFATDTTFTVGGQ | FAAGTT | + |
| MKKJFLLLJFL | MLSGVRNLLISALLLGAGNCM | 4 | 0.000226488 | 0.0113244 | 0.0113244 | 11 | MKKLFLLLLFL | MLSGVRNLLISALLLGAGNCM | + |
| MKLAIALFLAYILLLVPSTHASQEQFKSATNQLEPTKGQLS | MQQRRAFALLTLGSLLLAPCT | 2 | 0.000308401 | 0.0154201 | 0.0154201 | 19 | MKLAIALFLAYILLLVPSTHASQEQFKSATNQLEPTKGQLS | MQQRRAFALLTLGSLLLAPCT | + |
| MRIAIRGCAALMIGAIANAQA | MRFTITALALLLIGQ | 0 | 0.00016618 | 0.00830898 | 0.00830898 | 15 | MRIAISGCAALMIGAIANAQA | MRLTITALALLLIGQ | + |
| SDEISSNTFELADGV | HIAKSILAGTFFLLLGTSHGF | 2 | 0.000314017 | 0.0157009 | 0.0157009 | 15 | SDEISSNTFELADGV | HIAKSILAGTFFLLLGTSHGF | + |

**NDM vs AmpC**

| Query_ID | Target_ID | Optimal_offset | p-value | E-value | q-value | Overlap | Query_consensus | Target_consensus | Orientation |
| --- | --- | --- | --- | --- | --- | --- | --- | --- | --- |
| IGRCLA | MQQRRAFALLTLGSLLLAPCT | 12 | 0.000445087 | 0.0222544 | 0.0222544 | 6 | IGLLLA | MQQRRAFALLTLGSLLLAPCT | + |
| KLSTALAAALMLSGC | DRAASMLS | -5 | 0.000873212 | 0.0436606 | 0.0436606 | 8 | KLSTALAAALMLSGC | DRAASMLS | + |
| MELPNIMHPVAKLSTALAAALMLSGCMPG | DRAASMLS | -16 | 0.000958715 | 0.0479358 | 0.0309436 | 8 | MELPNIMHPVAKLSTALAAALMLSGCMPG | DRAASMLS | + |
| MELPNIMHPVAKLSTALAAALMLSGCMPG | MHQKTNLKPGILAAFTLSL | -6 | 0.00123774 | 0.0618872 | 0.0309436 | 19 | MELPNIMHPVAKLSTALAAALMLSGCMPG | MHQKTNLKPGILAAFTLSL | + |
| RRALCR | DRAASMLS | 1 | 0.000525305 | 0.0262653 | 0.0262653 | 6 | AAALML | DRAASMLS | + |
| WVEPATAPNFGPLKVFYPGPGHTSDNITVGIDGTDIAFGGCLIKDSKAKS.1 | LSLSDKASQHLPALKGSAFDHISVLQLGTYTAGGLPLQFPDEADSADKML | -5 | 0.000122028 | 0.00610142 | 0.00610142 | 45 | WVEPATAPNFGPLKVFYPGPGHTSDNITVGIDGTDIAFGGCLIKDSKAKS | LSLSDKASQHLPALKGSAFDHISVLQLGTYTAGGLPLQFPDEADSADKML | + |
| WVEPATAPNFGPLKVFYPGPGHTSDNITVGIDGTDIAFGGCLIKDSKAKS.2 | LSLSDKASQHLPALKGSAFDHISVLQLGTYTAGGLPLQFPDEADSADKML | -5 | 0.000114959 | 0.00574795 | 0.00574795 | 45 | WVEPATAPNFGPLKVFYPGPGHTSDNITVGIDGTDIAFGGCLIKDSKAKS | LSLSDKASQHLPALKGSAFDHISVLQLGTYTAGGLPLQFPDEADSADKML | + |

**VIM vs AmpC**

| Query_ID | Target_ID | Optimal_offset | p-value | E-value | q-value | Overlap | Query_consensus | Target_consensus | Orientation |
| --- | --- | --- | --- | --- | --- | --- | --- | --- | --- |
| MSLLRGAVFALLVGVAGCRVSSAAPQPTA.1 | RSYSAFALLLGAGTC | -3 | 6.80493e-05 | 0.00340247 | 0.00340247 | 15 | MSLLRGAVFALLVGVAACRVSSAAAQATA | RSYSAFALLLGAGTC | + |
| MSLLRGAVFALLVGVAGCRVSSAAPQPTA.1 | RPLLAALALLLASQS | -2 | 0.000273872 | 0.0136936 | 0.00684681 | 15 | MSLLRGAVFALLVGVAACRVSSAAAQATA | RPLLAALALLLASQS | + |
| MSLLRGAVFALLVGVAGCRVSSAAPQPTA.1 | CALALLLIGQSAC | -5 | 0.00100125 | 0.0500623 | 0.0166874 | 13 | MSLLRGAVFALLVGVAACRVSSAAAQATA | FALALLLSGATAM | + |
| MSLLRGAVFALLVGVAGCRVSSAAPQPTA.2 | RSYSAFALLLGAGTC | -3 | 6.80493e-05 | 0.00340247 | 0.00340247 | 15 | MSLLRGAVFALLVGVAACRVSSAAAQATA | RSYSAFALLLGAGTC | + |
| MSLLRGAVFALLVGVAGCRVSSAAPQPTA.2 | RPLLAALALLLASQS | -2 | 0.000273872 | 0.0136936 | 0.00684681 | 15 | MSLLRGAVFALLVGVAACRVSSAAAQATA | RPLLAALALLLASQS | + |
| MSLLRGAVFALLVGVAGCRVSSAAPQPTA.2 | CALALLLIGQSAC | -5 | 0.00100125 | 0.0500623 | 0.0166874 | 13 | MSLLRGAVFALLVGVAACRVSSAAAQATA | FALALLLSGATAM | + |
| TYPGGT.1 | MLDYYRNWQPVYPPGTQRLYSNPSIGLFGYLAARSLGQPFDQLMEQDLFP | 10 | 0.000815818 | 0.0407909 | 0.030262 | 6 | TYPGGT | MLDYYRNWQPVYPPGTQRLYSNPSIGLFGYLAARSLGQPFDQLMEQDLFP | + |
| TYPGGT.1 | YFQQWKPTYAPGTHRLYSNPSJGLFGYLAAQSLGQPFBQLMEKTLLPKLG | 7 | 0.00121048 | 0.0605239 | 0.030262 | 6 | TYPGGT | YFQQWKPTYAPGTHRLYSNPSIGLFGYLAAQSLGQPFDQLMEKTLLPKLG | + |
| TYPGGT.2 | MLDYYRNWQPVYPPGTQRLYSNPSIGLFGYLAARSLGQPFDQLMEQDLFP | 10 | 0.000815818 | 0.0407909 | 0.030262 | 6 | TYPGGT | MLDYYRNWQPVYPPGTQRLYSNPSIGLFGYLAARSLGQPFDQLMEQDLFP | + |
| TYPGGT.2 | YFQQWKPTYAPGTHRLYSNPSJGLFGYLAAQSLGQPFBQLMEKTLLPKLG | 7 | 0.00121048 | 0.0605239 | 0.030262 | 6 | TYPGGT | YFQQWKPTYAPGTHRLYSNPSIGLFGYLAAQSLGQPFDQLMEKTLLPKLG | + |
| CPLLLLLTACASTPS | RNJLAGALLLAAGTC | 3 | 0.000973113 | 0.0486556 | 0.0486556 | 12 | CPLLLLLTACASTPS | RNLLAGALLLAAGTC | + |
| MKRJJLLLLJL | RNJLAGALLLAAGTC | -1 | 4.29811e-05 | 0.00214906 | 0.00214906 | 10 | MKRILLLLLLL | RNLLAGALLLAAGTC | + |
| MRSRNWSRTLTERSGGNGAVAVFMACYDCFFVQSMPRASKQQARYAVGRC | PANVLLNKTGSTGGFGAYVAFVPSKDIGIVILANKNYPNAERVKIAHAIL | -2 | 0.00187761 | 0.0938807 | 0.0486051 | 48 | MRSRNWSRTLTERSGGNGAVAVFMACYDCFFVQSMPRASKQQARYAVGRC | PANVLLNKTGSTGGFGAYVAFVPSKDIGIVILANKNYPNAERVKIAHAIL | + |
| MRSRNWSRTLTERSGGNGAVAVFMACYDCFFVQSMPRASKQQARYAVGRC | LYNKTGSTGGFGAYVAYVPSKDIGIVILANKNYPNAERVKAAHAILSALD | -6 | 0.0019442 | 0.0972102 | 0.0486051 | 44 | MRSRNWSRTLTERSGGNGAVAVFMACYDCFFVQSMPRASKQQARYAVGRC | LYNKTGSTGGFGAYVAYVPSKDIGIVILANKNYPNAERVKAAHAILSALD | + |
| AEDTJGLPVRAAVVT | AADRLEALVDAAVQPVMQQQD | 0 | 0.000347705 | 0.0173853 | 0.0104088 | 15 | AEDTLGLPVRAAVVT | AADRLEALVDAAVQPVMQQQD | + |
| AEDTJGLPVRAAVVT | AAAQLRAVVDAAVKPLMQQQG | 0 | 0.00041635 | 0.0208175 | 0.0104088 | 15 | AEDTLGLPVRAAVVT | AAAQLRAVVDAAVKPLMQQQG | + |
| EWIKTKLKKPVKKAI | MHQKTNLKPGILAAFTLSL | 0 | 0.000588399 | 0.0294199 | 0.0294199 | 15 | EWIKTKLKKPVKKAI | MHQKTNLKPGILAAFTLSL | + |
| KLAKEQGYEVPBPILDELTTL | IPGMAVAITHKGQRHYFNYGVASKETGQAVTEDTLFEIGSVSKTFTATLG | 20 | 0.000310069 | 0.0155035 | 0.0155035 | 21 | KLAKEQGYEVPNPILDELTTL | IPGMAVAITHKGQRHYFNYGVASKETGQAVTEDTLFEIGSVSKTFTATLG | + |
| KLAKEQGYEVPBPILDELTTL | IPGLAVAVTVBGKAHYFNYGVASKETGQPVSEBTLFEIGSVSKTFTATLA | 20 | 0.000911119 | 0.0455559 | 0.022778 | 21 | KLAKEQGYEVPNPILDELTTL | IPGLAVAVTVNGKAHYFNYGVASKETGQPVSENTLFEIGSVSKTFTATLA | + |
| MRSRNWSRTLTERSGGNGAVAVFMACYDC | QQRQSILWGAVATLMWAGLAH | -9 | 0.000712548 | 0.0356274 | 0.0356274 | 20 | MRSRNWSRTLTERSGGNGAVAVFMACYDC | QQRQSILWGAVATLMWAGLAH | + |
| MTPLRL | MCRRRL | 0 | 0.000658183 | 0.0329092 | 0.0329092 | 6 | MTSLRL | MTRRRL | + |
| NCYAGFAH | QQRQSILWGAVATLMWAGLAH | 13 | 0.00129214 | 0.064607 | 0.0446131 | 8 | NCYAGFAH | QQRQSILWGAVATLMWAGLAH | + |
| NCYAGFAH | CLCGFA | -1 | 0.00178452 | 0.0892261 | 0.0446131 | 6 | NCYAGFAH | TLCLIA | + |
| RIFISIVFFMQTFGLVFAEPD | MSSIKLSKVFSYSAFGLFFGAAACLAATP | 2 | 0.000269689 | 0.0134844 | 0.0134844 | 21 | RIFISIVFFMQTFGLVFAEPD | MSSIKLSKVFSYSAFGLFFGAAACLAATP | + |

**IMP vs NDM**

| Query_ID | Target_ID | Optimal_offset | p-value | E-value | q-value | Overlap | Query_consensus | Target_consensus | Orientation |
| --- | --- | --- | --- | --- | --- | --- | --- | --- | --- |
| ADADLDAWPASIAKVQARYPDAKIVVPGHGK | LGDADTEHYAASARAFGAAFPKASMIVMSHSAPDSRAAITHTARMADKLR.1 | 1 | 9.90197e-07 | 3.26765e-05 | 1.72686e-05 | 31 | ADADLDAWPASIAKVQARYPDAKIVVPGHGK | LGDADTEHYAASARAFGAAFPKASMIVMSHSAPDSRAAITHTARMADKLR | + |
| ADADLDAWPASIAKVQARYPDAKIVVPGHGK | LGDADTEHYAASARAFGAAFPKASMIVMSHSAPDSRAAITHTARMADKLR.2 | 1 | 1.04658e-06 | 3.45372e-05 | 1.72686e-05 | 31 | ADADLDAWPASIAKVQARYPDAKIVVPGHGK | LGDADTEHYAASARAFGAAFPKASMIVMSHSAPDSRAAITHTARMADKLR | + |
| DQQEAIIFDTPATDQASEELI | WQHTSYLDMPGFGAVASNGLIVRDGGRVLVVDTAWTDDQTAQILNWIKQE | 23 | 0.00042186 | 0.0139214 | 0.00733233 | 21 | DQQEAIIFDTPATDQASEELI | WQHTSYLDMPGFGAVASNGLIVRDGGRVLVVDTAWTDDQTAQILNWIKQE | + |
| DQQEAIIFDTPATDQASEELI | VRDGGRVLVVDTAWTDDQTAQILNWIKQEINLPVALAVVTHAHQDKMGGM | 2 | 0.000444383 | 0.0146647 | 0.00733233 | 21 | DQQEAIIFDTPATDQASEELI | VRDGGRVLVVDTAWTDDQTAQILNWIKQEINLPVALAVVTHAHQDKMGGM | + |
| DQQEAIIFDTPATDQASEELI | ASNGLI | -15 | 0.00103468 | 0.0341446 | 0.0113815 | 6 | DQQEAIIFDTPATDQASEELI | ASNGLI | + |
| EVFYPGAGHTKDNVVVWLPEEKILFGGCFVKS | WVEPATAPNFGPLKVFYPGPGHTSDNITVGIDGTDIAFGGCLIKDSKAKS.2 | 13 | 1.22787e-18 | 4.05195e-17 | 2.19445e-17 | 32 | EVFYPGAGHTKDNVVVWLPEEKILFGGCFVKS | WVEPATAPNFGPLKVFYPGPGHTSDNITVGIDGTDIAFGGCLIKDSKAKS | + |
| EVFYPGAGHTKDNVVVWLPEEKILFGGCFVKS | WVEPATAPNFGPLKVFYPGPGHTSDNITVGIDGTDIAFGGCLIKDSKAKS.1 | 13 | 1.32997e-18 | 4.3889e-17 | 2.19445e-17 | 32 | EVFYPGAGHTKDNVVVWLPEEKILFGGCFVKS | WVEPATAPNFGPLKVFYPGPGHTSDNITVGIDGTDIAFGGCLIKDSKAKS | + |
| EVFYPGAGHTKDNVVVWLPEEKILFGGCFVKS | GCFGKD | -26 | 0.00366917 | 0.121083 | 0.0403609 | 6 | EVFYPGAGHTKDNVVVWLPEEKILFGGCFVKS | AGLGAV | + |
| GKDAYJIDTPWTEADTEKLVDWIEQQGLTLKASVSTHSHEDRTGGIGYLN | VRDGGRVLVVDTAWTDDQTAQILNWIKQEINLPVALAVVTHAHQDKMGGM | 3 | 7.66675e-06 | 0.000253003 | 0.000253003 | 47 | GKDAYIIDTPWTEADTEKLVDWIEQQGLTLKASVSTHSHEDRTGGIGYLN | VRDGGRVLVVDTAWTDDQTAQILNWIKQEINLPVALAVVTHAHQDKMGGM | + |
| GKDAYJIDTPWTEADTEKLVDWIEQQGLTLKASVSTHSHEDRTGGIGYLN | WQHTSYLDMPGFGAVASNGLIVRDGGRVLVVDTAWTDDQTAQILNWIKQE | 24 | 0.0011003 | 0.0363099 | 0.018155 | 26 | GKDAYIIDTPWTEADTEKLVDWIEQQGLTLKASVSTHSHEDRTGGIGYLN | WQHTSYLDMPGFGAVASNGLIVRDGGRVLVVDTAWTDDQTAQILNWIKQE | + |
| GVLFPSHGLVVSTKDGAVLVDTGWGNEPTEQLLA | WQHTSYLDMPGFGAVASNGLIVRDGGRVLVVDTAWTDDQTAQILNWIKQE | 11 | 6.58382e-10 | 2.17266e-08 | 2.17266e-08 | 34 | GVLFPSHGLVVSTKGGAVLVDTGWGNEPTEQLLA | WQHTSYLDMPGFGAVASNGLIVRDGGRVLVVDTAWTDDQTAQILNWIKQE | + |
| GVLFPSHGLVVSTKDGAVLVDTGWGNEPTEQLLA | VRDGGRVLVVDTAWTDDQTAQILNWIKQEINLPVALAVVTHAHQDKMGGM | -10 | 0.000140342 | 0.0046313 | 0.00231565 | 24 | GVLFPSHGLVVSTKGGAVLVDTGWGNEPTEQLLA | VRDGGRVLVVDTAWTDDQTAQILNWIKQEINLPVALAVVTHAHQDKMGGM | + |
| GVLFPSHGLVVSTKDGAVLVDTGWGNEPTEQLLA | ASNGLI | -4 | 0.00448518 | 0.148011 | 0.0493369 | 6 | GVLFPSHGLVVSTKGGAVLVDTGWGNEPTEQLLA | ASNGLI | + |
| GYKJKGSISTHFHEDSTGGJEWLNSHSIPTYASELTNELLK | VVTHAHQDKMGGMDALHAAGIATYANALSNQLAPQEGMVAAQHSLTFAAN | -7 | 8.32228e-11 | 2.74635e-09 | 2.74635e-09 | 34 | GYKIKGSISTHFHEDSTGGIEWLNSHSIPTYASELTNELLK | VVTHAHQDKMGGMDALHAAGIATYANALSNQLAPQEGMVAAQHSLTFAAN | + |
| IEVYYPGAGHTKDNLVVWLPKQKJLFGGCLVKSLESKSLGY | WVEPATAPNFGPLKVFYPGPGHTSDNITVGIDGTDIAFGGCLIKDSKAKS.2 | 12 | 6.62699e-17 | 2.18691e-15 | 1.17176e-15 | 38 | IEVFYPGAGHTKDNLVVWLPKQKILFGGCLVKSLESKSLGY | WVEPATAPNFGPLKVFYPGPGHTSDNITVGIDGTDIAFGGCLIKDSKAKS | + |
| IEVYYPGAGHTKDNLVVWLPKQKJLFGGCLVKSLESKSLGY | WVEPATAPNFGPLKVFYPGPGHTSDNITVGIDGTDIAFGGCLIKDSKAKS.1 | 12 | 7.10157e-17 | 2.34352e-15 | 1.17176e-15 | 38 | IEVFYPGAGHTKDNLVVWLPKQKILFGGCLVKSLESKSLGY | WVEPATAPNFGPLKVFYPGPGHTSDNITVGIDGTDIAFGGCLIKDSKAKS | + |
| IPTYASELTNALLAQNGKPLA | VVTHAHQDKMGGMDALHAAGIATYANALSNQLAPQEGMVAAQHSLTFAAN | 20 | 1.0391e-07 | 3.42902e-06 | 3.42902e-06 | 21 | IPTYASELTNALLAQNGKPLA | VVTHAHQDKMGGMDALHAAGIATYANALSNQLAPQEGMVAAQHSLTFAAN | + |
| IPTYASELTNALLAQNGKPLA | TYANALSNQLAPQEGMVAAQHSLTFAANG | -2 | 2.49348e-06 | 8.22848e-05 | 4.11424e-05 | 19 | IPTYASELTNALLAQNGKPLA | TYANALSNQLAPQEGMVAAQHSLTFAANG | + |
| KGLGNLGDANIEAWPKSAKKVKSK | NLGDPDC | -4 | 0.000213345 | 0.00704039 | 0.00704039 | 7 | KGLGNLGDANIEAWPKSAKKVKSK | NLGDADT | + |
| KGLGNLGDANIEAWPKSAKKVKSK | LGDADTEHYAASARAFGAAFPKASMIVMSHSAPDSRAAITHTARMADKLR.1 | -5 | 0.00175314 | 0.0578535 | 0.0212165 | 19 | KGLGNLGDANIEAWPKSAKKVKSK | LGDADTEHYAASARAFGAAFPKASMIVMSHSAPDSRAAITHTARMADKLR | + |
| KGLGNLGDANIEAWPKSAKKVKSK | LGDADTEHYAASARAFGAAFPKASMIVMSHSAPDSRAAITHTARMADKLR.2 | -5 | 0.00192877 | 0.0636495 | 0.0212165 | 19 | KGLGNLGDANIEAWPKSAKKVKSK | LGDADTEHYAASARAFGAAFPKASMIVMSHSAPDSRAAITHTARMADKLR | + |
| LHNLPQASAAIKSGV | INLPVALA | -1 | 0.000552515 | 0.018233 | 0.018233 | 8 | LHNLPQASAAIKSGV | INLPVALA | + |
| LVPANGLVVLDNKEAYJIDTPWTAKDTEKLVTWIEER | WQHTSYLDMPGFGAVASNGLIVRDGGRVLVVDTAWTDDQTAQILNWIKQE | 13 | 2.13248e-12 | 7.03718e-11 | 7.03718e-11 | 37 | LVPANGLVVLDNKEAYLIDTPWTAKDTEKLVTWIVER | WQHTSYLDMPGFGAVASNGLIVRDGGRVLVVDTAWTDDQTAQILNWIKQE | + |
| LVPANGLVVLDNKEAYJIDTPWTAKDTEKLVTWIEER | VRDGGRVLVVDTAWTDDQTAQILNWIKQEINLPVALAVVTHAHQDKMGGM | -8 | 1.17778e-05 | 0.000388668 | 0.000194334 | 29 | LVPANGLVVLDNKEAYLIDTPWTAKDTEKLVTWIVER | VRDGGRVLVVDTAWTDDQTAQILNWIKQEINLPVALAVVTHAHQDKMGGM | + |
| LVPANGLVVLDNKEAYJIDTPWTAKDTEKLVTWIEER | ASNGLI | -2 | 0.000211723 | 0.00698687 | 0.00232896 | 6 | LVPANGLVVLDNKEAYLIDTPWTAKDTEKLVTWIVER | ASNGLI | + |
| MHPPLRPHPGWKRLGPALLALLLGVFGAR | MELPNIMHPVAKLSTALAAALMLSGCMPG | 0 | 0.0008237 | 0.0271821 | 0.0271821 | 29 | MHPPLFKHPLFKSLGLALLALLLGAAGAR | MELPNIMHPVAKLSTALAAALMLSGCMPG | + |
| MKKLJLLLLLLLILL | IGRCLA | -4 | 0.000745891 | 0.0246144 | 0.0246144 | 6 | MKKLLLLLLLLLILL | IGLLLA | + |
| MKKLJLLLLLLLILL | KLSTALAAALMLSGC | -2 | 0.00171918 | 0.056733 | 0.0283665 | 13 | MKKLLLLLLLLLILL | KLSTALAAALMLSGC | + |
| MSIQHFRVALIPFFAAFCLPV | SIQHFRVALIPFFAAFCLPVF | -1 | 3.34486e-26 | 1.1038e-24 | 1.1038e-24 | 20 | MSIQHFRVALIPFFAAFCLPV | SIQHFRVALIPFFAAFCLPVF | + |
| PJADGVYLHTSYKZVEGFGLVDSNGLVVV | WQHTSYLDMPGFGAVASNGLIVRDGGRVLVVDTAWTDDQTAQILNWIKQE | -6 | 1.67134e-10 | 5.51544e-09 | 5.51544e-09 | 23 | PLADGVYLHTSYKQVEGFGLVDSNGLVVV | WQHTSYLDMPGFGAVASNGLIVRDGGRVLVVDTAWTDDQTAQILNWIKQE | + |
| PJADGVYLHTSYKZVEGFGLVDSNGLVVV | ASNGLI | -21 | 3.63861e-05 | 0.00120074 | 0.00060037 | 6 | PLADGVYLHTSYKQVEGFGLVDSNGLVVV | ASNGLI | + |
| WVETNLKQPVKAVVATHFHEDCLGGLQAF | VRDGGRVLVVDTAWTDDQTAQILNWIKQEINLPVALAVVTHAHQDKMGGM | 24 | 3.17081e-09 | 1.04637e-07 | 1.04637e-07 | 26 | WVETNLKQPVKAVVATHFHDDCLGGLGAF | VRDGGRVLVVDTAWTDDQTAQILNWIKQEINLPVALAVVTHAHQDKMGGM | + |
| WVETNLKQPVKAVVATHFHEDCLGGLQAF | VVTHAHQDKMGGMDALHAAGIATYANALSNQLAPQEGMVAAQHSLTFAAN | -13 | 1.25398e-05 | 0.000413815 | 0.000206907 | 16 | WVETNLKQPVKAVVATHFHDDCLGGLGAF | VVTHAHQDKMGGMDALHAAGIATYANALSNQLAPQEGMVAAQHSLTFAAN | + |
| ASNKSIQPTAEASAD | ASNGLI | 0 | 0.000124397 | 0.0041051 | 0.0041051 | 6 | ASNKSIQPTAEASAD | ASNGLI | + |
| AYJIDTPWTEKDTEKLVDWIEAQGLTLKASISTHSHZDRTGGIGYLNSKG | VRDGGRVLVVDTAWTDDQTAQILNWIKQEINLPVALAVVTHAHQDKMGGM | 6 | 0.000209927 | 0.00692761 | 0.00692761 | 44 | AYIIDTPWTEKDTEKLVDWIEAQGLTLKASISTHSHEDRTGGIGYLNSKG | VRDGGRVLVVDTAWTDDQTAQILNWIKQEINLPVALAVVTHAHQDKMGGM | + |
| FDTPWTBEPTEQLLAWVKDNLKAPVKAFVPTHWHDDCLGGL | VRDGGRVLVVDTAWTDDQTAQILNWIKQEINLPVALAVVTHAHQDKMGGM | 9 | 8.59328e-19 | 2.83578e-17 | 2.83578e-17 | 41 | FDTPWTDEQTEQLLAWVKDNLKAPVKAFVPTHWHDDCLGGL | VRDGGRVLVVDTAWTDDQTAQILNWIKQEINLPVALAVVTHAHQDKMGGM | + |
| FDTPWTBEPTEQLLAWVKDNLKAPVKAFVPTHWHDDCLGGL | WQHTSYLDMPGFGAVASNGLIVRDGGRVLVVDTAWTDDQTAQILNWIKQE | 30 | 0.000162233 | 0.00535369 | 0.00267684 | 20 | FDTPWTDEQTEQLLAWVKDNLKAPVKAFVPTHWHDDCLGGL | WQHTSYLDMPGFGAVASNGLIVRDGGRVLVVDTAWTDDQTAQILNWIKQE | + |
| GDABVSAWPNSVEKVKKK | LGDADTEHYAASARAFGAAFPKASMIVMSHSAPDSRAAITHTARMADKLR.1 | 1 | 0.00271391 | 0.0895589 | 0.0473723 | 18 | GDANVSAWPNSVEKVKKK | LGDADTEHYAASARAFGAAFPKASMIVMSHSAPDSRAAITHTARMADKLR | + |
| GDABVSAWPNSVEKVKKK | LGDADTEHYAASARAFGAAFPKASMIVMSHSAPDSRAAITHTARMADKLR.2 | 1 | 0.00287105 | 0.0947446 | 0.0473723 | 18 | GDANVSAWPNSVEKVKKK | LGDADTEHYAASARAFGAAFPKASMIVMSHSAPDSRAAITHTARMADKLR | + |
| GPGHTKDNVVVWLPKEKILFGGCLVKSLGAGNLGL | WVEPATAPNFGPLKVFYPGPGHTSDNITVGIDGTDIAFGGCLIKDSKAKS.2 | 18 | 1.28422e-12 | 4.23793e-11 | 2.41541e-11 | 32 | GPGHTKDNVVVWLPKEKILFGGCLVKSLGAGNLGL | WVEPATAPNFGPLKVFYPGPGHTSDNITVGIDGTDIAFGGCLIKDSKAKS | + |
| GPGHTKDNVVVWLPKEKILFGGCLVKSLGAGNLGL | WVEPATAPNFGPLKVFYPGPGHTSDNITVGIDGTDIAFGGCLIKDSKAKS.1 | 18 | 1.46388e-12 | 4.83081e-11 | 2.41541e-11 | 32 | GPGHTKDNVVVWLPKEKILFGGCLVKSLGAGNLGL | WVEPATAPNFGPLKVFYPGPGHTSDNITVGIDGTDIAFGGCLIKDSKAKS | + |
| GPGHTKDNVVVWLPKEKILFGGCLVKSLGAGNLGL | GCFGKD | -21 | 0.00362732 | 0.119702 | 0.0399006 | 6 | GPGHTKDNVVVWLPKEKILFGGCLVKSLGAGNLGL | AGLGAV | + |
| IEKJSENVYLHTSFKZTNGWG | WQHTSYLDMPGFGAVASNGLIVRDGGRVLVVDTAWTDDQTAQILNWIKQE | -8 | 0.00034325 | 0.0113273 | 0.0113273 | 13 | IEKLSENVYLHTSFKETNGWG | WQHTSYLDMPGFGAVASNGLIVRDGGRVLVVDTAWTDDQTAQILNWIKQE | + |
| IEVFYPGAGHTKDNIVVWLPKQKILFGGCLVKS | WVEPATAPNFGPLKVFYPGPGHTSDNITVGIDGTDIAFGGCLIKDSKAKS.2 | 12 | 1.66127e-17 | 5.48218e-16 | 3.19102e-16 | 33 | IEVFYPGAGHTKDNIVVWLPKQKILFGGCLVKS | WVEPATAPNFGPLKVFYPGPGHTSDNITVGIDGTDIAFGGCLIKDSKAKS | + |
| IEVFYPGAGHTKDNIVVWLPKQKILFGGCLVKS | WVEPATAPNFGPLKVFYPGPGHTSDNITVGIDGTDIAFGGCLIKDSKAKS.1 | 12 | 1.93395e-17 | 6.38204e-16 | 3.19102e-16 | 33 | IEVFYPGAGHTKDNIVVWLPKQKILFGGCLVKS | WVEPATAPNFGPLKVFYPGPGHTSDNITVGIDGTDIAFGGCLIKDSKAKS | + |
| IPTYASELTNELLKKKGKPQA | VVTHAHQDKMGGMDALHAAGIATYANALSNQLAPQEGMVAAQHSLTFAAN | 20 | 5.22904e-08 | 1.72558e-06 | 1.72558e-06 | 21 | IPTYASELTNELLKKKGKPQA | VVTHAHQDKMGGMDALHAAGIATYANALSNQLAPQEGMVAAQHSLTFAAN | + |
| IPTYASELTNELLKKKGKPQA | TYANALSNQLAPQEGMVAAQHSLTFAANG | -2 | 1.14031e-06 | 3.76303e-05 | 1.88151e-05 | 19 | IPTYASELTNELLKKKGKPQA | TYANALSNQLAPQEGMVAAQHSLTFAANG | + |
| KJADGVYLHTSYKEVEGFGLVSSNGLVVV | WQHTSYLDMPGFGAVASNGLIVRDGGRVLVVDTAWTDDQTAQILNWIKQE | -6 | 2.4718e-10 | 8.15695e-09 | 8.15695e-09 | 23 | KLADGVYLHTSYKEVEGFGLVSSNGLVVV | WQHTSYLDMPGFGAVASNGLIVRDGGRVLVVDTAWTDDQTAQILNWIKQE | + |
| KJADGVYLHTSYKEVEGFGLVSSNGLVVV | ASNGLI | -21 | 3.98603e-05 | 0.00131539 | 0.000657695 | 6 | KLADGVYLHTSYKEVEGFGLVSSNGLVVV | ASNGLI | + |
| LHNLPQASAAIKSGV | INLPVALA | -1 | 0.000556885 | 0.0183772 | 0.0183772 | 8 | LHNLPQASAAIKSGV | INLPVALA | + |
| MSIQHFRVALIPFFAAFCLPVFAGEQGH | SIQHFRVALIPFFAAFCLPVF | -1 | 1.81357e-28 | 5.98478e-27 | 5.98478e-27 | 21 | MSIQHFRVALIPFFAAFCLPVFALSQGH | SIQHFRVALIPFFAAFCLPVF | + |
| PCNGLVVIDNKEAYJIDTPWNAKDTE | WQHTSYLDMPGFGAVASNGLIVRDGGRVLVVDTAWTDDQTAQILNWIKQE | 15 | 9.61411e-10 | 3.17266e-08 | 3.17266e-08 | 26 | PCNGLVVIDNKEAYLIDTPWNAKDTE | WQHTSYLDMPGFGAVASNGLIVRDGGRVLVVDTAWTDDQTAQILNWIKQE | + |
| PCNGLVVIDNKEAYJIDTPWNAKDTE | ASNGLI | 0 | 0.000219274 | 0.00723604 | 0.00353053 | 6 | PCNGLVVIDNKEAYLIDTPWNAKDTE | ASNGLI | + |
| PCNGLVVIDNKEAYJIDTPWNAKDTE | VRDGGRVLVVDTAWTDDQTAQILNWIKQEINLPVALAVVTHAHQDKMGGM | -6 | 0.000320957 | 0.0105916 | 0.00353053 | 20 | PCNGLVVIDNKEAYLIDTPWNAKDTE | VRDGGRVLVVDTAWTDDQTAQILNWIKQEINLPVALAVVTHAHQDKMGGM | + |
| RGYKJKASISTHFHEDSTGGJEYLNSHSIPTYASELTNELL | VVTHAHQDKMGGMDALHAAGIATYANALSNQLAPQEGMVAAQHSLTFAAN | -8 | 1.64349e-11 | 5.42351e-10 | 5.42351e-10 | 33 | RGYKIKASISTHFHEDSTGGLEYLNSHSIPTYASELTNELL | VVTHAHQDKMGGMDALHAAGIATYANALSNQLAPQEGMVAAQHSLTFAAN | + |
| SLGNTGDADLEAWPASIAKVQAR | NLGDPDC | -3 | 0.000191536 | 0.0063207 | 0.0063207 | 7 | SLGNTGDADLEAWPASIAKVQAR | NLGDADT | + |
| SLGNTGDADLEAWPASIAKVQAR | LGDADTEHYAASARAFGAAFPKASMIVMSHSAPDSRAAITHTARMADKLR.2 | -4 | 0.0016715 | 0.0551593 | 0.0193395 | 19 | SLGNTGDADLEAWPASIAKVQAR | LGDADTEHYAASARAFGAAFPKASMIVMSHSAPDSRAAITHTARMADKLR | + |
| SLGNTGDADLEAWPASIAKVQAR | LGDADTEHYAASARAFGAAFPKASMIVMSHSAPDSRAAITHTARMADKLR.1 | -4 | 0.00175813 | 0.0580184 | 0.0193395 | 19 | SLGNTGDADLEAWPASIAKVQAR | LGDADTEHYAASARAFGAAFPKASMIVMSHSAPDSRAAITHTARMADKLR | + |

**IMP vs VIM**

| ADADLDAWPASIAKVQARYPDAKIVVPGHGK | RARYPEARVVVPGHGAPGGPELLDHTEAL.1 | -15 | 1.33414e-08 | 6.67068e-07 | 3.30203e-07 | 16 | ADADLDAWPASIAKVQARYPDAKIVVPGHGK | RARYPEARVVVPGHGAPGGPELLDHTEAL | + |
| --- | --- | --- | --- | --- | --- | --- | --- | --- | --- |
| ADADLDAWPASIAKVQARYPDAKIVVPGHGK | RARYPEARVVVPGHGAPGGPELLDHTEAL.2 | -15 | 1.33414e-08 | 6.67068e-07 | 3.30203e-07 | 16 | ADADLDAWPASIAKVQARYPDAKIVVPGHGK | RARYPEARVVVPGHGAPGGPELLDHTEAL | + |
| ADADLDAWPASIAKVQARYPDAKIVVPGHGK | AKDLGNLADADVAAWPASLER.1 | 7 | 2.64163e-08 | 1.32081e-06 | 3.30203e-07 | 14 | ADADLDAWPASIAKVQARYPDAKIVVPGHGK | AKDLGNLADADVAAWPASLER | + |
| ADADLDAWPASIAKVQARYPDAKIVVPGHGK | AKDLGNLADADVAAWPASLER.2 | 7 | 2.64163e-08 | 1.32081e-06 | 3.30203e-07 | 14 | ADADLDAWPASIAKVQARYPDAKIVVPGHGK | AKDLGNLADADVAAWPASLER | + |
| AESLPDLKIEKJSENVYLHTSFEZVNGWG | LAEDVRVRRJAPGVWLHVTLA.1 | -2 | 1.5508e-07 | 7.75401e-06 | 3.87701e-06 | 21 | AESLPDLKIEKLSENVYLHTSFEEVNGWG | LAEDVRVRRLAPGVWLHVTLA | + |
| AESLPDLKIEKJSENVYLHTSFEZVNGWG | LAEDVRVRRJAPGVWLHVTLA.2 | -2 | 1.5508e-07 | 7.75401e-06 | 3.87701e-06 | 21 | AESLPDLKIEKLSENVYLHTSFEEVNGWG | LAEDVRVRRLAPGVWLHVTLA | + |
| AKJVVPGHGEVGDASLLK | RARYPEARVVVPGHGAPGGPELLDHTEAL.1 | 6 | 1.27762e-11 | 6.38811e-10 | 3.19405e-10 | 18 | AKLVVPGHGEVGDASLLK | RARYPEARVVVPGHGAPGGPELLDHTEAL | + |
| AKJVVPGHGEVGDASLLK | RARYPEARVVVPGHGAPGGPELLDHTEAL.2 | 6 | 1.27762e-11 | 6.38811e-10 | 3.19405e-10 | 18 | AKLVVPGHGEVGDASLLK | RARYPEARVVVPGHGAPGGPELLDHTEAL | + |
| AKJVVPGHGEVGDASLLK | LVDTGWGPRQTEALLDWARDTL.1 | -2 | 0.00270446 | 0.135223 | 0.0338058 | 16 | AKLVVPGHGEVGDASLLK | LVDTGWGPRQTEALLDWARDTL | + |
| AKJVVPGHGEVGDASLLK | LVDTGWGPRQTEALLDWARDTL.2 | -2 | 0.00270446 | 0.135223 | 0.0338058 | 16 | AKLVVPGHGEVGDASLLK | LVDTGWGPRQTEALLDWARDTL | + |
| AYTLQAQEZEJKVTKJAPGVWVHTSYSTY | LAEDVRVRRJAPGVWLHVTLA.1 | -6 | 8.36433e-14 | 4.18216e-12 | 2.09108e-12 | 21 | AYTLQAQTQELKVTKLAPGVWVHTSYSTY | LAEDVRVRRLAPGVWLHVTLA | + |
| AYTLQAQEZEJKVTKJAPGVWVHTSYSTY | LAEDVRVRRJAPGVWLHVTLA.2 | -6 | 8.36433e-14 | 4.18216e-12 | 2.09108e-12 | 21 | AYTLQAQTQELKVTKLAPGVWVHTSYSTY | LAEDVRVRRLAPGVWLHVTLA | + |
| DQQEAIIFDTPATDQASEELI | LVDTGWGPRQTEALLDWARDTL.1 | -6 | 0.00102561 | 0.0512804 | 0.0256402 | 15 | DQQEAIIFDTPATDQASEELI | LVDTGWGPRQTEALLDWARDTL | + |
| DQQEAIIFDTPATDQASEELI | LVDTGWGPRQTEALLDWARDTL.2 | -6 | 0.00102561 | 0.0512804 | 0.0256402 | 15 | DQQEAIIFDTPATDQASEELI | LVDTGWGPRQTEALLDWARDTL | + |
| EVFYPGAGHTKDNVVVWLPEEKILFGGCFVKS | DAQTLGPLEVFFPGAGHAPDNJVVWHPASGVLFGGCFVKDA.1 | 8 | 5.61362e-31 | 2.80681e-29 | 1.40341e-29 | 32 | EVFYPGAGHTKDNVVVWLPEEKILFGGCFVKS | DAQTLGPLEVFFPGAGHAPDNLVVWHPASGVLFGGCFVKDA | + |
| EVFYPGAGHTKDNVVVWLPEEKILFGGCFVKS | DAQTLGPLEVFFPGAGHAPDNJVVWHPASGVLFGGCFVKDA.2 | 8 | 5.61362e-31 | 2.80681e-29 | 1.40341e-29 | 32 | EVFYPGAGHTKDNVVVWLPEEKILFGGCFVKS | DAQTLGPLEVFFPGAGHAPDNLVVWHPASGVLFGGCFVKDA | + |
| GKDAYJIDTPWTEADTEKLVDWIEQQGLTLKASVSTHSHEDRTGGIGYLN | LVDTGWGPRQTEALLDWARDTL.1 | -5 | 8.67538e-11 | 4.33769e-09 | 2.16885e-09 | 22 | GKDAYIIDTPWTEADTEKLVDWIEQQGLTLKASVSTHSHEDRTGGIGYLN | LVDTGWGPRQTEALLDWARDTL | + |
| GKDAYJIDTPWTEADTEKLVDWIEQQGLTLKASVSTHSHEDRTGGIGYLN | LVDTGWGPRQTEALLDWARDTL.2 | -5 | 8.67538e-11 | 4.33769e-09 | 2.16885e-09 | 22 | GKDAYIIDTPWTEADTEKLVDWIEQQGLTLKASVSTHSHEDRTGGIGYLN | LVDTGWGPRQTEALLDWARDTL | + |
| GKDAYJIDTPWTEADTEKLVDWIEQQGLTLKASVSTHSHEDRTGGIGYLN | HRPVRAAVVTHFHDDRTGGIPALVARGIPVYALED.1 | -26 | 4.71743e-09 | 2.35872e-07 | 5.89679e-08 | 24 | GKDAYIIDTPWTEADTEKLVDWIEQQGLTLKASVSTHSHEDRTGGIGYLN | HRPVRAAVVTHFHDDRTGGIPALVARGIPVHALED | + |
| GKDAYJIDTPWTEADTEKLVDWIEQQGLTLKASVSTHSHEDRTGGIGYLN | HRPVRAAVVTHFHDDRTGGIPALVARGIPVYALED.2 | -26 | 4.71743e-09 | 2.35872e-07 | 5.89679e-08 | 24 | GKDAYIIDTPWTEADTEKLVDWIEQQGLTLKASVSTHSHEDRTGGIGYLN | HRPVRAAVVTHFHDDRTGGIPALVARGIPVHALED | + |
| GVLFPSHGLVVSTKDGAVLVDTGWGNEPTEQLLA | LVDTGWGPRQTEALLDWARDTL.1 | -18 | 5.16288e-10 | 2.58144e-08 | 1.29072e-08 | 16 | GVLFPSHGLVVSTKGGAVLVDTGWGNEPTEQLLA | LVDTGWGPRQTEALLDWARDTL | + |
| GVLFPSHGLVVSTKDGAVLVDTGWGNEPTEQLLA | LVDTGWGPRQTEALLDWARDTL.2 | -18 | 5.16288e-10 | 2.58144e-08 | 1.29072e-08 | 16 | GVLFPSHGLVVSTKGGAVLVDTGWGNEPTEQLLA | LVDTGWGPRQTEALLDWARDTL | + |
| GVLFPSHGLVVSTKDGAVLVDTGWGNEPTEQLLA | YPANGLJVEDGDGSJ.1 | -3 | 6.60974e-08 | 3.30487e-06 | 8.26217e-07 | 15 | GVLFPSHGLVVSTKGGAVLVDTGWGNEPTEQLLA | YPANGLIVEDGDGSL | + |
| GVLFPSHGLVVSTKDGAVLVDTGWGNEPTEQLLA | YPANGLJVEDGDGSJ.2 | -3 | 6.60974e-08 | 3.30487e-06 | 8.26217e-07 | 15 | GVLFPSHGLVVSTKGGAVLVDTGWGNEPTEQLLA | YPANGLIVEDGDGSL | + |
| GYKJKGSISTHFHEDSTGGJEWLNSHSIPTYASELTNELLK | HRPVRAAVVTHFHDDRTGGIPALVARGIPVYALED.1 | 0 | 2.47032e-18 | 1.23516e-16 | 6.1758e-17 | 35 | GYKIKGSISTHFHEDSTGGIEWLNSHSIPTYASELTNELLK | HRPVRAAVVTHFHDDRTGGIPALVARGIPVHALED | + |
| GYKJKGSISTHFHEDSTGGJEWLNSHSIPTYASELTNELLK | HRPVRAAVVTHFHDDRTGGIPALVARGIPVYALED.2 | 0 | 2.47032e-18 | 1.23516e-16 | 6.1758e-17 | 35 | GYKIKGSISTHFHEDSTGGIEWLNSHSIPTYASELTNELLK | HRPVRAAVVTHFHDDRTGGIPALVARGIPVHALED | + |
| HTIKLAKK | TARLAAAQGNPVPSQ.1 | -1 | 0.00112653 | 0.0563264 | 0.0281632 | 7 | HTIKLAKK | TARLAAAQGNPVPSQ | + |
| HTIKLAKK | TARLAAAQGNPVPSQ.2 | -1 | 0.00112653 | 0.0563264 | 0.0281632 | 7 | HTIKLAKK | TARLAAAQGNPVPSQ | + |
| IEVYYPGAGHTKDNLVVWLPKQKJLFGGCLVKSLESKSLGY | DAQTLGPLEVFFPGAGHAPDNJVVWHPASGVLFGGCFVKDA.1 | 7 | 7.48477e-27 | 3.74238e-25 | 1.87119e-25 | 34 | IEVFYPGAGHTKDNLVVWLPKQKILFGGCLVKSLESKSLGY | DAQTLGPLEVFFPGAGHAPDNLVVWHPASGVLFGGCFVKDA | + |
| IEVYYPGAGHTKDNLVVWLPKQKJLFGGCLVKSLESKSLGY | DAQTLGPLEVFFPGAGHAPDNJVVWHPASGVLFGGCFVKDA.2 | 7 | 7.48477e-27 | 3.74238e-25 | 1.87119e-25 | 34 | IEVFYPGAGHTKDNLVVWLPKQKILFGGCLVKSLESKSLGY | DAQTLGPLEVFFPGAGHAPDNLVVWHPASGVLFGGCFVKDA | + |
| IPTYASELTNALLAQNGKPLA | TARLAAAQGNPVPSQ.1 | -8 | 0.000110494 | 0.00552471 | 0.00276235 | 13 | IPTYASELTNALLAQNGKPLA | TARLAAAQGNPVPSQ | + |
| IPTYASELTNALLAQNGKPLA | TARLAAAQGNPVPSQ.2 | -8 | 0.000110494 | 0.00552471 | 0.00276235 | 13 | IPTYASELTNALLAQNGKPLA | TARLAAAQGNPVPSQ | + |
| KGLGNLGDANIEAWPKSAKKVKSK | AKDLGNLADADVAAWPASLER.1 | 1 | 2.2947e-13 | 1.14735e-11 | 5.73675e-12 | 20 | KGLGNLGDANIEAWPKSAKKVKSK | AKDLGNLADADVAAWPASLER | + |
| KGLGNLGDANIEAWPKSAKKVKSK | AKDLGNLADADVAAWPASLER.2 | 1 | 2.2947e-13 | 1.14735e-11 | 5.73675e-12 | 20 | KGLGNLGDANIEAWPKSAKKVKSK | AKDLGNLADADVAAWPASLER | + |
| KHKGLSATHIIEEDKFSLLKD | YPANGLJVEDGDGSJ.1 | -4 | 0.00146031 | 0.0730154 | 0.0365077 | 15 | KHKGLSATHIIEEDKFSLLKD | YPANGLIVEDGDGSL | + |
| KHKGLSATHIIEEDKFSLLKD | YPANGLJVEDGDGSJ.2 | -4 | 0.00146031 | 0.0730154 | 0.0365077 | 15 | KHKGLSATHIIEEDKFSLLKD | YPANGLIVEDGDGSL | + |
| LVPANGLVVLDNKEAYJIDTPWTAKDTEKLVTWIEER | LVDTGWGPRQTEALLDWARDTL.1 | -16 | 3.67797e-09 | 1.83899e-07 | 9.19494e-08 | 21 | LVPANGLVVLDNKEAYLIDTPWTAKDTEKLVTWIVER | LVDTGWGPRQTEALLDWARDTL | + |
| LVPANGLVVLDNKEAYJIDTPWTAKDTEKLVTWIEER | LVDTGWGPRQTEALLDWARDTL.2 | -16 | 3.67797e-09 | 1.83899e-07 | 9.19494e-08 | 21 | LVPANGLVVLDNKEAYLIDTPWTAKDTEKLVTWIVER | LVDTGWGPRQTEALLDWARDTL | + |
| LVPANGLVVLDNKEAYJIDTPWTAKDTEKLVTWIEER | YPANGLJVEDGDGSJ.1 | -1 | 9.6433e-08 | 4.82165e-06 | 1.20541e-06 | 15 | LVPANGLVVLDNKEAYLIDTPWTAKDTEKLVTWIVER | YPANGLIVEDGDGSL | + |
| LVPANGLVVLDNKEAYJIDTPWTAKDTEKLVTWIEER | YPANGLJVEDGDGSJ.2 | -1 | 9.6433e-08 | 4.82165e-06 | 1.20541e-06 | 15 | LVPANGLVVLDNKEAYLIDTPWTAKDTEKLVTWIVER | YPANGLIVEDGDGSL | + |
| MFKITLALLSAVIGF | MSLLRGAVFALLVGVAGCRVSSAAPQPTA.1 | 3 | 0.00103369 | 0.0516844 | 0.0258422 | 15 | MFKITLALLSAVIGF | MSLLRGAVFALLVGVAACSVSSAAAQATA | + |
| MFKITLALLSAVIGF | MSLLRGAVFALLVGVAGCRVSSAAPQPTA.2 | 3 | 0.00103369 | 0.0516844 | 0.0258422 | 15 | MFKITLALLSAVIGF | MSLLRGAVFALLVGVAACSVSSAAAQATA | + |
| MHPPLRPHPGWKRLGPALLALLLGVFGAR | MSLLRGAVFALLVGVAGCRVSSAAPQPTA.1 | -10 | 2.94132e-07 | 1.47066e-05 | 7.35329e-06 | 19 | MHPPLFKHPLFKSLGLALLALLLGAAGAR | MSLLRGAVFALLVGVAACSVSSAAAQATA | + |
| MHPPLRPHPGWKRLGPALLALLLGVFGAR | MSLLRGAVFALLVGVAGCRVSSAAPQPTA.2 | -10 | 2.94132e-07 | 1.47066e-05 | 7.35329e-06 | 19 | MHPPLFKHPLFKSLGLALLALLLGAAGAR | MSLLRGAVFALLVGVAACSVSSAAAQATA | + |
| MKFLAPFLFLLPCVTVATEAS | CPLLLLLTACASTPS.1 | -4 | 0.000331912 | 0.0165956 | 0.00829779 | 15 | MKFLAPFLFLLPCVTVATEAS | CPLLLLLTACASTPS | + |
| MKFLAPFLFLLPCVTVATEAS | CPLLLLLTACASTPS.2 | -4 | 0.000331912 | 0.0165956 | 0.00829779 | 15 | MKFLAPFLFLLPCVTVATEAS | CPLLLLLTACASTPS | + |
| MKIILKNILVILFSFVILSCSSQNKDHFK | MSLLRGAVFALLVGVAGCRVSSAAPQPTA.1 | 0 | 0.00123438 | 0.0617189 | 0.0308595 | 29 | MKIILKNILVILFSFVILSCSSQNKDHFK | MSLLRGAVFALLVGVAACSVSSAAAQATA | + |
| MKIILKNILVILFSFVILSCSSQNKDHFK | MSLLRGAVFALLVGVAGCRVSSAAPQPTA.2 | 0 | 0.00123438 | 0.0617189 | 0.0308595 | 29 | MKIILKNILVILFSFVILSCSSQNKDHFK | MSLLRGAVFALLVGVAACSVSSAAAQATA | + |
| MKKLJLLLLLLLILL | MSLLRGAVFALLVGVAGCRVSSAAPQPTA.1 | 0 | 0.00161896 | 0.0809479 | 0.0293505 | 15 | MKKLLLLLLLLLILL | MSLLRGAVFALLVGVAACSVSSAAAQATA | + |
| MKKLJLLLLLLLILL | MSLLRGAVFALLVGVAGCRVSSAAPQPTA.2 | 0 | 0.00161896 | 0.0809479 | 0.0293505 | 15 | MKKLLLLLLLLLILL | MSLLRGAVFALLVGVAACSVSSAAAQATA | + |
| MKKLJLLLLLLLILL | MKILILAAMLFAAQP.1 | 0 | 0.00234804 | 0.117402 | 0.0293505 | 15 | MKKLLLLLLLLLILL | MKILILAAMLFAAQP | + |
| MKKLJLLLLLLLILL | MKILILAAMLFAAQP.2 | 0 | 0.00234804 | 0.117402 | 0.0293505 | 15 | MKKLLLLLLLLLILL | MKILILAAMLFAAQP | + |
| MKKLJLLLLLLLILL | CPLLLLLTACASTPS.1 | -2 | 0.00517568 | 0.258784 | 0.0431307 | 13 | MKKLLLLLLLLLILL | CPLLLLLTACASTPS | + |
| MKKLJLLLLLLLILL | CPLLLLLTACASTPS.2 | -2 | 0.00517568 | 0.258784 | 0.0431307 | 13 | MKKLLLLLLLLLILL | CPLLLLLTACASTPS | + |
| MKYJJLJJLLL | MKILILAAMLFAAQP.1 | 0 | 0.000401681 | 0.0200841 | 0.010042 | 11 | MKYILLLLLLL | MKILILAAMLFAAQP | + |
| MKYJJLJJLLL | MKILILAAMLFAAQP.2 | 0 | 0.000401681 | 0.0200841 | 0.010042 | 11 | MKYILLLLLLL | MKILILAAMLFAAQP | + |
| MLKKFLLLIQVFCVVTISAQS | CPLLLLLTACASTPS.1 | -3 | 0.00105171 | 0.0525856 | 0.0262928 | 15 | MLKKFLLLIQVFCVVTISAQS | CPLLLLLTACASTPS | + |
| MLKKFLLLIQVFCVVTISAQS | CPLLLLLTACASTPS.2 | -3 | 0.00105171 | 0.0525856 | 0.0262928 | 15 | MLKKFLLLIQVFCVVTISAQS | CPLLLLLTACASTPS | + |
| MLKKFLLLIQVFCVVTISAQS | LAFLACAATASAAPP.1 | -7 | 0.00309012 | 0.154506 | 0.0323878 | 14 | MLKKFLLLIQVFCVVTISAQS | LAFLACAATASAAPP | + |
| MLKKFLLLIQVFCVVTISAQS | LAFLACAATASAAPP.2 | -7 | 0.00309012 | 0.154506 | 0.0323878 | 14 | MLKKFLLLIQVFCVVTISAQS | LAFLACAATASAAPP | + |
| MLKKFLLLIQVFCVVTISAQS | MSLLRGAVFALLVGVAGCRVSSAAPQPTA.1 | 4 | 0.00388654 | 0.194327 | 0.0323878 | 21 | MLKKFLLLIQVFCVVTISAQS | MSLLRGAVFALLVGVAACSVSSAAAQATA | + |
| MLKKFLLLIQVFCVVTISAQS | MSLLRGAVFALLVGVAGCRVSSAAPQPTA.2 | 4 | 0.00388654 | 0.194327 | 0.0323878 | 21 | MLKKFLLLIQVFCVVTISAQS | MSLLRGAVFALLVGVAACSVSSAAAQATA | + |
| MLKKFLLLIQVFCVVTISAQS | TPSMQAEYT.1 | -15 | 0.00764868 | 0.382434 | 0.0478043 | 6 | MLKKFLLLIQVFCVVTISAQS | TPSMQAEYT | + |
| MLKKFLLLIQVFCVVTISAQS | TPSMQAEYT.2 | -15 | 0.00764868 | 0.382434 | 0.0478043 | 6 | MLKKFLLLIQVFCVVTISAQS | TPSMQAEYT | + |
| PJADGVYLHTSYKZVEGFGLVDSNGLVVV | YPANGLJVEDGDGSJ.1 | -20 | 0.000180961 | 0.00904804 | 0.00303423 | 9 | PLADGVYLHTSYKQVEGFGLVDSNGLVVV | YPANGLIVEDGDGSL | + |
| PJADGVYLHTSYKZVEGFGLVDSNGLVVV | YPANGLJVEDGDGSJ.2 | -20 | 0.000180961 | 0.00904804 | 0.00303423 | 9 | PLADGVYLHTSYKQVEGFGLVDSNGLVVV | YPANGLIVEDGDGSL | + |
| PJADGVYLHTSYKZVEGFGLVDSNGLVVV | LAEDVRVRRJAPGVWLHVTLA.1 | 8 | 0.000242739 | 0.0121369 | 0.00303423 | 13 | PLADGVYLHTSYKQVEGFGLVDSNGLVVV | LAEDVRVRRLAPGVWLHVTLA | + |
| PJADGVYLHTSYKZVEGFGLVDSNGLVVV | LAEDVRVRRJAPGVWLHVTLA.2 | 8 | 0.000242739 | 0.0121369 | 0.00303423 | 13 | PLADGVYLHTSYKQVEGFGLVDSNGLVVV | LAEDVRVRRLAPGVWLHVTLA | + |
| VGDASLLDHTIELAE | RARYPEARVVVPGHGAPGGPELLDHTEAL.1 | 16 | 1.54892e-08 | 7.7446e-07 | 3.8723e-07 | 13 | VGDASLLDHTIELAE | RARYPEARVVVPGHGAPGGPELLDHTEAL | + |
| VGDASLLDHTIELAE | RARYPEARVVVPGHGAPGGPELLDHTEAL.2 | 16 | 1.54892e-08 | 7.7446e-07 | 3.8723e-07 | 13 | VGDASLLDHTIELAE | RARYPEARVVVPGHGAPGGPELLDHTEAL | + |
| WVETNLKQPVKAVVATHFHEDCLGGLQAF | HRPVRAAVVTHFHDDRTGGIPALVARGIPVYALED.1 | -6 | 5.07018e-13 | 2.53509e-11 | 1.26754e-11 | 23 | WVETNLKQPVKAVVATHFHDDCLGGLGAF | HRPVRAAVVTHFHDDRTGGIPALVARGIPVHALED | + |
| WVETNLKQPVKAVVATHFHEDCLGGLQAF | HRPVRAAVVTHFHDDRTGGIPALVARGIPVYALED.2 | -6 | 5.07018e-13 | 2.53509e-11 | 1.26754e-11 | 23 | WVETNLKQPVKAVVATHFHDDCLGGLGAF | HRPVRAAVVTHFHDDRTGGIPALVARGIPVHALED | + |
| AKSKNLPIPDHGFKD | AKLAKEQGYEVPBPTLDELTT | 3 | 0.000968776 | 0.0484388 | 0.0484388 | 15 | AKSKNLPIPAHGFKD | AKLAKEQGYEVPNPSLDELTT | + |
| DASLLKHTLELAVKG | VVPGHGEWGGKELLSHTLELL | 9 | 7.33954e-07 | 3.66977e-05 | 3.66977e-05 | 12 | DASLLKHTLELAVKG | VVPGHGEWGGKELLSHTLELL | + |
| DASLLKHTLELAVKG | VVPGHGLPGGPELLDHTEALL | 9 | 0.000436768 | 0.0218384 | 0.0109192 | 12 | DASLLKHTLELAVKG | VVPGHGLPGGPELLDHTEALL | + |
| DASLLKHTLELAVKG | MKFHKCWIKKDVSKLFLTVEVVLTSFSED | 10 | 0.000782833 | 0.0391417 | 0.0130472 | 15 | DASLLKHTLELAVKG | MKFHKCWIKKDVSKLFLTVEVVLTSGSED | + |
| DSLVLKVGNEK | LKFGNTKV | -4 | 0.000207727 | 0.0103864 | 0.0103864 | 7 | DSLVLKVGNEK | LKFGNTKV | + |
| EGVYVHTSYEEVEGWGLVPANGLVVLDNKEAYJIDTPWTAK | YGLVDANGLVVLDGQGAYIIDTPWSZQDT | -14 | 8.61419e-23 | 4.30709e-21 | 4.30709e-21 | 27 | EGVYVHTSYEEVEGWGLVPANGLVVLDNKEAYLIDTPWTAK | YGLVDANGLVVLDGQGAYIIDTPWSEQDT | + |
| EGVYVHTSYEEVEGWGLVPANGLVVLDNKEAYJIDTPWTAK | FPSNGLIVETGDGLVLIDTAWGEEQTEZL | -17 | 2.62703e-10 | 1.31352e-08 | 6.56759e-09 | 24 | EGVYVHTSYEEVEGWGLVPANGLVVLDNKEAYLIDTPWTAK | FPSNGLIVETGDGLVLIDTAWGELQTEEL | + |
| EGVYVHTSYEEVEGWGLVPANGLVVLDNKEAYJIDTPWTAK | ANGLJVEDGDESLLVDTAWGARQTAALLAW | -19 | 7.50667e-09 | 3.75333e-07 | 1.25111e-07 | 22 | EGVYVHTSYEEVEGWGLVPANGLVVLDNKEAYLIDTPWTAK | ANGLIVEDGDESLLVDTAWGARQTAALLAW | + |
| EGVYVHTSYEEVEGWGLVPANGLVVLDNKEAYJIDTPWTAK | VEITKJAPGVWVHTSYYTYPG | 7 | 5.28018e-06 | 0.000264009 | 6.60023e-05 | 14 | EGVYVHTSYEEVEGWGLVPANGLVVLDNKEAYLIDTPWTAK | VEITKIAPGVWVHTSYYTYPG | + |
| ELLDYTIDLF | VVPGHGLPGGPELLDHTEALL | 11 | 1.41536e-05 | 0.000707678 | 0.000381655 | 10 | ELLDYTIDLF | VVPGHGLPGGPELLDHTEALL | + |
| ELLDYTIDLF | VVPGHGEWGGKELLSHTLELL | 11 | 1.52662e-05 | 0.00076331 | 0.000381655 | 10 | ELLDYTIDLF | VVPGHGEWGGKELLSHTLELL | + |
| ELLKENGKPLAKHTF | AKLAKEQGYEVPBPTLDELTT | 1 | 1.40501e-09 | 7.02507e-08 | 7.02507e-08 | 15 | ELLKENGKPLAKHTF | AKLAKEQGYEVPNPSLDELTT | + |
| ELLKENGKPLAKHTF | YASPSTRRLAEAEGNEIPTHSLEGLSSSG | 7 | 3.33356e-06 | 0.000166678 | 8.33389e-05 | 15 | ELLKENGKPLAKHTF | YASPSTRRLAEAEGNEIPTHSLEGLSSSG | + |
| ELLKENGKPLAKHTF | HGLEDTARLATEQGNPVPTQR | 7 | 2.43773e-05 | 0.00121886 | 0.000406288 | 14 | ELLKENGKPLAKHTF | HGLEDTARLATEQGNPVPTQR | + |
| ELLKENGKPLAKHTF | YALPLSNELAPERGLPPAEFL | 7 | 3.72493e-05 | 0.00186247 | 0.000465616 | 14 | ELLKENGKPLAKHTF | YALPLSNQLAASNGLPPAEFL | + |
| ELLKENGKPLAKHTF | IVTHAHDDRIGGIDVLKKRGIPVYSTPLT | 13 | 0.00030282 | 0.015141 | 0.0030282 | 15 | ELLKENGKPLAKHTF | IVTHAHDDRIGGIDVLKKRGIPVYSTPLT | + |
| ELLKKDGKVQATNSF | AKLAKEQGYEVPBPTLDELTT | 1 | 2.95335e-05 | 0.00147668 | 0.00147668 | 15 | ELLKKDGKVQATNSF | AKLAKEQGYEVPNPSLDELTT | + |
| ELLKKDGKVQATNSF | YASPSTRRLAEAEGNEIPTHSLEGLSSSG | 7 | 0.000469468 | 0.0234734 | 0.0117367 | 15 | ELLKKDGKVQATNSF | YASPSTRRLAEAEGNEIPTHSLEGLSSSG | + |
| ENGKGTPDITFATDT | AKLAKEQGYEVPBPTLDELTT | 5 | 5.98261e-06 | 0.00029913 | 0.00029913 | 15 | ENKKGTPDVTFATDT | AKLAKEQGYEVPNPSLDELTT | + |
| EYESPELEITPJSDNVYLHTSYKZVEGFG | VEITKJAPGVWVHTSYYTYPG | -6 | 2.03479e-17 | 1.01739e-15 | 1.01739e-15 | 21 | EYESPELEITPLSDNVYLHTSYKQVEGFG | VEITKIAPGVWVHTSYYTYPG | + |
| EYESPELEITPJSDNVYLHTSYKZVEGFG | DVRVRRJAPGVWLHVTTAGFD | -5 | 4.45498e-08 | 2.22749e-06 | 1.11374e-06 | 21 | EYESPELEITPLSDNVYLHTSYKQVEGFG | DVRVRRIAPGVWLHVTTAGFD | + |
| GLGNLGDANIEAWPKSIKKVKSKYPKAKLVVPGHGKVG | KSLGNTADADLKEWPKSIKRVQQRYPKAK | 1 | 3.80907e-26 | 1.90453e-24 | 1.90453e-24 | 28 | GLGNLGDANIEAWPKSIKKVKSKYPKAKLVVPGHGKVG | KSLGNTADADLKEWPKSIKRVQQRYPKAK | + |
| GLGNLGDANIEAWPKSIKKVKSKYPKAKLVVPGHGKVG | VLFGGCFVKDASAKSLGNVADADVAAWPASJERIRQRYPEARV | 14 | 3.08935e-13 | 1.54468e-11 | 7.72338e-12 | 29 | GLGNLGDANIEAWPKSIKKVKSKYPKAKLVVPGHGKVG | VLFGGCFVKDASAKSLGNVADADVAAWPASLERIRQRYPEARV | + |
| GLGNLGDANIEAWPKSIKKVKSKYPKAKLVVPGHGKVG | VVPGHGEWGGKELLSHTLELL | -29 | 0.000131073 | 0.00655365 | 0.00218455 | 9 | GLGNLGDANIEAWPKSIKKVKSKYPKAKLVVPGHGKVG | VVPGHGEWGGKELLSHTLELL | + |
| GLGNLGDANIEAWPKSIKKVKSKYPKAKLVVPGHGKVG | VVPGHGLPGGPELLDHTEALL | -29 | 0.00394491 | 0.197246 | 0.0493114 | 9 | GLGNLGDANIEAWPKSIKKVKSKYPKAKLVVPGHGKVG | VVPGHGLPGGPELLDHTEALL | + |
| IEVFYPGAGHTKDNLVVWLPKEKILFGGCLVKS | EVFYPGPGHSPDNJVVWLPZYKILFGGCLVKSLQA | -1 | 1.89722e-39 | 9.48612e-38 | 9.48612e-38 | 32 | IEVFYPGAGHTKDNLVVWLPKEKILFGGCLVKS | EVFYPGPGHSPDNIVVWLPQYKILFGGCLVKSLQA | + |
| IEVFYPGAGHTKDNLVVWLPKEKILFGGCLVKS | GPLELFFPGAGHSPDNLVVWH | 2 | 2.4838e-17 | 1.2419e-15 | 6.20949e-16 | 19 | IEVFYPGAGHTKDNLVVWLPKEKILFGGCLVKS | GPLELFFPGAGHSPDNLVVWH | + |
| ISTHFHEDRTGGJGYLNSHGIPTYASELT | IVTHAHDDRIGGIDVLKKRGIPVYSTPLT | 0 | 8.78285e-24 | 4.39142e-22 | 4.39142e-22 | 29 | ISTHFHEDRTGGLGYLNSHGIPTYASELT | IVTHAHDDRIGGIDVLKKRGIPVYSTPLT | + |
| ISTHFHEDRTGGJGYLNSHGIPTYASELT | ARDTJGLPVRAAVVTHFHDDRTGGVPVLRARGIPV | 12 | 1.39162e-15 | 6.95809e-14 | 3.47904e-14 | 23 | ISTHFHEDRTGGLGYLNSHGIPTYASELT | ARDTLGLPVRAAVVTHFHDDRTGGVPVLRARGIPV | + |
| IVQTAPHKAVLIDTPWDNSDVDTLFSWLE | YGLVDANGLVVLDGQGAYIIDTPWSZQDT | 8 | 1.08021e-05 | 0.000540107 | 0.000540107 | 21 | IVQTAPHKAVLIDTPWDNSDVDTLFSWLE | YGLVDANGLVVLDGQGAYIIDTPWSEQDT | + |
| IVQTAPHKAVLIDTPWDNSDVDTLFSWLE | FPSNGLIVETGDGLVLIDTAWGEEQTEZL | 5 | 9.49087e-05 | 0.00474543 | 0.00237272 | 24 | IVQTAPHKAVLIDTPWDNSDVDTLFSWLE | FPSNGLIVETGDGLVLIDTAWGELQTEEL | + |
| IVQTAPHKAVLIDTPWDNSDVDTLFSWLE | ANGLJVEDGDESLLVDTAWGARQTAALLAW | 3 | 0.000256053 | 0.0128026 | 0.00426754 | 27 | IVQTAPHKAVLIDTPWDNSDVDTLFSWLE | ANGLIVEDGDESLLVDTAWGARQTAALLAW | + |
| LGNLGDAVLEAWPTSAKKLISKYGEAKJV | KSLGNTADADLKEWPKSIKRVQQRYPKAK | 2 | 5.0056e-16 | 2.5028e-14 | 2.5028e-14 | 27 | LGNLGDAVLEAWPTSAKKLISKYGKAKIV | KSLGNTADADLKEWPKSIKRVQQRYPKAK | + |
| LGNLGDAVLEAWPTSAKKLISKYGEAKJV | VLFGGCFVKDASAKSLGNVADADVAAWPASJERIRQRYPEARV | 15 | 2.94649e-13 | 1.47324e-11 | 7.36622e-12 | 28 | LGNLGDAVLEAWPTSAKKLISKYGKAKIV | VLFGGCFVKDASAKSLGNVADADVAAWPASLERIRQRYPEARV | + |
| LVPSNGLVVVDGKEAYJIDTPWSDKDTEKLVDWI | YGLVDANGLVVLDGQGAYIIDTPWSZQDT | 2 | 8.1861e-27 | 4.09305e-25 | 4.09305e-25 | 27 | LVPSNGLVVVDGKEAYIIDTPWSDKDTEKLVDWI | YGLVDANGLVVLDGQGAYIIDTPWSEQDT | + |
| LVPSNGLVVVDGKEAYJIDTPWSDKDTEKLVDWI | ANGLJVEDGDESLLVDTAWGARQTAALLAW | -3 | 1.33702e-19 | 6.68509e-18 | 3.34255e-18 | 30 | LVPSNGLVVVDGKEAYIIDTPWSDKDTEKLVDWI | ANGLIVEDGDESLLVDTAWGARQTAALLAW | + |
| LVPSNGLVVVDGKEAYJIDTPWSDKDTEKLVDWI | FPSNGLIVETGDGLVLIDTAWGEEQTEZL | -1 | 2.72911e-19 | 1.36455e-17 | 4.54851e-18 | 29 | LVPSNGLVVVDGKEAYIIDTPWSDKDTEKLVDWI | FPSNGLIVETGDGLVLIDTAWGELQTEEL | + |
| MFKJTLALLSAVIGF | MFKLJSKLLVY | 0 | 0.00133869 | 0.0669345 | 0.0374603 | 11 | MFKITLALLSAVIGF | MFKLLSKLLVY | + |
| MFKJTLALLSAVIGF | MKRJJLLLLJL | 0 | 0.00149841 | 0.0749206 | 0.0374603 | 11 | MFKITLALLSAVIGF | MKRILLLLLLL | + |
| MKIRIKNIVAJLLSFVILSCSSZKTDNFK | MMKFRRIGGFLFFLLIGSFTQSAQENIPN | 1 | 0.00123318 | 0.0616589 | 0.0411282 | 28 | MKIRIKNIVALLLSFVILSCSSQKTTNFK | MMKFRRIGGFLFFLLIGSFTQSAQANIPN | + |
| MKIRIKNIVAJLLSFVILSCSSZKTDNFK | MKRJJLLLLJL | -4 | 0.00164513 | 0.0822564 | 0.0411282 | 11 | MKIRIKNIVALLLSFVILSCSSQKTTNFK | MKRILLLLLLL | + |
| MKKLJLLLLJL | MKRJJLLLLJL | 0 | 1.16466e-12 | 5.82328e-11 | 5.82328e-11 | 11 | MKKLILLLLIL | MKRILLLLLLL | + |
| MKKLJLLLLJL | MKFLFAALFVVMFCGLARASD | 0 | 0.000105567 | 0.00527833 | 0.00263916 | 11 | MKKLILLLLIL | MKSLFAALFVVMFCGLARASD | + |
| MKKLJLLLLJL | MFKLJSKLLVY | 0 | 0.000230745 | 0.0115373 | 0.00384575 | 11 | MKKLILLLLIL | MFKLLSKLLVY | + |
| MKKLLVFFJFLFCSI | MKRJJLLLLJL | 0 | 5.66371e-10 | 2.83185e-08 | 2.83185e-08 | 11 | MKKLLVFFIFLFCSI | MKRILLLLLLL | + |
| MKKLLVFFJFLFCSI | KLCRTHPHGVLMFKLLSKLLVYLTASIMAIASPLAFSVDSSGEYPTVSEI | 12 | 0.00132872 | 0.0664361 | 0.033218 | 15 | MKKLLVFFIFLFCSI | KLCRTHPHGVLMFKLLSKLLVYLTASIMAIASPLAFSVDSSGEYPTVSEI | + |
| MKYIJLLLLLI | MKRJJLLLLJL | 0 | 1.4027e-12 | 7.01349e-11 | 7.01349e-11 | 11 | MKYIILLLLLI | MKRILLLLLLL | + |
| MKYIJLLLLLI | MKFLFAALFVVMFCGLARASD | 0 | 0.00113371 | 0.0566855 | 0.0283427 | 11 | MKYIILLLLLI | MKSLFAALFVVMFCGLARASD | + |
| MKYIJLLLLLI | MMKFRRIGGFLFFLLIGSFTQSAQENIPN | 1 | 0.0019826 | 0.0991301 | 0.0330434 | 11 | MKYIILLLLLI | MMKFRRIGGFLFFLLIGSFTQSAQANIPN | + |
| MPNRALWQRIAILLJ | MKRJJLLLLJL | -6 | 0.000977803 | 0.0488902 | 0.0488902 | 9 | MPNRALWQRIAILLL | MKRILLLLLLL | + |
| MSIQHFRVALIPFFAAFCLP | MSIQHFRVALI | 0 | 1.79381e-15 | 8.96903e-14 | 8.96903e-14 | 11 | MSIQHFRVALIPFFAAFCLP | MSIQHFRVALI | + |
| PAYTLLAQEQZJKVTKIAPBVYVHTSYSTYQGVLVPSHGLV | VEITKJAPGVWVHTSYYTYPG | -11 | 2.45156e-18 | 1.22578e-16 | 1.22578e-16 | 21 | PAYTLLAQEQELKVTKIAPNVYVHTSYSTYQGVLVPSHGLV | VEITKIAPGVWVHTSYYTYPG | + |
| PAYTLLAQEQZJKVTKIAPBVYVHTSYSTYQGVLVPSHGLV | LLLVPAYALLAQSQZ | 4 | 4.90897e-09 | 2.45449e-07 | 1.22724e-07 | 11 | PAYTLLAQEQELKVTKIAPNVYVHTSYSTYQGVLVPSHGLV | LLLVPAYALLAQSQE | + |
| PAYTLLAQEQZJKVTKIAPBVYVHTSYSTYQGVLVPSHGLV | DVRVRRJAPGVWLHVTTAGFD | -10 | 1.46811e-07 | 7.34054e-06 | 2.44685e-06 | 21 | PAYTLLAQEQELKVTKIAPNVYVHTSYSTYQGVLVPSHGLV | DVRVRRIAPGVWLHVTTAGFD | + |
| PGHGKVGDGELLDHT | VVPGHGEWGGKELLSHTLELL | 2 | 2.11517e-16 | 1.05758e-14 | 1.05758e-14 | 15 | PGHGKVGDGELLDHT | VVPGHGEWGGKELLSHTLELL | + |
| PGHGKVGDGELLDHT | VVPGHGLPGGPELLDHTEALL | 2 | 7.53772e-14 | 3.76886e-12 | 1.88443e-12 | 15 | PGHGKVGDGELLDHT | VVPGHGLPGGPELLDHTEALL | + |
| PGHGKVGDGELLDHT | ANGLJVEDGDESLLVDTAWGARQTAALLAW | 2 | 0.00210648 | 0.105324 | 0.035108 | 15 | PGHGKVGDGELLDHT | ANGLIVEDGDESLLVDTAWGARQTAALLAW | + |
| PLKIEAJSSKVYLVKSYKEILNVYESDTPKIIDANALJYI | VEITKJAPGVWVHTSYYTYPG | -1 | 9.36304e-07 | 4.68152e-05 | 4.68152e-05 | 21 | PLKIEALSSKVYLVKSYKTILNVYESDTPKIIDANSLLYI | VEITKIAPGVWVHTSYYTYPG | + |
| SKSLGYTGDABLSAWPNSVEKVKAKYPDA | KSLGNTADADLKEWPKSIKRVQQRYPKAK | -1 | 6.11003e-29 | 3.05501e-27 | 3.05501e-27 | 28 | SKSLGYTGDADLSAWPNSVEKVKAKYPDA | KSLGNTADADLKEWPKSIKRVQQRYPKAK | + |
| SKSLGYTGDABLSAWPNSVEKVKAKYPDA | VLFGGCFVKDASAKSLGNVADADVAAWPASJERIRQRYPEARV | 12 | 6.59462e-24 | 3.29731e-22 | 1.64866e-22 | 29 | SKSLGYTGDADLSAWPNSVEKVKAKYPDA | VLFGGCFVKDASAKSLGNVADADVAAWPASLERIRQRYPEARV | + |
| STHFHEDSTGGIEWLNSRSIPTYASELTN | IVTHAHDDRIGGIDVLKKRGIPVYSTPLT | 1 | 9.27576e-20 | 4.63788e-18 | 4.63788e-18 | 28 | STHFHEDSTGGIEWLNSRSIPTYASELTN | IVTHAHDDRIGGIDVLKKRGIPVYSTPLT | + |
| STHFHEDSTGGIEWLNSRSIPTYASELTN | ARDTJGLPVRAAVVTHFHDDRTGGVPVLRARGIPV | 13 | 5.48616e-12 | 2.74308e-10 | 1.37154e-10 | 22 | STHFHEDSTGGIEWLNSRSIPTYASELTN | ARDTLGLPVRAAVVTHFHDDRTGGVPVLRARGIPV | + |
| TKEGAVLIDTGWGNE | FPSNGLIVETGDGLVLIDTAWGEEQTEZL | 9 | 9.59638e-13 | 4.79819e-11 | 4.79819e-11 | 15 | TKEGAVLIDTGWGNE | FPSNGLIVETGDGLVLIDTAWGELQTEEL | + |
| TKEGAVLIDTGWGNE | ANGLJVEDGDESLLVDTAWGARQTAALLAW | 7 | 6.33447e-08 | 3.16723e-06 | 1.58362e-06 | 15 | TKEGAVLIDTGWGNE | ANGLIVEDGDESLLVDTAWGARQTAALLAW | + |
| TKEGAVLIDTGWGNE | YGLVDANGLVVLDGQGAYIIDTPWSZQDT | 12 | 2.38356e-06 | 0.000119178 | 3.97259e-05 | 15 | TKEGAVLIDTGWGNE | YGLVDANGLVVLDGQGAYIIDTPWSEQDT | + |
| TSVPAPRPAAEPFTSITEDLRIQPLAPGVWRLVALSGEEWG | DVRVRRJAPGVWLHVTTAGFD | -18 | 8.90756e-09 | 4.45378e-07 | 4.45378e-07 | 21 | TSVPAPRPAAEPFTSITEDLRIQPLAPGVWRLVALSGEEWG | DVRVRRIAPGVWLHVTTAGFD | + |
| YWLVKNKIEVFYPGAGHTPDNVVVWLPEEKILFGGCFVKSL | EVFYPGPGHSPDNJVVWLPZYKILFGGCLVKSLQA | -8 | 1.79573e-38 | 8.97865e-37 | 8.97865e-37 | 33 | FWLVKGKIEVFYPGAGHTPDNVVVWLPESKILFGGCFVKSL | EVFYPGPGHSPDNIVVWLPQYKILFGGCLVKSLQA | + |
| YWLVKNKIEVFYPGAGHTPDNVVVWLPEEKILFGGCFVKSL | GPLELFFPGAGHSPDNLVVWH | -5 | 5.51457e-18 | 2.75728e-16 | 1.37864e-16 | 21 | FWLVKGKIEVFYPGAGHTPDNVVVWLPESKILFGGCFVKSL | GPLELFFPGAGHSPDNLVVWH | + |
| AKJVVPGHGEVGDASLL | KIVVPGHGEWGGKDLLSHTJKLLK | -1 | 5.82088e-15 | 2.91044e-13 | 2.91044e-13 | 16 | AKLVVPGHGEVGDASLL | KIVVPGHGEWGGKDLLSHTLKLLK | + |
| AKJVVPGHGEVGDASLL | VVPGHGLPGGPELLDHTEALL | -3 | 1.09057e-08 | 5.45287e-07 | 2.72643e-07 | 14 | AKLVVPGHGEVGDASLL | VVPGHGLPGGPELLDHTEALL | + |
| ASNKSIQPTAEASAD | DLAPSNKLLPAEFDLSFDNNNKSSDISP | 3 | 0.00048757 | 0.0243785 | 0.0243785 | 15 | ASNKSIQPTAEASAD | DLAPSNKLLPAEFDLSFDSNNKSSDISP | + |
| AYJIDTPWTEKDTEKLVDWIEAQGLTLKASISTHSHZDRTGGIGYLNSKG | VTHAHDDRIGGIDVLKKRGIPVYSTPLTA | -31 | 2.04068e-09 | 1.02034e-07 | 1.02034e-07 | 19 | AYIIDTPWTEKDTEKLVDWIEAQGLTLKASISTHSHEDRTGGIGYLNSKG | VTHAHDDRIGGIDVLKKRGIPVYSTPLTA | + |
| AYJIDTPWTEKDTEKLVDWIEAQGLTLKASISTHSHZDRTGGIGYLNSKG | HFHDDRTGGVPVLRARGIPVYALPDTARL | -33 | 3.28403e-06 | 0.000164201 | 8.21007e-05 | 17 | AYIIDTPWTEKDTEKLVDWIEAQGLTLKASISTHSHEDRTGGIGYLNSKG | HFHDDRTGGVPVLRARGIPVYALPDTARL | + |
| AYJIDTPWTEKDTEKLVDWIEAQGLTLKASISTHSHZDRTGGIGYLNSKG | FPSNGLIVETGKGLVLIDTAWGEEQTEZL | 13 | 4.30389e-05 | 0.00215195 | 0.000717315 | 16 | AYIIDTPWTEKDTEKLVDWIEAQGLTLKASISTHSHEDRTGGIGYLNSKG | FPSNGLIVETGKGLVLIDTAWGEEQTEEL | + |
| AYJIDTPWTEKDTEKLVDWIEAQGLTLKASISTHSHZDRTGGIGYLNSKG | YPANGLIVEDGDELLLVDTAWGARQTAALL | 13 | 0.000153177 | 0.00765887 | 0.00191472 | 17 | AYIIDTPWTEKDTEKLVDWIEAQGLTLKASISTHSHEDRTGGIGYLNSKG | YPANGLIVEDGDELLLVDTAWGARQTAALL | + |
| AYJIDTPWTEKDTEKLVDWIEAQGLTLKASISTHSHZDRTGGIGYLNSKG | AEDTJGLPVRAAVVT | -18 | 0.00165777 | 0.0828886 | 0.0165777 | 15 | AYIIDTPWTEKDTEKLVDWIEAQGLTLKASISTHSHEDRTGGIGYLNSKG | AEDTLGLPVRAAVVT | + |
| DHTIKLFK | KIVVPGHGEWGGKDLLSHTJKLLK | 16 | 2.79658e-07 | 1.39829e-05 | 1.39829e-05 | 8 | DHTIKLFK | KIVVPGHGEWGGKDLLSHTLKLLK | + |
| DHTIKLFK | VVPGHGLPGGPELLDHTEALL | 14 | 0.000403185 | 0.0201593 | 0.0100796 | 7 | DHTIKLFK | VVPGHGLPGGPELLDHTEALL | + |
| FDTPWTBEPTEQLLAWVKDNLKAPVKAFVPTHWHDDCLGGL | EWIKTKLKKPVKKAI | -14 | 5.88801e-10 | 2.94401e-08 | 2.94401e-08 | 15 | FDTPWTDEQTEQLLAWVKDNLKAPVKAFVPTHWHDDCLGGL | EWIKTKLKKPVKKAI | + |
| FDTPWTBEPTEQLLAWVKDNLKAPVKAFVPTHWHDDCLGGL | AEDTJGLPVRAAVVT | -16 | 2.63472e-07 | 1.31736e-05 | 6.5868e-06 | 15 | FDTPWTDEQTEQLLAWVKDNLKAPVKAFVPTHWHDDCLGGL | AEDTLGLPVRAAVVT | + |
| FDTPWTBEPTEQLLAWVKDNLKAPVKAFVPTHWHDDCLGGL | FPSNGLIVETGKGLVLIDTAWGEEQTEZL | 16 | 5.22527e-06 | 0.000261263 | 8.70878e-05 | 13 | FDTPWTDEQTEQLLAWVKDNLKAPVKAFVPTHWHDDCLGGL | FPSNGLIVETGKGLVLIDTAWGEEQTEEL | + |
| FDTPWTBEPTEQLLAWVKDNLKAPVKAFVPTHWHDDCLGGL | YPANGLIVEDGDELLLVDTAWGARQTAALL | 16 | 3.05257e-05 | 0.00152628 | 0.000381571 | 14 | FDTPWTDEQTEQLLAWVKDNLKAPVKAFVPTHWHDDCLGGL | YPANGLIVEDGDELLLVDTAWGARQTAALL | + |
| FDTPWTBEPTEQLLAWVKDNLKAPVKAFVPTHWHDDCLGGL | VTHAHDDRIGGIDVLKKRGIPVYSTPLTA | -29 | 0.00103112 | 0.0515559 | 0.0103112 | 12 | FDTPWTDEQTEQLLAWVKDNLKAPVKAFVPTHWHDDCLGGL | VTHAHDDRIGGIDVLKKRGIPVYSTPLTA | + |
| GDABVSAWPNSVEKVKKK | VWHPSSGVLFGGCAVKDASAKSLGNVADADLAAWPASJERIRQRYPEARV | 26 | 5.68197e-10 | 2.84098e-08 | 2.84098e-08 | 18 | GDANVSAWPNSVEKVKKK | VWHPSSGVLFGGCAVKDASAKSLGNVADADLAAWPASLERIRQRYPEARV | + |
| GDABVSAWPNSVEKVKKK | SIKRVQQRYPK | -10 | 5.62658e-06 | 0.000281329 | 0.000140664 | 8 | GDANVSAWPNSVEKVKKK | SIKRVQQRYPK | + |
| GDABVSAWPNSVEKVKKK | EVFYPGPGHTMDNIVVWLPZQKILFGGCLVKSLQAKDLGNTADADLNEWP | 41 | 0.00195468 | 0.0977338 | 0.0325779 | 9 | GDANVSAWPNSVEKVKKK | EVFYPGAGHTMDNIVVWLPQQKILFGGCLVKSLQAKDLGNTADADLNSWP | + |
| GPGHTKDNVVVWLPKEKILFGGCLVKSLGAGNLGL | EVFYPGPGHTMDNIVVWLPZQKILFGGCLVKSLQAKDLGNTADADLNEWP | 5 | 2.11493e-36 | 1.05746e-34 | 1.05746e-34 | 35 | GPGHTKDNVVVWLPKEKILFGGCLVKSLGAGNLGL | EVFYPGAGHTMDNIVVWLPQQKILFGGCLVKSLQAKDLGNTADADLNSWP | + |
| GPGHTKDNVVVWLPKEKILFGGCLVKSLGAGNLGL | VWHPSSGVLFGGCAVKDASAKSLGNVADADLAAWPASJERIRQRYPEARV | -10 | 1.86489e-11 | 9.32447e-10 | 4.66223e-10 | 25 | GPGHTKDNVVVWLPKEKILFGGCLVKSLGAGNLGL | VWHPSSGVLFGGCAVKDASAKSLGNVADADLAAWPASLERIRQRYPEARV | + |
| GPGHTKDNVVVWLPKEKILFGGCLVKSLGAGNLGL | VRFGPVELFFPGAGHSPDNLV | 11 | 0.000331714 | 0.0165857 | 0.00552857 | 10 | GPGHTKDNVVVWLPKEKILFGGCLVKSLGAGNLGL | VRFGPVELFFPGAGHSPDNLV | + |
| IEKJSENVYLHTSFKZTNGWG | KNEENQVEITKJAEGVWVHTSYGTYNGGT | 8 | 2.36013e-16 | 1.18006e-14 | 1.18006e-14 | 21 | IEKLSENVYLHTSFKETNGWG | KNEENQVEITKIAEGVWVHTSYGTYNGGT | + |
| IEKJSENVYLHTSFKZTNGWG | DVRVRRJAPGVWLHTTTQGFD | 3 | 2.92611e-05 | 0.00146306 | 0.000731528 | 18 | IEKLSENVYLHTSFKETNGWG | DVRVRRIAPGVWLHTTTQGFD | + |
| IEVFYPGAGHTKDNIVVWLPKQKILFGGCLVKS | EVFYPGPGHTMDNIVVWLPZQKILFGGCLVKSLQAKDLGNTADADLNEWP | -1 | 6.65925e-41 | 3.32963e-39 | 3.32963e-39 | 32 | IEVFYPGAGHTKDNIVVWLPKQKILFGGCLVKS | EVFYPGAGHTMDNIVVWLPQQKILFGGCLVKSLQAKDLGNTADADLNSWP | + |
| IEVFYPGAGHTKDNIVVWLPKQKILFGGCLVKS | VRFGPVELFFPGAGHSPDNLV | 5 | 5.71006e-13 | 2.85503e-11 | 1.42751e-11 | 16 | IEVFYPGAGHTKDNIVVWLPKQKILFGGCLVKS | VRFGPVELFFPGAGHSPDNLV | + |
| IEVFYPGAGHTKDNIVVWLPKQKILFGGCLVKS | VWHPSSGVLFGGCAVKDASAKSLGNVADADLAAWPASJERIRQRYPEARV | -16 | 5.42456e-07 | 2.71228e-05 | 9.04093e-06 | 17 | IEVFYPGAGHTKDNIVVWLPKQKILFGGCLVKS | VWHPSSGVLFGGCAVKDASAKSLGNVADADLAAWPASLERIRQRYPEARV | + |
| IPTYASELTNELLKKKGKPQA | HFHDDRTGGVPVLRARGIPVYALPDTARL | 17 | 7.84187e-05 | 0.00392094 | 0.00392094 | 12 | IPTYASELTNELLKKKGKPQA | HFHDDRTGGVPVLRARGIPVYALPDTARL | + |
| IPTYASELTNELLKKKGKPQA | KLAKEQGYEVPBPILDELTTL | -10 | 0.00101254 | 0.0506268 | 0.0253134 | 11 | IPTYASELTNELLKKKGKPQA | KLAKEQGYEVPNPILDELTTL | + |
| IPTYASELTNELLKKKGKPQA | VTHAHDDRIGGIDVLKKRGIPVYSTPLTA | 19 | 0.00244233 | 0.122116 | 0.0407054 | 10 | IPTYASELTNELLKKKGKPQA | VTHAHDDRIGGIDVLKKRGIPVYSTPLTA | + |
| KENNFVVPQNSFNDSLVLKVGBEKVIAKF | KLAKEQGYEVPBPILDELTTL | 3 | 1.96776e-05 | 0.000983878 | 0.000983878 | 18 | KENNFVVPQNSFNDSLVLKVGNEKVIAKF | KLAKEQGYEVPNPILDELTTL | + |
| KENNFVVPQNSFNDSLVLKVGBEKVIAKF | FGNTKV | -19 | 0.00166511 | 0.0832553 | 0.0416276 | 6 | KENNFVVPQNSFNDSLVLKVGNEKVIAKF | FGNTKV | + |
| KJADGVYLHTSYKEVEGFGLVSSNGLVVV | KNEENQVEITKJAEGVWVHTSYGTYNGGT | 10 | 7.42327e-08 | 3.71163e-06 | 3.71163e-06 | 19 | KLADGVYLHTSYKEVEGFGLVSSNGLVVV | KNEENQVEITKIAEGVWVHTSYGTYNGGT | + |
| KJADGVYLHTSYKEVEGFGLVSSNGLVVV | DVRVRRJAPGVWLHTTTQGFD | 5 | 7.79023e-05 | 0.00389511 | 0.00194756 | 16 | KLADGVYLHTSYKEVEGFGLVSSNGLVVV | DVRVRRIAPGVWLHTTTQGFD | + |
| KKDGKVQATHSFSGVNFWLVKNKIEVFYP | AEAZGNPVPTHSLAG | 1 | 0.000197249 | 0.00986247 | 0.00986247 | 14 | KKDGKVQATHSFSGVNFWLVKNKIEVFYP | AEAEGNPVPTHSLAG | + |
| MFSRILILLAGFJCCPPYAHSAESSGQLTITPLSSGALVVA | MMKFRILGVFF | 1 | 0.000456316 | 0.0228158 | 0.0228158 | 10 | MFSRILILLAGFLCCVPLAHSAESSGQLTITPLLSGALVVA | MMKFRILGVFF | + |
| MFSRILILLAGFJCCPPYAHSAESSGQLTITPLSSGALVVA | AEAZGNPVPTHSLAG | -21 | 0.00105903 | 0.0529515 | 0.0264757 | 15 | MFSRILILLAGFLCCVPLAHSAESSGQLTITPLLSGALVVA | AEAEGNPVPTHSLAG | + |
| MFSSINALGKFTLYVLFIISFNLNAADPK | RIFISIVFFMQTFGLVFAEPD | -8 | 9.26907e-05 | 0.00463454 | 0.00463454 | 21 | MFSSINALGKFTLYVLFIISFNLNAADPK | RIFISIVFFMQTFGLVFAEPD | + |
| MKFLLLFLJ | MKRILLLLLLL | 0 | 3.77077e-10 | 1.88539e-08 | 1.88539e-08 | 9 | MKFLLLFLL | MKRILLLLLLL | + |
| MKFLLLFLJ | MKSLFAALFVVMFCGLARASD | 0 | 0.00025171 | 0.0125855 | 0.00481665 | 9 | MKFLLLFLL | MKSLFAALFVVMFCGLARASD | + |
| MKFLLLFLJ | MFKLJSKLLVY | 0 | 0.000288999 | 0.01445 | 0.00481665 | 9 | MKFLLLFLL | MFKLLSKLLVY | + |
| MKFLLLFLJ | MMKFRILGVFF | 1 | 0.000458157 | 0.0229079 | 0.00572697 | 9 | MKFLLLFLL | MMKFRILGVFF | + |
| MKKJFLLLJFL | MKRILLLLLLL | 0 | 3.39937e-11 | 1.69968e-09 | 1.69968e-09 | 11 | MKKLFLLLLFL | MKRILLLLLLL | + |
| MKKJFLLLJFL | MMKFRILGVFF | 1 | 0.000203492 | 0.0101746 | 0.00454014 | 10 | MKKLFLLLLFL | MMKFRILGVFF | + |
| MKKJFLLLJFL | MFKLJSKLLVY | 0 | 0.000272409 | 0.0136204 | 0.00454014 | 11 | MKKLFLLLLFL | MFKLLSKLLVY | + |
| MKKJFLLLJFL | MKSLFAALFVVMFCGLARASD | 0 | 0.00309297 | 0.154649 | 0.0386621 | 11 | MKKLFLLLLFL | MKSLFAALFVVMFCGLARASD | + |
| MSIQHFRVALIPFFAAFCLPVFAGEQGH | MSIQHFRVALI | 0 | 3.8932e-18 | 1.9466e-16 | 1.9466e-16 | 11 | MSIQHFRVALIPFFAAFCLPVFALSQGH | MSIQHFRVALI | + |
| MSIQHFRVALIPFFAAFCLPVFAGEQGH | HFHDDRTGGVPVLRARGIPVYALPDTARL | -1 | 0.000680129 | 0.0340065 | 0.0170032 | 27 | MSIQHFRVALIPFFAAFCLPVFALSQGH | HFHDDRTGGVPVLRARGIPVYALPDTARL | + |
| PAYTLQAQEQEJQVTKJAPGVWVHTSYSTYNGVLVPSHGLVVSTKEGAVL | KNEENQVEITKJAEGVWVHTSYGTYNGGT | -5 | 3.91273e-21 | 1.95637e-19 | 1.95637e-19 | 29 | PAYTLLAQEQELQVTKLAPGVWVHTSYSTYGGVLVPSHGLVVSTKGGAVL | KNEENQVEITKIAEGVWVHTSYGTYNGGT | + |
| PAYTLQAQEQEJQVTKJAPGVWVHTSYSTYNGVLVPSHGLVVSTKEGAVL | DVRVRRJAPGVWLHTTTQGFD | -10 | 7.45773e-12 | 3.72886e-10 | 1.86443e-10 | 21 | PAYTLLAQEQELQVTKLAPGVWVHTSYSTYGGVLVPSHGLVVSTKGGAVL | DVRVRRIAPGVWLHTTTQGFD | + |
| PAYTLQAQEQEJQVTKJAPGVWVHTSYSTYNGVLVPSHGLVVSTKEGAVL | FPSNGLIVETGKGLVLIDTAWGEEQTEZL | -34 | 3.86718e-08 | 1.93359e-06 | 6.4453e-07 | 16 | PAYTLLAQEQELQVTKLAPGVWVHTSYSTYGGVLVPSHGLVVSTKGGAVL | FPSNGLIVETGKGLVLIDTAWGEEQTEEL | + |
| PAYTLQAQEQEJQVTKJAPGVWVHTSYSTYNGVLVPSHGLVVSTKEGAVL | YPANGLIVEDGDELLLVDTAWGARQTAALL | -34 | 0.000544408 | 0.0272204 | 0.0068051 | 16 | PAYTLLAQEQELQVTKLAPGVWVHTSYSTYGGVLVPSHGLVVSTKGGAVL | YPANGLIVEDGDELLLVDTAWGARQTAALL | + |
| PCNGLVVIDNKEAYJIDTPWNAKDTE | FPSNGLIVETGKGLVLIDTAWGEEQTEZL | 1 | 6.17829e-17 | 3.08915e-15 | 1.71447e-15 | 26 | PCNGLVVIDNKEAYLIDTPWNAKDTE | FPSNGLIVETGKGLVLIDTAWGEEQTEEL | + |
| PCNGLVVIDNKEAYJIDTPWNAKDTE | YPANGLIVEDGDELLLVDTAWGARQTAALL | 1 | 6.85788e-17 | 3.42894e-15 | 1.71447e-15 | 26 | PCNGLVVIDNKEAYLIDTPWNAKDTE | YPANGLIVEDGDELLLVDTAWGARQTAALL | + |
| RGYKJKASISTHFHEDSTGGJEYLNSHSIPTYASELTNELL | VTHAHDDRIGGIDVLKKRGIPVYSTPLTA | -9 | 2.16118e-20 | 1.08059e-18 | 1.08059e-18 | 29 | RGYKIKASISTHFHEDSTGGLEYLNSHSIPTYASELTNELL | VTHAHDDRIGGIDVLKKRGIPVYSTPLTA | + |
| RGYKJKASISTHFHEDSTGGJEYLNSHSIPTYASELTNELL | HFHDDRTGGVPVLRARGIPVYALPDTARL | -11 | 6.38567e-19 | 3.19283e-17 | 1.59642e-17 | 29 | RGYKIKASISTHFHEDSTGGLEYLNSHSIPTYASELTNELL | HFHDDRTGGVPVLRARGIPVYALPDTARL | + |
| SLGNTGDADLEAWPASIAKVQAR | VWHPSSGVLFGGCAVKDASAKSLGNVADADLAAWPASJERIRQRYPEARV | 21 | 1.48588e-19 | 7.42942e-18 | 7.42942e-18 | 23 | SLGNTGDADLEAWPASIAKVQAR | VWHPSSGVLFGGCAVKDASAKSLGNVADADLAAWPASLERIRQRYPEARV | + |
| SLGNTGDADLEAWPASIAKVQAR | EVFYPGPGHTMDNIVVWLPZQKILFGGCLVKSLQAKDLGNTADADLNEWP | 36 | 7.88808e-09 | 3.94404e-07 | 1.97202e-07 | 14 | SLGNTGDADLEAWPASIAKVQAR | EVFYPGAGHTMDNIVVWLPQQKILFGGCLVKSLQAKDLGNTADADLNSWP | + |
| SLGNTGDADLEAWPASIAKVQAR | SIKRVQQRYPK | -15 | 2.37632e-06 | 0.000118816 | 3.96053e-05 | 8 | SLGNTGDADLEAWPASIAKVQAR | SIKRVQQRYPK | + |
| THSFSGBQFWLVNGK | ATGPEEYVLAE | -2 | 0.000609187 | 0.0304594 | 0.0304594 | 11 | THSFSGDQFWLVNGK | ATGQEEYVLAE | + |
| VVPGHGKVGDASLLKHTIELA | KIVVPGHGEWGGKDLLSHTJKLLK | 2 | 5.36433e-22 | 2.68217e-20 | 2.68217e-20 | 21 | VVPGHGKVGDASLLKHTIELA | KIVVPGHGEWGGKDLLSHTLKLLK | + |
| VVPGHGKVGDASLLKHTIELA | VVPGHGLPGGPELLDHTEALL | 0 | 4.96692e-16 | 2.48346e-14 | 1.24173e-14 | 21 | VVPGHGKVGDASLLKHTIELA | VVPGHGLPGGPELLDHTEALL | + |
| YPDAKJ | VWHPSSGVLFGGCAVKDASAKSLGNVADADLAAWPASJERIRQRYPEARV | 44 | 9.84827e-06 | 0.000492413 | 0.000492413 | 6 | YPDAKL | VWHPSSGVLFGGCAVKDASAKSLGNVADADLAAWPASLERIRQRYPEARV | + |

**NDM vs IMP**

| Query_ID | Target_ID | Optimal_offset | p-value | E-value | q-value | Overlap | Query_consensus | Target_consensus | Orientation |
| --- | --- | --- | --- | --- | --- | --- | --- | --- | --- |
| ASNGLI | PJADGVYLHTSYKZVEGFGLVDSNGLVVV | 21 | 0.000209503 | 0.0104752 | 0.0104752 | 6 | ASNGLI | PLADGVYLHTSYKQVEGFGLVDSNGLVVV | + |
| ASNGLI | LVPANGLVVLDNKEAYJIDTPWTAKDTEKLVTWIEER | 2 | 0.00120476 | 0.060238 | 0.030119 | 6 | ASNGLI | LVPANGLVVLDNKEAYLIDTPWTAKDTEKLVTWIVER | + |
| ASNGLI | DQQEAIIFDTPATDQASEELI | 15 | 0.00263963 | 0.131982 | 0.0439939 | 6 | ASNGLI | DQQEAIIFDTPATDQASEELI | + |
| GCFGKD | EVFYPGAGHTKDNVVVWLPEEKILFGGCFVKS | 26 | 0.00164724 | 0.0823618 | 0.0478666 | 6 | AGLGAD | EVFYPGAGHTKDNVVVWLPEEKILFGGCFVKS | + |
| GCFGKD | IEVYYPGAGHTKDNLVVWLPKQKJLFGGCLVKSLESKSLGY | 27 | 0.0026721 | 0.133605 | 0.0478666 | 6 | AGLGAD | IEVFYPGAGHTKDNLVVWLPKQKILFGGCLVKSLESKSLGY | + |
| GCFGKD | WVETNLKQPVKAVVATHFHEDCLGGLQAF | 23 | 0.002872 | 0.1436 | 0.0478666 | 6 | AGLGAD | WVETNLKQPVKAVVATHFHDDCLGGLGAF | + |
| INLPVALA | LHNLPQASAAIKSGV | 1 | 0.00022682 | 0.011341 | 0.011341 | 8 | INLPVALA | LHNLPQASAAIKSGV | + |
| LGDADTEHYAASARAFGAAFPKASMIVMSHSAPDSRAAITHTARMADKLR.1 | ADADLDAWPASIAKVQARYPDAKIVVPGHGK | -1 | 8.58674e-07 | 4.29337e-05 | 4.29337e-05 | 31 | LGDADTEHYAASARAFGAAFPKASMIVMSHSAPDSRAAITHTARMADKLR | ADADLDAWPASIAKVQARYPDAKIVVPGHGK | + |
| LGDADTEHYAASARAFGAAFPKASMIVMSHSAPDSRAAITHTARMADKLR.1 | KGLGNLGDANIEAWPKSAKKVKSK | 5 | 0.00165481 | 0.0827403 | 0.0413701 | 19 | LGDADTEHYAASARAFGAAFPKASMIVMSHSAPDSRAAITHTARMADKLR | KGLGNLGDANIEAWPKSAKKVKSK | + |
| LGDADTEHYAASARAFGAAFPKASMIVMSHSAPDSRAAITHTARMADKLR.2 | ADADLDAWPASIAKVQARYPDAKIVVPGHGK | -1 | 9.04521e-07 | 4.5226e-05 | 4.5226e-05 | 31 | LGDADTEHYAASARAFGAAFPKASMIVMSHSAPDSRAAITHTARMADKLR | ADADLDAWPASIAKVQARYPDAKIVVPGHGK | + |
| LGDADTEHYAASARAFGAAFPKASMIVMSHSAPDSRAAITHTARMADKLR.2 | KGLGNLGDANIEAWPKSAKKVKSK | 5 | 0.00181495 | 0.0907476 | 0.0453738 | 19 | LGDADTEHYAASARAFGAAFPKASMIVMSHSAPDSRAAITHTARMADKLR | KGLGNLGDANIEAWPKSAKKVKSK | + |
| NLGDPDC | KGLGNLGDANIEAWPKSAKKVKSK | 4 | 1.867e-05 | 0.000933498 | 0.000933498 | 7 | NLGDADT | KGLGNLGDANIEAWPKSAKKVKSK | + |
| SIQHFRVALIPFFAAFCLPVF | MSIQHFRVALIPFFAAFCLPV | 1 | 5.00448e-29 | 2.50224e-27 | 2.50224e-27 | 20 | SIQHFRVALIPFFAAFCLPVF | MSIQHFRVALIPFFAAFCLPV | + |
| TYANALSNQLAPQEGMVAAQHSLTFAANG | IPTYASELTNALLAQNGKPLA | 2 | 3.51281e-06 | 0.00017564 | 0.00017564 | 19 | TYANALSNQLAPQEGMVAAQHSLTFAANG | IPTYASELTNALLAQNGKPLA | + |
| VRDGGRVLVVDTAWTDDQTAQILNWIKQEINLPVALAVVTHAHQDKMGGM | WVETNLKQPVKAVVATHFHEDCLGGLQAF | -24 | 8.83028e-09 | 4.41514e-07 | 4.41514e-07 | 26 | VRDGGRVLVVDTAWTDDQTAQILNWIKQEINLPVALAVVTHAHQDKMGGM | WVETNLKQPVKAVVATHFHDDCLGGLGAF | + |
| VRDGGRVLVVDTAWTDDQTAQILNWIKQEINLPVALAVVTHAHQDKMGGM | GKDAYJIDTPWTEADTEKLVDWIEQQGLTLKASVSTHSHEDRTGGIGYLN | -3 | 4.56129e-06 | 0.000228064 | 0.000114032 | 47 | VRDGGRVLVVDTAWTDDQTAQILNWIKQEINLPVALAVVTHAHQDKMGGM | GKDAYIIDTPWTEADTEKLVDWIEQQGLTLKASVSTHSHEDRTGGIGYLN | + |
| VRDGGRVLVVDTAWTDDQTAQILNWIKQEINLPVALAVVTHAHQDKMGGM | LVPANGLVVLDNKEAYJIDTPWTAKDTEKLVTWIEER | 8 | 3.99022e-05 | 0.00199511 | 0.000665037 | 29 | VRDGGRVLVVDTAWTDDQTAQILNWIKQEINLPVALAVVTHAHQDKMGGM | LVPANGLVVLDNKEAYLIDTPWTAKDTEKLVTWIVER | + |
| VRDGGRVLVVDTAWTDDQTAQILNWIKQEINLPVALAVVTHAHQDKMGGM | GVLFPSHGLVVSTKDGAVLVDTGWGNEPTEQLLA | 10 | 6.06963e-05 | 0.00303481 | 0.000758704 | 24 | VRDGGRVLVVDTAWTDDQTAQILNWIKQEINLPVALAVVTHAHQDKMGGM | GVLFPSHGLVVSTKGGAVLVDTGWGNEPTEQLLA | + |
| VRDGGRVLVVDTAWTDDQTAQILNWIKQEINLPVALAVVTHAHQDKMGGM | DQQEAIIFDTPATDQASEELI | -2 | 0.000419804 | 0.0209902 | 0.00419804 | 21 | VRDGGRVLVVDTAWTDDQTAQILNWIKQEINLPVALAVVTHAHQDKMGGM | DQQEAIIFDTPATDQASEELI | + |
| VVTHAHQDKMGGMDALHAAGIATYANALSNQLAPQEGMVAAQHSLTFAAN | GYKJKGSISTHFHEDSTGGJEWLNSHSIPTYASELTNELLK | 7 | 8.03838e-10 | 4.01919e-08 | 4.01919e-08 | 34 | VVTHAHQDKMGGMDALHAAGIATYANALSNQLAPQEGMVAAQHSLTFAAN | GYKIKGSISTHFHEDSTGGIEWLNSHSIPTYASELTNELLK | + |
| VVTHAHQDKMGGMDALHAAGIATYANALSNQLAPQEGMVAAQHSLTFAAN | IPTYASELTNALLAQNGKPLA | -20 | 1.99004e-07 | 9.95018e-06 | 4.97509e-06 | 21 | VVTHAHQDKMGGMDALHAAGIATYANALSNQLAPQEGMVAAQHSLTFAAN | IPTYASELTNALLAQNGKPLA | + |
| VVTHAHQDKMGGMDALHAAGIATYANALSNQLAPQEGMVAAQHSLTFAAN | WVETNLKQPVKAVVATHFHEDCLGGLQAF | 13 | 1.73178e-05 | 0.000865889 | 0.00028863 | 16 | VVTHAHQDKMGGMDALHAAGIATYANALSNQLAPQEGMVAAQHSLTFAAN | WVETNLKQPVKAVVATHFHDDCLGGLGAF | + |
| WQHTSYLDMPGFGAVASNGLIVRDGGRVLVVDTAWTDDQTAQILNWIKQE | LVPANGLVVLDNKEAYJIDTPWTAKDTEKLVTWIEER | -13 | 4.32399e-11 | 2.162e-09 | 2.162e-09 | 37 | WQHTSYLDMPGFGAVASNGLIVRDGGRVLVVDTAWTDDQTAQILNWIKQE | LVPANGLVVLDNKEAYLIDTPWTAKDTEKLVTWIVER | + |
| WQHTSYLDMPGFGAVASNGLIVRDGGRVLVVDTAWTDDQTAQILNWIKQE | GVLFPSHGLVVSTKDGAVLVDTGWGNEPTEQLLA | -11 | 2.5492e-10 | 1.2746e-08 | 6.373e-09 | 34 | WQHTSYLDMPGFGAVASNGLIVRDGGRVLVVDTAWTDDQTAQILNWIKQE | GVLFPSHGLVVSTKGGAVLVDTGWGNEPTEQLLA | + |
| WQHTSYLDMPGFGAVASNGLIVRDGGRVLVVDTAWTDDQTAQILNWIKQE | PJADGVYLHTSYKZVEGFGLVDSNGLVVV | 6 | 4.28912e-10 | 2.14456e-08 | 7.14854e-09 | 23 | WQHTSYLDMPGFGAVASNGLIVRDGGRVLVVDTAWTDDQTAQILNWIKQE | PLADGVYLHTSYKQVEGFGLVDSNGLVVV | + |
| WQHTSYLDMPGFGAVASNGLIVRDGGRVLVVDTAWTDDQTAQILNWIKQE | GKDAYJIDTPWTEADTEKLVDWIEQQGLTLKASVSTHSHEDRTGGIGYLN | -24 | 0.000312607 | 0.0156304 | 0.00321735 | 26 | WQHTSYLDMPGFGAVASNGLIVRDGGRVLVVDTAWTDDQTAQILNWIKQE | GKDAYIIDTPWTEADTEKLVDWIEQQGLTLKASVSTHSHEDRTGGIGYLN | + |
| WQHTSYLDMPGFGAVASNGLIVRDGGRVLVVDTAWTDDQTAQILNWIKQE | DQQEAIIFDTPATDQASEELI | -23 | 0.000321735 | 0.0160868 | 0.00321735 | 21 | WQHTSYLDMPGFGAVASNGLIVRDGGRVLVVDTAWTDDQTAQILNWIKQE | DQQEAIIFDTPATDQASEELI | + |
| WVEPATAPNFGPLKVFYPGPGHTSDNITVGIDGTDIAFGGCLIKDSKAKS.1 | EVFYPGAGHTKDNVVVWLPEEKILFGGCFVKS | -13 | 3.98017e-17 | 1.99008e-15 | 1.99008e-15 | 32 | WVEPATAPNFGPLKVFYPGPGHTSDNITVGIDGTDIAFGGCLIKDSKAKS | EVFYPGAGHTKDNVVVWLPEEKILFGGCFVKS | + |
| WVEPATAPNFGPLKVFYPGPGHTSDNITVGIDGTDIAFGGCLIKDSKAKS.1 | IEVYYPGAGHTKDNLVVWLPKQKJLFGGCLVKSLESKSLGY | -12 | 6.52137e-16 | 3.26069e-14 | 1.63034e-14 | 38 | WVEPATAPNFGPLKVFYPGPGHTSDNITVGIDGTDIAFGGCLIKDSKAKS | IEVFYPGAGHTKDNLVVWLPKQKILFGGCLVKSLESKSLGY | + |
| WVEPATAPNFGPLKVFYPGPGHTSDNITVGIDGTDIAFGGCLIKDSKAKS.2 | EVFYPGAGHTKDNVVVWLPEEKILFGGCFVKS | -13 | 3.98652e-17 | 1.99326e-15 | 1.99326e-15 | 32 | WVEPATAPNFGPLKVFYPGPGHTSDNITVGIDGTDIAFGGCLIKDSKAKS | EVFYPGAGHTKDNVVVWLPEEKILFGGCFVKS | + |
| WVEPATAPNFGPLKVFYPGPGHTSDNITVGIDGTDIAFGGCLIKDSKAKS.2 | IEVYYPGAGHTKDNLVVWLPKQKJLFGGCLVKSLESKSLGY | -12 | 5.74196e-16 | 2.87098e-14 | 1.43549e-14 | 38 | WVEPATAPNFGPLKVFYPGPGHTSDNITVGIDGTDIAFGGCLIKDSKAKS | IEVFYPGAGHTKDNLVVWLPKQKILFGGCLVKSLESKSLGY | + |

| ASNGLI | ASNKSIQPTAEASAD | 0 | 0.000118888 | 0.00594438 | 0.00594438 | 6 | ASNGLI | ASNKSIQPTAEASAD | + |
| --- | --- | --- | --- | --- | --- | --- | --- | --- | --- |
| ASNGLI | KJADGVYLHTSYKEVEGFGLVSSNGLVVV | 21 | 0.000244136 | 0.0122068 | 0.0061034 | 6 | ASNGLI | KLADGVYLHTSYKEVEGFGLVSSNGLVVV | + |
| ASNGLI | PCNGLVVIDNKEAYJIDTPWNAKDTE | 0 | 0.00148816 | 0.0744081 | 0.0248027 | 6 | ASNGLI | PCNGLVVIDNKEAYLIDTPWNAKDTE | + |
| GCFGKD | GPGHTKDNVVVWLPKEKILFGGCLVKSLGAGNLGL | 21 | 0.00160923 | 0.0804613 | 0.0463973 | 6 | AGLGAD | GPGHTKDNVVVWLPKEKILFGGCLVKSLGAGNLGL | + |
| GCFGKD | IEVFYPGAGHTKDNIVVWLPKQKILFGGCLVKS | 27 | 0.00185589 | 0.0927946 | 0.0463973 | 6 | AGLGAD | IEVFYPGAGHTKDNIVVWLPKQKILFGGCLVKS | + |
| INLPVALA | LHNLPQASAAIKSGV | 1 | 0.000244561 | 0.0122281 | 0.0122281 | 8 | INLPVALA | LHNLPQASAAIKSGV | + |
| NLGDPDC | SLGNTGDADLEAWPASIAKVQAR | 3 | 7.62451e-05 | 0.00381225 | 0.00381225 | 7 | NLGDADT | SLGNTGDADLEAWPASIAKVQAR | + |
| SIQHFRVALIPFFAAFCLPVF | MSIQHFRVALIPFFAAFCLPVFAGEQGH | 1 | 2.26469e-35 | 1.13234e-33 | 1.13234e-33 | 21 | SIQHFRVALIPFFAAFCLPVF | MSIQHFRVALIPFFAAFCLPVFAGSQGH | + |
| TYANALSNQLAPQEGMVAAQHSLTFAANG | IPTYASELTNELLKKKGKPQA | 2 | 2.49802e-06 | 0.000124901 | 0.000124901 | 19 | TYANALSNQLAPQEGMVAAQHSLTFAANG | IPTYASELTNELLKKNGKPQA | + |
| VRDGGRVLVVDTAWTDDQTAQILNWIKQEINLPVALAVVTHAHQDKMGGM | FDTPWTBEPTEQLLAWVKDNLKAPVKAFVPTHWHDDCLGGL | -9 | 6.53328e-17 | 3.26664e-15 | 3.26664e-15 | 41 | VRDGGRVLVVDTAWTDDQTAQILNWIKQEINLPVALAVVTHAHQDKMGGM | FDTPWTDEPTEQLLAWVKDNLKAPVKAFVPTHWHDDCLGGL | + |
| VRDGGRVLVVDTAWTDDQTAQILNWIKQEINLPVALAVVTHAHQDKMGGM | AYJIDTPWTEKDTEKLVDWIEAQGLTLKASISTHSHZDRTGGIGYLNSKG | -6 | 0.000133401 | 0.00667007 | 0.00333503 | 44 | VRDGGRVLVVDTAWTDDQTAQILNWIKQEINLPVALAVVTHAHQDKMGGM | AYIIDTPWTEKDTEKLVDWIEAQGLTLKASISTHSHEDRTGGIGYLNSIG | + |
| VRDGGRVLVVDTAWTDDQTAQILNWIKQEINLPVALAVVTHAHQDKMGGM | PCNGLVVIDNKEAYJIDTPWNAKDTE | 6 | 0.000627887 | 0.0313944 | 0.0104648 | 20 | VRDGGRVLVVDTAWTDDQTAQILNWIKQEINLPVALAVVTHAHQDKMGGM | PCNGLVVIDNKEAYLIDTPWNAKDTE | + |
| VVTHAHQDKMGGMDALHAAGIATYANALSNQLAPQEGMVAAQHSLTFAAN | RGYKJKASISTHFHEDSTGGJEYLNSHSIPTYASELTNELL | 8 | 2.29145e-10 | 1.14572e-08 | 1.14572e-08 | 33 | VVTHAHQDKMGGMDALHAAGIATYANALSNQLAPQEGMVAAQHSLTFAAN | RGYKIKASISTHFHEDSTGGLEYLNSHSIPTYASELTNELL | + |
| VVTHAHQDKMGGMDALHAAGIATYANALSNQLAPQEGMVAAQHSLTFAAN | IPTYASELTNELLKKKGKPQA | -20 | 1.83476e-07 | 9.1738e-06 | 4.5869e-06 | 21 | VVTHAHQDKMGGMDALHAAGIATYANALSNQLAPQEGMVAAQHSLTFAAN | IPTYASELTNELLKKNGKPQA | + |
| WQHTSYLDMPGFGAVASNGLIVRDGGRVLVVDTAWTDDQTAQILNWIKQE | KJADGVYLHTSYKEVEGFGLVSSNGLVVV | 6 | 8.88003e-10 | 4.44001e-08 | 4.44001e-08 | 23 | WQHTSYLDMPGFGAVASNGLIVRDGGRVLVVDTAWTDDQTAQILNWIKQE | KLADGVYLHTSYKEVEGFGLVSSNGLVVV | + |
| WQHTSYLDMPGFGAVASNGLIVRDGGRVLVVDTAWTDDQTAQILNWIKQE | PCNGLVVIDNKEAYJIDTPWNAKDTE | -15 | 6.2259e-09 | 3.11295e-07 | 1.55647e-07 | 26 | WQHTSYLDMPGFGAVASNGLIVRDGGRVLVVDTAWTDDQTAQILNWIKQE | PCNGLVVIDNKEAYLIDTPWNAKDTE | + |
| WQHTSYLDMPGFGAVASNGLIVRDGGRVLVVDTAWTDDQTAQILNWIKQE | FDTPWTBEPTEQLLAWVKDNLKAPVKAFVPTHWHDDCLGGL | -30 | 0.000100953 | 0.00504767 | 0.00168256 | 20 | WQHTSYLDMPGFGAVASNGLIVRDGGRVLVVDTAWTDDQTAQILNWIKQE | FDTPWTDEPTEQLLAWVKDNLKAPVKAFVPTHWHDDCLGGL | + |
| WQHTSYLDMPGFGAVASNGLIVRDGGRVLVVDTAWTDDQTAQILNWIKQE | IEKJSENVYLHTSFKZTNGWG | 8 | 0.00189828 | 0.0949141 | 0.0237285 | 13 | WQHTSYLDMPGFGAVASNGLIVRDGGRVLVVDTAWTDDQTAQILNWIKQE | IEKLSENVYLHTSFKETNGWG | + |
| WQHTSYLDMPGFGAVASNGLIVRDGGRVLVVDTAWTDDQTAQILNWIKQE | AYJIDTPWTEKDTEKLVDWIEAQGLTLKASISTHSHZDRTGGIGYLNSKG | -27 | 0.0040497 | 0.202485 | 0.0361237 | 23 | WQHTSYLDMPGFGAVASNGLIVRDGGRVLVVDTAWTDDQTAQILNWIKQE | AYIIDTPWTEKDTEKLVDWIEAQGLTLKASISTHSHEDRTGGIGYLNSIG | + |
| WQHTSYLDMPGFGAVASNGLIVRDGGRVLVVDTAWTDDQTAQILNWIKQE | PAYTLQAQEQEJQVTKJAPGVWVHTSYSTYNGVLVPSHGLVVSTKEGAVL | 21 | 0.00433485 | 0.216742 | 0.0361237 | 29 | WQHTSYLDMPGFGAVASNGLIVRDGGRVLVVDTAWTDDQTAQILNWIKQE | PAYTLLAQEQELQVTKLAPGVWVHTSYSTYGGVLVPSHGLVVSTKGGAVL | + |
| WVEPATAPNFGPLKVFYPGPGHTSDNITVGIDGTDIAFGGCLIKDSKAKS.1 | IEVFYPGAGHTKDNIVVWLPKQKILFGGCLVKS | -12 | 2.86339e-16 | 1.43169e-14 | 1.43169e-14 | 33 | WVEPATAPNFGPLKVFYPGPGHTSDNITVGIDGTDIAFGGCLIKDSKAKS | IEVFYPGAGHTKDNIVVWLPKQKILFGGCLVKS | + |
| WVEPATAPNFGPLKVFYPGPGHTSDNITVGIDGTDIAFGGCLIKDSKAKS.1 | GPGHTKDNVVVWLPKEKILFGGCLVKSLGAGNLGL | -18 | 7.47604e-12 | 3.73802e-10 | 1.86901e-10 | 32 | WVEPATAPNFGPLKVFYPGPGHTSDNITVGIDGTDIAFGGCLIKDSKAKS | GPGHTKDNVVVWLPKEKILFGGCLVKSLGAGNLGL | + |
| WVEPATAPNFGPLKVFYPGPGHTSDNITVGIDGTDIAFGGCLIKDSKAKS.2 | IEVFYPGAGHTKDNIVVWLPKQKILFGGCLVKS | -12 | 2.60425e-16 | 1.30213e-14 | 1.30213e-14 | 33 | WVEPATAPNFGPLKVFYPGPGHTSDNITVGIDGTDIAFGGCLIKDSKAKS | IEVFYPGAGHTKDNIVVWLPKQKILFGGCLVKS | + |
| WVEPATAPNFGPLKVFYPGPGHTSDNITVGIDGTDIAFGGCLIKDSKAKS.2 | GPGHTKDNVVVWLPKEKILFGGCLVKSLGAGNLGL | -18 | 6.8757e-12 | 3.43785e-10 | 1.71893e-10 | 32 | WVEPATAPNFGPLKVFYPGPGHTSDNITVGIDGTDIAFGGCLIKDSKAKS | GPGHTKDNVVVWLPKEKILFGGCLVKSLGAGNLGL | + |

**NDM vs VIM**

| ASNGLI | YPANGLJVEDGDGSJ.1 | 1 | 6.54775e-05 | 0.00327388 | 0.00163694 | 6 | ASNGLI | YPANGLIVEDGDGSL | + |
| --- | --- | --- | --- | --- | --- | --- | --- | --- | --- |
| ASNGLI | YPANGLJVEDGDGSJ.2 | 1 | 6.54775e-05 | 0.00327388 | 0.00163694 | 6 | ASNGLI | YPANGLIVEDGDGSL | + |
| DALHAAGI | HRPVRAAVVTHFHDDRTGGIPALVARGIPVYALED.1 | 20 | 0.000993015 | 0.0496507 | 0.0248254 | 8 | DALHAAGI | HRPVRAAVVTHFHDDRTGGIPALVARGIPVHALED | + |
| DALHAAGI | HRPVRAAVVTHFHDDRTGGIPALVARGIPVYALED.2 | 20 | 0.000993015 | 0.0496507 | 0.0248254 | 8 | DALHAAGI | HRPVRAAVVTHFHDDRTGGIPALVARGIPVHALED | + |
| GCFGKD | DAQTLGPLEVFFPGAGHAPDNJVVWHPASGVLFGGCFVKDA.1 | 34 | 0.00116114 | 0.0580569 | 0.0290284 | 6 | AGLGAD | DAQTLGPLEVFFPGAGHAPDNLVVWHPASGVLFGGCFVKDA | + |
| GCFGKD | DAQTLGPLEVFFPGAGHAPDNJVVWHPASGVLFGGCFVKDA.2 | 34 | 0.00116114 | 0.0580569 | 0.0290284 | 6 | AGLGAD | DAQTLGPLEVFFPGAGHAPDNLVVWHPASGVLFGGCFVKDA | + |
| LGDADTEHYAASARAFGAAFPKASMIVMSHSAPDSRAAITHTARMADKLR.1 | RARYPEARVVVPGHGAPGGPELLDHTEAL.1 | -16 | 0.000230192 | 0.0115096 | 0.00575479 | 29 | LGDADTEHYAASARAFGAAFPKASMIVMSHSAPDSRAAITHTARMADKLR | RARYPEARVVVPGHGAPGGPELLDHTEAL | + |
| LGDADTEHYAASARAFGAAFPKASMIVMSHSAPDSRAAITHTARMADKLR.1 | RARYPEARVVVPGHGAPGGPELLDHTEAL.2 | -16 | 0.000230192 | 0.0115096 | 0.00575479 | 29 | LGDADTEHYAASARAFGAAFPKASMIVMSHSAPDSRAAITHTARMADKLR | RARYPEARVVVPGHGAPGGPELLDHTEAL | + |
| LGDADTEHYAASARAFGAAFPKASMIVMSHSAPDSRAAITHTARMADKLR.2 | RARYPEARVVVPGHGAPGGPELLDHTEAL.1 | -16 | 0.000228246 | 0.0114123 | 0.00570616 | 29 | LGDADTEHYAASARAFGAAFPKASMIVMSHSAPDSRAAITHTARMADKLR | RARYPEARVVVPGHGAPGGPELLDHTEAL | + |
| LGDADTEHYAASARAFGAAFPKASMIVMSHSAPDSRAAITHTARMADKLR.2 | RARYPEARVVVPGHGAPGGPELLDHTEAL.2 | -16 | 0.000228246 | 0.0114123 | 0.00570616 | 29 | LGDADTEHYAASARAFGAAFPKASMIVMSHSAPDSRAAITHTARMADKLR | RARYPEARVVVPGHGAPGGPELLDHTEAL | + |
| NLGDPDC | AKDLGNLADADVAAWPASLER.1 | 5 | 0.000124795 | 0.00623976 | 0.00311988 | 7 | NLGDADT | AKDLGNLADADVAAWPASLER | + |
| NLGDPDC | AKDLGNLADADVAAWPASLER.2 | 5 | 0.000124795 | 0.00623976 | 0.00311988 | 7 | NLGDADT | AKDLGNLADADVAAWPASLER | + |
| VRDGGRVLVVDTAWTDDQTAQILNWIKQEINLPVALAVVTHAHQDKMGGM | LVDTGWGPRQTEALLDWARDTL.1 | -8 | 6.30473e-08 | 3.15236e-06 | 1.57618e-06 | 22 | VRDGGRVLVVDTAWTDDQTAQILNWIKQEINLPVALAVVTHAHQDKMGGM | LVDTGWGPRQTEALLDWARDTL | + |
| VRDGGRVLVVDTAWTDDQTAQILNWIKQEINLPVALAVVTHAHQDKMGGM | LVDTGWGPRQTEALLDWARDTL.2 | -8 | 6.30473e-08 | 3.15236e-06 | 1.57618e-06 | 22 | VRDGGRVLVVDTAWTDDQTAQILNWIKQEINLPVALAVVTHAHQDKMGGM | LVDTGWGPRQTEALLDWARDTL | + |
| VRDGGRVLVVDTAWTDDQTAQILNWIKQEINLPVALAVVTHAHQDKMGGM | HRPVRAAVVTHFHDDRTGGIPALVARGIPVYALED.1 | -30 | 5.70504e-06 | 0.000285252 | 7.1313e-05 | 20 | VRDGGRVLVVDTAWTDDQTAQILNWIKQEINLPVALAVVTHAHQDKMGGM | HRPVRAAVVTHFHDDRTGGIPALVARGIPVHALED | + |
| VRDGGRVLVVDTAWTDDQTAQILNWIKQEINLPVALAVVTHAHQDKMGGM | HRPVRAAVVTHFHDDRTGGIPALVARGIPVYALED.2 | -30 | 5.70504e-06 | 0.000285252 | 7.1313e-05 | 20 | VRDGGRVLVVDTAWTDDQTAQILNWIKQEINLPVALAVVTHAHQDKMGGM | HRPVRAAVVTHFHDDRTGGIPALVARGIPVHALED | + |
| VVTHAHQDKMGGMDALHAAGIATYANALSNQLAPQEGMVAAQHSLTFAAN | HRPVRAAVVTHFHDDRTGGIPALVARGIPVYALED.1 | 7 | 3.64942e-09 | 1.82471e-07 | 9.12355e-08 | 28 | VVTHAHQDKMGGMDALHAAGIATYANALSNQLAPQEGMVAAQHSLTFAAN | HRPVRAAVVTHFHDDRTGGIPALVARGIPVHALED | + |
| VVTHAHQDKMGGMDALHAAGIATYANALSNQLAPQEGMVAAQHSLTFAAN | HRPVRAAVVTHFHDDRTGGIPALVARGIPVYALED.2 | 7 | 3.64942e-09 | 1.82471e-07 | 9.12355e-08 | 28 | VVTHAHQDKMGGMDALHAAGIATYANALSNQLAPQEGMVAAQHSLTFAAN | HRPVRAAVVTHFHDDRTGGIPALVARGIPVHALED | + |
| WQHTSYLDMPGFGAVASNGLIVRDGGRVLVVDTAWTDDQTAQILNWIKQE | YPANGLJVEDGDGSJ.1 | -14 | 1.75734e-07 | 8.78671e-06 | 4.39336e-06 | 15 | WQHTSYLDMPGFGAVASNGLIVRDGGRVLVVDTAWTDDQTAQILNWIKQE | YPANGLIVEDGDGSL | + |
| WQHTSYLDMPGFGAVASNGLIVRDGGRVLVVDTAWTDDQTAQILNWIKQE | YPANGLJVEDGDGSJ.2 | -14 | 1.75734e-07 | 8.78671e-06 | 4.39336e-06 | 15 | WQHTSYLDMPGFGAVASNGLIVRDGGRVLVVDTAWTDDQTAQILNWIKQE | YPANGLIVEDGDGSL | + |
| WQHTSYLDMPGFGAVASNGLIVRDGGRVLVVDTAWTDDQTAQILNWIKQE | LVDTGWGPRQTEALLDWARDTL.1 | -29 | 3.85211e-07 | 1.92606e-05 | 4.81514e-06 | 21 | WQHTSYLDMPGFGAVASNGLIVRDGGRVLVVDTAWTDDQTAQILNWIKQE | LVDTGWGPRQTEALLDWARDTL | + |
| WQHTSYLDMPGFGAVASNGLIVRDGGRVLVVDTAWTDDQTAQILNWIKQE | LVDTGWGPRQTEALLDWARDTL.2 | -29 | 3.85211e-07 | 1.92606e-05 | 4.81514e-06 | 21 | WQHTSYLDMPGFGAVASNGLIVRDGGRVLVVDTAWTDDQTAQILNWIKQE | LVDTGWGPRQTEALLDWARDTL | + |
| WVEPATAPNFGPLKVFYPGPGHTSDNITVGIDGTDIAFGGCLIKDSKAKS.1 | DAQTLGPLEVFFPGAGHAPDNJVVWHPASGVLFGGCFVKDA.1 | -5 | 7.69633e-17 | 3.84817e-15 | 1.92408e-15 | 41 | WVEPATAPNFGPLKVFYPGPGHTSDNITVGIDGTDIAFGGCLIKDSKAKS | DAQTLGPLEVFFPGAGHAPDNLVVWHPASGVLFGGCFVKDA | + |
| WVEPATAPNFGPLKVFYPGPGHTSDNITVGIDGTDIAFGGCLIKDSKAKS.1 | DAQTLGPLEVFFPGAGHAPDNJVVWHPASGVLFGGCFVKDA.2 | -5 | 7.69633e-17 | 3.84817e-15 | 1.92408e-15 | 41 | WVEPATAPNFGPLKVFYPGPGHTSDNITVGIDGTDIAFGGCLIKDSKAKS | DAQTLGPLEVFFPGAGHAPDNLVVWHPASGVLFGGCFVKDA | + |
| WVEPATAPNFGPLKVFYPGPGHTSDNITVGIDGTDIAFGGCLIKDSKAKS.2 | DAQTLGPLEVFFPGAGHAPDNJVVWHPASGVLFGGCFVKDA.1 | -5 | 7.18357e-17 | 3.59179e-15 | 1.79589e-15 | 41 | WVEPATAPNFGPLKVFYPGPGHTSDNITVGIDGTDIAFGGCLIKDSKAKS | DAQTLGPLEVFFPGAGHAPDNLVVWHPASGVLFGGCFVKDA | + |
| WVEPATAPNFGPLKVFYPGPGHTSDNITVGIDGTDIAFGGCLIKDSKAKS.2 | DAQTLGPLEVFFPGAGHAPDNJVVWHPASGVLFGGCFVKDA.2 | -5 | 7.18357e-17 | 3.59179e-15 | 1.79589e-15 | 41 | WVEPATAPNFGPLKVFYPGPGHTSDNITVGIDGTDIAFGGCLIKDSKAKS | DAQTLGPLEVFFPGAGHAPDNLVVWHPASGVLFGGCFVKDA | + |
| ASNGLI | FPSNGLIVETGKGLVLIDTAWGEEQTEZL | 1 | 4.74036e-05 | 0.00237018 | 0.00237018 | 6 | ASNGLI | FPSNGLIVETGKGLVLIDTAWGEEQTEEL | + |
| ASNGLI | YPANGLIVEDGDELLLVDTAWGARQTAALL | 1 | 0.000101945 | 0.00509723 | 0.00254862 | 6 | ASNGLI | YPANGLIVEDGDELLLVDTAWGARQTAALL | + |
| DALHAAGI | HFHDDRTGGVPVLRARGIPVYALPDTARL | 10 | 3.82655e-05 | 0.00191328 | 0.00191328 | 8 | DALHAAGI | HFHDDRTGGVPVLRARGIPVYALPDTARL | + |
| DALHAAGI | VTHAHDDRIGGIDVLKKRGIPVYSTPLTA | 12 | 0.00139913 | 0.0699566 | 0.0349783 | 8 | DALHAAGI | VTHAHDDRIGGIDVLKKRGIPVYSTPLTA | + |
| NLGDPDC | VWHPSSGVLFGGCAVKDASAKSLGNVADADLAAWPASJERIRQRYPEARV | 24 | 0.000625759 | 0.0312879 | 0.0312879 | 7 | NLGDADT | VWHPSSGVLFGGCAVKDASAKSLGNVADADLAAWPASLERIRQRYPEARV | + |
| QEGNLP | AAAVERNIPVK | 3 | 0.000374555 | 0.0187278 | 0.0187278 | 6 | VTGNLA | AAAVERNIPVK | + |
| SIQHFRVALIPFFAAFCLPVF | MSIQHFRVALI | 1 | 4.28423e-13 | 2.14211e-11 | 2.14211e-11 | 10 | SIQHFRVALIPFFAAFCLPVF | MSIQHFRVALI | + |
| VRDGGRVLVVDTAWTDDQTAQILNWIKQEINLPVALAVVTHAHQDKMGGM | YPANGLIVEDGDELLLVDTAWGARQTAALL | 7 | 5.50663e-09 | 2.75332e-07 | 2.75332e-07 | 23 | VRDGGRVLVVDTAWTDDQTAQILNWIKQEINLPVALAVVTHAHQDKMGGM | YPANGLIVEDGDELLLVDTAWGARQTAALL | + |
| VRDGGRVLVVDTAWTDDQTAQILNWIKQEINLPVALAVVTHAHQDKMGGM | AEDTJGLPVRAAVVT | -25 | 9.63592e-07 | 4.81796e-05 | 2.40898e-05 | 15 | VRDGGRVLVVDTAWTDDQTAQILNWIKQEINLPVALAVVTHAHQDKMGGM | AEDTLGLPVRAAVVT | + |
| VRDGGRVLVVDTAWTDDQTAQILNWIKQEINLPVALAVVTHAHQDKMGGM | FPSNGLIVETGKGLVLIDTAWGEEQTEZL | 7 | 1.09477e-05 | 0.000547383 | 0.000182461 | 22 | VRDGGRVLVVDTAWTDDQTAQILNWIKQEINLPVALAVVTHAHQDKMGGM | FPSNGLIVETGKGLVLIDTAWGEEQTEEL | + |
| VRDGGRVLVVDTAWTDDQTAQILNWIKQEINLPVALAVVTHAHQDKMGGM | EWIKTKLKKPVKKAI | -23 | 1.73607e-05 | 0.000868035 | 0.000217009 | 15 | VRDGGRVLVVDTAWTDDQTAQILNWIKQEINLPVALAVVTHAHQDKMGGM | EWIKTKLKKPVKKAI | + |
| VRDGGRVLVVDTAWTDDQTAQILNWIKQEINLPVALAVVTHAHQDKMGGM | VTHAHDDRIGGIDVLKKRGIPVYSTPLTA | -38 | 0.000929113 | 0.0464557 | 0.00929113 | 12 | VRDGGRVLVVDTAWTDDQTAQILNWIKQEINLPVALAVVTHAHQDKMGGM | VTHAHDDRIGGIDVLKKRGIPVYSTPLTA | + |
| VRDGGRVLVVDTAWTDDQTAQILNWIKQEINLPVALAVVTHAHQDKMGGM | QQRAKTEDLLLPVQSVVTLATGEKYQDGP | -22 | 0.00156518 | 0.0782588 | 0.0130431 | 28 | VRDGGRVLVVDTAWTDDQTAQILNWIKQEINLPVALAVVTHAHQDKMGGM | QQRAKTEDLLLPVQSVVTLATGEKYQDGP | + |
| VVTHAHQDKMGGMDALHAAGIATYANALSNQLAPQEGMVAAQHSLTFAAN | VTHAHDDRIGGIDVLKKRGIPVYSTPLTA | -1 | 7.93061e-13 | 3.9653e-11 | 3.9653e-11 | 29 | VVTHAHQDKMGGMDALHAAGIATYANALSNQLAPQEGMVAAQHSLTFAAN | VTHAHDDRIGGIDVLKKRGIPVYSTPLTA | + |
| VVTHAHQDKMGGMDALHAAGIATYANALSNQLAPQEGMVAAQHSLTFAAN | HFHDDRTGGVPVLRARGIPVYALPDTARL | -3 | 2.79725e-11 | 1.39862e-09 | 6.99312e-10 | 29 | VVTHAHQDKMGGMDALHAAGIATYANALSNQLAPQEGMVAAQHSLTFAAN | HFHDDRTGGVPVLRARGIPVYALPDTARL | + |
| WQHTSYLDMPGFGAVASNGLIVRDGGRVLVVDTAWTDDQTAQILNWIKQE | YPANGLIVEDGDELLLVDTAWGARQTAALL | -14 | 1.17153e-15 | 5.85767e-14 | 5.85767e-14 | 30 | WQHTSYLDMPGFGAVASNGLIVRDGGRVLVVDTAWTDDQTAQILNWIKQE | YPANGLIVEDGDELLLVDTAWGARQTAALL | + |
| WQHTSYLDMPGFGAVASNGLIVRDGGRVLVVDTAWTDDQTAQILNWIKQE | FPSNGLIVETGKGLVLIDTAWGEEQTEZL | -14 | 3.77902e-12 | 1.88951e-10 | 9.44755e-11 | 29 | WQHTSYLDMPGFGAVASNGLIVRDGGRVLVVDTAWTDDQTAQILNWIKQE | FPSNGLIVETGKGLVLIDTAWGEEQTEEL | + |
| WVEPATAPNFGPLKVFYPGPGHTSDNITVGIDGTDIAFGGCLIKDSKAKS.1 | EVFYPGPGHTMDNIVVWLPZQKILFGGCLVKSLQAKDLGNTADADLNEWP | -13 | 2.68649e-15 | 1.34324e-13 | 1.34324e-13 | 37 | WVEPATAPNFGPLKVFYPGPGHTSDNITVGIDGTDIAFGGCLIKDSKAKS | EVFYPGAGHTMDNIVVWLPQQKILFGGCLVKSLQAKDLGNTADADLNSWP | + |
| WVEPATAPNFGPLKVFYPGPGHTSDNITVGIDGTDIAFGGCLIKDSKAKS.1 | VRFGPVELFFPGAGHSPDNLV | -7 | 8.86087e-11 | 4.43044e-09 | 2.21522e-09 | 21 | WVEPATAPNFGPLKVFYPGPGHTSDNITVGIDGTDIAFGGCLIKDSKAKS | VRFGPVELFFPGAGHSPDNLV | + |
| WVEPATAPNFGPLKVFYPGPGHTSDNITVGIDGTDIAFGGCLIKDSKAKS.1 | FGNTKV | -9 | 0.00238686 | 0.119343 | 0.0397811 | 6 | WVEPATAPNFGPLKVFYPGPGHTSDNITVGIDGTDIAFGGCLIKDSKAKS | FGNTKV | + |
| WVEPATAPNFGPLKVFYPGPGHTSDNITVGIDGTDIAFGGCLIKDSKAKS.2 | EVFYPGPGHTMDNIVVWLPZQKILFGGCLVKSLQAKDLGNTADADLNEWP | -13 | 2.53456e-15 | 1.26728e-13 | 1.26728e-13 | 37 | WVEPATAPNFGPLKVFYPGPGHTSDNITVGIDGTDIAFGGCLIKDSKAKS | EVFYPGAGHTMDNIVVWLPQQKILFGGCLVKSLQAKDLGNTADADLNSWP | + |
| WVEPATAPNFGPLKVFYPGPGHTSDNITVGIDGTDIAFGGCLIKDSKAKS.2 | VRFGPVELFFPGAGHSPDNLV | -7 | 8.89924e-11 | 4.44962e-09 | 2.22481e-09 | 21 | WVEPATAPNFGPLKVFYPGPGHTSDNITVGIDGTDIAFGGCLIKDSKAKS | VRFGPVELFFPGAGHSPDNLV | + |

**VIM vs IMP**

| Query_ID | Target_ID | Optimal_offset | p-value | E-value | q-value | Overlap | Query_consensus | Target_consensus | Orientation |
| --- | --- | --- | --- | --- | --- | --- | --- | --- | --- |
| AKDLGNLADADVAAWPASLER.1 | KGLGNLGDANIEAWPKSAKKVKSK | -1 | 4.07514e-12 | 2.03757e-10 | 2.03757e-10 | 20 | AKDLGNLADADVAAWPASLER | KGLGNLGDANIEAWPKSAKKVKSK | + |
| AKDLGNLADADVAAWPASLER.1 | ADADLDAWPASIAKVQARYPDAKIVVPGHGK | -7 | 3.84104e-08 | 1.92052e-06 | 9.60261e-07 | 14 | AKDLGNLADADVAAWPASLER | ADADLDAWPASIAKVQARYPDAKIVVPGHGK | + |
| AKDLGNLADADVAAWPASLER.2 | KGLGNLGDANIEAWPKSAKKVKSK | -1 | 4.07514e-12 | 2.03757e-10 | 2.03757e-10 | 20 | AKDLGNLADADVAAWPASLER | KGLGNLGDANIEAWPKSAKKVKSK | + |
| AKDLGNLADADVAAWPASLER.2 | ADADLDAWPASIAKVQARYPDAKIVVPGHGK | -7 | 3.84104e-08 | 1.92052e-06 | 9.60261e-07 | 14 | AKDLGNLADADVAAWPASLER | ADADLDAWPASIAKVQARYPDAKIVVPGHGK | + |
| CPLLLLLTACASTPS.1 | MKFLAPFLFLLPCVTVATEAS | 4 | 0.000160454 | 0.00802272 | 0.00802272 | 15 | CPLLLLLTACASTPS | MKFLAPFLFLLPCVTVATEAS | + |
| CPLLLLLTACASTPS.2 | MKFLAPFLFLLPCVTVATEAS | 4 | 0.000160454 | 0.00802272 | 0.00802272 | 15 | CPLLLLLTACASTPS | MKFLAPFLFLLPCVTVATEAS | + |
| DAQTLGPLEVFFPGAGHAPDNJVVWHPASGVLFGGCFVKDA.1 | EVFYPGAGHTKDNVVVWLPEEKILFGGCFVKS | -8 | 5.45554e-30 | 2.72777e-28 | 2.72777e-28 | 32 | DAQTLGPLEVFFPGAGHAPDNLVVWHPASGVLFGGCFVKDA | EVFYPGAGHTKDNVVVWLPEEKILFGGCFVKS | + |
| DAQTLGPLEVFFPGAGHAPDNJVVWHPASGVLFGGCFVKDA.1 | IEVYYPGAGHTKDNLVVWLPKQKJLFGGCLVKSLESKSLGY | -7 | 1.62204e-26 | 8.11022e-25 | 4.05511e-25 | 34 | DAQTLGPLEVFFPGAGHAPDNLVVWHPASGVLFGGCFVKDA | IEVFYPGAGHTKDNLVVWLPKQKILFGGCLVKSLESKSLGY | + |
| DAQTLGPLEVFFPGAGHAPDNJVVWHPASGVLFGGCFVKDA.2 | EVFYPGAGHTKDNVVVWLPEEKILFGGCFVKS | -8 | 5.45554e-30 | 2.72777e-28 | 2.72777e-28 | 32 | DAQTLGPLEVFFPGAGHAPDNLVVWHPASGVLFGGCFVKDA | EVFYPGAGHTKDNVVVWLPEEKILFGGCFVKS | + |
| DAQTLGPLEVFFPGAGHAPDNJVVWHPASGVLFGGCFVKDA.2 | IEVYYPGAGHTKDNLVVWLPKQKJLFGGCLVKSLESKSLGY | -7 | 1.62204e-26 | 8.11022e-25 | 4.05511e-25 | 34 | DAQTLGPLEVFFPGAGHAPDNLVVWHPASGVLFGGCFVKDA | IEVFYPGAGHTKDNLVVWLPKQKILFGGCLVKSLESKSLGY | + |
| HRPVRAAVVTHFHDDRTGGIPALVARGIPVYALED.1 | GYKJKGSISTHFHEDSTGGJEWLNSHSIPTYASELTNELLK | 0 | 3.63185e-18 | 1.81592e-16 | 1.81592e-16 | 35 | HRPVRAAVVTHFHDDRTGGIPALVARGIPVHALED | GYKIKGSISTHFHEDSTGGIEWLNSHSIPTYASELTNELLK | + |
| HRPVRAAVVTHFHDDRTGGIPALVARGIPVYALED.1 | WVETNLKQPVKAVVATHFHEDCLGGLQAF | 6 | 4.65863e-13 | 2.32931e-11 | 1.16466e-11 | 23 | HRPVRAAVVTHFHDDRTGGIPALVARGIPVHALED | WVETNLKQPVKAVVATHFHDDCLGGLGAF | + |
| HRPVRAAVVTHFHDDRTGGIPALVARGIPVYALED.1 | GKDAYJIDTPWTEADTEKLVDWIEQQGLTLKASVSTHSHEDRTGGIGYLN | 26 | 1.11217e-09 | 5.56083e-08 | 1.85361e-08 | 24 | HRPVRAAVVTHFHDDRTGGIPALVARGIPVHALED | GKDAYIIDTPWTEADTEKLVDWIEQQGLTLKASVSTHSHEDRTGGIGYLN | + |
| HRPVRAAVVTHFHDDRTGGIPALVARGIPVYALED.1 | MSIQHFRVALIPFFAAFCLPV | -9 | 0.00351392 | 0.175696 | 0.0439239 | 21 | HRPVRAAVVTHFHDDRTGGIPALVARGIPVHALED | MSIQHFRVALIPFFAAFCLPV | + |
| HRPVRAAVVTHFHDDRTGGIPALVARGIPVYALED.1 | GVLFPSHGLVVSTKDGAVLVDTGWGNEPTEQLLA | -4 | 0.00496714 | 0.248357 | 0.0496714 | 31 | HRPVRAAVVTHFHDDRTGGIPALVARGIPVHALED | GVLFPSHGLVVSTKGGAVLVDTGWGNEPTEQLLA | + |
| HRPVRAAVVTHFHDDRTGGIPALVARGIPVYALED.2 | GYKJKGSISTHFHEDSTGGJEWLNSHSIPTYASELTNELLK | 0 | 3.63185e-18 | 1.81592e-16 | 1.81592e-16 | 35 | HRPVRAAVVTHFHDDRTGGIPALVARGIPVHALED | GYKIKGSISTHFHEDSTGGIEWLNSHSIPTYASELTNELLK | + |
| HRPVRAAVVTHFHDDRTGGIPALVARGIPVYALED.2 | WVETNLKQPVKAVVATHFHEDCLGGLQAF | 6 | 4.65863e-13 | 2.32931e-11 | 1.16466e-11 | 23 | HRPVRAAVVTHFHDDRTGGIPALVARGIPVHALED | WVETNLKQPVKAVVATHFHDDCLGGLGAF | + |
| HRPVRAAVVTHFHDDRTGGIPALVARGIPVYALED.2 | GKDAYJIDTPWTEADTEKLVDWIEQQGLTLKASVSTHSHEDRTGGIGYLN | 26 | 1.11217e-09 | 5.56083e-08 | 1.85361e-08 | 24 | HRPVRAAVVTHFHDDRTGGIPALVARGIPVHALED | GKDAYIIDTPWTEADTEKLVDWIEQQGLTLKASVSTHSHEDRTGGIGYLN | + |
| HRPVRAAVVTHFHDDRTGGIPALVARGIPVYALED.2 | MSIQHFRVALIPFFAAFCLPV | -9 | 0.00351392 | 0.175696 | 0.0439239 | 21 | HRPVRAAVVTHFHDDRTGGIPALVARGIPVHALED | MSIQHFRVALIPFFAAFCLPV | + |
| HRPVRAAVVTHFHDDRTGGIPALVARGIPVYALED.2 | GVLFPSHGLVVSTKDGAVLVDTGWGNEPTEQLLA | -4 | 0.00496714 | 0.248357 | 0.0496714 | 31 | HRPVRAAVVTHFHDDRTGGIPALVARGIPVHALED | GVLFPSHGLVVSTKGGAVLVDTGWGNEPTEQLLA | + |
| LAEDVRVRRJAPGVWLHVTLA.1 | AYTLQAQEZEJKVTKJAPGVWVHTSYSTY | 6 | 3.53446e-13 | 1.76723e-11 | 1.76723e-11 | 21 | LAEDVRVRRLAPGVWLHVTLA | AYTLQAQTQELQVTKLAPGVWVHTSYSTY | + |
| LAEDVRVRRJAPGVWLHVTLA.1 | AESLPDLKIEKJSENVYLHTSFEZVNGWG | 2 | 1.38138e-06 | 6.9069e-05 | 3.45345e-05 | 21 | LAEDVRVRRLAPGVWLHVTLA | AESLPDLKIEKLSENVYLHTSFEEVNGWG | + |
| LAEDVRVRRJAPGVWLHVTLA.1 | PJADGVYLHTSYKZVEGFGLVDSNGLVVV | -8 | 0.000499237 | 0.0249619 | 0.00832062 | 13 | LAEDVRVRRLAPGVWLHVTLA | PLADGVYLHTSYKQVEGFGLVDSNGLVVV | + |
| LAEDVRVRRJAPGVWLHVTLA.2 | AYTLQAQEZEJKVTKJAPGVWVHTSYSTY | 6 | 3.53446e-13 | 1.76723e-11 | 1.76723e-11 | 21 | LAEDVRVRRLAPGVWLHVTLA | AYTLQAQTQELQVTKLAPGVWVHTSYSTY | + |
| LAEDVRVRRJAPGVWLHVTLA.2 | AESLPDLKIEKJSENVYLHTSFEZVNGWG | 2 | 1.38138e-06 | 6.9069e-05 | 3.45345e-05 | 21 | LAEDVRVRRLAPGVWLHVTLA | AESLPDLKIEKLSENVYLHTSFEEVNGWG | + |
| LAEDVRVRRJAPGVWLHVTLA.2 | PJADGVYLHTSYKZVEGFGLVDSNGLVVV | -8 | 0.000499237 | 0.0249619 | 0.00832062 | 13 | LAEDVRVRRLAPGVWLHVTLA | PLADGVYLHTSYKQVEGFGLVDSNGLVVV | + |
| LVDTGWGPRQTEALLDWARDTL.1 | GVLFPSHGLVVSTKDGAVLVDTGWGNEPTEQLLA | 18 | 2.43858e-11 | 1.21929e-09 | 8.15884e-10 | 16 | LVDTGWGPRQTEALLDWARDTL | GVLFPSHGLVVSTKGGAVLVDTGWGNEPTEQLLA | + |
| LVDTGWGPRQTEALLDWARDTL.1 | GKDAYJIDTPWTEADTEKLVDWIEQQGLTLKASVSTHSHEDRTGGIGYLN | 5 | 3.26353e-11 | 1.63177e-09 | 8.15884e-10 | 22 | LVDTGWGPRQTEALLDWARDTL | GKDAYIIDTPWTEADTEKLVDWIEQQGLTLKASVSTHSHEDRTGGIGYLN | + |
| LVDTGWGPRQTEALLDWARDTL.1 | LVPANGLVVLDNKEAYJIDTPWTAKDTEKLVTWIEER | 16 | 3.93823e-09 | 1.96912e-07 | 6.56372e-08 | 21 | LVDTGWGPRQTEALLDWARDTL | LVPANGLVVLDNKEAYLIDTPWTAKDTEKLVTWIVER | + |
| LVDTGWGPRQTEALLDWARDTL.1 | DQQEAIIFDTPATDQASEELI | 6 | 0.00046873 | 0.0234365 | 0.00585913 | 15 | LVDTGWGPRQTEALLDWARDTL | DQQEAIIFDTPATDQASEELI | + |
| LVDTGWGPRQTEALLDWARDTL.1 | AKJVVPGHGEVGDASLLK | 2 | 0.00121146 | 0.0605731 | 0.0121146 | 16 | LVDTGWGPRQTEALLDWARDTL | AKLVVPGHGEVGDASLLK | + |
| LVDTGWGPRQTEALLDWARDTL.1 | IPTYASELTNALLAQNGKPLA | -2 | 0.00338663 | 0.169332 | 0.0282219 | 20 | LVDTGWGPRQTEALLDWARDTL | IPTYASELTNALLAQNGKPLA | + |
| LVDTGWGPRQTEALLDWARDTL.2 | GVLFPSHGLVVSTKDGAVLVDTGWGNEPTEQLLA | 18 | 2.43858e-11 | 1.21929e-09 | 8.15884e-10 | 16 | LVDTGWGPRQTEALLDWARDTL | GVLFPSHGLVVSTKGGAVLVDTGWGNEPTEQLLA | + |
| LVDTGWGPRQTEALLDWARDTL.2 | GKDAYJIDTPWTEADTEKLVDWIEQQGLTLKASVSTHSHEDRTGGIGYLN | 5 | 3.26353e-11 | 1.63177e-09 | 8.15884e-10 | 22 | LVDTGWGPRQTEALLDWARDTL | GKDAYIIDTPWTEADTEKLVDWIEQQGLTLKASVSTHSHEDRTGGIGYLN | + |
| LVDTGWGPRQTEALLDWARDTL.2 | LVPANGLVVLDNKEAYJIDTPWTAKDTEKLVTWIEER | 16 | 3.93823e-09 | 1.96912e-07 | 6.56372e-08 | 21 | LVDTGWGPRQTEALLDWARDTL | LVPANGLVVLDNKEAYLIDTPWTAKDTEKLVTWIVER | + |
| LVDTGWGPRQTEALLDWARDTL.2 | DQQEAIIFDTPATDQASEELI | 6 | 0.00046873 | 0.0234365 | 0.00585913 | 15 | LVDTGWGPRQTEALLDWARDTL | DQQEAIIFDTPATDQASEELI | + |
| LVDTGWGPRQTEALLDWARDTL.2 | AKJVVPGHGEVGDASLLK | 2 | 0.00121146 | 0.0605731 | 0.0121146 | 16 | LVDTGWGPRQTEALLDWARDTL | AKLVVPGHGEVGDASLLK | + |
| LVDTGWGPRQTEALLDWARDTL.2 | IPTYASELTNALLAQNGKPLA | -2 | 0.00338663 | 0.169332 | 0.0282219 | 20 | LVDTGWGPRQTEALLDWARDTL | IPTYASELTNALLAQNGKPLA | + |
| MKILILAAMLFAAQP.1 | MKKLJLLLLLLLILL | 0 | 0.000828488 | 0.0414244 | 0.0265749 | 15 | MKILILAAMLFAAQP | MKRLLLLLLLLLILL | + |
| MKILILAAMLFAAQP.1 | MKYJJLJJLLL | 0 | 0.001063 | 0.0531499 | 0.0265749 | 11 | MKILILAAMLFAAQP | MKYILLLLLLL | + |
| MKILILAAMLFAAQP.2 | MKKLJLLLLLLLILL | 0 | 0.000828488 | 0.0414244 | 0.0265749 | 15 | MKILILAAMLFAAQP | MKRLLLLLLLLLILL | + |
| MKILILAAMLFAAQP.2 | MKYJJLJJLLL | 0 | 0.001063 | 0.0531499 | 0.0265749 | 11 | MKILILAAMLFAAQP | MKYILLLLLLL | + |
| MSLLRGAVFALLVGVAGCRVSSAAPQPTA.1 | MHPPLRPHPGWKRLGPALLALLLGVFGAR | 10 | 1.52664e-07 | 7.63318e-06 | 7.63318e-06 | 19 | MSLLRGAVFALLVGVAACRVSSAAAQATA | MHPPLRPHPLFKRLGLALLALLLGAAGAR | + |
| MSLLRGAVFALLVGVAGCRVSSAAPQPTA.1 | MKKLJLLLLLLLILL | 0 | 0.000556821 | 0.027841 | 0.0139205 | 15 | MSLLRGAVFALLVGVAACRVSSAAAQATA | MKRLLLLLLLLLILL | + |
| MSLLRGAVFALLVGVAGCRVSSAAPQPTA.1 | MFKITLALLSAVIGF | -3 | 0.00127534 | 0.0637672 | 0.0212557 | 15 | MSLLRGAVFALLVGVAACRVSSAAAQATA | MFKITLALLSAVIGF | + |
| MSLLRGAVFALLVGVAGCRVSSAAPQPTA.2 | MHPPLRPHPGWKRLGPALLALLLGVFGAR | 10 | 1.52664e-07 | 7.63318e-06 | 7.63318e-06 | 19 | MSLLRGAVFALLVGVAACRVSSAAAQATA | MHPPLRPHPLFKRLGLALLALLLGAAGAR | + |
| MSLLRGAVFALLVGVAGCRVSSAAPQPTA.2 | MKKLJLLLLLLLILL | 0 | 0.000556821 | 0.027841 | 0.0139205 | 15 | MSLLRGAVFALLVGVAACRVSSAAAQATA | MKRLLLLLLLLLILL | + |
| MSLLRGAVFALLVGVAGCRVSSAAPQPTA.2 | MFKITLALLSAVIGF | -3 | 0.00127534 | 0.0637672 | 0.0212557 | 15 | MSLLRGAVFALLVGVAACRVSSAAAQATA | MFKITLALLSAVIGF | + |
| RARYPEARVVVPGHGAPGGPELLDHTEAL.1 | AKJVVPGHGEVGDASLLK | -6 | 2.63984e-12 | 1.31992e-10 | 1.31992e-10 | 18 | RARYPEARVVVPGHGAPGGPELLDHTEAL | AKLVVPGHGEVGDASLLK | + |
| RARYPEARVVVPGHGAPGGPELLDHTEAL.1 | ADADLDAWPASIAKVQARYPDAKIVVPGHGK | 15 | 1.67916e-09 | 8.39578e-08 | 4.19789e-08 | 16 | RARYPEARVVVPGHGAPGGPELLDHTEAL | ADADLDAWPASIAKVQARYPDAKIVVPGHGK | + |
| RARYPEARVVVPGHGAPGGPELLDHTEAL.1 | VGDASLLDHTIELAE | -16 | 2.09504e-08 | 1.04752e-06 | 3.49173e-07 | 13 | RARYPEARVVVPGHGAPGGPELLDHTEAL | VGDASLLDHTIELAE | + |
| RARYPEARVVVPGHGAPGGPELLDHTEAL.2 | AKJVVPGHGEVGDASLLK | -6 | 2.63984e-12 | 1.31992e-10 | 1.31992e-10 | 18 | RARYPEARVVVPGHGAPGGPELLDHTEAL | AKLVVPGHGEVGDASLLK | + |
| RARYPEARVVVPGHGAPGGPELLDHTEAL.2 | ADADLDAWPASIAKVQARYPDAKIVVPGHGK | 15 | 1.67916e-09 | 8.39578e-08 | 4.19789e-08 | 16 | RARYPEARVVVPGHGAPGGPELLDHTEAL | ADADLDAWPASIAKVQARYPDAKIVVPGHGK | + |
| RARYPEARVVVPGHGAPGGPELLDHTEAL.2 | VGDASLLDHTIELAE | -16 | 2.09504e-08 | 1.04752e-06 | 3.49173e-07 | 13 | RARYPEARVVVPGHGAPGGPELLDHTEAL | VGDASLLDHTIELAE | + |
| TARLAAAQGNPVPSQ.1 | IPTYASELTNALLAQNGKPLA | 8 | 3.22488e-05 | 0.00161244 | 0.00161244 | 13 | TARLAAAQGNPVPSQ | IPTYASELTNALLAQNGKPLA | + |
| TARLAAAQGNPVPSQ.2 | IPTYASELTNALLAQNGKPLA | 8 | 3.22488e-05 | 0.00161244 | 0.00161244 | 13 | TARLAAAQGNPVPSQ | IPTYASELTNALLAQNGKPLA | + |
| YPANGLJVEDGDGSJ.1 | GVLFPSHGLVVSTKDGAVLVDTGWGNEPTEQLLA | 3 | 2.81149e-08 | 1.40575e-06 | 1.40575e-06 | 15 | YPANGLIVEDGDGSL | GVLFPSHGLVVSTKGGAVLVDTGWGNEPTEQLLA | + |
| YPANGLJVEDGDGSJ.1 | LVPANGLVVLDNKEAYJIDTPWTAKDTEKLVTWIEER | 1 | 5.07363e-07 | 2.53682e-05 | 1.26841e-05 | 15 | YPANGLIVEDGDGSL | LVPANGLVVLDNKEAYLIDTPWTAKDTEKLVTWIVER | + |
| YPANGLJVEDGDGSJ.1 | PJADGVYLHTSYKZVEGFGLVDSNGLVVV | 20 | 0.000428943 | 0.0214471 | 0.00714905 | 9 | YPANGLIVEDGDGSL | PLADGVYLHTSYKQVEGFGLVDSNGLVVV | + |
| YPANGLJVEDGDGSJ.2 | GVLFPSHGLVVSTKDGAVLVDTGWGNEPTEQLLA | 3 | 2.81149e-08 | 1.40575e-06 | 1.40575e-06 | 15 | YPANGLIVEDGDGSL | GVLFPSHGLVVSTKGGAVLVDTGWGNEPTEQLLA | + |
| YPANGLJVEDGDGSJ.2 | LVPANGLVVLDNKEAYJIDTPWTAKDTEKLVTWIEER | 1 | 5.07363e-07 | 2.53682e-05 | 1.26841e-05 | 15 | YPANGLIVEDGDGSL | LVPANGLVVLDNKEAYLIDTPWTAKDTEKLVTWIVER | + |
| YPANGLJVEDGDGSJ.2 | PJADGVYLHTSYKZVEGFGLVDSNGLVVV | 20 | 0.000428943 | 0.0214471 | 0.00714905 | 9 | YPANGLIVEDGDGSL | PLADGVYLHTSYKQVEGFGLVDSNGLVVV | + |
| AKLAKEQGYEVPBPTLDELTT | ELLKENGKPLAKHTF | -1 | 2.14368e-09 | 1.07184e-07 | 1.07184e-07 | 15 | AKLAKEQGYEVPNPTLDELTT | ELLKENGKPLAKHTF | + |
| AKLAKEQGYEVPBPTLDELTT | ENGKGTPDITFATDT | -5 | 1.6095e-05 | 0.000804751 | 0.000355685 | 15 | AKLAKEQGYEVPNPTLDELTT | ENGKGTPDVTFATDT | + |
| AKLAKEQGYEVPBPTLDELTT | ELLKKDGKVQATNSF | -1 | 2.13411e-05 | 0.00106706 | 0.000355685 | 15 | AKLAKEQGYEVPNPTLDELTT | ELLKKDGKVQATNSF | + |
| AKLAKEQGYEVPBPTLDELTT | AKSKNLPIPDHGFKD | -3 | 0.00299758 | 0.149879 | 0.0374698 | 15 | AKLAKEQGYEVPNPTLDELTT | AKSKNLPIPAHGFRD | + |
| ANGLJVEDGDESLLVDTAWGARQTAALLAW | LVPSNGLVVVDGKEAYJIDTPWSDKDTEKLVDWI | 3 | 2.6489e-18 | 1.32445e-16 | 1.32445e-16 | 30 | ANGLIVEDGDESLLVDTAWGARQTAALLAW | LVPSNGLVVVDGKEAYIIDTPWSDKDTEKLVDWI | + |
| ANGLJVEDGDESLLVDTAWGARQTAALLAW | EGVYVHTSYEEVEGWGLVPANGLVVLDNKEAYJIDTPWTAK | 19 | 3.54327e-08 | 1.77164e-06 | 8.85818e-07 | 22 | ANGLIVEDGDESLLVDTAWGARQTAALLAW | EGVYVHTSYEEVEGWGLVPANGLVVLDNKEAYLIDTPWTAK | + |
| ANGLJVEDGDESLLVDTAWGARQTAALLAW | TKEGAVLIDTGWGNE | -7 | 2.25253e-07 | 1.12626e-05 | 3.75422e-06 | 15 | ANGLIVEDGDESLLVDTAWGARQTAALLAW | TKEGAVLIDTGWGNE | + |
| ANGLJVEDGDESLLVDTAWGARQTAALLAW | IVQTAPHKAVLIDTPWDNSDVDTLFSWLE | -3 | 0.001209 | 0.0604502 | 0.0151126 | 27 | ANGLIVEDGDESLLVDTAWGARQTAALLAW | IVQTAPHKAVLIDTPWDNSDVDTLFSWLE | + |
| ANGLJVEDGDESLLVDTAWGARQTAALLAW | DASLLKHTLELAVKG | -9 | 0.00167974 | 0.0839872 | 0.0167974 | 15 | ANGLIVEDGDESLLVDTAWGARQTAALLAW | DASLLKHTLELAVKG | + |
| ANGLJVEDGDESLLVDTAWGARQTAALLAW | PGHGKVGDGELLDHT | -2 | 0.00397009 | 0.198504 | 0.0330841 | 15 | ANGLIVEDGDESLLVDTAWGARQTAALLAW | PGHGKVGDGELLDHT | + |
| ARDTJGLPVRAAVVTHFHDDRTGGVPVLRARGIPV | ISTHFHEDRTGGJGYLNSHGIPTYASELT | -12 | 1.19673e-15 | 5.98365e-14 | 5.98365e-14 | 23 | ARDTLGLPVRAAVVTHFHDDRTGGVPVLRARGIPV | ISTHFHEDRTGGLGYLNSHGIPTYASELT | + |
| ARDTJGLPVRAAVVTHFHDDRTGGVPVLRARGIPV | STHFHEDSTGGIEWLNSRSIPTYASELTN | -13 | 3.20891e-12 | 1.60446e-10 | 8.02228e-11 | 22 | ARDTLGLPVRAAVVTHFHDDRTGGVPVLRARGIPV | STHFHEDSTGGIEWLNSRSIPTYASELTN | + |
| DVRVRRJAPGVWLHVTTAGFD | TSVPAPRPAAEPFTSITEDLRIQPLAPGVWRLVALSGEEWG | 18 | 7.84301e-11 | 3.9215e-09 | 3.9215e-09 | 21 | DVRVRRIAPGVWLHVTTAGFD | TSVPAPRPAAEPFTSITEDLRIQPLAPGVWRLVALSGEEWG | + |
| DVRVRRJAPGVWLHVTTAGFD | EYESPELEITPJSDNVYLHTSYKZVEGFG | 5 | 1.70536e-08 | 8.52678e-07 | 4.26339e-07 | 21 | DVRVRRIAPGVWLHVTTAGFD | EYESPELEITPLSDNVYLHTSYKQVEGFG | + |
| DVRVRRJAPGVWLHVTTAGFD | PAYTLLAQEQZJKVTKIAPBVYVHTSYSTYQGVLVPSHGLV | 10 | 3.2265e-08 | 1.61325e-06 | 5.3775e-07 | 21 | DVRVRRIAPGVWLHVTTAGFD | PAYTLLAQEQELKVTKIAPNVWVHTSYSTYQGVLVPSHGLV | + |
| EVFYPGPGHSPDNJVVWLPZYKILFGGCLVKSLQA | IEVFYPGAGHTKDNLVVWLPKEKILFGGCLVKS | 1 | 4.74354e-41 | 2.37177e-39 | 2.37177e-39 | 32 | EVFYPGPGHSPDNIVVWLPQYKILFGGCLVKSLQA | IEVFYPGAGHTKDNLVVWLPKEKILFGGCLVKS | + |
| EVFYPGPGHSPDNJVVWLPZYKILFGGCLVKSLQA | YWLVKNKIEVFYPGAGHTPDNVVVWLPEEKILFGGCFVKSL | 8 | 1.66897e-39 | 8.34487e-38 | 4.17243e-38 | 33 | EVFYPGPGHSPDNIVVWLPQYKILFGGCLVKSLQA | FWLVKGKIEVFYPGAGHTPDNVVVWLPESKILFGGCFVKSL | + |
| FPSNGLIVETGDGLVLIDTAWGEEQTEZL | LVPSNGLVVVDGKEAYJIDTPWSDKDTEKLVDWI | 1 | 6.06099e-18 | 3.03049e-16 | 3.03049e-16 | 29 | FPSNGLIVETGDGLVLIDTAWGELQTEEL | LVPSNGLVVVDGKEAYIIDTPWSDKDTEKLVDWI | + |
| FPSNGLIVETGDGLVLIDTAWGEEQTEZL | TKEGAVLIDTGWGNE | -9 | 1.05185e-11 | 5.25927e-10 | 2.62963e-10 | 15 | FPSNGLIVETGDGLVLIDTAWGELQTEEL | TKEGAVLIDTGWGNE | + |
| FPSNGLIVETGDGLVLIDTAWGEEQTEZL | EGVYVHTSYEEVEGWGLVPANGLVVLDNKEAYJIDTPWTAK | 17 | 1.31466e-09 | 6.57331e-08 | 2.1911e-08 | 24 | FPSNGLIVETGDGLVLIDTAWGELQTEEL | EGVYVHTSYEEVEGWGLVPANGLVVLDNKEAYLIDTPWTAK | + |
| FPSNGLIVETGDGLVLIDTAWGEEQTEZL | IVQTAPHKAVLIDTPWDNSDVDTLFSWLE | -5 | 0.000674643 | 0.0337322 | 0.00843304 | 24 | FPSNGLIVETGDGLVLIDTAWGELQTEEL | IVQTAPHKAVLIDTPWDNSDVDTLFSWLE | + |
| GPLELFFPGAGHSPDNLVVWH | YWLVKNKIEVFYPGAGHTPDNVVVWLPEEKILFGGCFVKSL | 5 | 2.61948e-18 | 1.30974e-16 | 1.30974e-16 | 21 | GPLELFFPGAGHSPDNLVVWH | FWLVKGKIEVFYPGAGHTPDNVVVWLPESKILFGGCFVKSL | + |
| GPLELFFPGAGHSPDNLVVWH | IEVFYPGAGHTKDNLVVWLPKEKILFGGCLVKS | -2 | 9.30129e-18 | 4.65064e-16 | 2.32532e-16 | 19 | GPLELFFPGAGHSPDNLVVWH | IEVFYPGAGHTKDNLVVWLPKEKILFGGCLVKS | + |
| HGLEDTARLATEQGNPVPTQR | ELLKENGKPLAKHTF | -7 | 1.63174e-05 | 0.000815868 | 0.000815868 | 14 | HGLEDTARLATEQGNPVPTQR | ELLKENGKPLAKHTF | + |
| IVTHAHDDRIGGIDVLKKRGIPVYSTPLT | ISTHFHEDRTGGJGYLNSHGIPTYASELT | 0 | 2.33455e-23 | 1.16728e-21 | 1.16728e-21 | 29 | IVTHAHDDRIGGIDVLKKRGIPVYSTPLT | ISTHFHEDRTGGLGYLNSHGIPTYASELT | + |
| IVTHAHDDRIGGIDVLKKRGIPVYSTPLT | STHFHEDSTGGIEWLNSRSIPTYASELTN | -1 | 8.75281e-20 | 4.3764e-18 | 2.1882e-18 | 28 | IVTHAHDDRIGGIDVLKKRGIPVYSTPLT | STHFHEDSTGGIEWLNSRSIPTYASELTN | + |
| IVTHAHDDRIGGIDVLKKRGIPVYSTPLT | ELLKENGKPLAKHTF | -13 | 0.000287521 | 0.014376 | 0.00479201 | 15 | IVTHAHDDRIGGIDVLKKRGIPVYSTPLT | ELLKENGKPLAKHTF | + |
| KSLGNTADADLKEWPKSIKRVQQRYPKAK | SKSLGYTGDABLSAWPNSVEKVKAKYPDA | 1 | 3.42198e-28 | 1.71099e-26 | 1.71099e-26 | 28 | KSLGNTADADLKEWPKSIKRVQQRYPKAK | SKSLGYTGDADLSAWPNSVEKVKAKYPDA | + |
| KSLGNTADADLKEWPKSIKRVQQRYPKAK | GLGNLGDANIEAWPKSIKKVKSKYPKAKLVVPGHGKVG | -1 | 1.06245e-27 | 5.31223e-26 | 2.65612e-26 | 28 | KSLGNTADADLKEWPKSIKRVQQRYPKAK | GLGNLGDANIEAWPKSIKKVKSKYPKAKLVVPGHGKVG | + |
| KSLGNTADADLKEWPKSIKRVQQRYPKAK | LGNLGDAVLEAWPTSAKKLISKYGEAKJV | -2 | 5.00826e-17 | 2.50413e-15 | 8.3471e-16 | 27 | KSLGNTADADLKEWPKSIKRVQQRYPKAK | LGNLGDAVLEAWPTSAKKLISKYGAAKIV | + |
| LKFGNTKV | DSLVLKVGNEK | 4 | 0.000131412 | 0.0065706 | 0.0065706 | 7 | LKFGNTKV | DSLVLKVGNEK | + |
| LLLVPAYALLAQSQZ | PAYTLLAQEQZJKVTKIAPBVYVHTSYSTYQGVLVPSHGLV | -4 | 2.51322e-08 | 1.25661e-06 | 1.25661e-06 | 11 | LLLVPAYALLAQSQE | PAYTLLAQEQELKVTKIAPNVWVHTSYSTYQGVLVPSHGLV | + |
| MFKLJSKLLVY | MKKLJLLLLJL | 0 | 0.000204892 | 0.0102446 | 0.0102446 | 11 | MFKLLSKLLVY | MKSLILLLLIL | + |
| MFKLJSKLLVY | MFKJTLALLSAVIGF | 0 | 0.00110727 | 0.0553633 | 0.0276816 | 11 | MFKLLSKLLVY | MFKITLALLSAVIGF | + |
| MKFHKCWIKKDVSKLFLTVEVVLTSFSED | DASLLKHTLELAVKG | -10 | 0.000237808 | 0.0118904 | 0.0118904 | 15 | MKFHKCWIKKDVSKLFLTVEVVLTSGSED | DASLLKHTLELAVKG | + |
| MKFLFAALFVVMFCGLARASD | MKKLJLLLLJL | 0 | 1.54512e-05 | 0.000772558 | 0.000772558 | 11 | MKSLFAALFVVMFCGLARASD | MKSLILLLLIL | + |
| MKFLFAALFVVMFCGLARASD | MKYIJLLLLLI | 0 | 0.000388025 | 0.0194013 | 0.00970063 | 11 | MKSLFAALFVVMFCGLARASD | MKYIILLLLLI | + |
| MKRJJLLLLJL | MKKLJLLLLJL | 0 | 3.37525e-12 | 1.68762e-10 | 8.43812e-11 | 11 | MKRILLLLLLL | MKSLILLLLIL | + |
| MKRJJLLLLJL | MKYIJLLLLLI | 0 | 3.37525e-12 | 1.68762e-10 | 8.43812e-11 | 11 | MKRILLLLLLL | MKYIILLLLLI | + |
| MKRJJLLLLJL | MKKLLVFFJFLFCSI | 0 | 3.85786e-09 | 1.92893e-07 | 6.42976e-08 | 11 | MKRILLLLLLL | MKKLLVFFIFLFCSI | + |
| MKRJJLLLLJL | MFKJTLALLSAVIGF | 0 | 0.00153215 | 0.0766074 | 0.0191519 | 11 | MKRILLLLLLL | MFKITLALLSAVIGF | + |
| MMKFRRIGGFLFFLLIGSFTQSAQENIPN | MKYIJLLLLLI | -1 | 0.000354301 | 0.0177151 | 0.0166407 | 11 | MMKFRRIGGFLFFLLIGSFTQSAQANIPN | MKYIILLLLLI | + |
| MMKFRRIGGFLFFLLIGSFTQSAQENIPN | MKKLLVFFJFLFCSI | -3 | 0.000873167 | 0.0436584 | 0.0166407 | 15 | MMKFRRIGGFLFFLLIGSFTQSAQANIPN | MKKLLVFFIFLFCSI | + |
| MMKFRRIGGFLFFLLIGSFTQSAQENIPN | MKIRIKNIVAJLLSFVILSCSSZKTDNFK | -1 | 0.000998443 | 0.0499221 | 0.0166407 | 28 | MMKFRRIGGFLFFLLIGSFTQSAQANIPN | MKIRIKNIVALLLSFVILSCSSQKTTNFK | + |
| MMKFRRIGGFLFFLLIGSFTQSAQENIPN | MKKLJLLLLJL | -1 | 0.00133533 | 0.0667664 | 0.0166916 | 11 | MMKFRRIGGFLFFLLIGSFTQSAQANIPN | MKSLILLLLIL | + |
| MSIQHFRVALI | MSIQHFRVALIPFFAAFCLP | 0 | 1.7043e-19 | 8.5215e-18 | 8.5215e-18 | 11 | MSIQHFRVALI | MSIQHFRVALIPFFAAFCLP | + |
| VEITKJAPGVWVHTSYYTYPG | PAYTLLAQEQZJKVTKIAPBVYVHTSYSTYQGVLVPSHGLV | 11 | 2.35806e-19 | 1.17903e-17 | 1.17903e-17 | 21 | VEITKIAPGVWVHTSYYTYPG | PAYTLLAQEQELKVTKIAPNVWVHTSYSTYQGVLVPSHGLV | + |
| VEITKJAPGVWVHTSYYTYPG | EYESPELEITPJSDNVYLHTSYKZVEGFG | 6 | 1.89533e-16 | 9.47663e-15 | 4.73832e-15 | 21 | VEITKIAPGVWVHTSYYTYPG | EYESPELEITPLSDNVYLHTSYKQVEGFG | + |
| VEITKJAPGVWVHTSYYTYPG | PLKIEAJSSKVYLVKSYKEILNVYESDTPKIIDANALJYI | 1 | 9.94747e-07 | 4.97373e-05 | 1.65791e-05 | 21 | VEITKIAPGVWVHTSYYTYPG | PLKIEALSSKVYLVTSYKTILNVYESDTPKIIDANALLYI | + |
| VEITKJAPGVWVHTSYYTYPG | EGVYVHTSYEEVEGWGLVPANGLVVLDNKEAYJIDTPWTAK | -7 | 3.71853e-06 | 0.000185927 | 4.64817e-05 | 14 | VEITKIAPGVWVHTSYYTYPG | EGVYVHTSYEEVEGWGLVPANGLVVLDNKEAYLIDTPWTAK | + |
| VEITKJAPGVWVHTSYYTYPG | TSVPAPRPAAEPFTSITEDLRIQPLAPGVWRLVALSGEEWG | 19 | 0.00205666 | 0.102833 | 0.0205666 | 21 | VEITKIAPGVWVHTSYYTYPG | TSVPAPRPAAEPFTSITEDLRIQPLAPGVWRLVALSGEEWG | + |
| VLFGGCFVKDASAKSLGNVADADVAAWPASJERIRQRYPEARV | SKSLGYTGDABLSAWPNSVEKVKAKYPDA | -12 | 2.11476e-21 | 1.05738e-19 | 1.05738e-19 | 29 | VLFGGCFVKDAAAKSLGNVADADVAAWPASLERIRQRYPEARV | SKSLGYTGDADLSAWPNSVEKVKAKYPDA | + |
| VLFGGCFVKDASAKSLGNVADADVAAWPASJERIRQRYPEARV | GLGNLGDANIEAWPKSIKKVKSKYPKAKLVVPGHGKVG | -14 | 1.28519e-13 | 6.42593e-12 | 3.21297e-12 | 29 | VLFGGCFVKDAAAKSLGNVADADVAAWPASLERIRQRYPEARV | GLGNLGDANIEAWPKSIKKVKSKYPKAKLVVPGHGKVG | + |
| VLFGGCFVKDASAKSLGNVADADVAAWPASJERIRQRYPEARV | LGNLGDAVLEAWPTSAKKLISKYGEAKJV | -15 | 2.31868e-13 | 1.15934e-11 | 3.86447e-12 | 28 | VLFGGCFVKDAAAKSLGNVADADVAAWPASLERIRQRYPEARV | LGNLGDAVLEAWPTSAKKLISKYGAAKIV | + |
| VVPGHGEWGGKELLSHTLELL | PGHGKVGDGELLDHT | -2 | 1.3708e-16 | 6.85399e-15 | 6.85399e-15 | 15 | VVPGHGEWGGKELLSHTLELL | PGHGKVGDGELLDHT | + |
| VVPGHGEWGGKELLSHTLELL | DASLLKHTLELAVKG | -9 | 2.06679e-07 | 1.0334e-05 | 5.16698e-06 | 12 | VVPGHGEWGGKELLSHTLELL | DASLLKHTLELAVKG | + |
| VVPGHGEWGGKELLSHTLELL | ELLDYTIDLF | -11 | 1.75589e-05 | 0.000877947 | 0.000292649 | 10 | VVPGHGEWGGKELLSHTLELL | ELLDYTIDLF | + |
| VVPGHGEWGGKELLSHTLELL | GLGNLGDANIEAWPKSIKKVKSKYPKAKLVVPGHGKVG | 29 | 0.00013986 | 0.00699302 | 0.00174825 | 9 | VVPGHGEWGGKELLSHTLELL | GLGNLGDANIEAWPKSIKKVKSKYPKAKLVVPGHGKVG | + |
| VVPGHGLPGGPELLDHTEALL | PGHGKVGDGELLDHT | -2 | 2.85786e-13 | 1.42893e-11 | 1.42893e-11 | 15 | VVPGHGLPGGPELLDHTEALL | PGHGKVGDGELLDHT | + |
| VVPGHGLPGGPELLDHTEALL | ELLDYTIDLF | -11 | 2.4741e-05 | 0.00123705 | 0.000618525 | 10 | VVPGHGLPGGPELLDHTEALL | ELLDYTIDLF | + |
| VVPGHGLPGGPELLDHTEALL | DASLLKHTLELAVKG | -9 | 0.000207808 | 0.0103904 | 0.00346347 | 12 | VVPGHGLPGGPELLDHTEALL | DASLLKHTLELAVKG | + |
| VVPGHGLPGGPELLDHTEALL | GLGNLGDANIEAWPKSIKKVKSKYPKAKLVVPGHGKVG | 29 | 0.00322781 | 0.16139 | 0.0403476 | 9 | VVPGHGLPGGPELLDHTEALL | GLGNLGDANIEAWPKSIKKVKSKYPKAKLVVPGHGKVG | + |
| YALPLSNELAPERGLPPAEFL | ELLKENGKPLAKHTF | -7 | 6.2689e-05 | 0.00313445 | 0.00313445 | 14 | YALPLTNQLAASRGLPPAEFL | ELLKENGKPLAKHTF | + |
| YASPSTRRLAEAEGNEIPTHSLEGLSSSG | ELLKENGKPLAKHTF | -7 | 8.57583e-06 | 0.000428791 | 0.000428791 | 15 | YASPSTRRLAEAEGNEIPTHSLEGLSSSG | ELLKENGKPLAKHTF | + |
| YASPSTRRLAEAEGNEIPTHSLEGLSSSG | ELLKKDGKVQATNSF | -7 | 0.000520299 | 0.026015 | 0.0130075 | 15 | YASPSTRRLAEAEGNEIPTHSLEGLSSSG | ELLKKDGKVQATNSF | + |
| YGLVDANGLVVLDGQGAYIIDTPWSZQDT | LVPSNGLVVVDGKEAYJIDTPWSDKDTEKLVDWI | -2 | 7.58285e-25 | 3.79142e-23 | 3.79142e-23 | 27 | YGLVDANGLVVLDGQGAYIIDTPWSERDT | LVPSNGLVVVDGKEAYIIDTPWSDKDTEKLVDWI | + |
| YGLVDANGLVVLDGQGAYIIDTPWSZQDT | EGVYVHTSYEEVEGWGLVPANGLVVLDNKEAYJIDTPWTAK | 14 | 2.87322e-21 | 1.43661e-19 | 7.18304e-20 | 27 | YGLVDANGLVVLDGQGAYIIDTPWSERDT | EGVYVHTSYEEVEGWGLVPANGLVVLDNKEAYLIDTPWTAK | + |
| YGLVDANGLVVLDGQGAYIIDTPWSZQDT | TKEGAVLIDTGWGNE | -12 | 4.05999e-06 | 0.000202999 | 6.76664e-05 | 15 | YGLVDANGLVVLDGQGAYIIDTPWSERDT | TKEGAVLIDTGWGNE | + |
| YGLVDANGLVVLDGQGAYIIDTPWSZQDT | IVQTAPHKAVLIDTPWDNSDVDTLFSWLE | -8 | 1.64586e-05 | 0.000822929 | 0.000205732 | 21 | YGLVDANGLVVLDGQGAYIIDTPWSERDT | IVQTAPHKAVLIDTPWDNSDVDTLFSWLE | + |
| AEAZGNPVPTHSLAG | KKDGKVQATHSFSGVNFWLVKNKIEVFYP | -1 | 0.000448492 | 0.0224246 | 0.0224246 | 14 | AEAEGNPVPTHSLAG | KKDGKVQATHSFSGVNFWLVKNKIEVFYP | + |
| AEAZGNPVPTHSLAG | MFSRILILLAGFJCCPPYAHSAESSGQLTITPLSSGALVVA | 21 | 0.00188136 | 0.0940678 | 0.0470339 | 15 | AEAEGNPVPTHSLAG | MFSRILILLAGFLCCVPLAHSAESSGQLTITPLLSGALVVA | + |
| AEDTJGLPVRAAVVT | FDTPWTBEPTEQLLAWVKDNLKAPVKAFVPTHWHDDCLGGL | 16 | 5.03506e-07 | 2.51753e-05 | 2.51753e-05 | 15 | AEDTLGLPVRAAVVT | FDTPWTDEPTEQLLAWVKDNLKAPVKAFVPTHWHDDCLGGL | + |
| AEDTJGLPVRAAVVT | MPNRALWQRIAILLLMPILSLPALAANLT | 14 | 0.000505879 | 0.0252939 | 0.012647 | 15 | AEDTLGLPVRAAVVT | MPNRALWQRIAILLLMPILSLPALAANLT | + |
| AEDTJGLPVRAAVVT | AYJIDTPWTEKDTEKLVDWIEAQGLTLKASISTHSHZDRTGGIGYLNSKG | 18 | 0.00164955 | 0.0824774 | 0.0274925 | 15 | AEDTLGLPVRAAVVT | AYIIDTPWTEKDTEKLVDWIEAQGLTLKASISTHSHEDRTGGIGYLNSIG | + |
| DLAPSNKLLPAEFDLSFDNNNKSSDISP | ASNKSIQPTAEASAD | -3 | 0.000501119 | 0.025056 | 0.025056 | 15 | DLAPSNKLLPAEFDLSFDSNNRSSDISP | ASNKSIQPTAEASAD | + |
| DVRVRRJAPGVWLHTTTQGFD | PAYTLQAQEQEJQVTKJAPGVWVHTSYSTYNGVLVPSHGLVVSTKEGAVL | 10 | 1.20832e-13 | 6.04159e-12 | 6.04159e-12 | 21 | DVRVRRIAPGVWLHTTTQGFD | PAYTLLAQEQELQVTKLAPGVWVHTSYSTYGGVLVPSHGLVVSTKGGAVL | + |
| DVRVRRJAPGVWLHTTTQGFD | KJADGVYLHTSYKEVEGFGLVSSNGLVVV | -5 | 1.80064e-05 | 0.000900322 | 0.000376413 | 16 | DVRVRRIAPGVWLHTTTQGFD | KLADGVYLHTSYKEVEGFGLVSSNGLVVV | + |
| DVRVRRJAPGVWLHTTTQGFD | IEKJSENVYLHTSFKZTNGWG | -3 | 2.25848e-05 | 0.00112924 | 0.000376413 | 18 | DVRVRRIAPGVWLHTTTQGFD | IEKLSENVYLHTSFKETNGWG | + |
| EVFYPGPGHTMDNIVVWLPZQKILFGGCLVKSLQAKDLGNTADADLNEWP | IEVFYPGAGHTKDNIVVWLPKQKILFGGCLVKS | 1 | 8.82818e-44 | 4.41409e-42 | 4.41409e-42 | 32 | EVFYPGAGHTMDNIVVWLPQQKILFGGCLVKSLQAKDLGNTADADLNSWP | IEVFYPGAGHTKDNIVVWLPKQKILFGGCLVKS | + |
| EVFYPGPGHTMDNIVVWLPZQKILFGGCLVKSLQAKDLGNTADADLNEWP | GPGHTKDNVVVWLPKEKILFGGCLVKSLGAGNLGL | -5 | 1.17285e-35 | 5.86423e-34 | 2.93211e-34 | 35 | EVFYPGAGHTMDNIVVWLPQQKILFGGCLVKSLQAKDLGNTADADLNSWP | GPGHTKDNVVVWLPKEKILFGGCLVKSLGAGNLGL | + |
| EVFYPGPGHTMDNIVVWLPZQKILFGGCLVKSLQAKDLGNTADADLNEWP | SLGNTGDADLEAWPASIAKVQAR | -36 | 3.42906e-08 | 1.71453e-06 | 5.7151e-07 | 14 | EVFYPGAGHTMDNIVVWLPQQKILFGGCLVKSLQAKDLGNTADADLNSWP | SLGNTGDADLEAWPASIAKVQAR | + |
| EWIKTKLKKPVKKAI | FDTPWTBEPTEQLLAWVKDNLKAPVKAFVPTHWHDDCLGGL | 14 | 1.93367e-09 | 9.66836e-08 | 9.66836e-08 | 15 | EWIKTKLKKPVKKAI | FDTPWTDEPTEQLLAWVKDNLKAPVKAFVPTHWHDDCLGGL | + |
| FPSNGLIVETGKGLVLIDTAWGEEQTEZL | PCNGLVVIDNKEAYJIDTPWNAKDTE | -1 | 1.16242e-14 | 5.81209e-13 | 5.81209e-13 | 26 | FPSNGLIVETGKGLVLIDTAWGEEQTEEL | PCNGLVVIDNKEAYLIDTPWNAKDTE | + |
| FPSNGLIVETGKGLVLIDTAWGEEQTEZL | PAYTLQAQEQEJQVTKJAPGVWVHTSYSTYNGVLVPSHGLVVSTKEGAVL | 34 | 7.43403e-08 | 3.71702e-06 | 1.85851e-06 | 16 | FPSNGLIVETGKGLVLIDTAWGEEQTEEL | PAYTLLAQEQELQVTKLAPGVWVHTSYSTYGGVLVPSHGLVVSTKGGAVL | + |
| FPSNGLIVETGKGLVLIDTAWGEEQTEZL | FDTPWTBEPTEQLLAWVKDNLKAPVKAFVPTHWHDDCLGGL | -16 | 4.62195e-05 | 0.00231098 | 0.000770325 | 13 | FPSNGLIVETGKGLVLIDTAWGEEQTEEL | FDTPWTDEPTEQLLAWVKDNLKAPVKAFVPTHWHDDCLGGL | + |
| FPSNGLIVETGKGLVLIDTAWGEEQTEZL | AYJIDTPWTEKDTEKLVDWIEAQGLTLKASISTHSHZDRTGGIGYLNSKG | -13 | 0.000132628 | 0.00663139 | 0.00165785 | 16 | FPSNGLIVETGKGLVLIDTAWGEEQTEEL | AYIIDTPWTEKDTEKLVDWIEAQGLTLKASISTHSHEDRTGGIGYLNSIG | + |
| HFHDDRTGGVPVLRARGIPVYALPDTARL | RGYKJKASISTHFHEDSTGGJEYLNSHSIPTYASELTNELL | 11 | 1.35918e-19 | 6.79589e-18 | 6.79589e-18 | 29 | HFHDDRTGGVPVLRARGIPVYALPDTARL | RGYKIKASISTHFHEDSTGGLEYLNSHSIPTYASELTNELL | + |
| HFHDDRTGGVPVLRARGIPVYALPDTARL | AYJIDTPWTEKDTEKLVDWIEAQGLTLKASISTHSHZDRTGGIGYLNSKG | 33 | 8.63999e-07 | 4.32e-05 | 2.16e-05 | 17 | HFHDDRTGGVPVLRARGIPVYALPDTARL | AYIIDTPWTEKDTEKLVDWIEAQGLTLKASISTHSHEDRTGGIGYLNSIG | + |
| HFHDDRTGGVPVLRARGIPVYALPDTARL | IPTYASELTNELLKKKGKPQA | -17 | 2.42791e-05 | 0.00121395 | 0.000404651 | 12 | HFHDDRTGGVPVLRARGIPVYALPDTARL | IPTYASELTNELLKKNGKPQA | + |
| HFHDDRTGGVPVLRARGIPVYALPDTARL | MSIQHFRVALIPFFAAFCLPVFAGEQGH | 1 | 0.000464742 | 0.0232371 | 0.00580927 | 27 | HFHDDRTGGVPVLRARGIPVYALPDTARL | MSIQHFRVALIPFFAAFCLPVFALSQGH | + |
| KIVVPGHGEWGGKDLLSHTJKLLK | VVPGHGKVGDASLLKHTIELA | -2 | 4.17819e-23 | 2.0891e-21 | 2.0891e-21 | 21 | KIVVPGHGEWGGKDLLSHTLKLLK | VVPGHGKVGDASLLKHTIELA | + |
| KIVVPGHGEWGGKDLLSHTJKLLK | AKJVVPGHGEVGDASLL | 1 | 2.71668e-16 | 1.35834e-14 | 6.79169e-15 | 16 | KIVVPGHGEWGGKDLLSHTLKLLK | AKLVVPGHGEVGDASLL | + |
| KIVVPGHGEWGGKDLLSHTJKLLK | DHTIKLFK | -16 | 4.66893e-06 | 0.000233446 | 7.78155e-05 | 8 | KIVVPGHGEWGGKDLLSHTLKLLK | DHTIDLFK | + |
| KLAKEQGYEVPBPILDELTTL | KENNFVVPQNSFNDSLVLKVGBEKVIAKF | -3 | 3.24581e-05 | 0.0016229 | 0.0016229 | 18 | KLAKEQGYEVPNPILDELTTL | KENNFVVPQNSFNDSLVLKVGNEKVIAKF | + |
| KLAKEQGYEVPBPILDELTTL | IPTYASELTNELLKKKGKPQA | 10 | 0.000375603 | 0.0187801 | 0.00939007 | 11 | KLAKEQGYEVPNPILDELTTL | IPTYASELTNELLKKNGKPQA | + |
| KNEENQVEITKJAEGVWVHTSYGTYNGGT | PAYTLQAQEQEJQVTKJAPGVWVHTSYSTYNGVLVPSHGLVVSTKEGAVL | 5 | 1.97091e-23 | 9.85455e-22 | 9.85455e-22 | 29 | KNEENQVEITKIAEGVWVHTSYGTYNGGT | PAYTLLAQEQELQVTKLAPGVWVHTSYSTYGGVLVPSHGLVVSTKGGAVL | + |
| KNEENQVEITKJAEGVWVHTSYGTYNGGT | IEKJSENVYLHTSFKZTNGWG | -8 | 9.25604e-16 | 4.62802e-14 | 2.31401e-14 | 21 | KNEENQVEITKIAEGVWVHTSYGTYNGGT | IEKLSENVYLHTSFKETNGWG | + |
| KNEENQVEITKJAEGVWVHTSYGTYNGGT | KJADGVYLHTSYKEVEGFGLVSSNGLVVV | -10 | 5.55167e-09 | 2.77584e-07 | 9.25279e-08 | 19 | KNEENQVEITKIAEGVWVHTSYGTYNGGT | KLADGVYLHTSYKEVEGFGLVSSNGLVVV | + |
| MFKLJSKLLVY | MKKJFLLLJFL | 0 | 0.000449543 | 0.0224771 | 0.0202347 | 11 | MFKLLSKLLVY | MKKLFLLLLFL | + |
| MFKLJSKLLVY | MKFLLLFLJ | 0 | 0.000809389 | 0.0404694 | 0.0202347 | 9 | MFKLLSKLLVY | MKFLLLFLL | + |
| MKRILLLLLLL | MKKJFLLLJFL | 0 | 1.83526e-10 | 9.17631e-09 | 9.17631e-09 | 11 | MKRILLLLLLL | MKKLFLLLLFL | + |
| MKRILLLLLLL | MKFLLLFLJ | 0 | 3.97835e-09 | 1.98918e-07 | 9.94589e-08 | 9 | MKRILLLLLLL | MKFLLLFLL | + |
| MKSLFAALFVVMFCGLARASD | MKFLLLFLJ | 0 | 0.000208406 | 0.0104203 | 0.0104203 | 9 | MKSLFAALFVVMFCGLARASD | MKFLLLFLL | + |
| MKSLFAALFVVMFCGLARASD | MKKJFLLLJFL | 0 | 0.00156742 | 0.0783711 | 0.0391855 | 11 | MKSLFAALFVVMFCGLARASD | MKKLFLLLLFL | + |
| MMKFRILGVFF | MKKJFLLLJFL | -1 | 9.2528e-06 | 0.00046264 | 0.00046264 | 10 | MMKFRILGVFF | MKKLFLLLLFL | + |
| MMKFRILGVFF | MKFLLLFLJ | -1 | 4.32072e-05 | 0.00216036 | 0.000910895 | 9 | MMKFRILGVFF | MKFLLLFLL | + |
| MMKFRILGVFF | MFSRILILLAGFJCCPPYAHSAESSGQLTITPLSSGALVVA | -1 | 5.46537e-05 | 0.00273269 | 0.000910895 | 10 | MMKFRILGVFF | MFSRILILLAGFLCCVPLAHSAESSGQLTITPLLSGALVVA | + |
| MSIQHFRVALI | MSIQHFRVALIPFFAAFCLPVFAGEQGH | 0 | 6.9457e-19 | 3.47285e-17 | 3.47285e-17 | 11 | MSIQHFRVALI | MSIQHFRVALIPFFAAFCLPVFALSQGH | + |
| RIFISIVFFMQTFGLVFAEPD | MFSSINALGKFTLYVLFIISFNLNAADPK | 8 | 1.82447e-05 | 0.000912237 | 0.000912237 | 21 | RIFISIVFFMQTFGLVFAEPD | MFSSINALGKFTLYVLFIISFNLNAADPK | + |
| SIKRVQQRYPK | SLGNTGDADLEAWPASIAKVQAR | 15 | 8.67874e-07 | 4.33937e-05 | 4.33937e-05 | 8 | SIKRVQQRYPK | SLGNTGDADLEAWPASIAKVQAR | + |
| SIKRVQQRYPK | GDABVSAWPNSVEKVKKK | 10 | 4.50288e-06 | 0.000225144 | 0.000112572 | 8 | SIKRVQQRYPK | GDANVSAWPNSVEKVKKK | + |
| VRFGPVELFFPGAGHSPDNLV | IEVFYPGAGHTKDNIVVWLPKQKILFGGCLVKS | -5 | 2.11411e-13 | 1.05706e-11 | 1.05706e-11 | 16 | VRFGPVELFFPGAGHSPDNLV | IEVFYPGAGHTKDNIVVWLPKQKILFGGCLVKS | + |
| VRFGPVELFFPGAGHSPDNLV | GPGHTKDNVVVWLPKEKILFGGCLVKSLGAGNLGL | -11 | 0.000145065 | 0.00725323 | 0.00362662 | 10 | VRFGPVELFFPGAGHSPDNLV | GPGHTKDNVVVWLPKEKILFGGCLVKSLGAGNLGL | + |
| VTHAHDDRIGGIDVLKKRGIPVYSTPLTA | RGYKJKASISTHFHEDSTGGJEYLNSHSIPTYASELTNELL | 9 | 1.20168e-19 | 6.00839e-18 | 6.00839e-18 | 29 | VTHAHDDRIGGIDVLKKRGIPVYSTPLTA | RGYKIKASISTHFHEDSTGGLEYLNSHSIPTYASELTNELL | + |
| VTHAHDDRIGGIDVLKKRGIPVYSTPLTA | AYJIDTPWTEKDTEKLVDWIEAQGLTLKASISTHSHZDRTGGIGYLNSKG | 31 | 1.69716e-09 | 8.48581e-08 | 4.2429e-08 | 19 | VTHAHDDRIGGIDVLKKRGIPVYSTPLTA | AYIIDTPWTEKDTEKLVDWIEAQGLTLKASISTHSHEDRTGGIGYLNSIG | + |
| VTHAHDDRIGGIDVLKKRGIPVYSTPLTA | IPTYASELTNELLKKKGKPQA | -19 | 0.00105376 | 0.0526881 | 0.0161204 | 10 | VTHAHDDRIGGIDVLKKRGIPVYSTPLTA | IPTYASELTNELLKKNGKPQA | + |
| VTHAHDDRIGGIDVLKKRGIPVYSTPLTA | FDTPWTBEPTEQLLAWVKDNLKAPVKAFVPTHWHDDCLGGL | 29 | 0.00128963 | 0.0644817 | 0.0161204 | 12 | VTHAHDDRIGGIDVLKKRGIPVYSTPLTA | FDTPWTDEPTEQLLAWVKDNLKAPVKAFVPTHWHDDCLGGL | + |
| VTHAHDDRIGGIDVLKKRGIPVYSTPLTA | MPNRALWQRIAILLLMPILSLPALAANLT | 1 | 0.00363586 | 0.181793 | 0.0363586 | 28 | VTHAHDDRIGGIDVLKKRGIPVYSTPLTA | MPNRALWQRIAILLLMPILSLPALAANLT | + |
| VVPGHGLPGGPELLDHTEALL | VVPGHGKVGDASLLKHTIELA | 0 | 4.901e-16 | 2.4505e-14 | 2.4505e-14 | 21 | VVPGHGLPGGPELLDHTEALL | VVPGHGKVGDASLLKHTIELA | + |
| VVPGHGLPGGPELLDHTEALL | AKJVVPGHGEVGDASLL | 3 | 2.7573e-09 | 1.37865e-07 | 6.89326e-08 | 14 | VVPGHGLPGGPELLDHTEALL | AKLVVPGHGEVGDASLL | + |
| VVPGHGLPGGPELLDHTEALL | DHTIKLFK | -14 | 0.00174629 | 0.0873142 | 0.0291048 | 7 | VVPGHGLPGGPELLDHTEALL | DHTIDLFK | + |
| VWHPSSGVLFGGCAVKDASAKSLGNVADADLAAWPASJERIRQRYPEARV | SLGNTGDADLEAWPASIAKVQAR | -21 | 2.86539e-17 | 1.4327e-15 | 1.4327e-15 | 23 | VWHPSSGVLFGGCAVKDASAKSLGNVADADLAAWPASLERIRQRYPEARV | SLGNTGDADLEAWPASIAKVQAR | + |
| VWHPSSGVLFGGCAVKDASAKSLGNVADADLAAWPASJERIRQRYPEARV | GPGHTKDNVVVWLPKEKILFGGCLVKSLGAGNLGL | 10 | 5.07047e-11 | 2.53523e-09 | 1.26762e-09 | 25 | VWHPSSGVLFGGCAVKDASAKSLGNVADADLAAWPASLERIRQRYPEARV | GPGHTKDNVVVWLPKEKILFGGCLVKSLGAGNLGL | + |
| VWHPSSGVLFGGCAVKDASAKSLGNVADADLAAWPASJERIRQRYPEARV | GDABVSAWPNSVEKVKKK | -26 | 6.47365e-09 | 3.23682e-07 | 1.07894e-07 | 18 | VWHPSSGVLFGGCAVKDASAKSLGNVADADLAAWPASLERIRQRYPEARV | GDANVSAWPNSVEKVKKK | + |
| VWHPSSGVLFGGCAVKDASAKSLGNVADADLAAWPASJERIRQRYPEARV | IEVFYPGAGHTKDNIVVWLPKQKILFGGCLVKS | 16 | 1.36838e-06 | 6.8419e-05 | 1.71047e-05 | 17 | VWHPSSGVLFGGCAVKDASAKSLGNVADADLAAWPASLERIRQRYPEARV | IEVFYPGAGHTKDNIVVWLPKQKILFGGCLVKS | + |
| VWHPSSGVLFGGCAVKDASAKSLGNVADADLAAWPASJERIRQRYPEARV | YPDAKJ | -44 | 3.47558e-05 | 0.00173779 | 0.000347558 | 6 | VWHPSSGVLFGGCAVKDASAKSLGNVADADLAAWPASLERIRQRYPEARV | YPDAKL | + |
| YPANGLIVEDGDELLLVDTAWGARQTAALL | PCNGLVVIDNKEAYJIDTPWNAKDTE | -1 | 6.90071e-15 | 3.45036e-13 | 3.45036e-13 | 26 | YPANGLIVEDGDELLLVDTAWGARQTAALL | PCNGLVVIDNKEAYLIDTPWNAKDTE | + |
| YPANGLIVEDGDELLLVDTAWGARQTAALL | FDTPWTBEPTEQLLAWVKDNLKAPVKAFVPTHWHDDCLGGL | -16 | 0.000196831 | 0.00984153 | 0.00492076 | 14 | YPANGLIVEDGDELLLVDTAWGARQTAALL | FDTPWTDEPTEQLLAWVKDNLKAPVKAFVPTHWHDDCLGGL | + |
| YPANGLIVEDGDELLLVDTAWGARQTAALL | AYJIDTPWTEKDTEKLVDWIEAQGLTLKASISTHSHZDRTGGIGYLNSKG | -13 | 0.000396403 | 0.0198201 | 0.00660671 | 17 | YPANGLIVEDGDELLLVDTAWGARQTAALL | AYIIDTPWTEKDTEKLVDWIEAQGLTLKASISTHSHEDRTGGIGYLNSIG | + |
| YPANGLIVEDGDELLLVDTAWGARQTAALL | PAYTLQAQEQEJQVTKJAPGVWVHTSYSTYNGVLVPSHGLVVSTKEGAVL | 34 | 0.000809864 | 0.0404932 | 0.0101233 | 16 | YPANGLIVEDGDELLLVDTAWGARQTAALL | PAYTLLAQEQELQVTKLAPGVWVHTSYSTYGGVLVPSHGLVVSTKGGAVL | + |

**VIM vs NDM**

| Query_ID | Target_ID | Optimal_offset | p-value | E-value | q-value | Overlap | Query_consensus | Target_consensus | Orientation |
| --- | --- | --- | --- | --- | --- | --- | --- | --- | --- |
| AKDLGNLADADVAAWPASLER.1 | NLGDPDC | -5 | 0.00146101 | 0.0482132 | 0.0482132 | 7 | AKDLGNLADADVAAWPASLER | NLGDADT | + |
| AKDLGNLADADVAAWPASLER.2 | NLGDPDC | -5 | 0.00146101 | 0.0482132 | 0.0482132 | 7 | AKDLGNLADADVAAWPASLER | NLGDADT | + |
| DAQTLGPLEVFFPGAGHAPDNJVVWHPASGVLFGGCFVKDA.1 | WVEPATAPNFGPLKVFYPGPGHTSDNITVGIDGTDIAFGGCLIKDSKAKS.2 | 5 | 1.00998e-16 | 3.33293e-15 | 1.78178e-15 | 41 | DAQTLGPLEVFFPGAGHAPDNLVVWHPASGVLFGGCFVKDA | WVEPATAPNFGPLKVFYPGPGHTSDNITVGIDGTDIAFGGCLIKDSKAKS | + |
| DAQTLGPLEVFFPGAGHAPDNJVVWHPASGVLFGGCFVKDA.1 | WVEPATAPNFGPLKVFYPGPGHTSDNITVGIDGTDIAFGGCLIKDSKAKS.1 | 5 | 1.07987e-16 | 3.56356e-15 | 1.78178e-15 | 41 | DAQTLGPLEVFFPGAGHAPDNLVVWHPASGVLFGGCFVKDA | WVEPATAPNFGPLKVFYPGPGHTSDNITVGIDGTDIAFGGCLIKDSKAKS | + |
| DAQTLGPLEVFFPGAGHAPDNJVVWHPASGVLFGGCFVKDA.2 | WVEPATAPNFGPLKVFYPGPGHTSDNITVGIDGTDIAFGGCLIKDSKAKS.2 | 5 | 1.00998e-16 | 3.33293e-15 | 1.78178e-15 | 41 | DAQTLGPLEVFFPGAGHAPDNLVVWHPASGVLFGGCFVKDA | WVEPATAPNFGPLKVFYPGPGHTSDNITVGIDGTDIAFGGCLIKDSKAKS | + |
| DAQTLGPLEVFFPGAGHAPDNJVVWHPASGVLFGGCFVKDA.2 | WVEPATAPNFGPLKVFYPGPGHTSDNITVGIDGTDIAFGGCLIKDSKAKS.1 | 5 | 1.07987e-16 | 3.56356e-15 | 1.78178e-15 | 41 | DAQTLGPLEVFFPGAGHAPDNLVVWHPASGVLFGGCFVKDA | WVEPATAPNFGPLKVFYPGPGHTSDNITVGIDGTDIAFGGCLIKDSKAKS | + |
| HRPVRAAVVTHFHDDRTGGIPALVARGIPVYALED.1 | VVTHAHQDKMGGMDALHAAGIATYANALSNQLAPQEGMVAAQHSLTFAAN | -7 | 1.06288e-09 | 3.50749e-08 | 3.50749e-08 | 28 | HRPVRAAVVTHFHDDRTGGIPALVARGIPVHALED | VVTHAHQDKMGGMDALHAAGIATYANALSNQLAPQEGMVAAQHSLTFAAN | + |
| HRPVRAAVVTHFHDDRTGGIPALVARGIPVYALED.1 | VRDGGRVLVVDTAWTDDQTAQILNWIKQEINLPVALAVVTHAHQDKMGGM | 30 | 2.9351e-05 | 0.000968583 | 0.000484292 | 20 | HRPVRAAVVTHFHDDRTGGIPALVARGIPVHALED | VRDGGRVLVVDTAWTDDQTAQILNWIKQEINLPVALAVVTHAHQDKMGGM | + |
| HRPVRAAVVTHFHDDRTGGIPALVARGIPVYALED.1 | DALHAAGI | -20 | 0.000227752 | 0.00751582 | 0.00250527 | 8 | HRPVRAAVVTHFHDDRTGGIPALVARGIPVHALED | DALHAAGI | + |
| HRPVRAAVVTHFHDDRTGGIPALVARGIPVYALED.2 | VVTHAHQDKMGGMDALHAAGIATYANALSNQLAPQEGMVAAQHSLTFAAN | -7 | 1.06288e-09 | 3.50749e-08 | 3.50749e-08 | 28 | HRPVRAAVVTHFHDDRTGGIPALVARGIPVHALED | VVTHAHQDKMGGMDALHAAGIATYANALSNQLAPQEGMVAAQHSLTFAAN | + |
| HRPVRAAVVTHFHDDRTGGIPALVARGIPVYALED.2 | VRDGGRVLVVDTAWTDDQTAQILNWIKQEINLPVALAVVTHAHQDKMGGM | 30 | 2.9351e-05 | 0.000968583 | 0.000484292 | 20 | HRPVRAAVVTHFHDDRTGGIPALVARGIPVHALED | VRDGGRVLVVDTAWTDDQTAQILNWIKQEINLPVALAVVTHAHQDKMGGM | + |
| HRPVRAAVVTHFHDDRTGGIPALVARGIPVYALED.2 | DALHAAGI | -20 | 0.000227752 | 0.00751582 | 0.00250527 | 8 | HRPVRAAVVTHFHDDRTGGIPALVARGIPVHALED | DALHAAGI | + |
| LVDTGWGPRQTEALLDWARDTL.1 | VRDGGRVLVVDTAWTDDQTAQILNWIKQEINLPVALAVVTHAHQDKMGGM | 8 | 8.98768e-08 | 2.96593e-06 | 2.20742e-06 | 22 | LVDTGWGPRQTEALLDWARDTL | VRDGGRVLVVDTAWTDDQTAQILNWIKQEINLPVALAVVTHAHQDKMGGM | + |
| LVDTGWGPRQTEALLDWARDTL.1 | WQHTSYLDMPGFGAVASNGLIVRDGGRVLVVDTAWTDDQTAQILNWIKQE | 29 | 1.33783e-07 | 4.41485e-06 | 2.20742e-06 | 21 | LVDTGWGPRQTEALLDWARDTL | WQHTSYLDMPGFGAVASNGLIVRDGGRVLVVDTAWTDDQTAQILNWIKQE | + |
| LVDTGWGPRQTEALLDWARDTL.2 | VRDGGRVLVVDTAWTDDQTAQILNWIKQEINLPVALAVVTHAHQDKMGGM | 8 | 8.98768e-08 | 2.96593e-06 | 2.20742e-06 | 22 | LVDTGWGPRQTEALLDWARDTL | VRDGGRVLVVDTAWTDDQTAQILNWIKQEINLPVALAVVTHAHQDKMGGM | + |
| LVDTGWGPRQTEALLDWARDTL.2 | WQHTSYLDMPGFGAVASNGLIVRDGGRVLVVDTAWTDDQTAQILNWIKQE | 29 | 1.33783e-07 | 4.41485e-06 | 2.20742e-06 | 21 | LVDTGWGPRQTEALLDWARDTL | WQHTSYLDMPGFGAVASNGLIVRDGGRVLVVDTAWTDDQTAQILNWIKQE | + |
| RARYPEARVVVPGHGAPGGPELLDHTEAL.1 | LGDADTEHYAASARAFGAAFPKASMIVMSHSAPDSRAAITHTARMADKLR.2 | 16 | 6.15713e-05 | 0.00203185 | 0.00106494 | 29 | RARYPEARVVVPGHGAPGGPELLDHTEAL | LGDADTEHYAASARAFGAAFPKASMIVMSHSAPDSRAAITHTARMADKLR | + |
| RARYPEARVVVPGHGAPGGPELLDHTEAL.1 | LGDADTEHYAASARAFGAAFPKASMIVMSHSAPDSRAAITHTARMADKLR.1 | 16 | 6.45417e-05 | 0.00212988 | 0.00106494 | 29 | RARYPEARVVVPGHGAPGGPELLDHTEAL | LGDADTEHYAASARAFGAAFPKASMIVMSHSAPDSRAAITHTARMADKLR | + |
| RARYPEARVVVPGHGAPGGPELLDHTEAL.2 | LGDADTEHYAASARAFGAAFPKASMIVMSHSAPDSRAAITHTARMADKLR.2 | 16 | 6.15713e-05 | 0.00203185 | 0.00106494 | 29 | RARYPEARVVVPGHGAPGGPELLDHTEAL | LGDADTEHYAASARAFGAAFPKASMIVMSHSAPDSRAAITHTARMADKLR | + |
| RARYPEARVVVPGHGAPGGPELLDHTEAL.2 | LGDADTEHYAASARAFGAAFPKASMIVMSHSAPDSRAAITHTARMADKLR.1 | 16 | 6.45417e-05 | 0.00212988 | 0.00106494 | 29 | RARYPEARVVVPGHGAPGGPELLDHTEAL | LGDADTEHYAASARAFGAAFPKASMIVMSHSAPDSRAAITHTARMADKLR | + |
| YPANGLJVEDGDGSJ.1 | WQHTSYLDMPGFGAVASNGLIVRDGGRVLVVDTAWTDDQTAQILNWIKQE | 14 | 4.77199e-08 | 1.57476e-06 | 1.57476e-06 | 15 | YPANGLIVEDGDGSL | WQHTSYLDMPGFGAVASNGLIVRDGGRVLVVDTAWTDDQTAQILNWIKQE | + |
| YPANGLJVEDGDGSJ.1 | ASNGLI | -1 | 0.000109236 | 0.00360479 | 0.0018024 | 6 | YPANGLIVEDGDGSL | ASNGLI | + |
| YPANGLJVEDGDGSJ.2 | WQHTSYLDMPGFGAVASNGLIVRDGGRVLVVDTAWTDDQTAQILNWIKQE | 14 | 4.77199e-08 | 1.57476e-06 | 1.57476e-06 | 15 | YPANGLIVEDGDGSL | WQHTSYLDMPGFGAVASNGLIVRDGGRVLVVDTAWTDDQTAQILNWIKQE | + |
| YPANGLJVEDGDGSJ.2 | ASNGLI | -1 | 0.000109236 | 0.00360479 | 0.0018024 | 6 | YPANGLIVEDGDGSL | ASNGLI | + |
| AEDTJGLPVRAAVVT | VRDGGRVLVVDTAWTDDQTAQILNWIKQEINLPVALAVVTHAHQDKMGGM | 25 | 7.79675e-07 | 2.57293e-05 | 2.57293e-05 | 15 | AEDTLGLPVRAAVVT | VRDGGRVLVVDTAWTDDQTAQILNWIKQEINLPVALAVVTHAHQDKMGGM | + |
| AEDTJGLPVRAAVVT | INLPVALA | -4 | 0.000613133 | 0.0202334 | 0.0101167 | 8 | AEDTLGLPVRAAVVT | INLPVALA | + |
| EVFYPGPGHTMDNIVVWLPZQKILFGGCLVKSLQAKDLGNTADADLNEWP | WVEPATAPNFGPLKVFYPGPGHTSDNITVGIDGTDIAFGGCLIKDSKAKS.1 | 13 | 7.97441e-16 | 2.63156e-14 | 1.49092e-14 | 37 | EVFYPGAGHTMDNIVVWLPQQKILFGGCLVKSLQAKDLGNTADADLNSWP | WVEPATAPNFGPLKVFYPGPGHTSDNITVGIDGTDIAFGGCLIKDSKAKS | + |
| EVFYPGPGHTMDNIVVWLPZQKILFGGCLVKSLQAKDLGNTADADLNEWP | WVEPATAPNFGPLKVFYPGPGHTSDNITVGIDGTDIAFGGCLIKDSKAKS.2 | 13 | 9.03587e-16 | 2.98184e-14 | 1.49092e-14 | 37 | EVFYPGAGHTMDNIVVWLPQQKILFGGCLVKSLQAKDLGNTADADLNSWP | WVEPATAPNFGPLKVFYPGPGHTSDNITVGIDGTDIAFGGCLIKDSKAKS | + |
| EWIKTKLKKPVKKAI | VRDGGRVLVVDTAWTDDQTAQILNWIKQEINLPVALAVVTHAHQDKMGGM | 23 | 3.22318e-07 | 1.06365e-05 | 1.06365e-05 | 15 | EWIKTKLKKPVKKAI | VRDGGRVLVVDTAWTDDQTAQILNWIKQEINLPVALAVVTHAHQDKMGGM | + |
| EWIKTKLKKPVKKAI | INLPVALA | -6 | 0.000107904 | 0.00356083 | 0.00178042 | 8 | EWIKTKLKKPVKKAI | INLPVALA | + |
| FGNTKV | WVEPATAPNFGPLKVFYPGPGHTSDNITVGIDGTDIAFGGCLIKDSKAKS.1 | 9 | 0.000629134 | 0.0207614 | 0.0103807 | 6 | FGNTKV | WVEPATAPNFGPLKVFYPGPGHTSDNITVGIDGTDIAFGGCLIKDSKAKS | + |
| FGNTKV | WVEPATAPNFGPLKVFYPGPGHTSDNITVGIDGTDIAFGGCLIKDSKAKS.2 | 9 | 0.000629134 | 0.0207614 | 0.0103807 | 6 | FGNTKV | WVEPATAPNFGPLKVFYPGPGHTSDNITVGIDGTDIAFGGCLIKDSKAKS | + |
| FPSNGLIVETGKGLVLIDTAWGEEQTEZL | WQHTSYLDMPGFGAVASNGLIVRDGGRVLVVDTAWTDDQTAQILNWIKQE | 14 | 1.00805e-13 | 3.32655e-12 | 3.32655e-12 | 29 | FPSNGLIVETGKGLVLIDTAWGEEQTEEL | WQHTSYLDMPGFGAVASNGLIVRDGGRVLVVDTAWTDDQTAQILNWIKQE | + |
| FPSNGLIVETGKGLVLIDTAWGEEQTEZL | VRDGGRVLVVDTAWTDDQTAQILNWIKQEINLPVALAVVTHAHQDKMGGM | -7 | 8.819e-06 | 0.000291027 | 0.000116143 | 22 | FPSNGLIVETGKGLVLIDTAWGEEQTEEL | VRDGGRVLVVDTAWTDDQTAQILNWIKQEINLPVALAVVTHAHQDKMGGM | + |
| FPSNGLIVETGKGLVLIDTAWGEEQTEZL | ASNGLI | -1 | 1.05584e-05 | 0.000348428 | 0.000116143 | 6 | FPSNGLIVETGKGLVLIDTAWGEEQTEEL | ASNGLI | + |
| HFHDDRTGGVPVLRARGIPVYALPDTARL | VVTHAHQDKMGGMDALHAAGIATYANALSNQLAPQEGMVAAQHSLTFAAN | 3 | 1.69752e-11 | 5.60182e-10 | 5.60182e-10 | 29 | HFHDDRTGGVPVLRARGIPVYALPDTARL | VVTHAHQDKMGGMDALHAAGIATYANALSNQLAPQEGMVAAQHSLTFAAN | + |
| HFHDDRTGGVPVLRARGIPVYALPDTARL | DALHAAGI | -10 | 1.1864e-05 | 0.000391511 | 0.000195755 | 8 | HFHDDRTGGVPVLRARGIPVYALPDTARL | DALHAAGI | + |
| MSIQHFRVALI | SIQHFRVALIPFFAAFCLPVF | -1 | 3.41107e-12 | 1.12565e-10 | 1.12565e-10 | 10 | MSIQHFRVALI | SIQHFRVALIPFFAAFCLPVF | + |
| VRFGPVELFFPGAGHSPDNLV | WVEPATAPNFGPLKVFYPGPGHTSDNITVGIDGTDIAFGGCLIKDSKAKS.1 | 7 | 5.95818e-11 | 1.9662e-09 | 9.831e-10 | 21 | VRFGPVELFFPGAGHSPDNLV | WVEPATAPNFGPLKVFYPGPGHTSDNITVGIDGTDIAFGGCLIKDSKAKS | + |
| VRFGPVELFFPGAGHSPDNLV | WVEPATAPNFGPLKVFYPGPGHTSDNITVGIDGTDIAFGGCLIKDSKAKS.2 | 7 | 5.95818e-11 | 1.9662e-09 | 9.831e-10 | 21 | VRFGPVELFFPGAGHSPDNLV | WVEPATAPNFGPLKVFYPGPGHTSDNITVGIDGTDIAFGGCLIKDSKAKS | + |
| VTHAHDDRIGGIDVLKKRGIPVYSTPLTA | VVTHAHQDKMGGMDALHAAGIATYANALSNQLAPQEGMVAAQHSLTFAAN | 1 | 2.05474e-13 | 6.78064e-12 | 6.78064e-12 | 29 | VTHAHDDRIGGIDVLKKRGIPVYSTPLTA | VVTHAHQDKMGGMDALHAAGIATYANALSNQLAPQEGMVAAQHSLTFAAN | + |
| VTHAHDDRIGGIDVLKKRGIPVYSTPLTA | DALHAAGI | -12 | 0.000311532 | 0.0102806 | 0.00514029 | 8 | VTHAHDDRIGGIDVLKKRGIPVYSTPLTA | DALHAAGI | + |
| YPANGLIVEDGDELLLVDTAWGARQTAALL | WQHTSYLDMPGFGAVASNGLIVRDGGRVLVVDTAWTDDQTAQILNWIKQE | 14 | 6.40183e-16 | 2.1126e-14 | 2.1126e-14 | 30 | YPANGLIVEDGDELLLVDTAWGARQTAALL | WQHTSYLDMPGFGAVASNGLIVRDGGRVLVVDTAWTDDQTAQILNWIKQE | + |
| YPANGLIVEDGDELLLVDTAWGARQTAALL | VRDGGRVLVVDTAWTDDQTAQILNWIKQEINLPVALAVVTHAHQDKMGGM | -7 | 4.48981e-08 | 1.48164e-06 | 7.40818e-07 | 23 | YPANGLIVEDGDELLLVDTAWGARQTAALL | VRDGGRVLVVDTAWTDDQTAQILNWIKQEINLPVALAVVTHAHQDKMGGM | + |
| YPANGLIVEDGDELLLVDTAWGARQTAALL | ASNGLI | -1 | 4.15991e-05 | 0.00137277 | 0.00045759 | 6 | YPANGLIVEDGDELLLVDTAWGARQTAALL | ASNGLI | + |
