## Supplementary File 9 for "Phylogenetic Analysis of Beta-Lactamases Reveals Distinct Evolutionary Patterns of Chromosomal and Plasmid-Encoded BLs and the Mosaic Role of VIM Linking NDM and IMP"

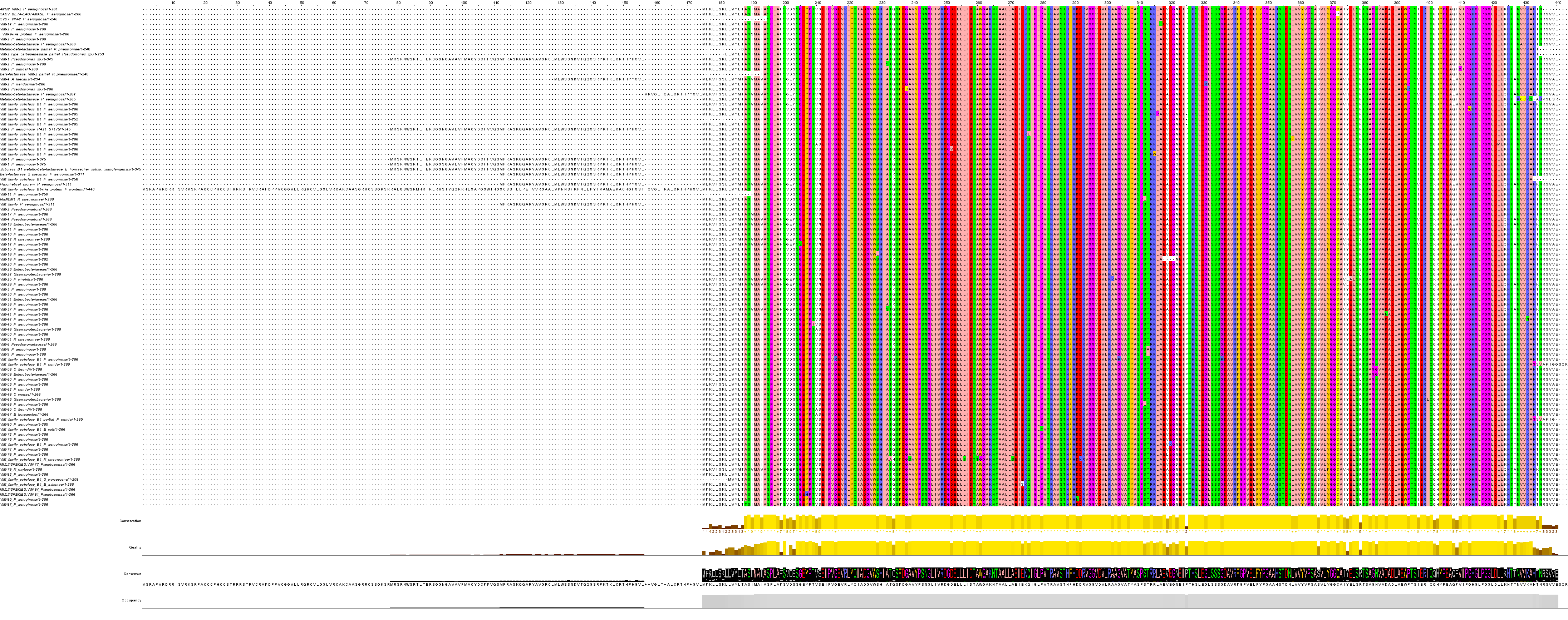

**Figure S1a**. Multiple sequence alignment of the VIM protein displayed with Jalview's color scheme, which highlights residues based on their chemical properties when conserved or similar across sequences. The figure also displays occupancy, consensus, quality, and sequence conservation levels.

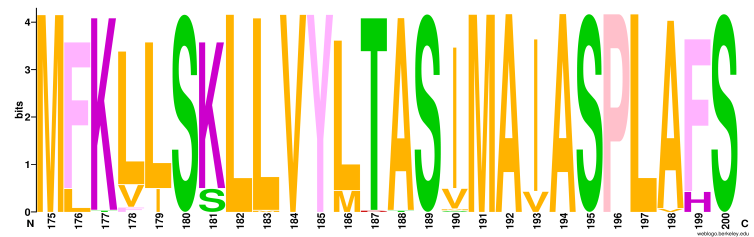

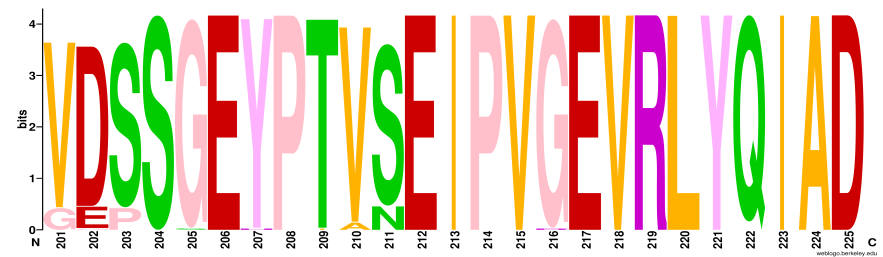

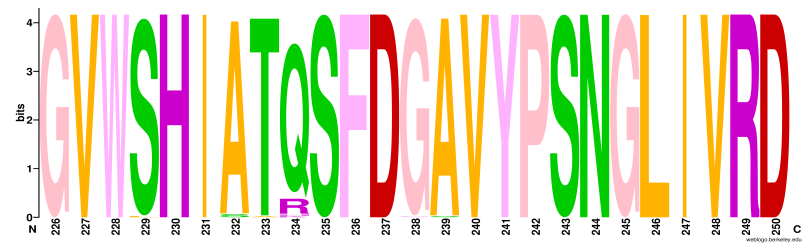

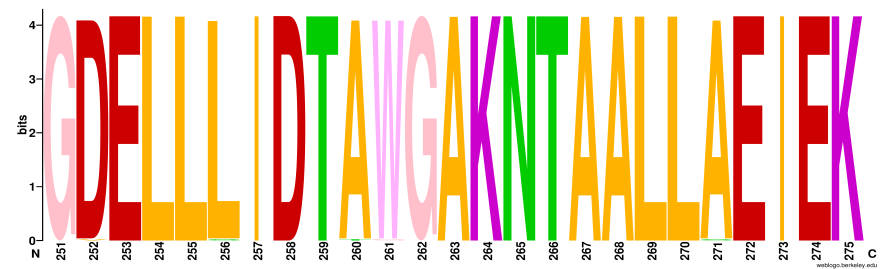

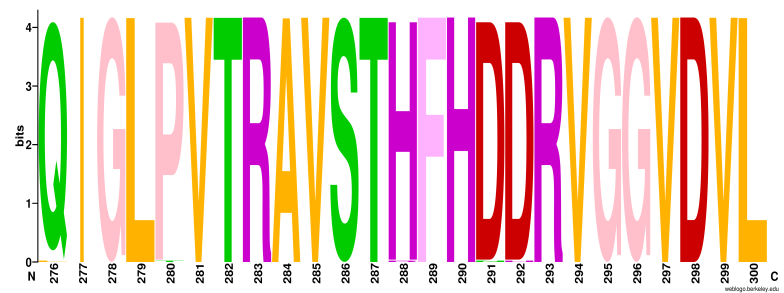

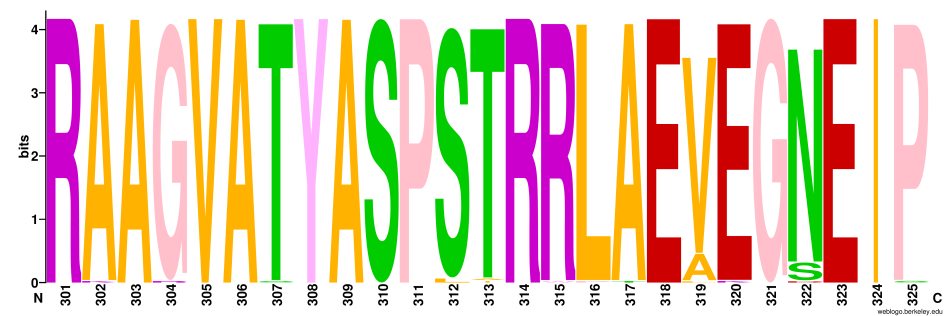

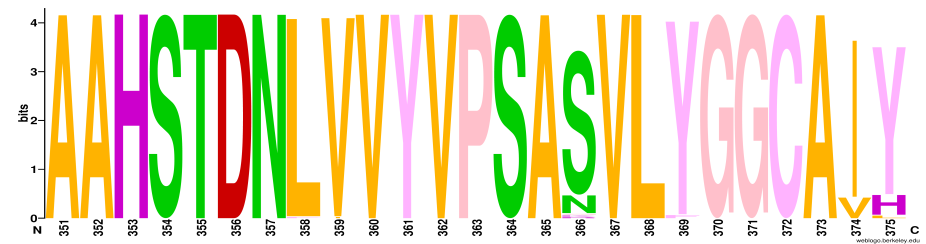

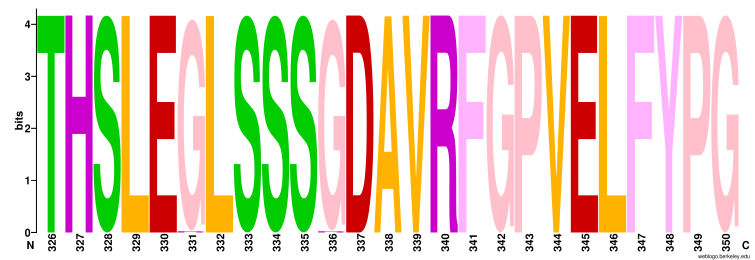

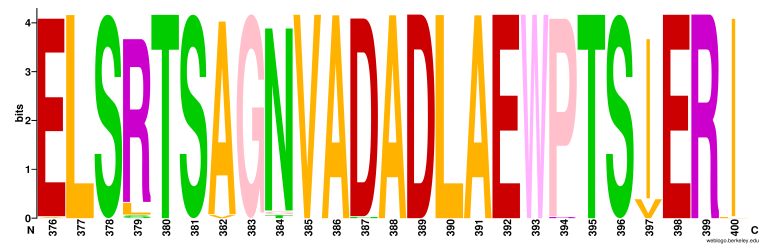

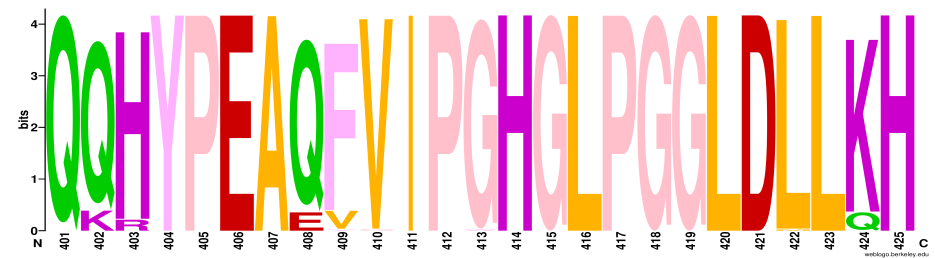

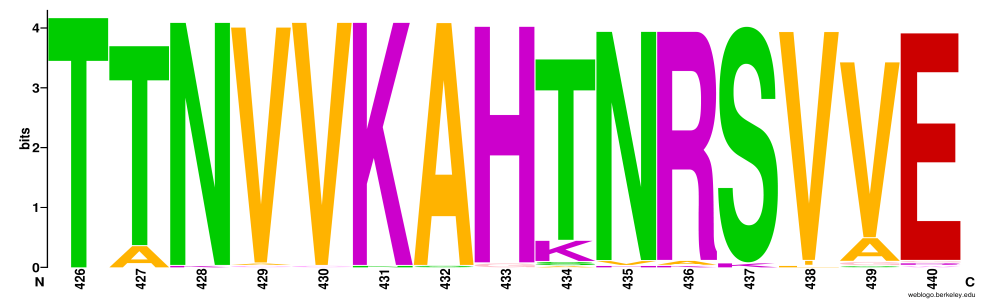

**Figure S1b.** Sequence logos illustrating residue conservation in VIM were generated for all aligned blocks of the MSA using the WebLogo web server (<https://weblogo.berkeley.edu/logo.cgi>). The x-axis represents the column numbers from the MSA.

**­­­­Table S1:** List of mutations in VIM occurring in highly conserved columns (approximately 98% conserved across all species) identified through multiple sequence alignment (MSA).

| **Protein and Bacteria** | **Position of Mutation (Amino Acid)** | **Amino Acid Mutation** |
| --- | --- | --- |
| VIM B1 Beta lactamase - *Klebsiella pneumoniae* | 198 Position | Alanine (non polar) Valine (non polar) |
| VIM 62- *Pseudomonas putida* | 198 Position | Alanine (non polar) Valine (non polar) |
| VIM 14- *Pseudomonas aeruginosa* | 205 Position | Glycine (non polar) Serine (polar) |
| VIM 81- *Pseudomonas* (Multispecies) | 207 Position | Tyrosine (uncharged) Histidine (charged) |
| VIM45- - *Pseudomonas aeruginosa* | 209 Position | Threonine (polar) Isoleucine (nonpolar) |
| VIM family subclass B1 -*Pseudomonas aeruginosa* | 216 Position | Glycine (non polar, uncharged) Arginine (polar, charged) |
| VIM16 - *Pseudomonas aeruginosa* | 229 Position | Serine (polar) Leucine (non polar) |
| VIM2 - *Pseudomonas aeruginosa* | 232 Position | Alanine (non polar) Serine (polar) |
| VIM37 - *Pseudomonas aeruginosa* | 232 Position | Alanine (non polar) Serine (polar) |
| VIM family subclass B1 *Klebsiella pneumoniae* | 233 Position | Tyrosine (polar) Alanine (non polar) |
| VIM2 - *Pseudomonas aeruginosa* | 237 Position | Glycine (non polar) Cysteine (polar) |
| VIM family subclass B1 *Klebsiella pneumoniae* | 238 Position | Alanine (non polar) Serine (polar) |
| VIM family subclass B1 -*Pseudomonas aeruginosa* | 252 Position | Aspartate (-ve charged) Valine (neutral) |
| VIM family subclass B1 *Klebsiella pneumoniae* | 256 Position | Leucine (non polar) Serine (polar) |
| VIM family subclass B1 *Klebsiella pneumoniae* | 260 Position | Alanine (non polar) Threonine (polar) |
| VIM family subclass B1 *Klebsiella pneumoniae* | 261 Position | Trypthophan (aromatic) Glycine (aliphatic) |
| VIM family subclass B1 *Klebsiella pneumoniae* | 271 Position | Alanine (non polar) Serine (polar) |
| VIM family subclass B1 -*Pseudomonas aeruginosa* | 276 Position | Glutamine (polar) Leucine (non polar) |
| VIM family subclass B1 *Escherichia coli* | 280 Position | Proline (non polar) Threonine (polar) |
| VIM family subclass B1 -*Pseudomonas aeruginosa* | 288 Position | Histidine (+ve charged) Arginine (+ve charged) |
| VIM 85- *Pseudomonas aeruginosa* | 291 Position | Asparagine (Uncharged) Aspartate (-ve charged) |
| VIM family subclass B1 *Klebsiella pneumoniae* | 292 Position | Aspartate (-ve charged) Histidine (+ve charged) |
| VIM25 - *Proteus mirabilis* | 301 Position | Alanine (non polar) Lysine (+ve charged) |
| VIM family subclass B1 -*Pseudomonas aeruginosa* | 304 Position | Glycine (non polar) Arginine (+ve charged) |
| VIM family subclass B1 -*Pseudomonas aeruginosa* | 307 Postion | Threonine (polar) Serine (polar) |
| blaNDM1- *Klebsiella pneumoniae* | 312 Position | Serine (polar) Leucine (non polar) |
| VIM66- *Pseudomonas aeruginosa* | 312 Position | Serine (polar) Leucine (non polar) |
| VIM8- *Pseudomonas aeruginosa* | 313 Position | Threonine (polar) Alanine (non polar) |
| VIM8- *Pseudomonas aeruginosa* | 313 Position | Threonine (polar) Isoleucine (non polar) |
| VIM family subclass B1 -*Pseudomonas aeruginosa* | 315 Position | Arginine (+ve charged, aliphatic) Tryptophan (non polar, aromatic) |
| VIM family subclass B1 -*Pseudomonas aeruginosa* | 316 Position | Leucine (aliphatic) Proline (ring structure) |
| VIM14- *Pseudomonas aeruginosa* | 317 Position | Alanine (non polar) Threonine (polar) |
| VIM family subclass B1 -*Pseudomonas aeruginosa* | 320 Position | Glutamate (-ve charged) Lysine (+ve charged) |
| VIM2 - *Pseudomonas mendocina* | 321 Position | Glycine (non polar) Arginine (+ve charged) |
| VIM family subclass B1 -*Pseudomonas aeruginosa* | 336 Position | Glycine (non polar) Arginine (+ve charged) |
| VIM family subclass B1 -*Pseudomonas aeruginosa* | 358 Position | Leucine (aliphatic) Phenylalanine (aromatic) |
| VIM2 - *Pseudomonas aeruginosa* | 359 Position | Valine(non polar) Isoleucine (non polar) |
| VIM15- *Pseudomonas aeruginosa* | 369 Position | Tyrosine (aromatic) Phenylalanine (aromatic) |
| VIM25 - *Proteus mirabilis* | 376 Position | Glutamate (+charged) Alanine (non polar) |
| VIM51- *Klebsiella pneumoniae* | 382 Position | Alanine (non polar) Valine (Polar) |
| VIM65-*Citrobacter freundii* | 382 Position | Alanine (non polar) Valine (Polar) |
| VIM41- *Pseudomonas aeruginosa* | 387 Position | Aspartate (-ve charged) Asparagine (uncharged) |
| VIM2 - *Pseudomonas aeruginosa* | 394 Position | Proline (uncharged) Arginine (+ve charged) |
| VIM63- Gammaproteobacteria | 400 Position | Isoleucine (non polar) Leucine (polar) |
| VIM2- *Pseudomonas putida* | 410 Position | Valine (non polar) Glycine (non polar) |
| VIM family subclass B1 -*Pseudomonas aeruginosa* | 403 Position | Glycine (Aliphatic) Tryptophan (Aromatic) |
| VIM family subclass B1 -*Pseudomonas aeruginosa* | 422 Position | Leucine (non polar) Methionine (polar) |
| CAE46566.1-Metallo-beta-lactamase *Pseudomonas aeruginosa* | 428 Position | Asparagine (uncharged) Lysine (+ ve charged) |
| CCE14945.1-Metallo-beta-lactamase *Pseudomonas aeruginosa* | 429 Position | Valine (non polar) Cysteine (polar) |
| CCE14945.1-Metallo-beta-lactamase *Pseudomonas aeruginosa* | 430 Position | Valine (non polar) Cysteine (polar) |
| VIM44- *Pseudomonas aeruginosa* | 431 Position | Lysine (+ve charged) Asparagine (uncharged) |
| CCE14945.1-Metallo-beta-lactamase *Pseudomonas aeruginosa* | 432 Position | Alanine (non polar) Serine (polar) |

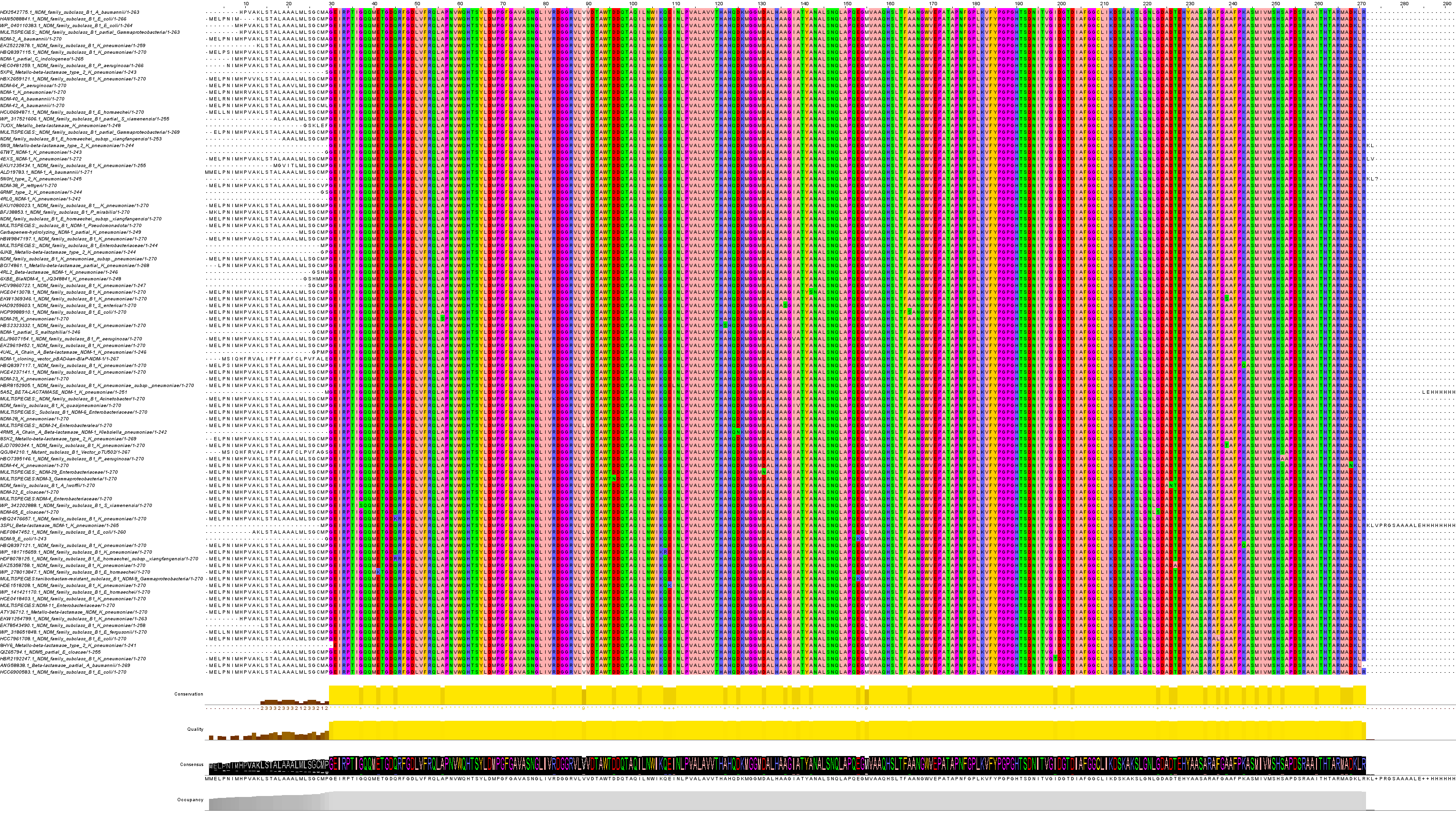

**Figure S2a**. Multiple sequence alignment of the NDM protein displayed with Jalview's color scheme, which highlights residues based on their chemical properties when conserved or similar across sequences. The figure also displays occupancy, consensus, quality, and sequence conservation levels.

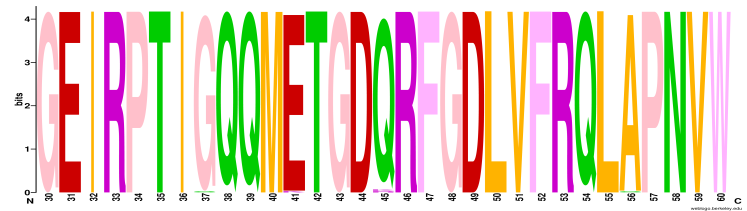

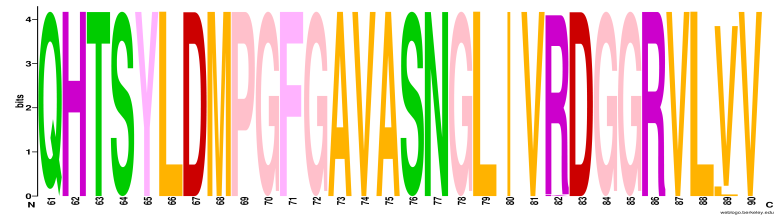

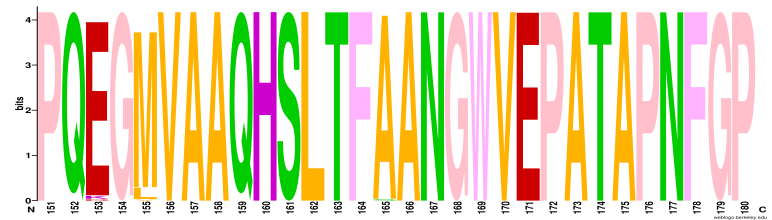

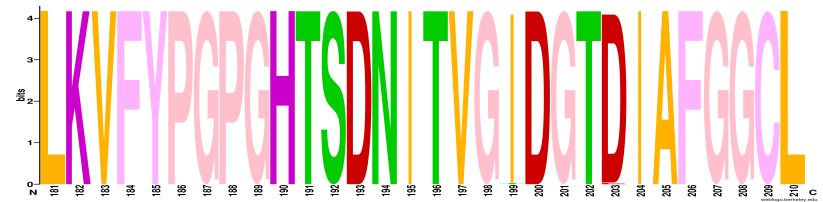

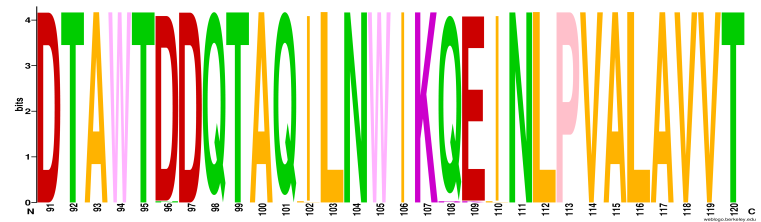

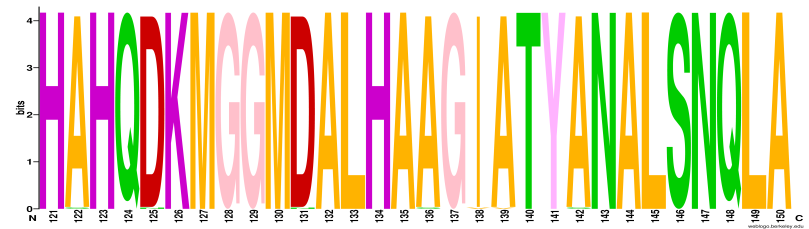

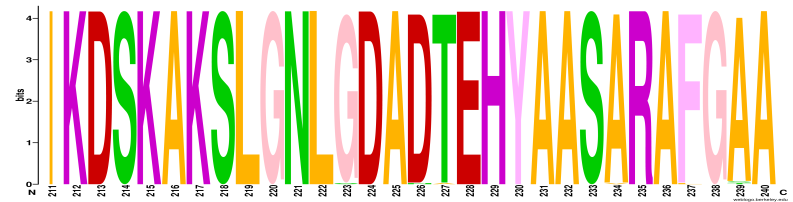

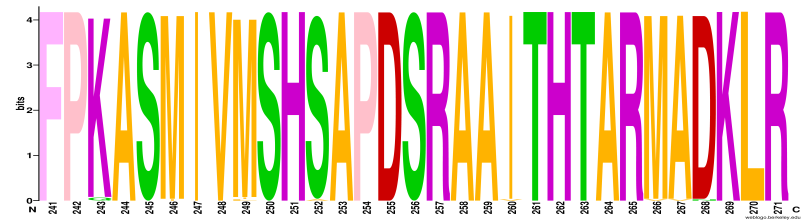

**Figure S2b.** Sequence logos illustrating residue conservation in NDM were generated for all aligned blocks of the MSA using the WebLogo web server (<https://weblogo.berkeley.edu/logo.cgi>). The x-axis represents the column numbers from the MSA.

**­­­­­Table S2:** List of mutations in NDM occurring in highly conserved columns (approximately 98% conserved across all species) identified through multiple sequence alignment (MSA).

| **Protein and Bacteria** | **Position of Mutation (Amino Acid)** | **Amino Acid Mutation** |
| --- | --- | --- |
| WP342202888.1 NDM family subclass B1 *Shewanella xiamenensis* | 37 Position | Glycine (non polar) Serine (polar) |
| WP278013847.1 NDM family subclass B1 *Enterobacter Hormaechei* | 41 Position | Glutamate (-ve charged) Lysine (+ve charged) |
| HBQ2476657.1 NDM family subclass B1 *Klebsiella pneumoniae* | 45 Position | Glutamine (uncharged) Lysine (+ve charged) |
| HDF8078125.1 NDM family subclass B1 *Enterobacter Hormaechei* subsp. *xiangfangensis* | 45 Position | Glutamine (uncharged) Arginine (+ve charged) |
| NDM-25 *Klebsiella pneumoniae* | 56 Position | Alanine (non polar) Serine (polar) |
| MULTISPECIES: NDM family subclass B1 *Acinetobacter* | 82 Position | Arginine (+ve charged) Lysine (+ve charged) |
| MULTISPECIES:NDM-3 *Gammaproteobacteria* | 96 Position | Glutamate (-ve charged) Asparagine (uncharged) |
| NDM-23 *Klebsiella pneumoniae* | 102 Position | Isoleucine (non polar) Leucine (polar) |
| WP181715659.1 NDM family subclass B1 *Klebsiella pneumoniae* | 108 Position | Glutamine (uncharged) Arginine (+ve charged) |
| HDE1518208.1 NDM family subclass B1 *Klebsiella pneumoniae* | 109 Position | Glutamate (-ve charged) Lysine (+ve charged) |
| HBR8152905.1 NDM family subclass B1 *Klebsiella pneumoniae* subsp. *pneumoniae* | 110 Position | Isoleucine (non polar) Leucine (polar) |
| HBS2323332.1 NDM family subclass B1 *Klebsiella pneumoniae* | 122 Position | Alanine (non polar) Serine (polar) |
| 4RM5 Beta-lactamase NDM-1 *Klebsiella_pneumoniae* | 125 Position | Aspartate (-ve charged) Asparagine (uncharged) |
| MULTISPECIES: NDM-29 *Enterobacteriaceae* | 131 Position | Aspartate (-ve charged) Asparagine (uncharged) |
| HAD9259603.1 NDM family subclass B1 *Salmonella enterica* | 136 Position | Alanine (non polar) Serine (polar) |
| ELJ9607154.1 NDM family subclass B1 *Pseudomonas aeruginosa* | 138 Position | Isoleucine (non polar) Leucine (polar) |
| NDM family subclass B1 *Klebsiella quasipneumoniae* | 139 Position | Alanine (non polar) Valine (non polar) |
| HCE0413078.1 NDM family subclass B1 *Klebsiella pneumoniae* | 142 Position | Alanine (non polar) Serine (polar) |
| HCP9988910.1 NDM family sub classB1 *Escherichia coli* | 165 Position | Alanine (non polar) Serine (polar) |
| HBR2192247.1 NDM_family subclass B1 *Klebsiella pneumoniae* | 199 Position | Isoleucine (non polar) Threonine (polar) |
| HCE4237141.1 NDM_family subclass B1 *Klebsiella pneumoniae* | 203 Position | Aspartate (-ve charged) Glutamate (-ve charged) |
| NDM-65 *Enterobacter cloacae* | 223 Position | Glycine (non polar) Serine (polar) |
| NDM family subclass B1 *Acinetobacter lwoffii* | 226 Position | Glutamate (-ve charged) Asparagine (uncharged) |
| EKZ5358758.1 NDM family subclass B1 *Klebsiella pneumoniae* | 227 Position | Threonine (polar) Alanine (non polar) |
| MULTISPECIES: Subclass B1 NDM-6 *Enterobacteriaceae* | 234 Position | Alanine (non polar) Valine (polar) |
| HCC7961708.1 NDM family subclass B1 *Escherichia coli* | 237 Position | Phenylalanine (aromatic) Leucine (aliphatic) |
| EKW1369346.1 NDM family subclass B1 *Klebsiella pneumoniae* | 239 Position | Alanine (non polar) Serine (polar) |
| EJD7090344.1 NDM family subclass B1 *Klebsiella pneumoniae* | 239 Position | Alanine (non polar) Threonine (polar) |
| HBQ8397117.1 NDM family subclass B1 *Klebsiella pneumoniae* | 243 Position | Glutamine (uncharged) Lysine (+ve charged) |
| HCE0418403.1 NDM family subclass B1 *Klebsiella pneumoniae* | 246 Position | Methionine (non polar) Isoleucine (non polar) |
| NDM-22 *Enterobacter cloacae* | 249 Position | Methionine (non polar) Leucine (polar) |
| WP141421170.1 NDM family subclass B1 *Enterobacter hormaechei* | 249 Position | Methionine (non polar) Leucine (polar) |
| HBO7395146.1 NDM family subclass B1 *Pseudomonas aeruginosa* | 252 Position | Serine (polar) Alanine (non polar) |
| EKZ9619452.1 NDM family subclass B1 *Klebsiella pneumoniae* | 260 Position | Isoleucine (non polar) Valine (polar) |
| ATY36712.1 Metallo-beta-lactamase NDM *Klebsiella pneumoniae* | 266 Position | Methionine (non polar) Valine (non polar) |
| NDM-28 *Klebsiella pneumoniae* | 267 Position | Alanine (non polar) Valine (non polar) |
| NDM-44 *Klebsiella pneumoniae* | 268 Position | Aspartate (-ve charged) Asparagine (uncharged) |

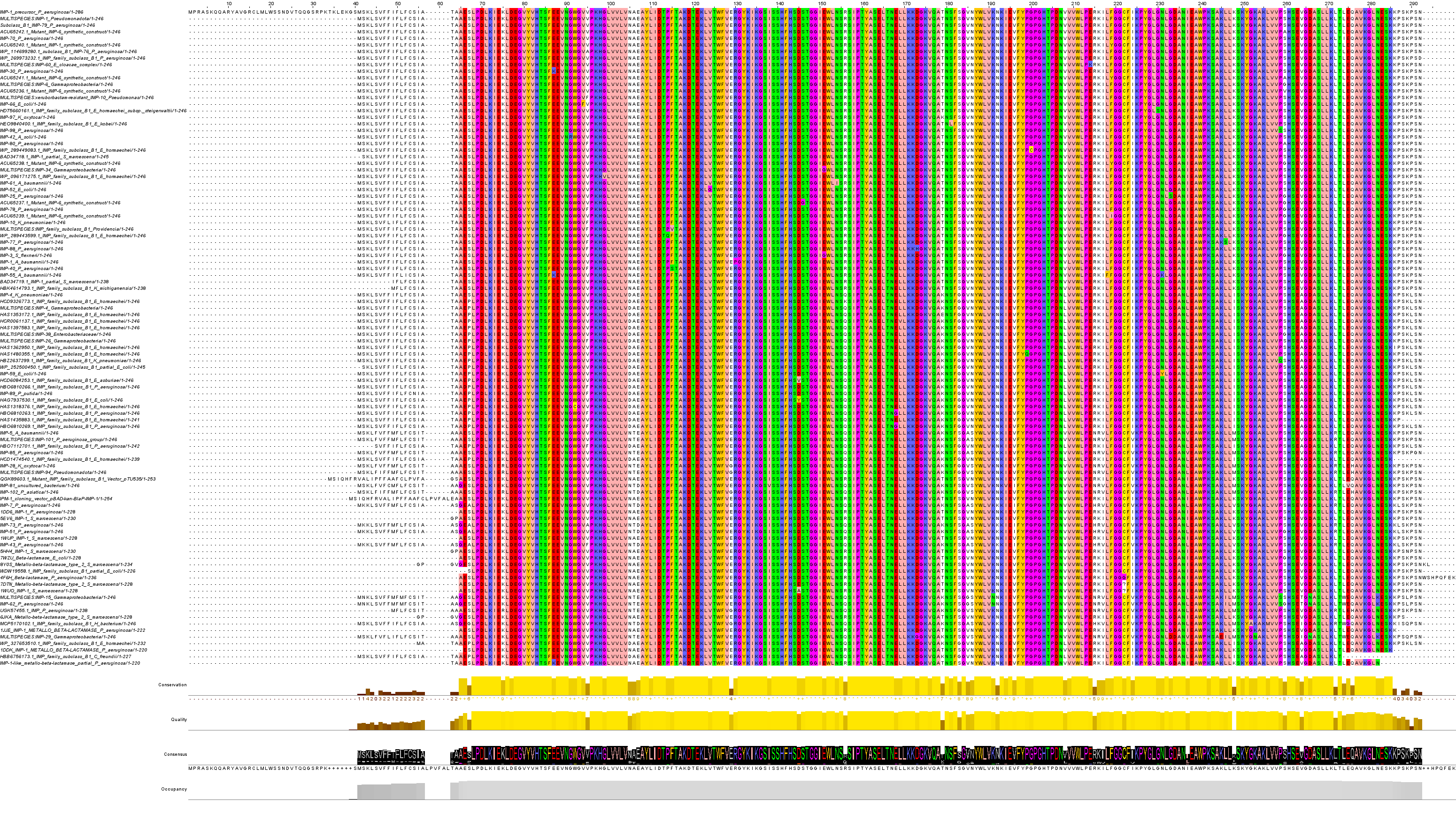
**Figure S3a**. Multiple sequence alignment of the IMP protein displayed with Jalview's color scheme, which highlights residues based on their chemical properties when conserved or similar across sequences. The figure also displays occupancy, consensus, quality, and sequence conservation levels.

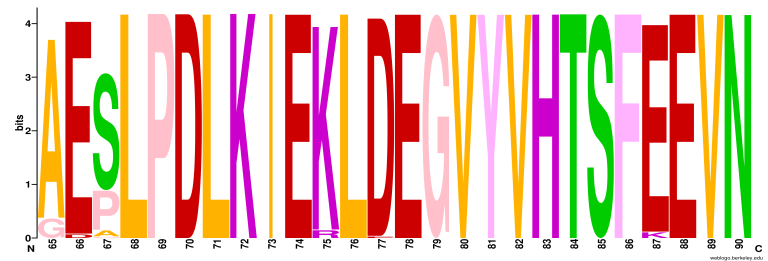

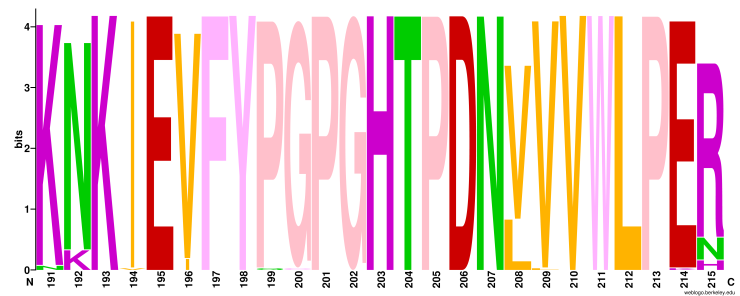

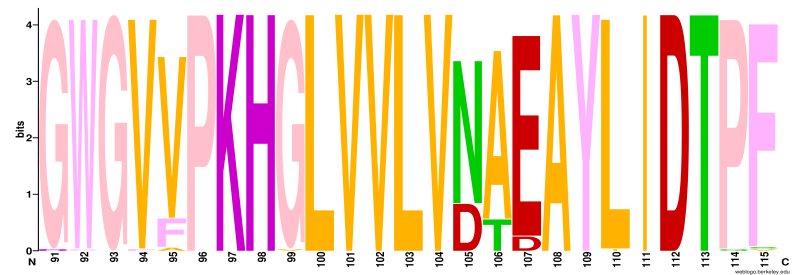

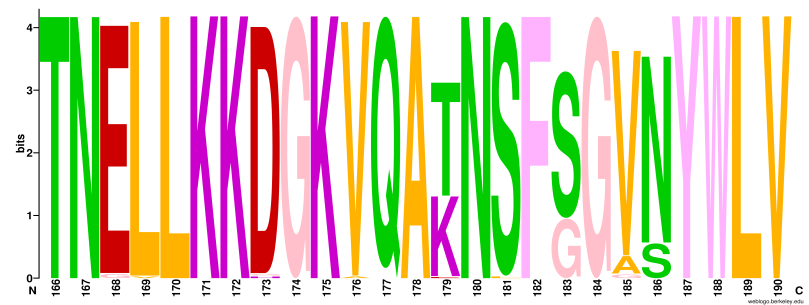

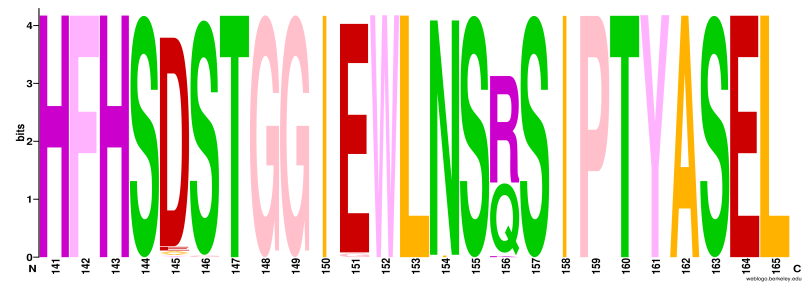

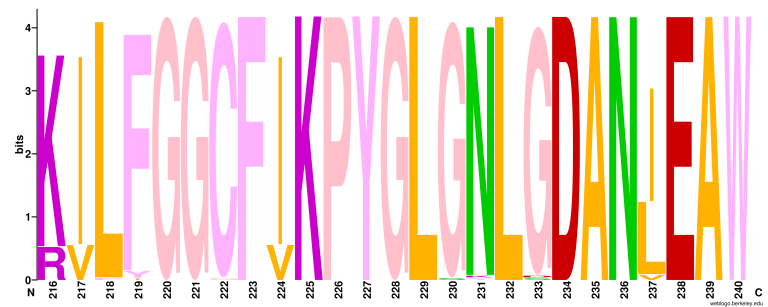

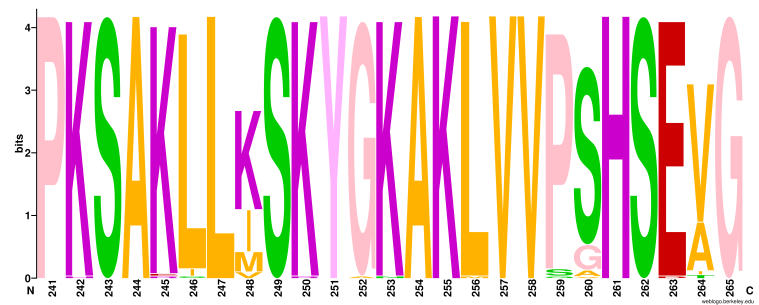

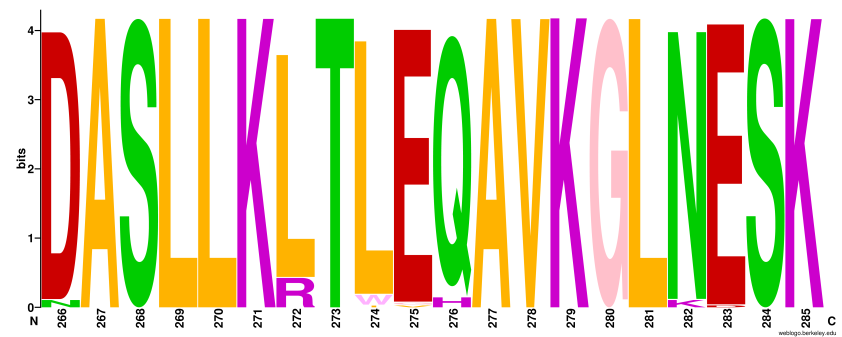

**Figure S3b.** Sequence logos illustrating residue conservation in IMP were generated for all aligned blocks of the MSA using the WebLogo web server (https://weblogo.berkeley.edu/logo.cgi). The x-axis represents the column numbers from the MSA.

**­­­­­**

**Table S3:** List of mutations in IMP occurring in highly conserved columns (approximately 98% conserved across all species) identified through multiple sequence alignment (MSA).

| **Protein and Bacteria** | **Position of Mutation (Amino Acid)** | **Amino Acid Mutation** |
| --- | --- | --- |
| IMP-55 *Acinetobacter baumannii* | 77 Position | Aspartate (-ve charged) Glutamate (-ve charged) |
| IMP-42 *Acinetobacter soli* | 91 Position | Glycine (non polar) Arginine (+ve charged) |
| HAS1318376.1 IMP family subclass B1 *Enterobacter Hormaechei* | 92 Position | Tryptophan (non polar, aromatic) Cysteine (polar, aliphatic) |
| IMP-66 *Escherichia coli* | 94 Position | Valine (aliphatic) Phenylalanine (aromatic) |
| WP 096171275.1 IMP family subclass B1 *Enterobacter Hormaechei* | 99 Position | Glycine (non polar) Valine (non polar) |
| IMP-52 *Escherichia coli* | 110 Position | Leucine (non polar) Isoleucine (non polar) |
| WP 289443599.1 IMP family subclass B1 *Enterobacter Hormaechei* | 114 Position | Proline (ring structure) Glutamine (normal chain structure) |
| MULTISPECIES:IMP family subclass B1 *Providencia* | 115 Position | Phenylalanine (aromatic) Valine (aliphatic) |
| IMP-40 *Pseudomonas aeruginosa* | 115 Position | Phenylalanine (aromatic) Serine (aliphatic) |
| IMP-52 *Escherichia coli* | 124 Position | Valine (non polar) Glycine (non polar) |
| IMP-1 *Acinetobacter baumannii* | 130 Position | Arginine (+ve charged) Proline (uncharged) |
| IMP-81 Uncultured bacterium | 138 Position | Isoleucine (non polar) Valine (polar) |
| ACU65237.1 Mutant IMP-6 Synthetic construct | 146 Position | Serine (polar) Glycine (non polar) |
| IMP-3 *Shigella flexneri* | 151 Position | Glutamate (-ve charged) Glycine (non polar) |
| MULTISPECIES:IMP-34 Gammaproteobacteria | 151 Position | Glutamate (-ve charged) Glycine (non polar) |
| IMP-61 *Acinetobacter baumannii* | 154 Position | Asparagine (polar) Isoleucine (non polar) |
| HBO6810263.1 IMP family subclass B1 *Pseudomonas aeruginosa* | 168 Position | Glutamate (-ve charged) Glycine (non polar) |
| HBO6810269.1 IMP family subclass B1 *Pseudomonas aeruginosa* | 168 Position | Glutamate (-ve charged) Glycine (non polar) |
| HCR0061137.1 IMP family subclass B1 *Enterobacter Hormaechei* | 169 Position | Leucine (non polar) Valine (non polar) |
| IMP-88 *Pseudomonas aeruginosa* | 173 Position | Aspartate (-ve charged) Histidine (+ve charged) |
| MULTISPECIES:IMP-29 Gammaproteobacteria | 173 Position | Aspartate (-ve charged) Glycine (non polar) |
| MCP5170102.1 IMP family subclass B1 *Helicobacter* bacterium | 176 Position | Valine (non polar) Alanine (non polar) |
| HEO9840400.1 IMP family subclass B1 *Enterobacter kobei* | 181 Position | Serine (polar) Leucine (non polar) |
| MULTISPECIES:IMP-15 Gammaproteobacteria | 191 Position | Lysine (charged) Asparagine (uncharged) |
| IMP-62 *Pseudomonas aeruginosa* | 191 Position | Lysine (charged) Asparagine (uncharged) |
| IMP-81 Uncultured bacterium | 194 Position | Isoleucine (non polar) Valine (non polar) |
| HAS1480355.1 IMP family subclass B1 *Enterobacter Hormaechei* | 199 Position | Proline (ring structure) Glutamine (normal chain structure) |
| WP 289449083.1 IMP family subclass B1 *Enterobacter Hormaechei* | 200 Position | Glycine (non polar) Cysteine (polar) |
| IMP-1 *Acinetobacter baumannii* | 209 Position | Valine (non polar) Leucine (non polar) |
| MULTISPECIES:IMP-60 *Enterobacter cloacae* complex | 214 Position | Glutamate (-ve charged) Lysine (+ve charged) |
| IMP-55 *Acinetobacter baumannii* | 218 Position | Leucine (aliphatic) Phenylalanine (aromatic) |
| HDT5660161.1 IMP family subclass B1 *Enterobacter Hormaechei* subsp. *steigerwaltii* | 230 Position | Glycine (non polar) Serine (polar) |
| IMP-55 *Acinetobacter baumannii* | 231 Position | Asparagine (uncharged) Lysine (+ve charged) |
| IMP-59 *Escherichia coli* | 231 Position | Asparagine (polar, aliphatic) Tyrosine (non polar, aromatic) |
| IMP-25 *Pseudomonas aeruginosa* | 233 Position | Glycine (non polar) Serine (polar) |
| MULTISPECIES:IMP-29 Gammaproteobacteria | 233 Position | Glycine (non polar) Aspartate (-ve charged) |
| MULTISPECIES:IMP-29 Gammaproteobacteria | 242 Position | Lysine (+ve charged) Histidine (+ve charged) |
| Subclass B1 IMP-79 *Pseudomonas aeruginosa* | 245 Position | Lysine (+ve charged) Arginine (+ve charged) |
| MULTISPECIES:IMP-29 Gammaproteobacteria | 245 Position | Lysine (+ve charged) Glutamate (-ve charged) |
| MULTISPECIES:IMP-29 Gammaproteobacteria | 250 Position | Lysine (+ve charged) Arginine (+ve charged) |
| MCP5170102.1 IMP family subclass B1 *Helicobacter* bacterium | 252 Position | Glycine (non polar) Alanine (non polar) |
| MULTISPECIES:IMP-29 Gammaproteobacteria | 253 Position | Lysine (+ve charged) Asparagine (uncharged) |
| MCP5170102.1 IMP family subclass B1 *Helicobacter* bacterium | 256 Position | Leucine (non polar) Methionine (non polar) |
| MULTISPECIES:IMP-29 Gammaproteobacteria | 263 Position | Glutamate (-ve charged) Aspartate (-ve charged) |

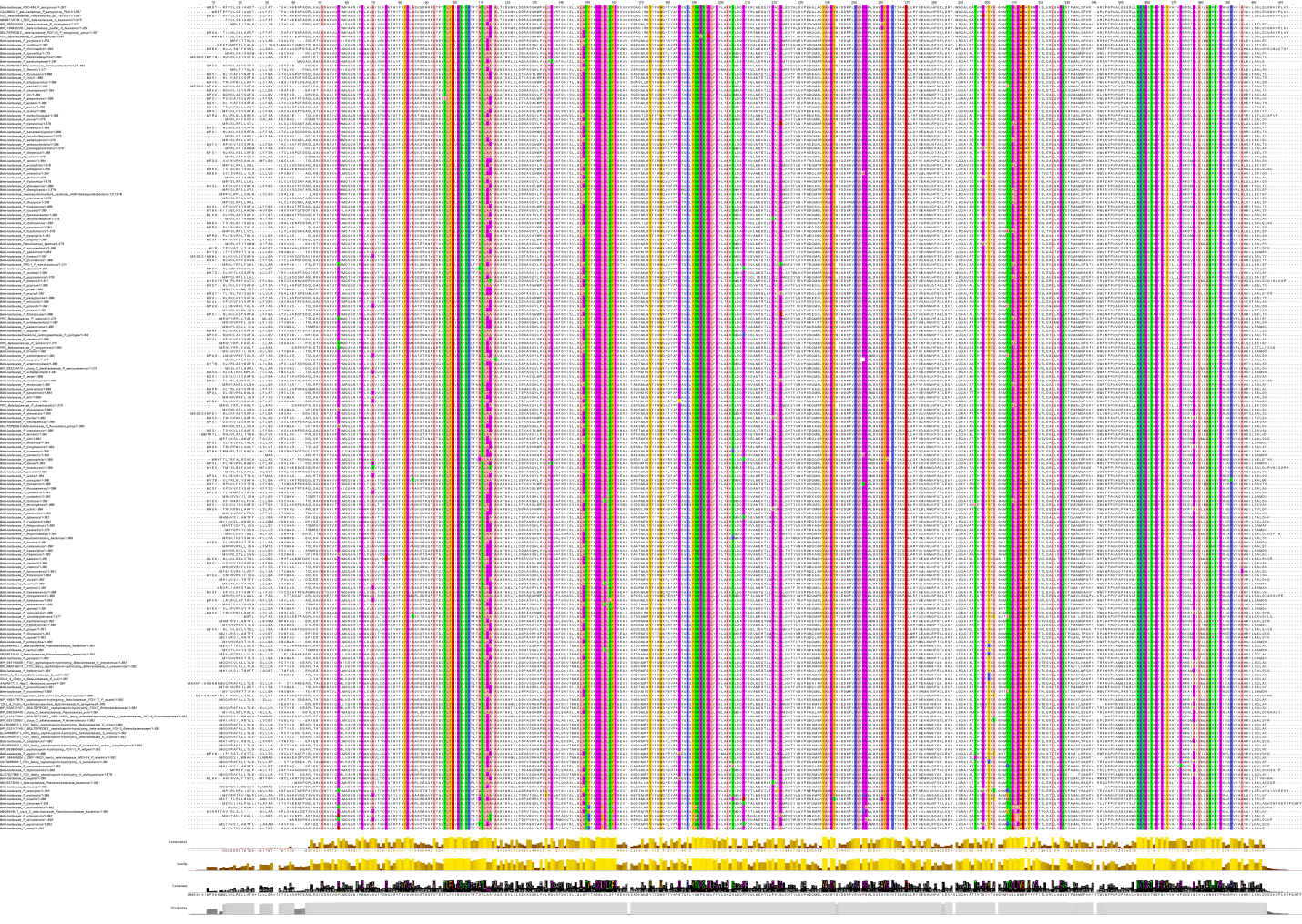
**Figure S4a**. Multiple sequence alignment of the AmpC protein displayed with Jalview's color scheme, which highlights residues based on their chemical properties when conserved or similar across sequences. The figure also displays occupancy, consensus, quality, and sequence conservation levels.

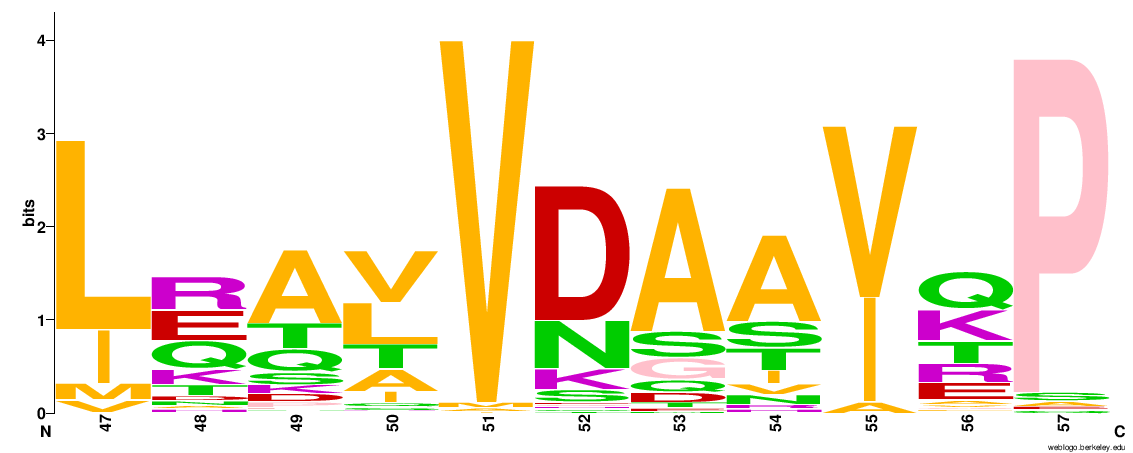

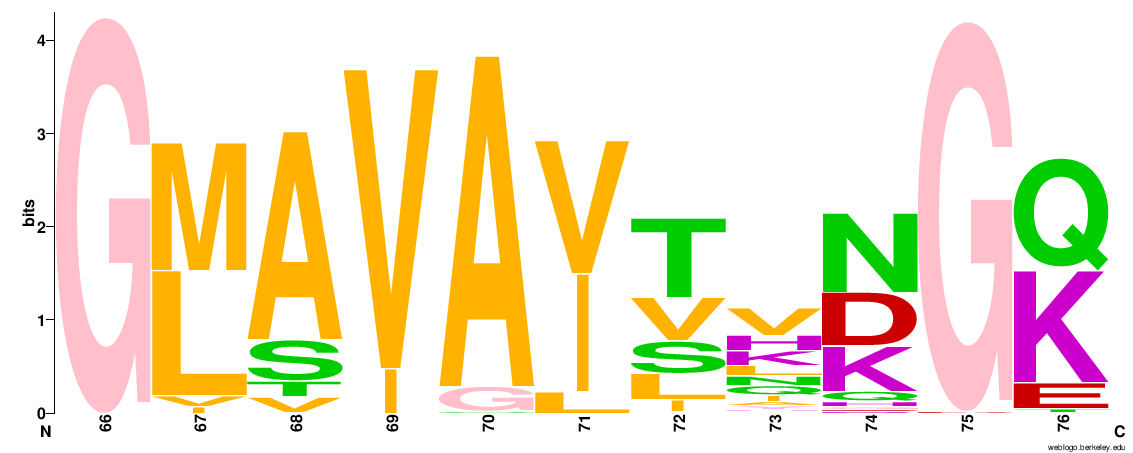

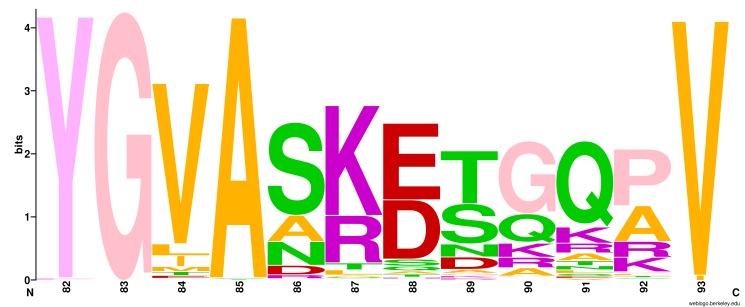

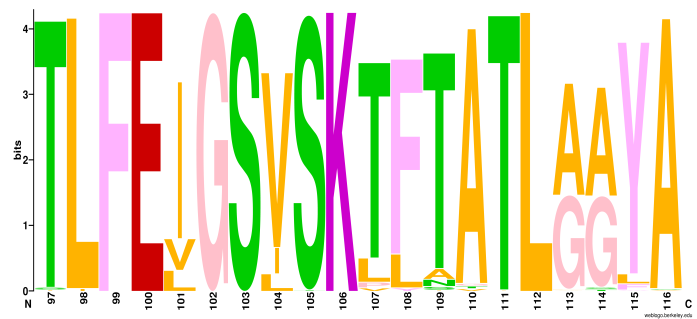

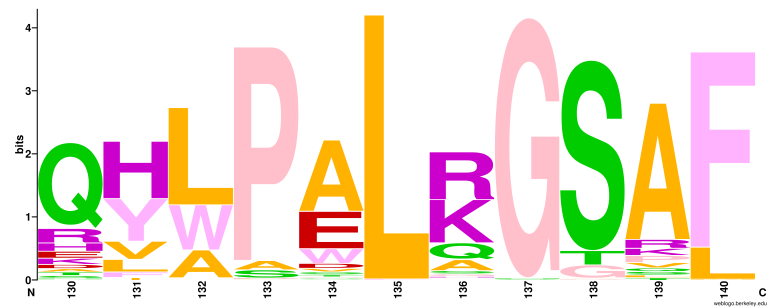

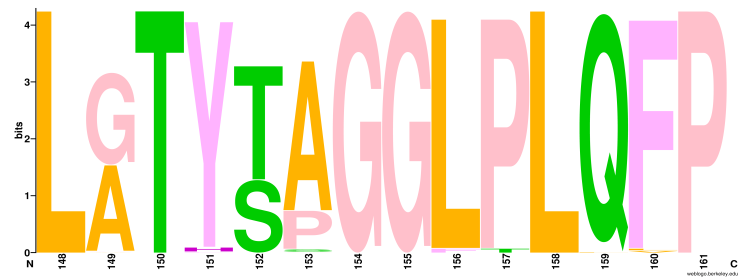

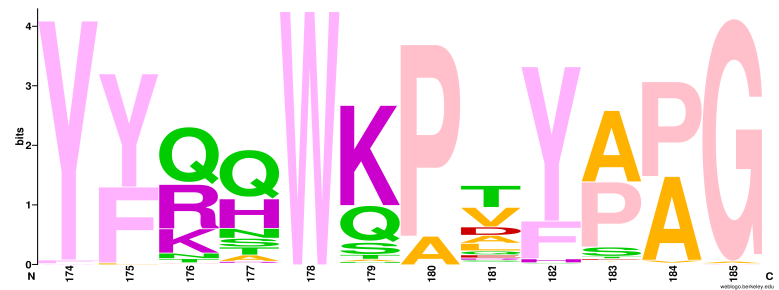

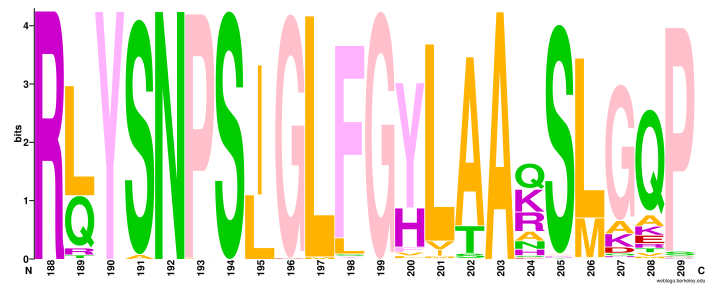

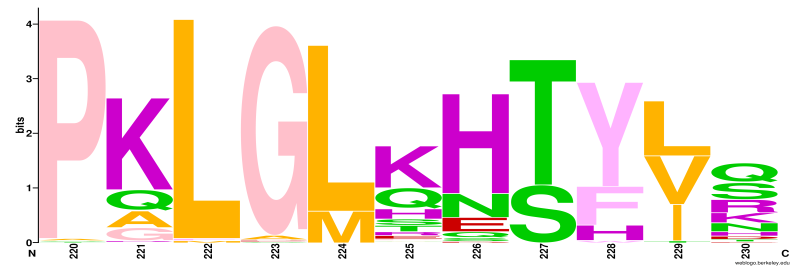

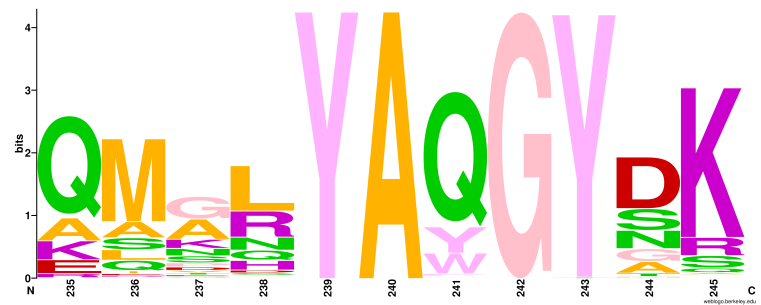

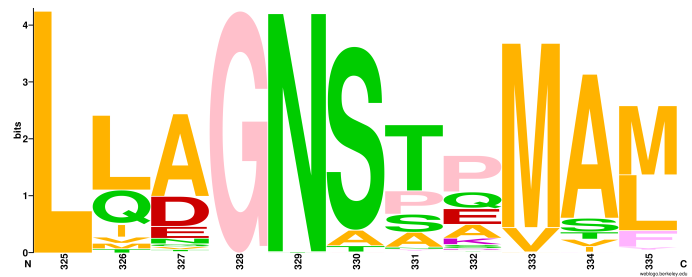

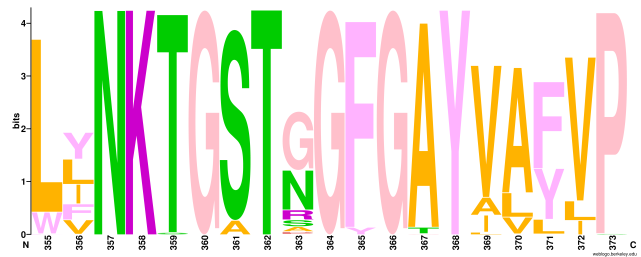

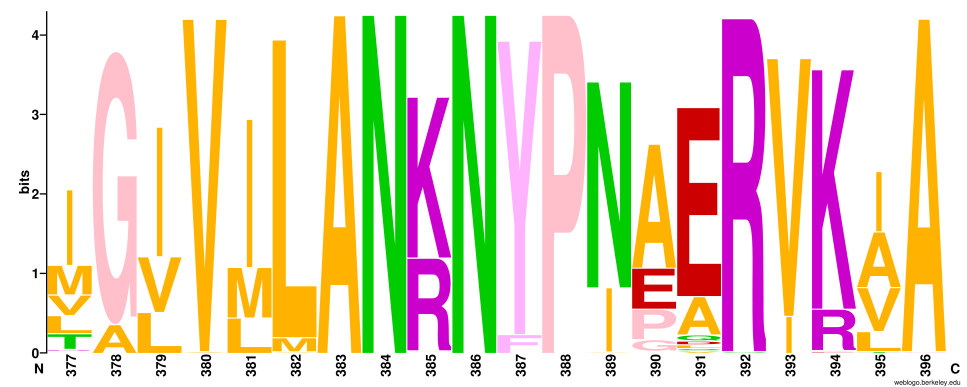

**Figure S4b.** Sequence logos illustrating residue conservation in AmpC were generated for all aligned blocks of the MSA using the WebLogo web server (<https://weblogo.berkeley.edu/logo.cgi>). The x-axis represents the column numbers from the MSA.

**­­­­­Table S4:** List of mutations in AmpC occurring in highly conserved columns (approximately 98% conserved across all species) identified through multiple sequence alignment (MSA).

| **Protein and Bacteria** | **Position of Mutation (Amino Acid)** | **Amino Acid Mutation** |
| --- | --- | --- |
| Beta-lactamase *Pseudomonas batumici* | 75 Position | Glycine (non polar) Aspartate (-ve charged) |
| Beta-lactamase *Pseudomonas resinovorans* | 82 Position | Tyrosine (aromatic) Phenylalanine (aromatic) |
| Beta-lactamase *Pseudomonas resinovorans* | 82 Position | Tyrosine (aromatic) Phenylalanine (aromatic) |
| Beta-lactamase *Pseudomonas* *corrugata* | 85 Position | Alanine (non polar) Threonine (polar) |
| Beta-lactamase *Pseudomonas* *vanderleydeniana* | 85 Position | Alanine (non polar) Serine (polar) |
| Beta-lactamase *Pseudomonas* *orientalis* | 105 Position | Serine (polar) Threonine (polar) |
| Beta-lactamase *Pseudomonas* *saponiphila* | 111 Position | Threonine (polar) Isoleucine (non polar) |
| Beta-lactamase *Pseudomonas* *marincola* | 135 Position | Leucine (aliphatic) Phenylalanine (aromatic) |
| Beta-lactamase *Pseudomonas* *paralcaligenes* | 137 Position | Glycine (non polar) Serine (polar) |
| Beta-lactamase *Pseudomonas* *simiae* | 137 Position | Glycine (non polar) Asparagine (polar) |
| PDC beta-lactamase *Pseudomonas* *paraeruginosa* | 193 Position | Proline (ring structure) Leucine (simple chain structure) |
| Beta-lactamase *Pseudomonas* *ullengensis* | 197 Position | Leucine (non polar) Methionine (non polar) |
| Beta-lactamase *Pseudomonas* *campi* | 197 Position | Leucine (non polar) Methionine (non polar) |
| Beta-lactamase *Pseudomonas* *dryadis* | 243 Position | Tyrosine (aromatic) Threonine (aliphatic) |
| Beta-lactamase *Pseudomonas* *resinovorans* | 255 Position | Glycine (non polar) Alanine (non polar) |
| Beta-lactamase *Pseudomonas* *taiwanensis* | 255 Position | Glycine (non polar) Alanine (non polar) |
| Beta-lactamase *Pseudomonas* *saponiphila* | 263 Position | Glycine (non polar) Alanine (non polar) |
| Penicillin-binding protein beta-lactamase *Pseudomonas* *fuscovaginae* | 329 Position | Asparagine (polar) Threonine (polar) |
| Beta-lactamase *Pseudomonas* *triticicola* | 359 Position | Threonine (polar) Serine (polar) |
| Beta-lactamase *Pseudomonas* *iranensis* | 359 Position | Threonine (polar) Serine (polar) |
| Beta-lactamase *Pseudomonas* *canavaninivorans* | 373 Position | Proline (ring structure) Serine (simple chain structure) |
| Beta-lactamase *Pseudomonas* *granadensis* | 380 Position | Valine (non polar) Alanine (non polar) |
| Beta-lactamase *Pseudomonas* *corrugata* | 392 Position | Arginine (+ve charged) Glutamine (uncharged) |
